# CellFishing.jl: an ultrafast and scalable cell search method for single-cell RNA-sequencing

**DOI:** 10.1101/374462

**Authors:** Kenta Sato, Koki Tsuyuzaki, Kentaro Shimizu, Itoshi Nikaido

## Abstract

Recent technical improvements in single-cell RNA sequencing (scRNA-seq) have enabled massively parallel profiling of transcriptomes, thereby promoting large-scale studies encompassing a wide range of cell types of multicellular organisms. With this background, we propose CellFishing.jl, a new method for searching atlas-scale datasets for similar cells and detecting noteworthy genes of query cells with high accuracy and throughput. Using multiple scRNA-seq datasets, we validate that our method demonstrates comparable accuracy to and is markedly faster than the state-of-the-art software. Moreover, CellFishing.jl is scalable to more than one million cells, and the throughput of the search is approximately 1,600 cells per second.

## Background

The development of high-throughput single-cell RNA sequencing (scRNA-seq) technology for the past several years has enabled massively parallel profiling of transcriptome expressions at the single-cell level. In contrast to traditional RNA sequencing methods that profile the average of bulk samples, scRNA-seq has the potential to reveal heterogeneity within phenotypes of individual cells as it can distinguish the transcriptome expression of each cell by attaching a distinct cellular barcode [1, 2]. In addition, several protocols have been developed that utilize unique molecular identifiers (UMIs) to more accurately quantify expression by removing duplicated counts resulting from the amplification of molecules [3–8]. The advent of library preparation for multiplexed sequencing with cellular barcoding and the refinement of cDNA amplification method with UMIs lead to a higher throughput and more reliable quantification of single-cell expression profiles.

These technologies have opened the door to research that comprehensively sequences and annotates massive numbers of cells to create a cell atlas for organs or multicellular organisms. Shekhar *et al*. [9] sequenced and performed unsupervised classification of 25,000 mouse retinal bipolar cells and identified novel cell types and marker genes, suggesting that sequencing a large number of cells is an essential factor for detecting under-represented cell types. Similarly, Plass *et al*. [10] sequenced more than 20,000 planarian cells and rendered a single lineage tree representing continuous differentiation. We also see collaborative efforts to create a comprehensive catalog covering all cells types composing an organism, such as the Human Cell Atlas [11] and the Tabula Muris [12] project. This trend of sequencing higher numbers of cells is expected to continue until a complete list of cell types is generated.

Emergence of these comprehensive single-cell sequencing studies shows a pressing demand for software to find similar cells by comparing their transcriptome expression patterns. Since discrete cell annotations are not always available or are even impossible to generate due to continuous cell state dynamics, software for cell-level searching is useful for comparative analysis. However, finding similar cells based on their transcriptome expression profiles is computationally challenging due to the unprecedented numbers of genes and cells. Recently, Kiselev *et al*. [13] developed a software package and web service named scmap to perform an approximate nearest neighbor search of cells using a product quantizer [14]. The scmap package contains two variations: scmap-cluster and scmap-cell. Scmap-cluster can be used to search for cell clusters that are defined by discrete cluster labels and hence requires cluster annotations in addition to expression profiles of reference cells. On the contrary, scmap-cell can be used to directly find similar cells only from their expression profiles and is applicable to scRNA-seq data without requiring cluster annotations for cells. The authors of scmap-cell claim that creating a search index is more rapid than employing machine-learning methods.

However, the scalability of scmap-cell is limited and is not applicable to extremely large data sets. Srivastava *et al*. [15] have also developed a web service named CellAtlasSearch that searches existing scRNA-seq experiments using locality-sensitive hashing (LSH) and graphical processing units (GPUs) to accelerate the search. In LSH, expression profiles are hashed into bit vectors, and their similarities are estimated from the Hamming distance between bit vectors calculated by LSH [16]. However, it requires GPUs to extract maximum performance, and its implementation details are neither openly accessible nor well-described in their paper.

We are also interested in determining cell state estimation. Although cell type estimation accomplished by matching query cells with similar cells found in annotated data sets provides important information concerning the query cells, relying on a single similarity score may result in overlooking significant differences in their gene expressions. For example, the developmental stages of the hematopoietic lineage from stem cells to completely differentiated cells are often characterized by the expression level of few marker genes. Additionally, using scRNA-seq Park *et al*. [17] revealed that genes related to Mendelian disease are differentially expressed in specific cell types. These facts indicate that mutually similar cells of the same type but under different conditions can be further distinguished by noting differentially expressed genes (DEGs) between these cells.

In this paper, we present CellFishing.jl (cell finder via hashing), a novel software package used to find similar cells from a prebuilt database based on their expression patterns with high accuracy and throughput. CellFishing.jl employs LSH, like CellAtlasSearch, to reduce the computational time and space required for searching; however, it does not require dedicated accelerators, and its implementation is freely available as an open-source software package written in the Julia programming language [18]. It also utilizes an indexing technique of bit vectors to rapidly narrow down candidates of similar cells. Moreover, a query cell can be compared with its neighboring cells in the database in order to prioritize noteworthy genes that are differentially expressed between the query and its neighbors, facilitating quick DEG analysis with single-cell resolution. Cell databases once created can be saved to a disk and quickly loaded for later searches. Here, we demonstrate the effectiveness and scalability of our approach using real scRNA-seq data sets, one of which includes more than one million cells.

## Results

### Workflow overview of CellFishing.jl

CellFishing.jl first creates a search database of reference cells from a matrix of transcriptome expression profiles of scRNA-seq experiments, and then searches the database for cells with an expression pattern similar to the query cells. The schematic workflow of CellFishing.jl is illustrated in Figure 1. When building a database, CellFishing.jl uses a digital gene expression (DGE) matrix as an input along with some metadata, if provided. It next applies preprocessing to the matrix, resulting in a reduced matrix, and subsequently hashes the column vectors of this reduced matrix to low-dimensional bit vectors. The preprocessing phase consists of five steps: feature (gene) selection, cell-wise normalization, variance stabilization, feature standardization, and dimensionality reduction. The information provided by these steps is stored in the database, and the same five steps are applied to the DGE matrix of query cells. In the hashing phase, random hyperplanes are generated from a pseudo-random number generator, and the column vectors of the reduced matrix are encoded into bit vectors according to the compartment in which the vector exists. This technique, termed LSH, is used to estimate the similarity between two data points by their hashed representation [16]. The bit vectors are indexed using multiple associative arrays that can be utilized for subsequent searches [19]. The implementation is written in the Julia language; the database object can be saved easily as a file, and the file can be transferred to other computers, which facilitates quick, comparative analyses across different scRNA-seq experiments.

**Figure 1:**
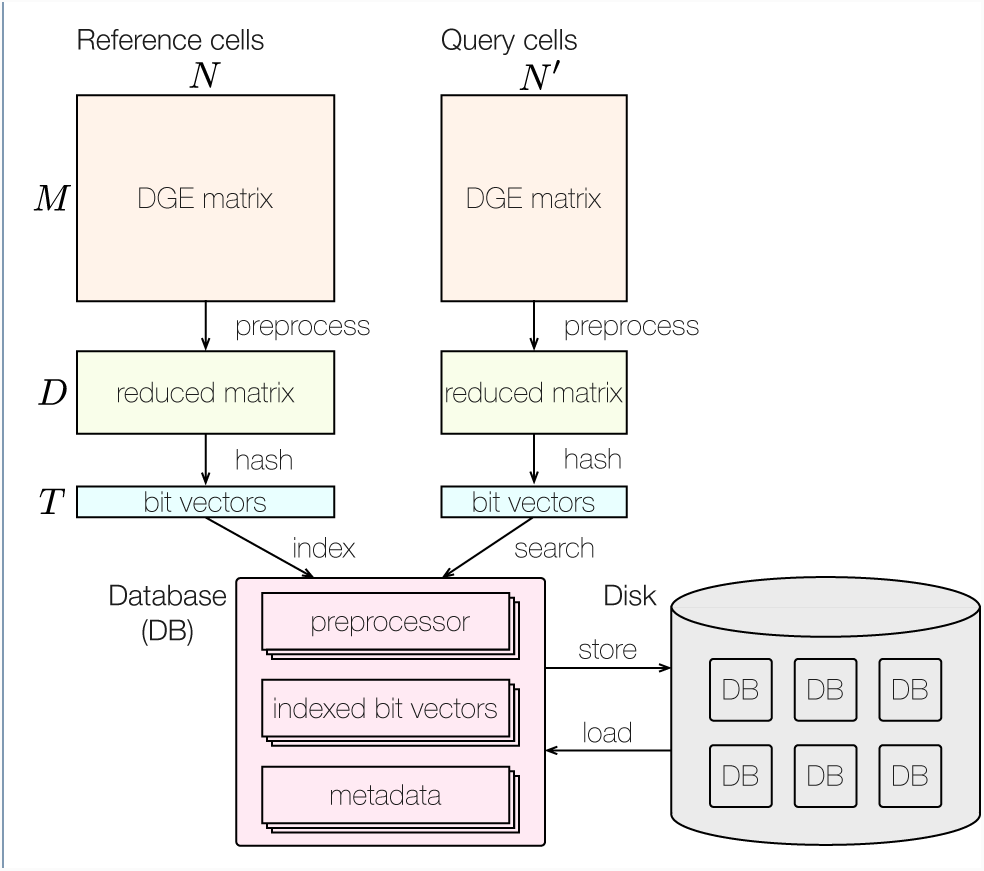
Schematic workflow of CellFishing.jl. CellFishing.jl first builds a database (DB) object that stores data preprocessors, indexed bit vectors, and cell metadata, if provided. The metadata can store any information including cell names, cell types, and transcript expressions of marker genes. When building a database, the DGE matrix of reference cells is preprocessed to extract important signals and then hashed into bit vectors by LSH. The preprocessors and the indexed bit vectors are stored in the database object. *M*, *D*, and *T* on the left side of the figure refer to the number of genes, number of reduced dimensions, and length of the bit vectors, respectively. *N* and *N*’ above the two DGE matrices represent the number of cells within the reference and query data, respectively. While searching the database for similar cells, the prebuilt preprocessors stored in the database are reused in a similar workflow that is involved in database building up to the hashing phase. The database object can be saved onto a disk and can be loaded from there.

### Data sets

We selected five data sets as benchmarks from scRNA-seq experiments, each including at least 10,000 cells, and one including more than 1.3 million cells, which was the largest publicly available data set. Cells without a cell type or cluster annotation were filtered out for evaluation. The data sets after filtering are summarized in Table 1.

**Table 1:** Summary of scRNA-seq data sets

| Data set | #Cells | #Clusters | Protocol | Cell types |
| --- | --- | --- | --- | --- |
| Baron2016 [22] | 10,455 | 14 | inDrop | pancreas cells of human and mouse |
| Shekhar2016 [9] | 27,499 | 19 | Drop-seq | retinal bipolar neurons of mouse |
| Plass2018 [10] | 21,612 | 51 | Drop-seq | mature and progenitor cells of planarian |
| TabulaMuris<sup>†</sup>[12] | 54,967 | 57 | Chromium | cell atlas of mouse |
| 1M_neurons [23] | 1,306,127 | 60 | Chromium | brain cells of mouse |

<sup>†</sup> Not including cells sequenced with Smart-Seq2.

Wagner *et al*. [20] recently reported that if there is no biological variation, excessive zero counts within a DGE matrix (dropouts) have not been observed in data generated from inDrop [5], Drop-seq [6], and Chromium [7] protocols. Similarly, Chen *et al*. [21] conducted a more thorough investigation and concluded that negative binomial models are preferred over zero-inflated negative binomial models for modeling scRNA-seq data with UMIs. We confirmed a similar observation using our control data generated from Quartz-Seq2 [8]. Therefore, we did not take into account the effects of dropout events in this study.

### Randomized singular value decomposition (SVD)

SVD is commonly used in scRNA-seq to enhance the signal-to-noise ratio by reducing the dimensions of the transcriptome expression matrix. However, computing the full SVD of an expression matrix or eigendecomposition of its covariance matrix is time consuming and requires large memory space especially when the matrix contains a large number of cells. Since researchers are usually interested in only a few dozen of the top singular vectors, it is common practice to compute only those important singular vectors. This technique is called low-rank matrix approximation, or truncated SVD. Recently, Halko *et al*. [24] developed approximated low-rank decomposition using randomization and were able to demonstrate its superior performance compared with other low-rank approximation methods. To determine the effectiveness of the randomized SVD, in this study, we benchmarked the performance of three SVD algorithms (full, truncated, and randomized) for real scRNA-seq data sets and evaluated the relative errors of singular values calculated using the randomized SVD. Full SVD is implemented using the svd function of Julia and the truncated SVD is implemented using the svds function of the Arpack.jl package, which computes the decomposition of a matrix using implicitly restarted Lanczos iterations; the same algorithm is used in Seurat [25] and CellRanger [7]. We implemented the randomized SVD as described in [26] and included the implementation in the CellFishing.jl package. We then computed the top 50 singular values and the corresponding singular vectors for the first four data sets listed in Table 1 and measured the elapsed time. All mouse cells (1,886 total) of the Baron2016 data set were excluded because merging expression profiles of human and mouse is neither trivial nor our focus here. The data sizes of the four data sets after feature selection were 2,190 × 8,569, 3,270 × 27,499, 3,099 × 21,612, and 2,363 × 54,967 in this order. From the benchmarks, we found that the randomized SVD remarkably accelerates the computation of low-rank approximation for scRNA-seq data without introducing large errors in the components corresponding to the largest singular values (Figure 2). It must be noted that in our application, obtaining exact singular vectors is not particularly important; rather computing the subspace with high variability spanned by approximated singular vectors is more important because each data point is eventually projected onto random hyperplanes during hashing. Therefore, evaluating relative errors of singular values suffices to quantify the precision of randomized SVD.

**Figure 2:**
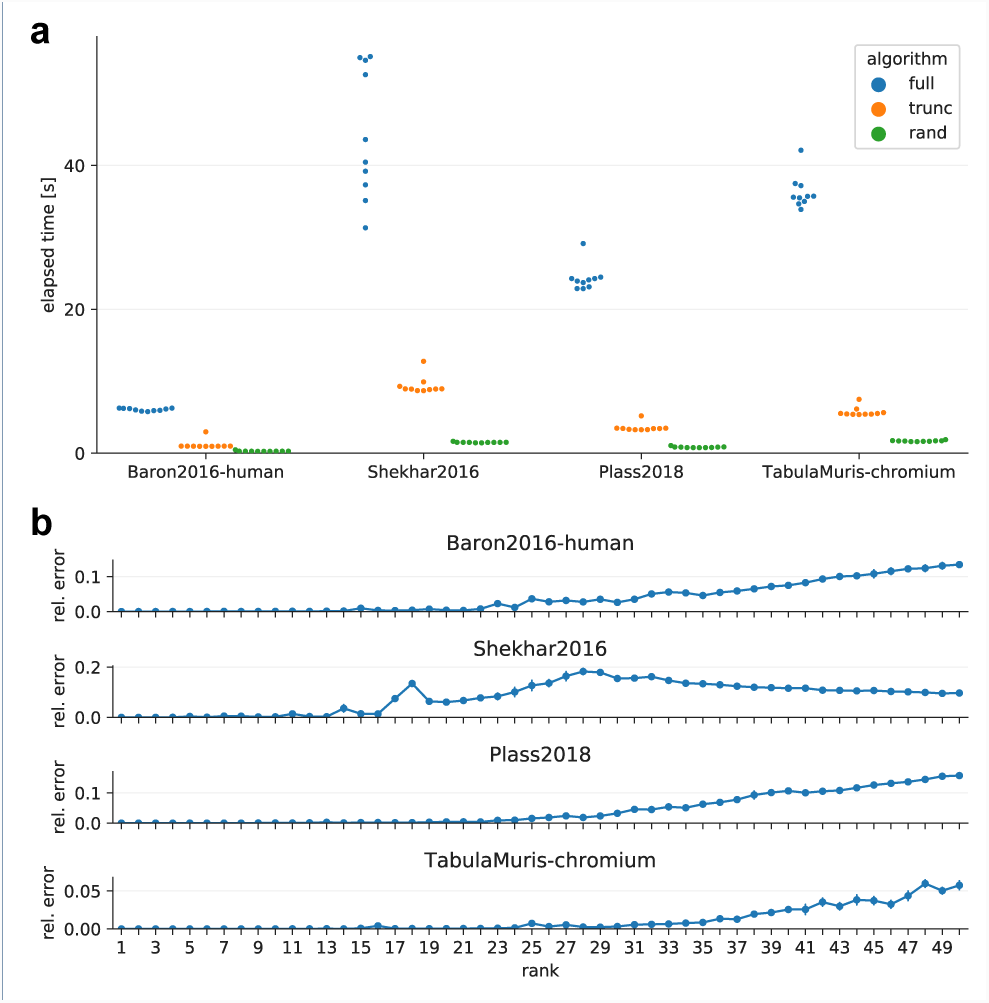
Benchmarks of randomized SVD. **a** Elapsed time of different SVD algorithms. The *blue, orange*, and *green points* indicate the elapsed time of the full, truncated, and randomized SVD, respectively. **b** Relative errors of the randomized SVD. The *error bars* denote the standard deviation of ten trials. The relative error of the *i*-th largest singular value *σ*_*i*_ is defined as 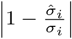, where 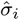 is an approximated value of *σ*_*i*_. The error bars denote the standard deviation of ten trials. The approximation error for a real matrix *A* with a low-rank matrix is bounded by a singular value as illustrated in the following formula: min_rank(*X*)*≤j*_ ‖*A − X*‖ = *σ*_*j*+1_, where ‖ · ‖ denotes the operator norm of a matrix.

### Similarity estimation using bit vectors

In LSH, the angular distance between two expression profiles can be estimated from the Hamming distance between their hashed bit vectors. Assuming *θ* is the angle between two numerical vectors representing the expression profiles, the estimator of *θ* derived from two bit vectors *p* and *q* is 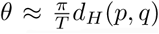 where *T* is the length of a bit vector and *d*_*H*_ (.,.)is a function that calculates the Hamming distance between the two bit vectors. This estimator is unbiased but occasionally suffers from its high variance, which can pose a problem. To counter this issue, CellFishing.jl employs an orthogonalization technique that creates more informative hyperplanes from random hyperplanes before hashing data points by orthogonalizing the normal vectors of these random hyperplanes [27]. We confirmed the variance reduction effect of this orthogonalization technique by comparing the estimators with the exact values using randomly sampled expression profiles, as shown in Figure 3a. The effect was consistent for all other data sets as expected (Additional file 1, Figure 12–14).

**Figure 3:**
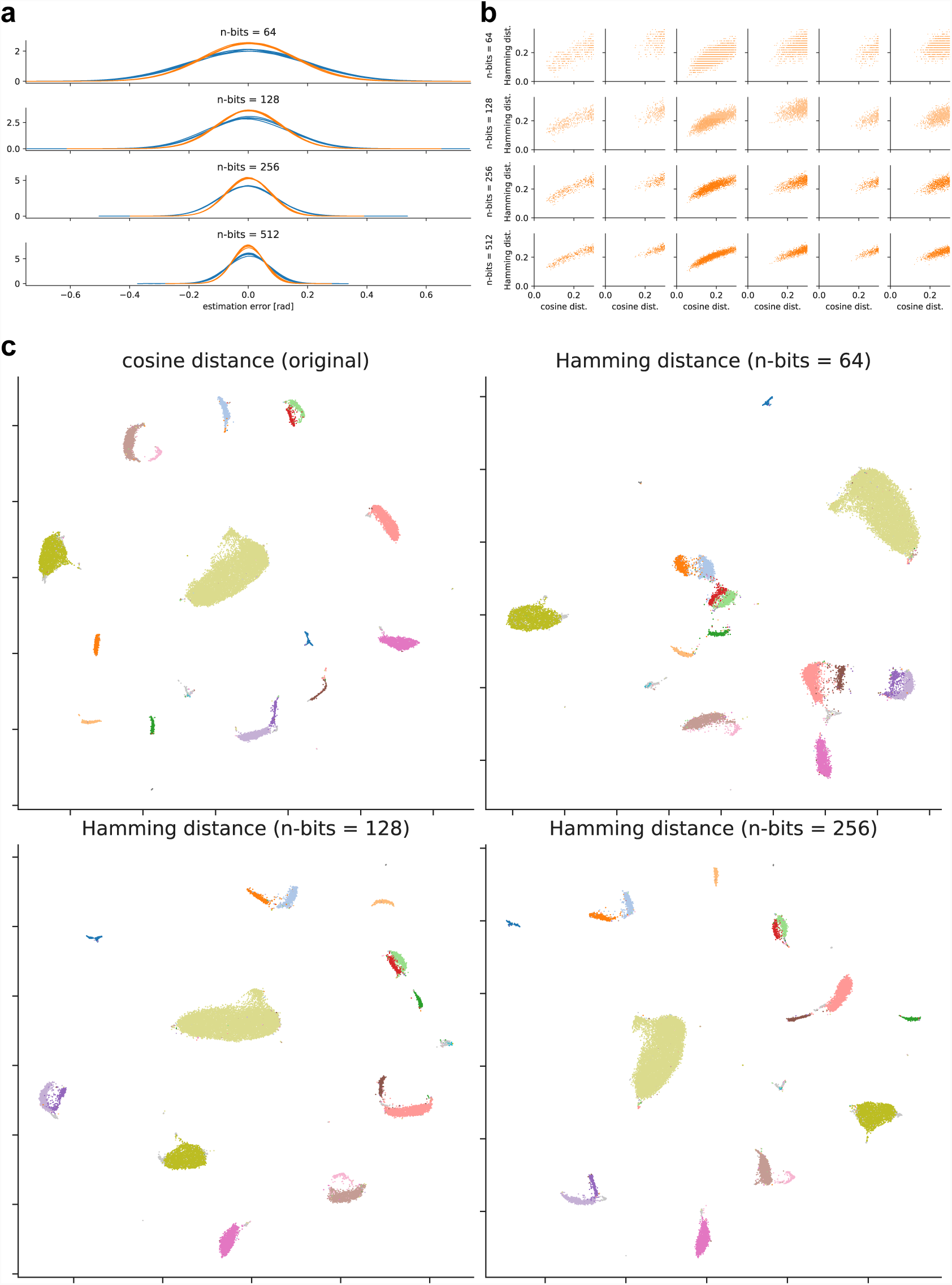
Locality-sensitive hashing of expression profiles (Shekhar2016). **a** Distributions of estimation errors for angles. The *blue lines* show the distributions without orthogonalization, and the *orange lines* show the distributions with orthogonalization. Five hash values were generated independently, and their estimation errors in radian were computed for 100 cells randomly sampled from Shekhar2016. **b** Scatter plots of the Hamming distance versus the cosine distance. The rows are four different bit lengths (64, 128, 256, and 512 bits) and the columns are six cells randomly sampled from Shekhar2016. The Hamming distance is normalized to [0, 1] for comparison across different bit lengths, and the cosine distance is truncated at 0.3 for brevity. **c** Two-dimensional embedding of expression profiles with UMAP. The upper-left plot was derived from the cosine distances following dimensionality reduction. The other three plots were derived from the Hamming distances after hashing with 64, 128, and 256 bits. Colors indicate the annotations (18 cell types and doublets/contaminants) of Shekhar2016.

As can be seen in Figure 3a, the estimator of the angular distance becomes less variable as the length of the bit vectors increases due to the central limit theorem. However, using more extended bit vectors requires more computational time and space. To investigate reasonable candidates for the length of bit vectors, we compared the Hamming and cosine distances of 100 random cells (Figure 3b). The Shekhar2016 data set shows that only 28% of random cells could find their true nearest neighbor in the top ten candidates nominated by 64-bit vectors, while 42%, 72%, and 85% could find their true neighbor by 128-, 256-, and 512-bit vectors, respectively. This result suggests that hashing expression profiles with 64-bit vectors is insufficient to find neighboring cells.

We next confirmed that the hashed expression profiles preserve the original differences among cell types by visualizing low-dimensional embedding of the data. Here we used Uniform Manifold Approximation and Projection (UMAP) [28], because it more explicitly preserves the global structure of the input data than tdistributed Stochastic Neighbor Embedding (t-SNE). The two-dimensional embedding of expression profiles of Shekhar2016 is visualized in Figure 3c. Comparing the embedding derived from the cosine distances (upper left) and the other three embeddings derived from the Hamming distances shows that the hashed expression profiles preserve the original structure of the cell types denoted by different colors. However, some cell-type clusters are more scattered with the 64-bit Hamming distance, which suggests that using 64-bit vectors is insufficient to discriminate cell types by their subtle expression differences. We also observed that the batch effects were considerably mitigated by projecting query cells onto the space spanned by variability derived from the database cells (Additional file 1, Figure 19–22), which is consistent with the observation of Li *et al*. [29].

CellFishing.jl indexes bit vectors in order to accelerate the cell searching process. The algorithm used in this bit search progressively expands the search space that is centered at the query bit vector; thus, using longer bit vectors is not feasible in practice because the search space rapidly expands as more bits are used. In our preliminary experiments, index searches using longer than or equal to 512 bits often consumed more time than linear searches for a wide range of database sizes due to this expansion of the search space. As a result, we limited the length of the bit vectors to 128 or 256 bits in the following experiments. To further reliably find similar cells with these limited bits, CellFishing.jl generates mutually independent bit indexes from a reduced matrix. When searching a database, CellFishing.jl separately searches these bit indexes within the database and aggregates the results by ranking candidate cells by the total Hamming distance from the query. This requires more time than using a single bit index, but as we show in the following section, it appreciably reduces the risk of overlooking potentially neighboring cells.

### Self-mapping evaluation

To compare the performance of CellFishing.jl with that of scmap-cell, we performed five-fold cross-validations by mapping one-fifth of cells randomly sampled from a data set to the remaining four-fifth of cells from the same data set, and computed the consistency and Cohen’s kappa score [30] of the neighboring cell’s label. A value of 1 in the consistency score indicates the perfect agreements of cluster (cell type) assignments and 0 indicates no agreements, while a value of 1 in Cohen’s kappa score indicates the perfect agreements and 0 indicates random assignments. We obtained the ten nearest neighbors for each cell, which is the default parameter of scmap-cell, but only the nearest neighbor was used to compute the scores. This evaluation assumes that cells with similar expression patterns belong to the same cell-type cluster, and hence a query cell and its nearest neighbors ought to have the same cluster assignment. In CellFishing.jl, we varied only the number of bits and number of indexes, which control the trade-off between estimation accuracy and computational cost; other parameters (i.e., the number of features and number of dimensions of a reduced matrix) were fixed to the defaults. In scmap-cell, DGE matrices were normalized and log-transformed using the `normalize` function of the scater package [31], and we varied two parameters: the number of centroids (landmark points calculated by k-means clustering of cells, used to approximate the similarity between cells) and the number of features, in order to find parameter sets that achieve better scores.

Figure 4a, b shows the consistency and Cohen’s kappa scores of CellFishing.jl and scmap-cell with different parameter sets. The overall scores were high (>0.94) for both methods, with the exception of the Plass2018 data set. With the default parameters (see Figure 4 for a description), CellFishing.jl consistently outperformed scmap-cell in both consistency and Cohen’s kappa score for all data sets. In CellFishing.jl, using multiple independent hashes significantly improved the scores, suggesting that using 128 or 256 bits is insufficient to reliably estimate the similarity between cells. Instead of log transformation, using the Freeman–Tukey transformation [20, 32] resulted in almost similar scores (Additional file 2). In scmap-cell, increasing the centroids or incorporating more features did not remarkably improve the scores.

**Figure 4:**
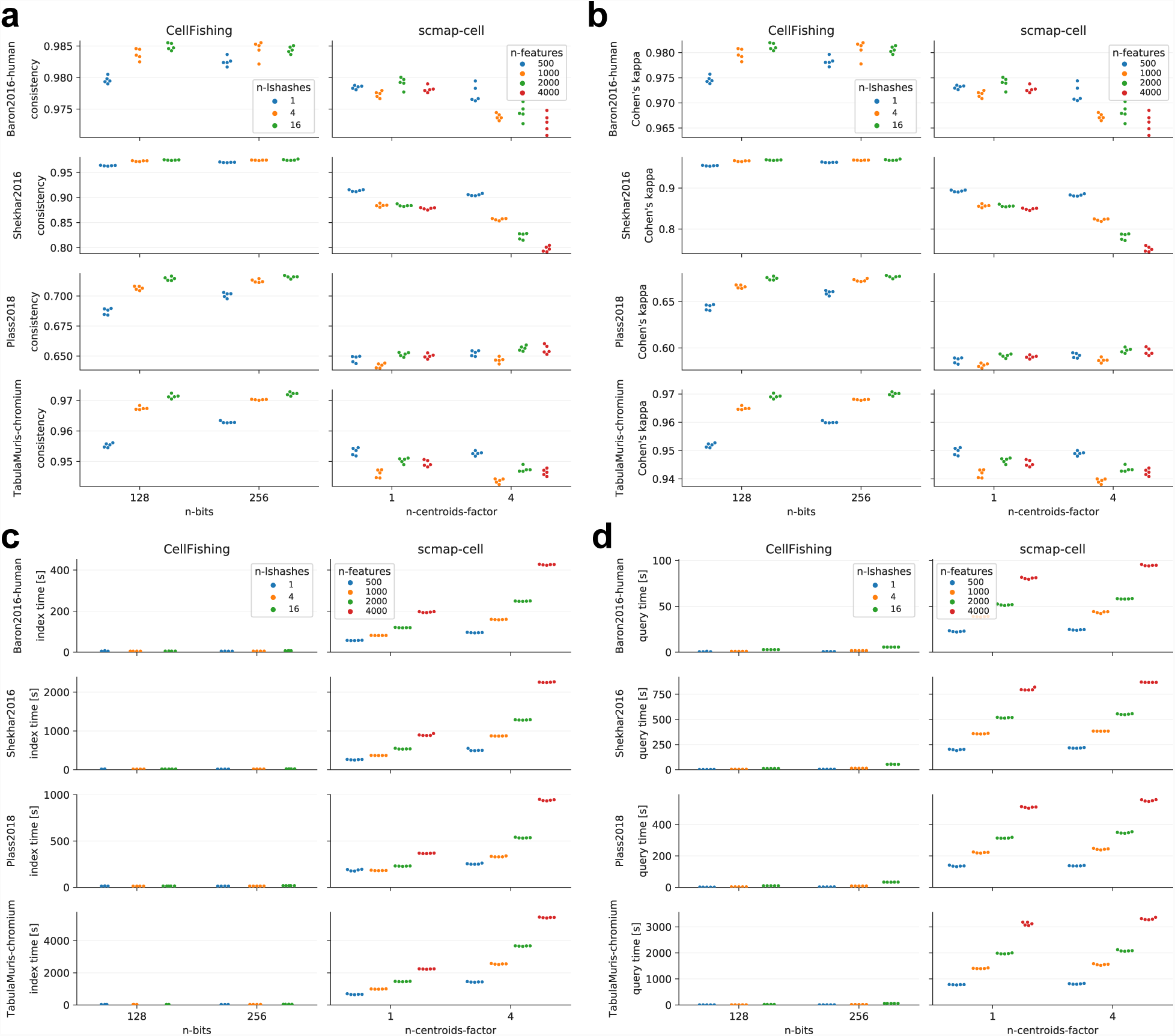
Comparison between CellFishing.jl and scmap-cell by self-mapping. **a** Consistency scores. **b** Cohen’s kappa scores. **c** Index times. **d** Query times. Each data point corresponds to a five-fold cross-validation. The n-bits and n-lshashes parameter of CellFishing.jl are the number of bits and the number of independent locality-sensitive hashes, respectively (n-bits=128 and n-lshashes=4 are the defaults). The n-centroids-factor and n-features parameter of scmap-cell are the multiplier of the default number of centroids (i.e, *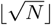*, where *N* refers to the number of reference cells) parameter and the number of features parameter, respectively (n-centroids-factor=1 and n-features=500 correspond to the defaults).

In this evaluation, we also measured the elapsed time of indexing and querying. The measured values do not include the cost of reading data from a disk because it varies depending on the file format and the disk type. In both indexing and querying, CellFishing.jl was faster than scmap-cell by a large margin, as shown in Figure 4c, d. For example, comparing the median of the elapsed time with the default parameters in the TabulaMuris data set, CellFishing.jl was 22 times faster for the indexing time (30.3 vs. 661.9 s) and 118 times faster for the querying time (6.6 vs. 780.6 s).

Since cell types have highly skewed or long-tail distributions (Additional file 1, Figure 1–7), the global evaluation scores used tend to be dominated by large subpopulations and could overlook underrepresented cell types. The cluster-specific consistency scores for each cluster assignment are visualized in Figure 5. Here, the parameters were fixed to the defaults for both CellFishing.jl and scmap-cell. From this figure, CellFishing.jl evidently includes minor cell types, although the scores are relatively unstable for those cell types. For example, while Baron2016 contains only eighteen Schwann cells and seven T cells, CellFishing.jl found these cell types with high consistency (>0.8). Also, CellFishing.jl shows comparable or better consistency scores than scmap-cell for the majority of the cell types.

**Figure 5:**
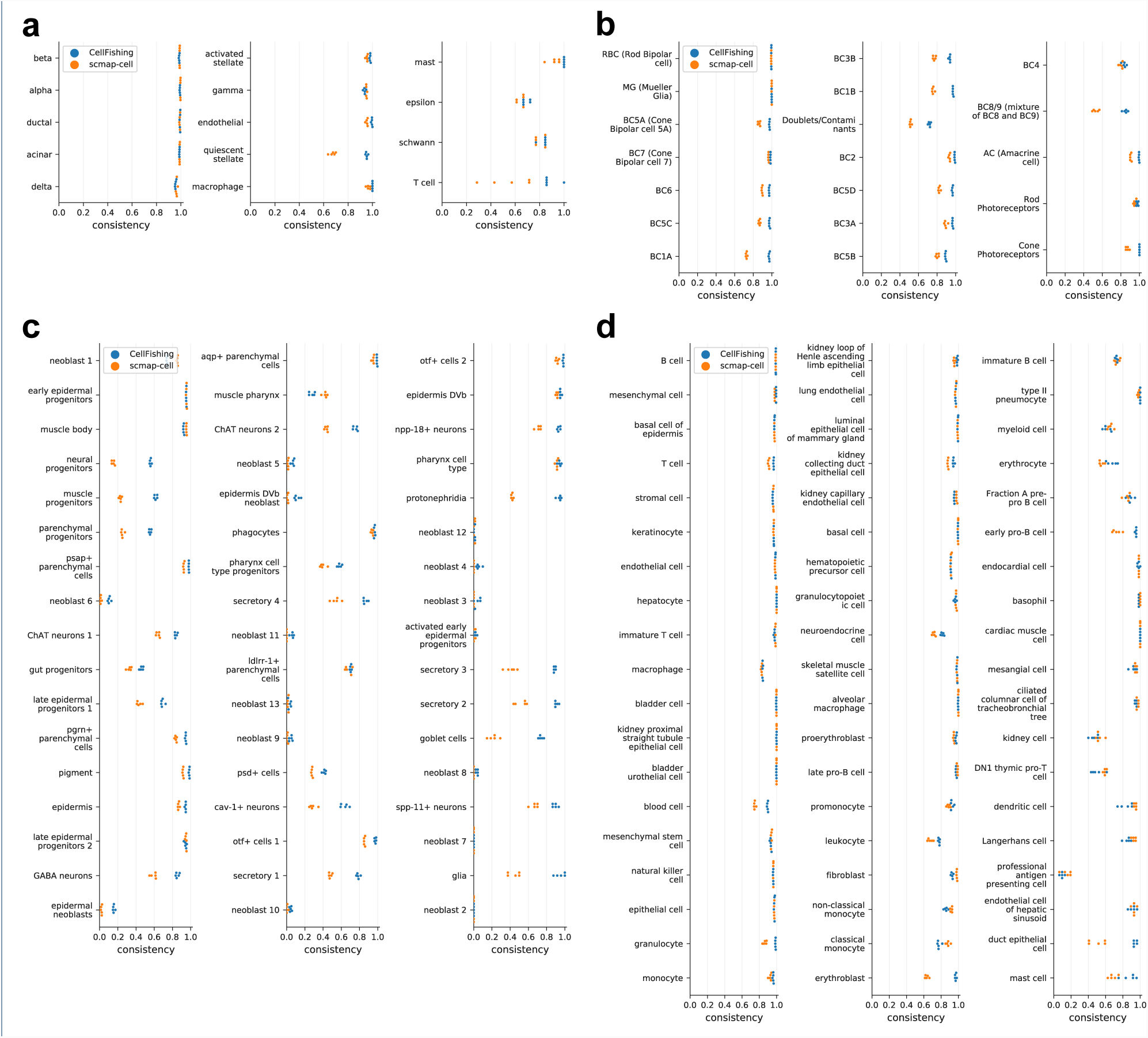
Cluster-specific consistency scores. **a**-**d** Cluster-specific consistency scores of Baron2016 (human), Shekhar2016, Plass2018, and TabulaMuris (Chromium). The *blue* and *orange points* denote the consistency scores of CellFishing.jl and scmap-cell, respectively. Cluster labels are ordered by the cluster size, from top to bottom and then left to right in decreasing order.

The consistency scores of Plass2018 are relatively lower than those of the other data sets. This could be because Plass2018 contains more progenitor or less-differentiated cells, and thus it is more difficult to distinguish these cell types from their expression profiles. As indicated in Figure 6, a significant number of neoblasts and progenitors are assigned to neoblast 1, which is the largest subpopulation (29.35%) of the data set. These subtypes of neoblasts and progenitors are almost indistinguishable from the t-SNE plot (see [10], Figure 1B), which suggests that these cell types have very slight differences in their expression profiles. Still, when comparing CellFishing.jl and scmap-cell, the former more clearly discriminates these cell types.

To see the effects of selected features, we evaluated the scores by exchanging features selected by CellFishing.jl and scmap-cell. When CellFishing.jl used features selected by scmap-cell, we could not detect meaningful differences in the consistency scores except within Plass2018, which improved the score by around 2.0% when `n-features` is 2,000. Likewise, when scmap-cell used features selected by CellFishing.jl, we could not detect meaningful differences except within Shekhar2016, which decreased the score by around 5.5% when `n-min-features` is 500. These results seem to indicate that while scmap-cell selects better features, it has limited effects on the performance of CellFishing.jl.

**Figure 6:**
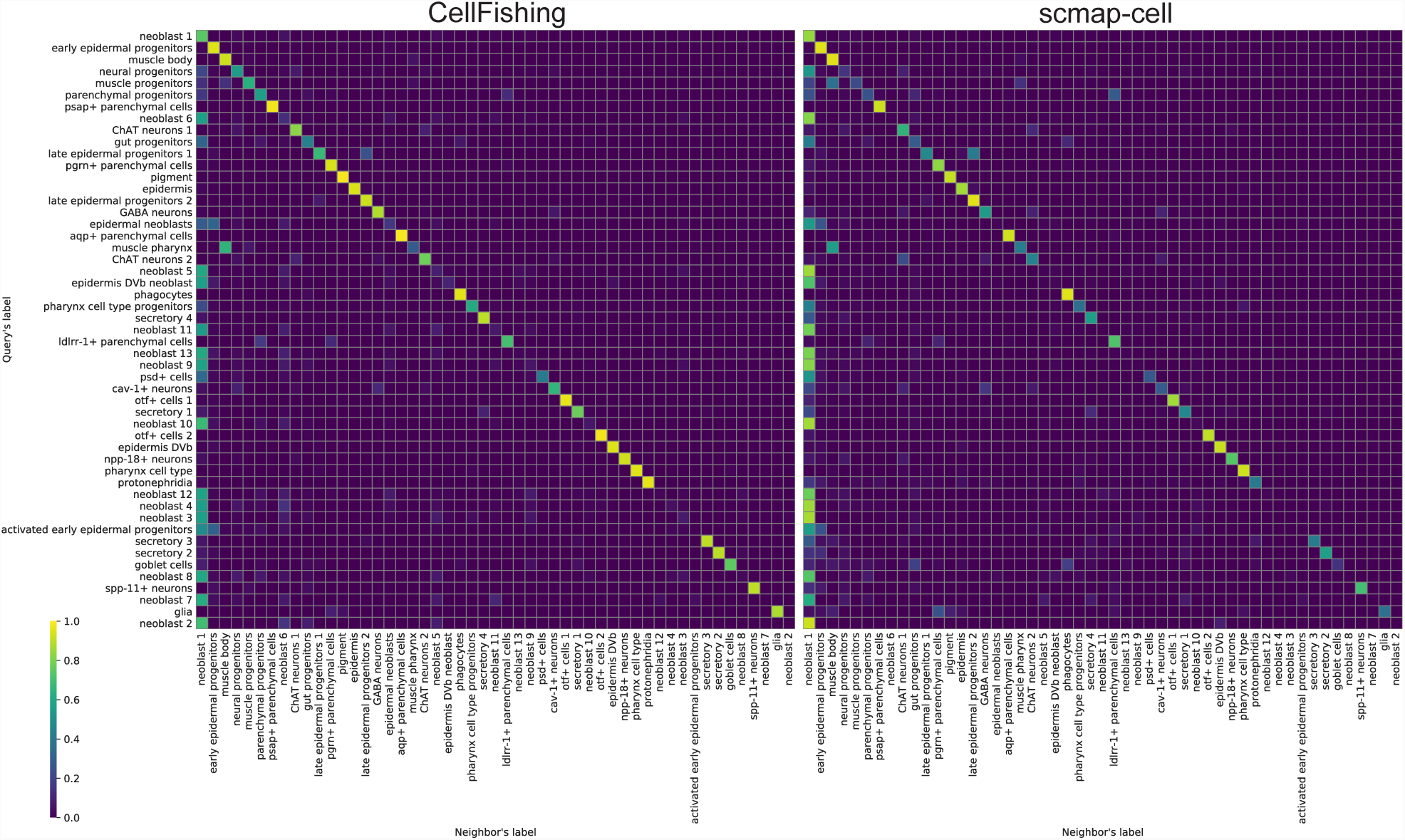
Proportions of cluster assignments for CellFishing.jl and scmap-cell. The rows are labels of query cells and the columns are labels of their nearest neighbor. Cluster labels are ordered by the cluster size, from top to bottom (rows) or left to right (columns) in decreasing order. The values are derived from a cross-validation with the default parameters and normalized row-wise (i.e., the sum of values in a row is 1).

### Evaluation of the search outcome and DEG detection

CellFishing.jl always retrieves the nearest neighboring cells of a query cell in a Hamming space, and its algorithm does not cease to search until all the nearest cells are found. Although this is an important feature because the user does not need to specify any parameters except the number of neighbors when commencing a search, it also means that CellFishing.jl may retrieve far distant cells that are virtually unrelated to the query cell. To evaluate the search outcomes, CellFishing.jl provides a function estimating the cosine similarity between two cells from their Hamming distance. Here we evaluated the cosine similarities of the nearest neighbors either with or without specific cell types in the database in order to simulate a situation wherein a cell type does not exist in the database. We performed five-fold cross-validations in the same manner as in the previous experiment using seven cell types arbitrarily selected from the minor cell types (comprising approximately 2% or less) of each Shekhar2016 and TabulaMuris.

The similarity distributions for the first three nearest neighbors are shown in Figure 7. In the majority of the cell types, the cosine similarity remarkably drops if cells of the query cell type are removed from the database. Notably, the similarity distribution of amacrine cells from Shekhar2016 (Figure 7a) and basal cells from TabulaMuris (Figure 7b) after removal hardly overlap with the distribution before removal. However, the differences are less evident in some cell types, such as immature B cells and kidney cells from TabulaMuris, which is not surprising because these cell types have similar cells with different labels. For example, 9%, 9%, and 81% of immature B cells were mapped to early pro-B cells, late pro-B cells, and B cells, respectively, when immature B cells were removed from the database. This result suggests that CellFishing.jl has the ability to identify similar cells even when a database does not include cells with the exact cell type of a query.

**Figure 7:**
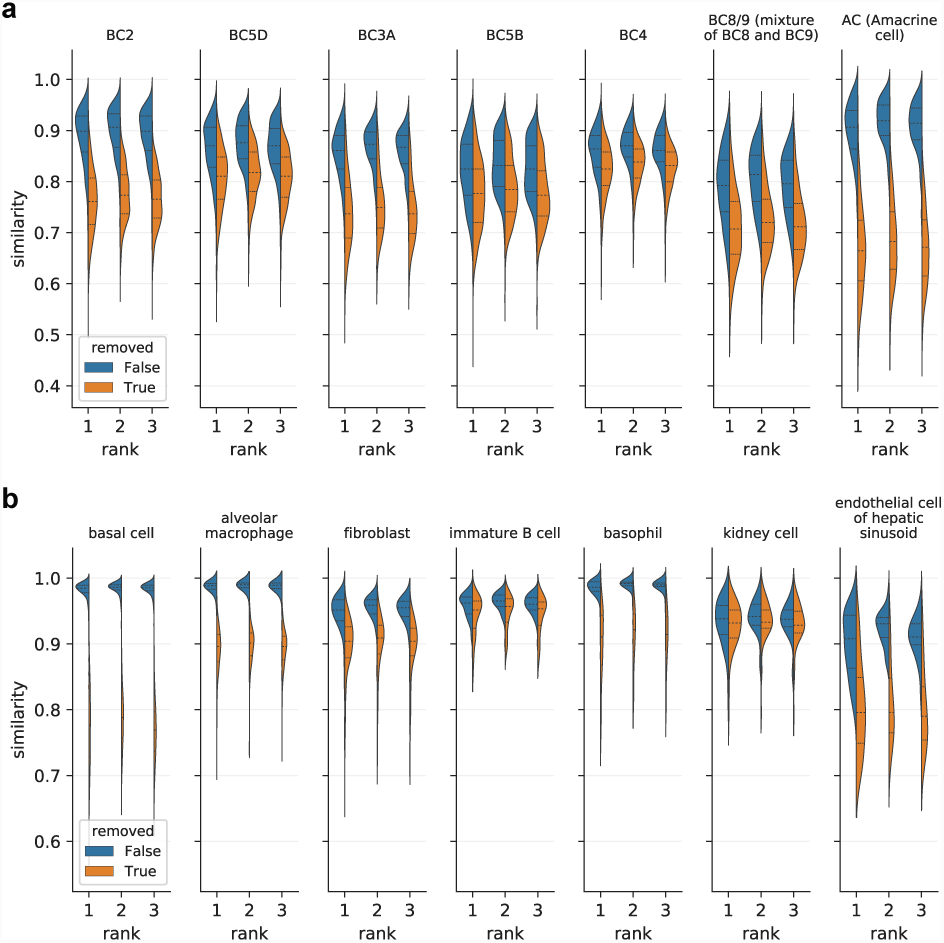
Distributions of similarities with or without a specific cell type. **a** Seven cell types from Shekhar2016. **b** Seven cell types from TabulaMuris. The x-axis (*rank*) refers to the rank of the nearest neighbors (e.g., rank = 1 refers to the nearest cell of a query), and the y-axis (*similarity)* refers to the cosine similarity estimated from the Hamming distance. The *blue* or *orange* regions denote the distribution of cosine similarities with or without the cell type shown in the title, respectively. The *lower dotted lines, dashed lines*, and *upper dotted lines* denote the first, second, and third quartiles, respectively.

Once the user identifies cells with high similarity, the next question will probably be concerning the differences between the query and its nearest neighbors. To answer this, CellFishing.jl provides a function to detect DEGs of the query cell compared to reference cells in the database. It estimates the average expression of neighboring cells using their raw UMI counts and then computes the probability of observing the count of the query or more extreme values for each gene. Although a CellFishing.jl database requires additional space to store raw counts for DEG detection, it efficiently compresses the raw counts and avoids loading the whole count matrix into memory, thereby saving disk space and memory. Here we focus on the performance evaluation of our DEG detection method, and the cost required for storing raw counts is presented in a following section.

For the performance evaluation, we selected immature B cells of TabulaMuris as an example because they are very similar to other subtypes of B cells, as can be seen in Figure 7b, and the development of B lymphocytes is well characterized. First, all immature B cells were removed from the data set and mapped to the remaining cells. Then, we compared the query cells with their nearest neighbors, and filtered genes with a probability of being less than 10^−4^(0.01%) as significant DEGs for each query cell (see Methods for details). Out of the 113 immature B cells mapped as queries, 9, 10, and 91 cells were mapped to early pro-B cells, late pro-B cells, and B cells, respectively, and the other 3 cells were mapped to separate subtypes of lymphocytes. We ignored these three cells here because they represent a relatively minor percentage of the results (less than 3%), and it is impossible to deduce meaningful conclusions only from a single sample mapped to a unique cell type. We summarize the detected DEGs in Table 2; the result was reasonably robust to the selection of the number of neighbors being between 5 and 20 (see Additional file 2); thus, here we show results when the number of neighbors is set to 10 (i.e., the default parameter).

**Table 2:** Top DEGs of immature B cells

| Early pro-B cell |  |  |  | Late pro-B cell |  |  |  | B cell |  |  |  |
| --- | --- | --- | --- | --- | --- | --- | --- | --- | --- | --- | --- |
| Negative |  | Positive |  | Negative |  | Positive |  | Negative |  | Positive |  |
| *Vpreb1* | 9 | *Dnajc7* | 8 | *Hmgb2* | 6 | *Malat1* | 7 | *Cd74* | 41 | *S100a8* | 83 |
| *Igll1* | 9 | *Ly6d* | 6 | *Ptma* | 2 | *Tmsb4x* | 4 | *H2-Ab1* | 19 | *Beta-s* | 81 |
| *Dntt* | 8 | *Malat1* | 5 | *H2afz* | 2 | *Actb* | 3 | *H2-Eb1* | 15 | *S100a9* | 76 |
| *Crip1* | 3 | *Tmsb10* | 5 | *Tuba1b* | 1 | *Dnajc7* | 3 | *Malat1* | 15 | *Fos* | 36 |
| *Rps24* | 1 | *Cd74* | 4 | *Stmn1* | 1 | *Tmsb10* | 3 | *H2-Aa* | 15 | *Camp* | 30 |
| *Ucp2* | 1 | *Herpud1* | 4 | *Hist1h2ao* | 1 | *Ly6d* | 2 | *Rps28* | 10 | *Klf2* | 25 |
| Total | 31 | Total | 91 | Total | 13 | Total | 47 | Total | 182 | Total | 752 |

The number of cells for each gene detected as significantly different from the nearest cell and its neighbors. The list is truncated for brevity at the top-six DEGs after sorting all detected DEGs by the number of occurrences for each cell type (ties are ordered arbitrarily). The last row (*Total*) shows the total number of DEGs including ones omitted from each list.

Next, we carefully examined these genes to validate that biologically meaningful results were obtained using our analysis method. For the nine cells mapped to early pro-B cells, *Vpreb1* and *Dntt* are negatively regulated when compared to immature B cells as shown in Table 2, which is consistent with the annotation by scientific experts because these two genes are used to distinguish immature B cells and early pro-B cells (see [12], Supplementary Information). Although *Igll1* was not used as a marker gene in the annotation [12], it is known to play a critical role in B-cell development (see UniProt accession: P20764). For the ten cells mapped to late pro-B cells, *Hmgb2* is involved in V(D)J recombination (see UniProt accession: P30681). We detected fewer DEGs from late pro-B cells in comparison with early pro-B cells and B cells with the same threshold, even if the number of cells for each cell type is considered, which may reflect the fact that late pro-B cells more closely resemble immature B cells than the other two cell types. For the 91 cells mapped to B cells, *Cd74* is negatively regulated, and was used as a marker gene to discriminate immature from mature B cells. Similarly, *H2-Ab1, H2-Eb1*, and *H2-Aa*, which encode components of the major histocompatibility complex present on the surface of antigen-presenting cells, are negatively regulated, suggesting that immature B cells do not actively express these genes as much as mature B cells. We also found that *S100a8, Betas, S100a9, Fos*, and *Camp* are positively regulated in immature B cells; although these genes appear to be upregulated in a tissue-specific manner rather than in a cell type-specific manner, cytokines encoded by *S100a8* and *S100a9* were recently reported to regulate B lymphopoiesis in rabbit bone marrow [33]. Overall, our method detected reasonable DEGs in many cases; however, we could not find evidence implying the relationship between *Malat1* (a long intergenic noncoding RNA) and B-cell development, and therefore, we hypothesize that this is a false positive due to its high variability regardless of cell type.

### Mapping cells across different batches

Researchers are often interested in searching for cells across different experiments or batches. To verify the robustness of our method in this situation, we performed cell mapping from one batch to other batches and evaluated the performance scores. Shekhar2016 consists of two different batches (1 and 2), which exhibit remarkable differences in their expression patterns (see [9], Figure S1H). We mapped cells from one batch to the other using CellFishing.jl and scmap-cell, and calculated the consistency and Cohen’s kappa score. Similarly, we selected two batches (plan1 and plan2, derived from wild-type samples) from Plass2018 and mapped cells from one batch to the remaining batches. This data set exhibits relatively weak batch effects. The default parameters were used in this experiment. Figure 8a shows the results of these two data sets for consistency scores. In both data sets, the consistency scores of CellFishing.jl were close to the mean score obtained from the self-mapping experiment. Moreover, their distances from the mean score in CellFishing.jl were smaller than those in scmap-cell, suggesting that CellFishing.jl is more robust to batch effects. The results of the Cohen’s kappa scores were consistent with these results (Additional file 1, Figure 44). We predict that the differences in scores between batches are due to differences in cluster sizes. For example, batch 1 of Shekhar2016 contains many more rod bipolar cells than batch 2, while the latter contains more minor cell types than the former (Figure 8b). The discrepancy in cluster sizes across batches leads to a difference in scores because each cluster has a different consistency score, as observed in the self-mapping experiment.

**Figure 8:**
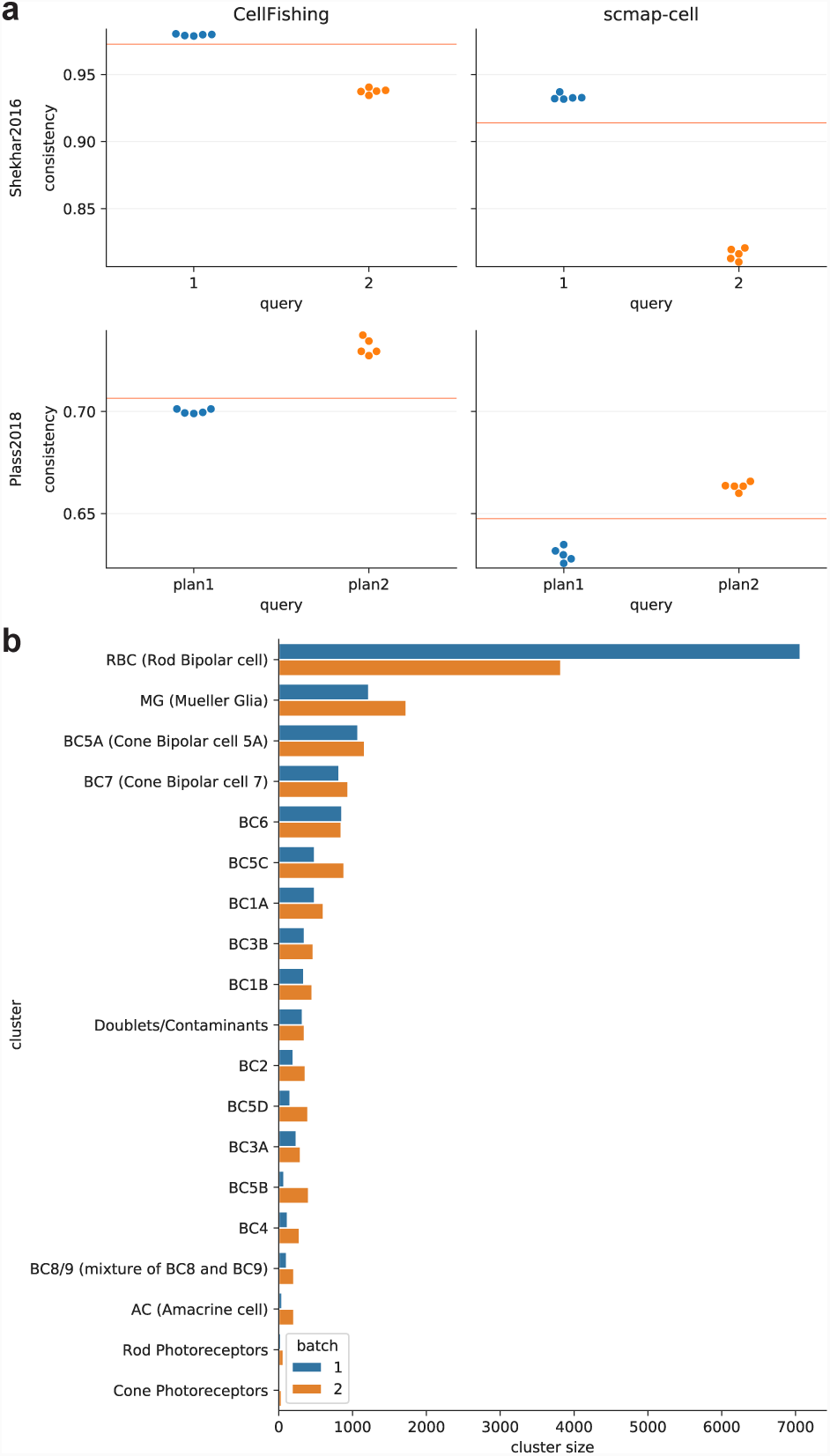
Cell mapping across batches. **a** Consistency scores of cell mapping (Shekhar2016 and Plass2018). The *red lines* denote the mean score of the corresponding self-mapping experiment with the same parameters. **b** Distribution of cluster sizes (Shekhar2016).

### Mapping cells across different species

Comparing transcriptome expressions across different species provides important information on the function of unknown cell types. Since the Baron2016 data set includes cells derived from human and mouse, we attempted to match cells between both species. To match genes from different species, we downloaded a list of homologous genes between human and mouse from the Vertebrate Homology database of Mouse Genome Informatics and removed non-unique relations from the list. A total of 12,413 one-to-one gene pairs were included. We compared the performance of CellFishing.jl and scmap-cell with the default parameters. In CellFishing.jl, the feature statistics were estimated only from the query cells because they were expected to be considerably different between species. The consistency scores of CellFishing.jl and scmap-cell mapping from human to mouse were 0.681 and 0.563, respectively, and from mouse to human were 0.787 and 0.832, respectively. The Cohen’s kappa scores of CellFishing.jl and scmap-cell mapping from human to mouse were 0.599 and 0.455, respectively, and from mouse to human were 0.715 and 0.753, respectively. These results show that CellFishing.jl and scmap-cell are roughly comparable for cell mapping accuracy across different species.

### Mapping cells across different protocols

Mapping cells across different sequencing protocols is also important. To validate the robustness of CellFishing.jl in this case, we used TabulaMuris, which consists of two data sets derived from different sequencing platforms, and mapped 44,807 cells sequenced with Smart-Seq2 [34] to 54,967 cells with Chromium. The default parameters were used in this experiment. Because cluster labels are not identical between the two data sets, it is not possible to compute the consistency or Cohen’s kappa score. Thus, the matrices of cluster assignments are visualized in Figure 9. On the whole, these two matrices show very similar assignment patterns, although CellFishing.jl failed to detect a large number of fibroblast cells.

**Figure 9:**
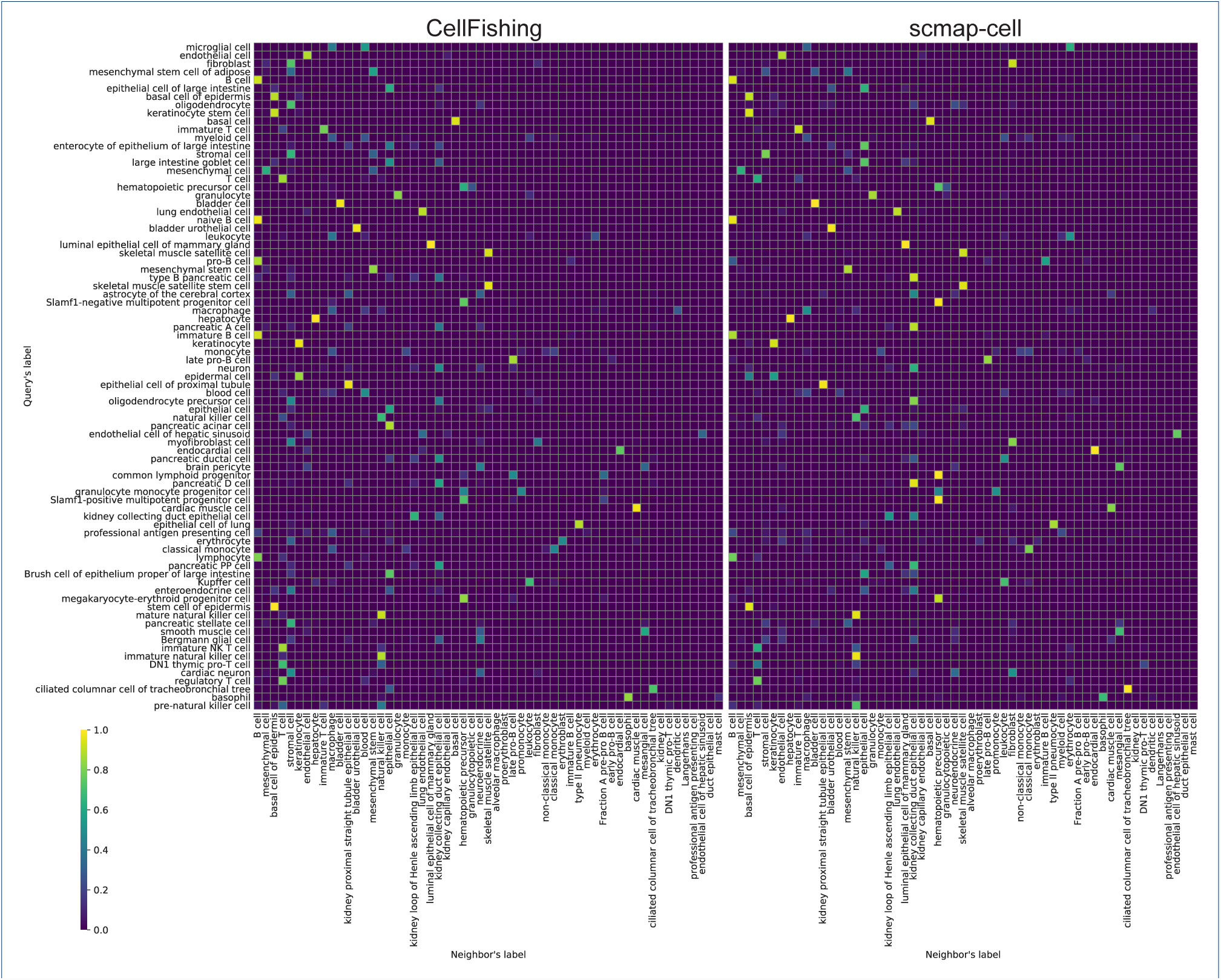
Cluster assignments across different protocols (TabulaMuris). **a** CellFishing.jl. **b** scmap-cell. The rows are the query’s cluster labels (Smart-Seq2) and the columns are its neighbor’s cluster labels (Chromium).

### Saving and loading databases

CellFishing.jl is designed to search multiple scRNA-seq experiments, and the database objects can be serialized into a disk and later deserialized. For this purpose, CellFishing.jl provides a pair of functions to save and load a database object into and from a file. To verify the feasibility of this approach, we measured the elapsed time of saving and loading a database object with cell names as metadata, as well as the memory and file size of the object. The results are summarized in Table 3. The memory and file size of database objects are reasonably small, even for current low-end laptop computers. Although the memory and file sizes become several times larger when raw UMI counts are stored in the database, the actual memory usage is usually much smaller because the raw counts are loaded on request as described in Methods. The elapsed time required for saving and loading a database is also small relative to the time required for querying. From these results, we predict that CellFishing.jl can be used to quickly search multiple scRNA-seq experiments by building and serializing their database objects in advance.

**Table 3:** Performance of saving and loading databases and their memory and file size.

| Data set | #bits | Raw counts | Size [MiB] |  | Time [ms] |  |
| --- | --- | --- | --- | --- | --- | --- |
|  |  |  | Memory | File | Save | Load |
| Baron2016-human | 128 | - | 3.1 | 2.9 | 2.3 | 2.4 |
|  |  | + | 10.9 | 10.7 | 6.5 | 3.1 |
|  | 256 | - | 5.4 | 5.2 | 3.7 | 5.3 |
|  |  | + | 13.2 | 13.0 | 8.0 | 3.6 |
| Shekhar2016 | 128 | - | 8.2 | 7.8 | 5.6 | 8.3 |
|  |  | + | 29.7 | 29.3 | 16.3 | 6.7 |
|  | 256 | - | 14.6 | 14.1 | 9.1 | 8.3 |
|  |  | + | 36.1 | 35.7 | 20.7 | 11.1 |
| Plass2018 | 128 | - | 6.7 | 6.4 | 4.7 | 5.0 |
|  |  | + | 17.5 | 17.2 | 10.4 | 6.2 |
|  | 256 | - | 12.0 | 11.6 | 8.0 | 9.6 |
|  |  | + | 22.8 | 22.4 | 14.9 | 9.2 |
| TabulaMuris-chromium | 128 | - | 13.8 | 13.0 | 8.6 | 12.5 |
|  |  | + | 67.2 | 66.5 | 35.3 | 16.9 |
|  | 256 | - | 24.9 | 24.2 | 13.8 | 18.0 |
|  |  | + | 78.3 | 77.6 | 42.4 | 21.9 |

Plus and minus signs in the raw counts column denote with or without raw UMI counts, respectively. The memory size includes the region read as a memory-mapped file. Each value is the median of ten measurements.

### Scalability for large data sets

To check the scalability of our approach for large data sets, we measured the index time, query time, and the memory size of a database by changing the number of cells within the database. In this benchmark, we randomly sampled 10,000 cells from the 1M neurons data set as queries and then randomly sampled *N*= 2^13^,2^14^, *…*,2^20^ cells from the remaining cells to create a database (2^13^ = 8,192 and 2^20^ = 1,048,576, which covers a wide range of high-throughput scRNA-seq experiments); there are no overlapping cells between or within the query and database sets. The number of bit indexes was fixed to the default (i.e., 4) in all cases. For comparison, we also benchmarked the performance of the linear search that scans all hash values in a database instead of using indexes. The elapsed time does not include the time of loading expression profiles from a file.

The benchmark results are summarized in Table 4. As for the index search, the query time is sublinear to the database size, while the index time and memory size are roughly linear. For example, when 128-bit vectors are used, the query time becomes only 2.8 times longer in the same range, as the database size becomes 128 times larger from *N*= 2^13^ to 2^20^. The linearities of the index time and memory size are expected, because when building a database all the reference cells need to be scanned and stored, though these are not wholly proportional to the database size because some overhead costs are included in the measured values (e.g., generating and storing projection matrices). Also, the memory usage per cell is only 183.3 bytes for 128-bit vectors and 365.9 bytes for 256-bit vectors when *N*= 2^20^, which is approximately 22 and 11 times smaller than storing UMI counts of 1,000 genes in 32-bit integers, respectively.

**Table 4:** Scalability of CellFishing.jl (1M neurons)

| | #bits | Index | Database size $N$ | | | | | | | |
| --- | --- | --- | --- | --- | --- | --- | --- | --- | --- | --- |
| | | | $2^{13}$ (1.0) | $2^{14}$ (2.0) | $2^{15}$ (4.0) | $2^{16}$ (8.0) | $2^{17}$ (16.0) | $2^{18}$ (32.0) | $2^{19}$ (64.0) | $2^{20}$ (128.0) |
| Index time [s] | 128 | - | 1.4 (1.0) | 2.5 (1.8) | 4.5 (3.3) | 8.5 (6.2) | 16.4 (12.0) | 31.9 (23.3) | 63.3 (46.3) | 121.3 (88.8) |
|  | 128 | + | 1.4 (1.0) | 2.5 (1.8) | 4.7 (3.2) | 8.6 (6.0) | 16.5 (11.4) | 32.0 (22.1) | 64.1 (44.3) | 125.0 (86.3) |
|  | 256 | - | 1.4 (1.0) | 2.7 (1.9) | 4.8 (3.4) | 8.9 (6.3) | 16.1 (11.4) | 32.3 (22.7) | 65.0 (45.7) | 125.4 (88.2) |
|  | 256 | + | 1.4 (1.0) | 2.7 (1.9) | 4.7 (3.3) | 9.0 (6.3) | 16.8 (11.7) | 33.7 (23.5) | 68.6 (47.9) | 132.2 (92.2) |
| Query time [s] | 128 | - | 1.8 (1.0) | 2.1 (1.2) | 2.9 (1.6) | 4.2 (2.4) | 7.0 (3.9) | 12.6 (7.0) | 23.7 (13.2) | 52.9 (29.4) |
|  | 128 | + | 2.2 (1.0) | 2.2 (1.0) | 2.4 (1.1) | 2.4 (1.1) | 2.9 (1.3) | 3.0 (1.3) | 4.2 (1.9) | 6.3 (2.8) |
|  | 256 | - | 2.0 (1.0) | 2.6 (1.3) | 3.8 (1.8) | 6.1 (3.0) | 10.5 (5.2) | 19.6 (9.6) | 46.7 (22.9) | 109.3 (53.7) |
|  | 256 | + | 3.4 (1.0) | 3.6 (1.1) | 3.9 (1.2) | 4.6 (1.4) | 5.2 (1.6) | 8.8 (2.6) | 13.4 (4.0) | 22.1 (6.6) |
| Memory size [MiB] | 128 | - | 1.3 (1.0) | 1.8 (1.4) | 2.8 (2.2) | 4.8 (3.7) | 8.8 (6.8) | 16.8 (12.9) | 32.8 (25.2) | 64.7 (49.7) |
|  | 128 | + | 3.0 (1.0) | 4.8 (1.6) | 8.3 (2.8) | 14.8 (5.0) | 27.1 (9.1) | 51.0 (17.1) | 93.0 (31.2) | 183.3 (61.6) |
|  | 256 | - | 1.9 (1.0) | 2.9 (1.6) | 4.9 (2.6) | 8.9 (4.8) | 16.9 (9.0) | 32.9 (17.6) | 64.9 (34.7) | 128.8 (68.9) |
|  | 256 | + | 5.2 (1.0) | 8.9 (1.7) | 15.8 (3.0) | 28.8 (5.5) | 53.7 (10.3) | 101.3 (19.3) | 192.9 (36.8) | 365.9 (69.9) |

Plus and minus signs in the index column denote the index search and the linear search, respectively.
The values enclosed by parentheses are relative to those of $N = 2^{13}$ . Each value is the median of five measurements.

The index time remains almost unchanged between the index and the linear search, suggesting that the computational cost of creating hash indexes is, in effect, negligible. In contrast, the gap of the query time between the two search methods expands as the database becomes larger, which can be attributed to the searching phase of bit vectors because the cost of the preprocessing phase is constant (Figure 10). In addition, even though indexing bit vectors requires extra memory, the relative difference from a database without indexes is approximately equal to or less than three times, and the absolute memory size of a database with indexes is small enough for modern computing environments, including laptops.

**Figure 10:**
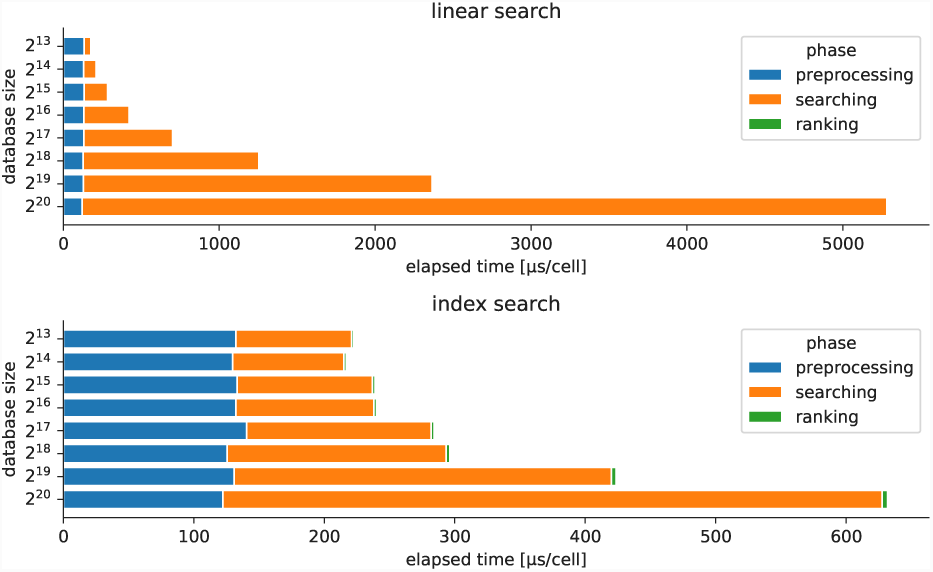
Computational costs of different phases. The upper and lower plots show the costs of the linear and index searches, respectively. The elapsed time of each phase was measured using the time_ns function and divided by the number of query cells to compute the average time per cell.

We also evaluated the consistency scores and found that they were slightly improved by incorporating more cells into a database (when using 128-bit vectors, the consistency scores were 0.775 and 0.830 for *N*= 2^13^ and *N*= 2^20^, respectively) (Additional file 1, Figure 51). This result suggests that building a database with more cells plays an important role in identifying cell types. The consistency scores did not vary significantly between the index and linear search as expected, because nearest neighbors found by the index search have the same Hamming distances as those found by the linear search. In summary, indexing bit vectors is effective in reducing the search time for high-throughput scRNA-seq data and is scalable for extremely large data sets containing more than one million cells.

## Discussion

Our LSH-based method is particularly suitable for middle or large scale scRNA-seq data sets because it circumvents costly brute-force search by using indexes for low-dimensional bit vectors. We considered relatively large data sets consisting of at least ten thousand cells, one of which contains more than one million cells, and confirmed our approach outperforms scmapcell, the state-of-the-art method for cell searching, in both accuracy and throughput using real scRNA-seq data sets. Searching across multiple experiments will be feasible because our method is reasonably robust to batch differences, and serialized database objects can be loaded quickly. In this paper, we did not compare our method with CellAtlasSearch, mainly because its source code is not freely available, and its algorithm is not well described in the original paper, which makes it difficult to compare the performance fairly. Moreover, CellAtlasSearch requires a GPU to achieve its maximum performance, but this is not always available on server machines.

The application of cell searching is not limited to mapping cells between different data sets. The task of finding similar cells within a data set is a subroutine in many analysis methods, such as data smoothing [20], clustering [35], community detection [36], and visualization [37]. As we have demonstrated in the self-mapping experiment, our LSH-based method can find similar cells within a data set with high accuracy and throughput; thus, it would be possible to speed up analysis by utilizing our cell search method in lieu of the currently available method.

The feature selection used in CellFishing.jl is relatively simple and rapid. We confirmed that it works well with our search method; however, we also found that the criterion based on the dropout rate used in scmap-cell performed slightly better in a data set. This fact suggests that our simple selection method is not necessarily suitable for all scRNA-seq data sets, and a more careful feature selection, such as adding marker genes selected by domain experts or more careful selection methods such as GiniClust [38], may significantly improve the accuracy of cell typing. For this purpose, CellFishing.jl provides for addition and removal of specific features to or from a feature set.

Handling batch effects is still a persistent problem in scRNA-seq. We performed cell mapping experiments across different batches and protocols and confirmed that the performance of our approach is at least comparable to scmap-cell. We consider that the robustness of CellFishing.jl comes from projecting expression profiles to a space with dataset-specific variability, as previously reported by Li *et al*. [29]. This type of technique is also discussed in the context of information retrieval and is termed *folding-in*[39]. Although substantial literature exists on the removal of batch effects in scRNA-seq data [25, 40–42], the existing methods require merging raw expression profiles of reference and query cells to obtain their batch-adjusted profiles, and the computational costs are relatively high; these characteristics are not suitable for our low-memory/high-throughput search method. In contrast, folding-in is very affordable because projection matrices can be computed when building a database and reused for any query.

The DEG detection method introduced in this paper assumes that the database encompasses enough cells to retrieve a small group of homogeneous neighbors containing no biological differences, and each UMI count follows a Poisson distribution. The former can be justified by considering the high-throughput characteristic of recent scRNA-seq experiments, the feasibility of which we have demonstrated using the TabulaMuris data set; the latter is experimentally verified by several works [8, 43, 44]. However, some highly expressed genes, such as *Malat1*, seem to be exceptions to these model assumptions, and as a consequence, despite it being unlikely that *Malat1* is related to biological differences between cells, it was falsely detected as a DEG in many cells within our experiment. We predict that this problem can be partially mitigated by replacing point estimation of the mean expression with some interval estimation, such as Bayesian inference.

In this work, we have focused on unsupervised cell searching: no cellular information is required except their transcriptome expression data. This makes our method even more useful because it is widely applicable to any scRNA-seq data with no cell annotations. However, incorporating cell annotations or prior knowledge of reference cells could remarkably improve our method’s performance. For example, if cell-type annotations are available, it would be possible to generate tailored hash values separating cell types more efficiently by focusing on their marker genes. Further research is needed in this direction.

## Conclusions

In summary, the new cell search method we propose in this manuscript outperforms the state-of-the-art method and is scalable to large data sets containing more than one million cells. We confirmed that our method considers very rare cell types and is reasonably robust in response to differences between batches, species, and protocols. The low-memory footprint and database serialization facilitate comparative analysis between different scRNA-seq experiments.

## Methods

### Preprocessing

In the preprocessing step, biological signals are extracted from a DGE matrix. When building a database of reference cells, CellFishing.jl takes a DGE matrix of *M* rows and *N* columns, with the rows being features (genes) and the columns being cells, along with some metadata (e.g., cell names). In scRNA-seq analysis, it is common practice to filter out low-abundance or low-variance features because these do not contain enough information to distinguish differences among cells [45]. In the filtering step of CellFishing.jl, features that have smaller maximum count across cells than a specific threshold are excluded. We found that this criterion is rapid and sometimes more robust than other gene filtering methods, such as selecting highly variable genes [46]. The optimal threshold depends on various factors such as sequencing protocol and depth. CellFishing.jl uses a somewhat conservative threshold that retains at least 10% of features by default; however, retaining an excessive amount of features would not be detrimental with regard to accuracy because CellFishing.jl uses the principal components of the data matrix in a later step. One parameter can change the threshold, and it is possible to specify a list of features to be retained or excluded if, for example, some marker genes are known *a priori*.

The filtered DGE matrix is then normalized so that the total counts are equal across all the cells, which reduces the differences in the library size among cells [43]. After normalization, each count *x* is transformed by log transformation log(*x*+ 1), which isa common transformation in scRNA-seq, or FTT 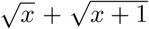 [32]. FTT is a variance-stabilizing transformation assuming the Poisson noise, which is observed as a technical noise in scRNA-seq [20, 43]. Also, the computation of FTT is significantly faster than that of log transformation. The user can specify a preferable transformation, and the choice will be saved in a preprocessor object. In this paper, we used log transformation if not otherwise specified. After transformation, the feature counts are standardized so that their mean and variance are equal to zero and one, respectively.

Finally, the column vectors in the matrix are projected onto a subspace to reduce the number of dimensions. The advantages of this projection are three-fold: (1) computational time and working space for preprocessing are saved; (2) the number of bits required for hash expression profiles is reduced; and (3) batch effects between the query and the database are mitigated. As recently reported [29], projecting data onto a subspace defined by variability in reference cells greatly reduces unwanted technical variation and improves the accuracy of cell-type clustering. A similar approach is found elsewhere [6]. In CellFishing.jl, a subspace with high variance is calculated by applying the SVD to the reference data matrix. Since the number of cells may be extremely large and singular vectors corresponding to small singular values are irrelevant, CellFishing.jl uses a randomized SVD algorithm [24, 26] that approximately computes singular vectors corresponding to the top *D* singular values. The dimension of the subspace *D* is set to 50 by default, but it can be changed easily by passing a parameter. The net result of the preprocessing phase is a matrix of *D* rows and *N* columns. The information of the pre-processing phase (e.g., gene names and projection matrices) is stored in the database object, and the same process is applied to the count vectors of query cells.

### Hashing expression profiles

LSH is a technique used to approximately compute the similarity between two data points [16], which hashes a numerical vector *x* ∈ ℝ ^*D*^ to a bit vector *p* ∈ {0,1}^*T*^ that preserves some similarity or distance between the original vectors. Hashed values (bits) of LSH collide with high probability if two original vectors are similar. This is the fundamental property of hash functions used in LSH, because by comparing binary hash values of numerical vectors their similarity can be estimated, which bypasses the time-consuming distance computation between the numerical vectors.

In LSH, it is common to use *T* hash functions to generates *T* bits; each hash function returns zero or one for an input vector, and the results of multiple hash functions for the vector are bundled into a bit vector of length *T.* More formally, given a similarity function sim(·, ·): ℝ^*D*^ × ℝ^*D*^ *→*[0, 1] that measures some similarity between two data points, an LSH function *h*(*·*): ℝ^*D*^ *→*{0, 1} is defined as a function that satisfies the following relation:

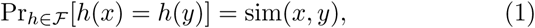

where Pr_*h∈F*_[*h*(*x*) = *h*(*y*)] is the probability of hash collision for a hash function *h*(*·*) generated from a family of hash functions *Ƒ* given a pair of points *x* and *y*. The existence of a family of hash functions that satisfies equation 1 depends on the similarity function sim(·, ·). Similarity, if exists, can be approximated by randomly generating hash functions from *Ƒ* as follows:

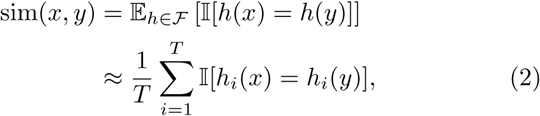

where 𝔼 _*h* ∈ *Ƒ*_ [.] is an expectation over *Ƒ*, 𝕝 [.] is the indicator function, and 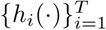 is a set of hash functions sampled from *Ƒ*.

In this work, we used a signed random projection LSH (SRP-LSH) [16] to hash expression profiles, as it estimates angular similarity between two numerical vectors, which is reasonable to measure the similarity between expression profiles, and is straightforward to implement. Briefly, SRP-LSH divides a set of data points in a space into two disjoint sets by drawing a random hyperplane on the space, then the data points in a set of the two are hashed to zeros, with the remaining points hashed to ones. This procedure is repeated *T* times to get a bit vector of length *T* for each data point. Intuitively, the closer two points are to each other, the more likely it is that they occur in the same half-space with respect to a random hyperplane. Therefore, we can stochastically estimate the similarity of two points by calculating the Hamming distance of their bit vectors. In SRP-LSH, the angular similarity function sim 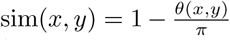, where *θ*(*x, y*) refers to the angle between two vectors *x* and *y*, is estimated from equation 2. In this way, we randomly generate *T* hyperplanes, calculate their hash values, and bundle their zero or one bits into a bit vector of length *T.*

While SRP-LSH dramatically reduces the computational time and space of the approximate search of neighboring cells, it suffers from its high variance of the similarity estimator. To alleviate the problem, CellFishing.jl orthogonalizes the vectors perpendicular to the random hyperplanes (normal vectors) because it reduces the variance of the estimator by removing the linear dependency among hyperplanes without introducing estimation bias [27]. Specifically, *T* normal vectors of length *D* are generated independently from the standard isotropic Gaussian distribution and then orthogonalized using the QR decomposition. If *T* is larger than *D*, the *T* vectors are divided into [*T/D*] batches of vectors, each of which contains at most *D* vectors, and the vectors in a batch are orthogonalized separately.

### Indexing hash values

Although comparing bit vectors is much faster than comparing numerical vectors, it is still lengthy process to scan all the bit vectors in a database, especially in a large database. To reduce the computational cost of hash searching, CellFishing.jl creates search indexes in the space of bit vectors. Specifically, CellFishing.jl creates multi-index hash (MIH) [19] tables to find the nearest neighbors quickly in the Hamming space ℋ_*T*_:= 0, 1 ^*T*^. Briefly, an MIH divides bit vectors into shorter subvectors and indexes them separately using multiple associative arrays (tables); when searching for the nearest neighbors of a query, it divides the query bit vector in the same way and picks candidates of neighbors from the tables of the subvectors. It then computes the full Hamming distances between the query and the candidates by scanning the list of the candidates and finally returns the *k*-nearest neighbors. The search algorithm progressively expands the search space in ℋ_*T*_ to find all *k*-nearest neighbors in the database; hence, the result is equivalent to that of the brute-force search, disregarding a possible difference in the order of ties with the same Hamming distance. The main point here is that dividing and separately indexing the bit vectors dramatically reduces the search space that needs to be explored. For example, when attempting to find all bit vectors within *r*-bit differences from a query using a table, the number of buckets of the table we need to check is 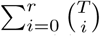 where *T* is the length of the bit vectors and 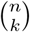.is the number of combinations choosing *k* distinct items from *n*. This value rapidly increases even for a small *r*, which would easily exceed the number of elements in the table. For instance, the total number of combinations for *T*= 128 and *r*= 9 is roughly 20.6 trillion, which is the same order of magnitude as the estimated number of cells in the human body [47]. To avoid the problem, CellAtlasSearch seems to stop searching at some cutoff distance from the query bit vector [15], but choosing a good threshold for each cell is rather difficult. Instead, by dividing a bit vector of length *T* into *m* subvectors of the same length (we assume *T* is divisible by *m* for brevity), when *r < m*, it is possible to find at least one subvector that perfectly matches the corresponding subvector of the query with the pigeonhole principle. This partial matching can be used to find candidate bit vectors quickly using a table data structure. Even when there are no perfect matches in subvectors, the search space greatly shrinks by the division. Our implementation uses a direct-address table to index subvectors, and because the buckets of the table are fairly sparse (i.e., mostly vacant), we devised a data structure, illustrated in Figure 11, in order to exploit the sparsity and CPU caches. In addition, we found that inserting a data prefetch instruction greatly improves the performance of scanning candidates in buckets, because it reduces cache misses when bit vectors do not fit into the CPU caches. Please refer to [19] and our source code for details of the algorithm and the data structure.

**Figure 11:**
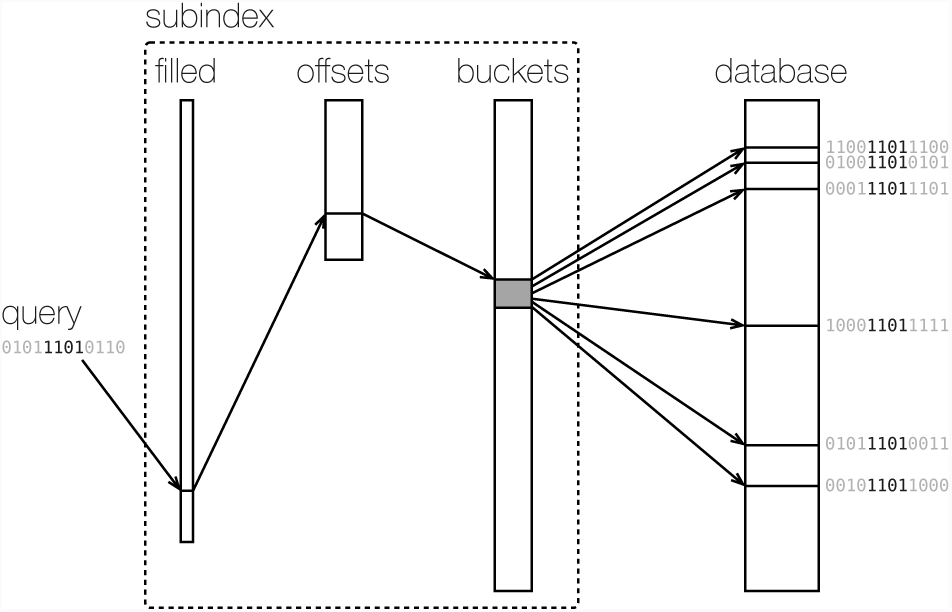
Data structure indexing subvectors. Given a subvector of the query bit vector, the subindex calculates the locations of the bit vectors in the database that contains the subvector in the same position. The subindex consists of three arrays: *filled, offsets*, and *buckets*. The *filled* array is a bit vector of length 2^*s*^, where *s* is the length of indexed subvectors (in this figure, *T*= 12 and *s*= 4), and supports bit counting in a specific range with a constant time, which is used to calculate the location of the *offsets* array for a given subvector (highlighted four bits of the query). The *offsets* and *buckets* arrays are jointly used to obtain the locations of the *database* array at which bit vectors with a given subvector are stored.

The technique of dividing and indexing bit vectors reduces the cost of the search. However, it is still difficult in practice using long bit vectors due to expansion of the search space. To overcome this problem, we use multiple MIH indexes that are independent of each other. We refer to the number of MIH indexes as *L*; thus, the number of bits stored in a database per cell is *TL*. The *L* indexes separately find their own *k*-nearest neighbors of a given query and thus collect *kL* possibly duplicated neighboring candidates for each query These candidates are passed to the next ranking phase. In our method, the two parameters *T* and *L* control the trade-off between accuracy and computational cost of the search.

### Ranking cells

After collecting *kL* candidates from the *L* indexes, CellFishing.jl orders the *kL* cells to return only the top *k* cells. The algorithm computes total Hamming distances from a query and retains the top *k* candidates with the smallest Hamming distances. Candidates with identical distances, if any, are ordered in an arbitrary but deterministic way.

### Similarity estimation

CellFishing.jl can retrieve all the *k*-nearest neighbors of a query cell without any cut-off distance. This is particularly important when the query cell and its nearest neighbors in the database are similar but considerably different due to various factors (e.g., batch effects). However, it also means that CellFishing.jl may retrieve unrelated cells with very low similarity. Using hashed bit vectors, we can estimate similarities between the query and its neighbors by their Hamming distances, but the range of this distance varies depending on the parameters *T* and *L*, and thus is therefore counter-intuitive. Accordingly, CellFishing.jl provides a utility function to estimate the cosine similarity between cells from their Hamming distance. The cosine similarity is normalized between −1 and 1 and is therefore easier to interpret than the Hamming distance.

### Single-cell DEG detection

CellFishing.jl implements a utility function to detect DEGs between two cells (e.g., a query cell and its nearest neighbor in a database), which can be used to evaluate the search outcome in a *post hoc* manner. Here we refer to a query and a reference cells as *u* and *v*, respectively. The DEG detection function first retrieves the *k*-nearest neighbors of *v* from the database and we collectively refer to the set of neighbors as *𝒱*= {*v*_1_, *v*_2_, *…, v_k_*} (note that *v* will be included in *𝒱* as it is also in the same database). Then the raw counts of *v*_1_, *v*_2_, *…, v_k_* are normalized so that their cell-wise total counts are equal to the total count of *u*. The arithmetic mean of the normalized counts for gene *i, λ_i_*, is used as an estimator of the mean parameter of the gene of *u*. Finally, the probability of observing a count *y*_*i*_ or more extremes for each gene *i* is calculated as 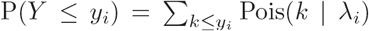 (negative) or 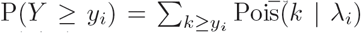 (positive), where Pois(*| λ*) is the probability mass function of the Poisson distribution with mean *λ*. This procedure assumes that all the cells in a local neighborhood, 𝒱, are not biologically different from each other, and therefore, the differences of their normalized counts are due to randomness. The parameter *k* controls the bias-variance tradeoff of this model assumption. In our method, *k* is set to ten by default.

The estimator of *λ*_*i*_ introduced above can be justified as follows. Here, we assume that the UMI count of a gene for *u* and *v* follows *y*_*u*_ *∼* Pois(*β*_*u*_*λ*) and *y*_*v*_ *∼* Pois(*β*_*v*_*λ*), where *β*_*u*_ and *β*_*v*_ denote the capture efficiency of *u* and *v* and *λ* denotes the true gene expression (the concentration of mRNA) of the two cells; the index *i* indicating genes is dropped for brevity. Note that the same expression level *λ* is shared between *u* and *v*, which is the fundamental assumption for DEG detection. We also assume that the ratio of *β*_*u*_ and *β*_*v*_ is equal to the ratio of the total counts (or the library sizes), denoted by *n*_*u*_ and *n*_*v*_. Under these assumptions, we can derive the expectation of *y*_*u*_ from the normalized count of *y*_*v*_ as follows:

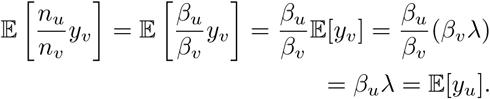

Therefore, the arithmetic mean of the normalized counts of nearby cells is an unbiased estimator of the expectation of *y*_*u*_.

DEG detection functionality is an optional utility of CellFishing.jl because it requires additional disk space to store the raw UMI counts of the database cells. If the database stores the raw counts, the count matrix is compressed by Blosc (http://blosc.org/), a high performance compressor optimized for binary data. CellFishing.jl uses LZ4HC (https://lz4.github.io/lz4/) as the backend compressor of Blosc with the maximum compression level (i.e., level=9) and shuffling, achieving high compression ratio and rapid decompression in our preliminary experiments using actual scRNA-seq data. When loading a database object with a count matrix from a file, CellFishing.jl does not directly load the matrix into memory. Instead, the compressed matrix is mapped to the memory space as a memory-mapped file using the mmap system call on a POSIX-compliant platform or its counterpart in Windows, and essential parts of the matrix are decompressed on request. This has several advantages such as reducing data loading time, avoiding unnecessary memory allocation, and sharing the same data among different processes without duplication.

### Implementation

CellFishing.jl is an open-source package written in the Julia language [18] and is distributed under the MIT License. Julia is a high-performance dynamic programming language for technical computing, which compiles the source code at run-time and makes it easier to install CellFishing.jl, since the user does not need to compile the source code during installation. The entire code of the package is written in Julia, as it makes the code simpler while its performance is closely comparable to other compiled programming languages such as C. The installation can be done using a package manager bundled with Julia. The source code and the documentation of CellFishing.jl are hosted on GitHub: https://github.com/bicycle1885/CellFishing.jl.

The maximum performance of CellFishing.jl is achieved by exploiting the characteristics of modern processors. For example, CellFishing.jl heavily uses the POPCNT instruction (to count the number of 1 bits) and the PREFETCHNTA instruction (to prefetch data into caches from memory) introduced by the Streaming SIMD Extensions to compute the Hamming distance between bit vectors. These instructions are available on most processors manufactured by Intel or AMD. Since Julia compiles the source code at run-time, suitable instructions for a processor are automatically selected. Also, the linear algebra libraries included in Julia, such as OpenBLAS and LAPACK, contribute to the performance of the preprocessing and hashing phases. We consider that using accelerators such as GPUs is not particularly important in CellFishing.jl because these phases do not represent a major bottleneck.

### Reproducibility

The script files used in this study are included in Additional file 3. To ensure reproducibility, all experiments were run using Snakemake [48], a Python-based workflow management tool for bioinformatics. We used Julia 1.0.1 to run CellFishing.jl 0.3.0 and R 3.5.0 to run scmap 1.2.0. R and scmap were installed in a Docker image built on top of Bioconductor’s Docker image (https://hub.docker.com/r/bioconductor/release_base2/, R3.5.0_Bioc3.7) [49], with the Dockerfile included in the additional file. All the plots and tables in this manuscript were generated in a Jupyter notebook [50], which is also included in the same additional file. We used Linux machines with Intel Xeon Gold 6126 CPU (629.4 GiB of memory, hard disk drive) or Intel Xeon CPU E5-2637 v4 (251.6 GiB of memory, hard disk drive) to benchmark the run-time performance. Performance comparisons between CellFishing.jl and scmap-cell were performed on the same machine.

### Ethics approval and consent to participate

Not applicable.

### Consent for publication

Not applicable.

### Availability of data and materials

CellFishing.jl is implemented in the Julia programming language, and the source code is freely available under the MIT license at https://github.com/bicycle1885/CellFishing.jl (DOI: https://doi.org/10.5281/zenodo.1495440). All analyses and figures in this paper can be reproduced using the scripts in Additional file 3. The Baron2016 and Shekhar2016 data sets were downloaded from the Gene Expression Omnibus with accession numbers GSE84133 and GSE81904, respectively. The list of homologous genes between human and mouse was downloaded from the Vertebrate Homology database of Mouse Genome Informatics at http://www.informatics.jax.org/homology.shtml. The cluster annotation file of Shekhar2016 was downloaded from https://portals.broadinstitute.org/single_cell/study/retinal-bipolar-neuron-drop-seq. The Plass2018 data set was downloaded from https://shiny.mdc-berlin.de/psca/. The TabulaMuris data set was downloaded from https://figshare.com/articles/Single-cell_RNA-seq_data_from_Smart-seq2_sequencing_of_FACS_sorted_cells_v2_/5829687 and https://figshare.com/articles/Single-cell_RNA-seq_data_from_microfluidic_emulsion_v2_/5968960. The 1M neurons data set was downloaded from https://support.10xgenomics.com/single-cell-gene-expression/datasets/1.3.0/1M_neurons.

## Competing interests

The authors declare that they have no competing interests.

## Funding

This work was supported by MEXT KAKENHI Grant Number 16K16152. This work was supported by the Projects for Technological Development, Research Center Network for Realization of Regenerative Medicine by Japan (18bm0404024h0001), the Japan Agency for Medical Research and Development (AMED), and JST CREST grant number JPMJCR16G3, Japan to I.N.

## Author’s contributions

KST, KT, KSH and IN designed the study. KST implemented all the software. KT validated all the R scripts. KST, KT, and IN wrote the manuscript. All authors have read and approved the final manuscript.

## Supporting information

## Acknowledgements

We thank Hirotaka Matsumoto for helpful discussions. We also thank Mr. Akihiro Matsushima and Mr. Manabu Ishii for their assistance with the IT infrastructure for the data analysis. We are also grateful to all members of the Laboratory for Bioinformatics Research, RIKEN Center for Biosystems Dynamics Research for giving us helpful advice.

