## Supplementary material for "CellFishing.jl: an ultrafast and scalable cell search method for single-cell RNA-sequencing"

### Additional and high-resolution figures of "CellFishing.jl: an ultrafast and scalable cell search method for single-cell RNA-sequencing"

Kenta Sato Koki Tsuyuzaki Kentaro Shimizu
Itoshi Nikaido

November 27, 2018

This file lists all the figures embedded in Analysis.html with higher resolution.

14 Distributions of estimation errors for angles (TabulaMuris, Chromium) 17
15 Scatter plots of the Hamming distance versus the cosine distance (Baron2016, human) . . . . . . . . . . . . . . . . . . . . . . . . . 18
16 Scatter plots of the Hamming distance versus the cosine distance (Shekhar2016) . . . . . . . . . . . . . . . . . . . . . . . . . . . . 19
17 Scatter plots of the Hamming distance versus the cosine distance (Plass2018) . . . . . . . . . . . . . . . . . . . . . . . . . . . . . 20
18 Scatter plots of the Hamming distance versus the cosine distance (TabulaMuris, Chromium) . . . . . . . . . . . . . . . . . . . . . . 21
19 Two-dimensional embedding of expression profiles with UMAP (Shekhar2016, query batch = 1, Euclidean distance) . . . . . . . 23

20 Two-dimensional embedding of expression profiles with UMAP (Shekhar2016, query batch = 1, cosine distance) . . . . . . . . . 24
21 Two-dimensional embedding of expression profiles with UMAP (Shekhar2016, query batch = 2, Euclidean distance) . . . . . . . 25
22 Two-dimensional embedding of expression profiles with UMAP (Shekhar2016, query batch = 2, cosine distance) . . . . . . . . . 26
23 Two-dimensional embedding of expression profiles with UMAP (Shekhar2016, query batch = 1) . . . . . . . . . . . . . . . . . . . 27
24 Two-dimensional embedding of expression profiles with UMAP (Shekhar2016, query batch = 2) . . . . . . . . . . . . . . . . . . . 28
25 Comparison of consistency scores . . . . . . . . . . . . . . . . . . . 29
26 Comparison of Cohen's kappa scores . . . . . . . . . . . . . . . . . 30
27 Comparison of index times . . . . . . . . . . . . . . . . . . . . . . 31
28 Comparison of query times . . . . . . . . . . . . . . . . . . . . . . 32
29 Proportions of cluster assignments (Baron2016, human, CellFishing) . . . . . . . . . . . . . . . . . . . . . . . . . . . . . . . . . 34
30 Proportions of cluster assignments (Baron2016, human, scmap-cell) 35
31 Cluster-specific consistency scores (Baron2016, human) . . . . . . 36
32 Proportions of cluster assignments (Shekhar2016, CellFishing) . . 37
33 Proportions of cluster assignments (Shekhar2016, scmap-cell) . . 38
34 Cluster-specific consistency scores (Shekhar2016) . . . . . . . . . 39
35 Proportions of cluster assignments (Plass2018, CellFishing) . . . 40
36 Proportions of cluster assignments (Plass2018, scmap-cell) . . . . 41
37 Cluster-specific consistency scores (Plass2018) . . . . . . . . . . . 42
38 Proportions of cluster assignments (TabulaMuris, Chromium, CellFishing) . . . . . . . . . . . . . . . . . . . . . . . . . . . . . . . 43
39 Proportions of cluster assignments (TabulaMuris, Chromium, scmap-cell) . . . . . . . . . . . . . . . . . . . . . . . . . . . . . . . . 44
40 Cluster-specific consistency scores (TabulaMuris, Chromium) . . 45
41 Distributions of similarities with or without a specific cell type (Shekhar2016) . . . . . . . . . . . . . . . . . . . . . . . . . . . . 46
42 Distributions of similarities with or without a specific cell type (TabulaMuris, Chromium) . . . . . . . . . . . . . . . . . . . . . . 47
43 Cell mapping across batches (consistency) . . . . . . . . . . . . . 48
44 Cell mapping across batches (Cohen's kappa) . . . . . . . . . . . 49
45 Distribution of cluster sizes (Shekhar2016) . . . . . . . . . . . . . 50
46 Distribution of cluster sizes (Plass2018) . . . . . . . . . . . . . . 51
47 Cluster assignments across different protocol (TabulaMuris, Chromium, CellFishing, All) . . . . . . . . . . . . . . . . . . . . . . . . 53
48 Cluster assignments across different protocol (TabulaMuris, Chromium, CellFishing, Overlapping cell types) . . . . . . . . . . . . . . . . 54
49 Cluster assignments across different protocol (TabulaMuris, Chromium, scmap-cell, All) . . . . . . . . . . . . . . . . . . . . . . . . 55
50 Cluster assignments across different protocol (TabulaMuris, Chromium, scmap-cell, Overlapping cell types) . . . . . . . . . . . . . . . . . 56
51 Scalability (1M_neurons) . . . . . . . . . . . . . . . . . . . . . . 57

52 Computational costs of different phases (1M_neurons) . . . . . . 58

### Cell-type distributions

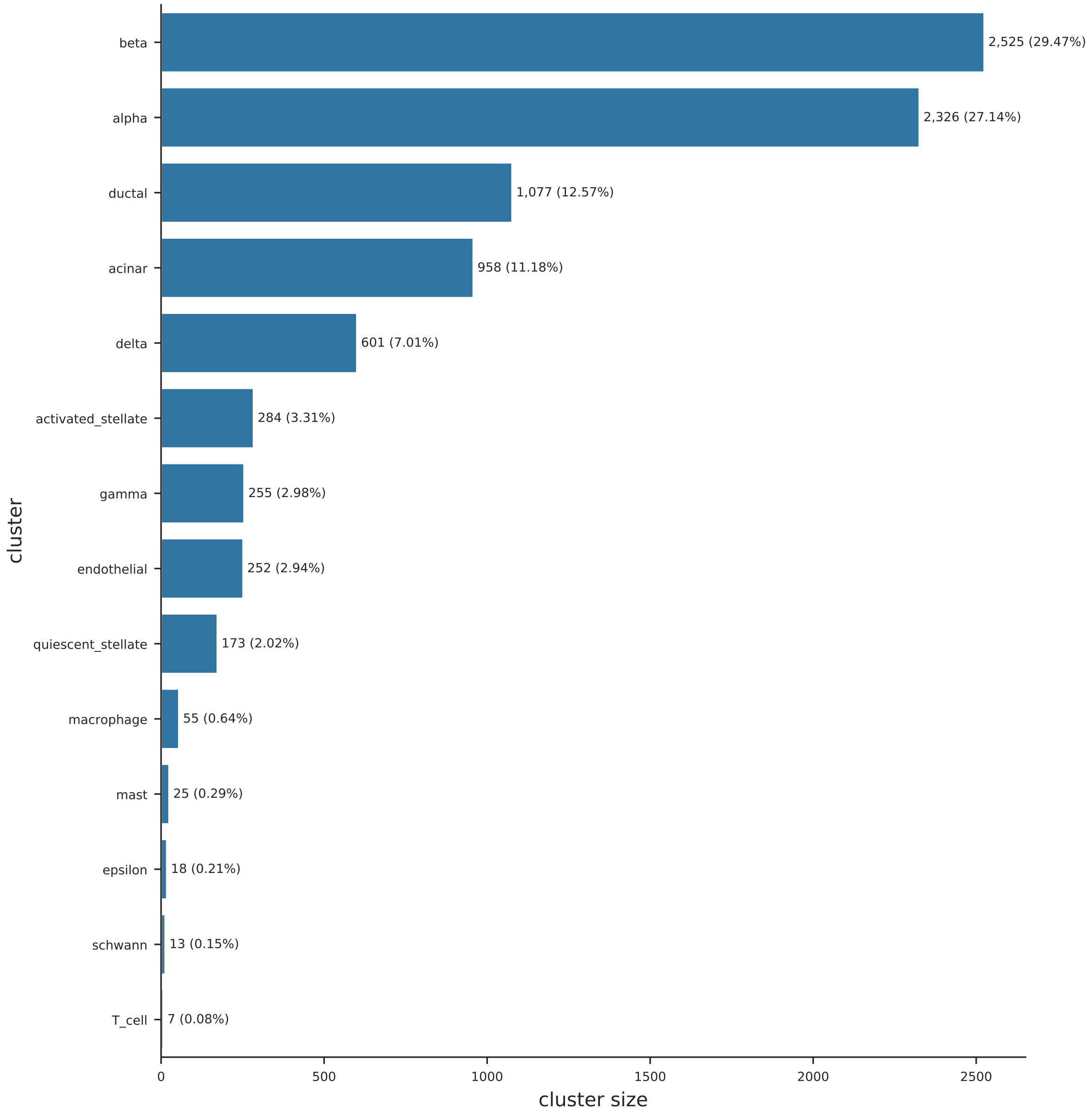

Figure 1: Distribution of cell types (Baron2016, human).

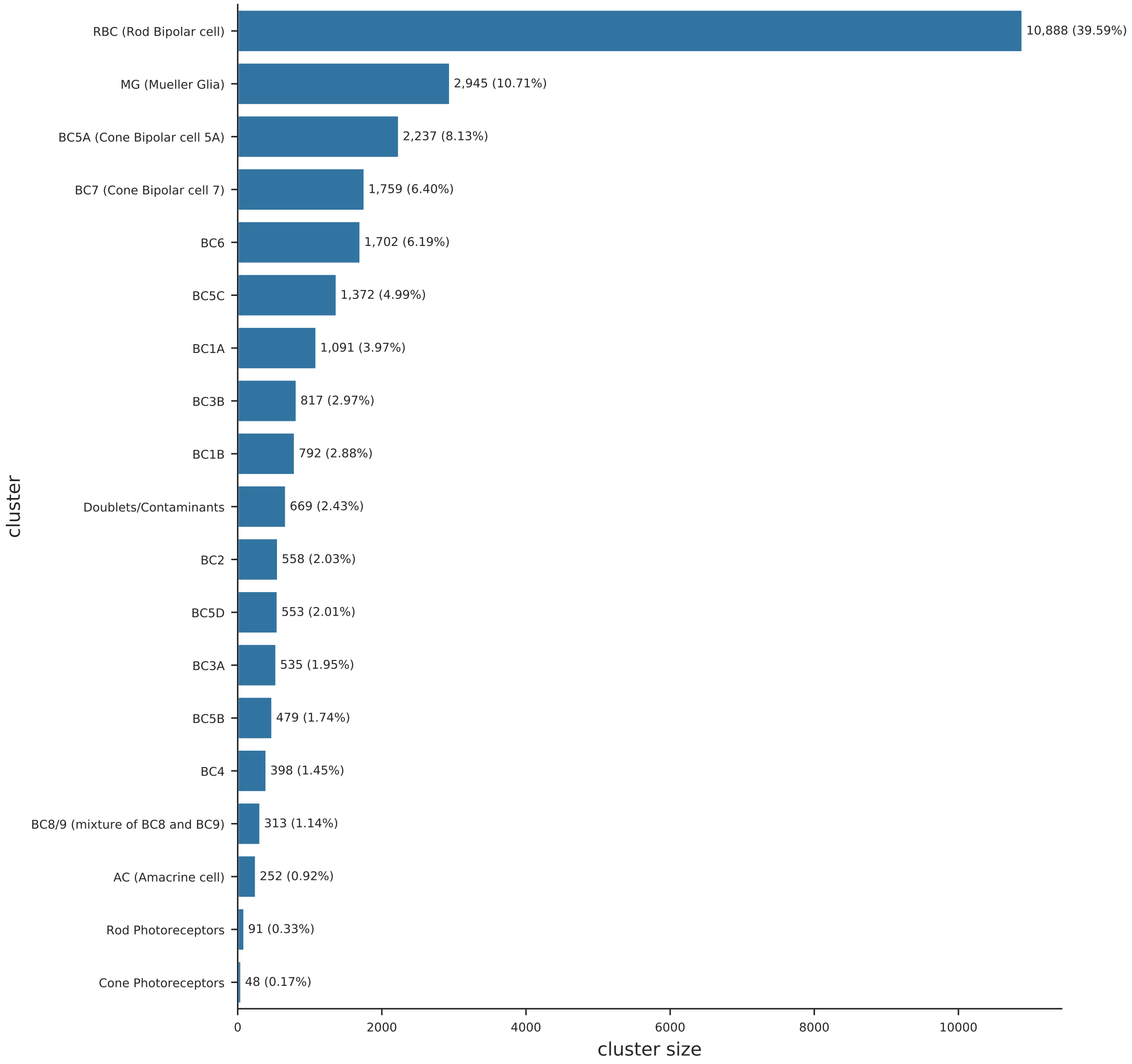

Figure 2: Distribution of cell types (Shekhar2016).

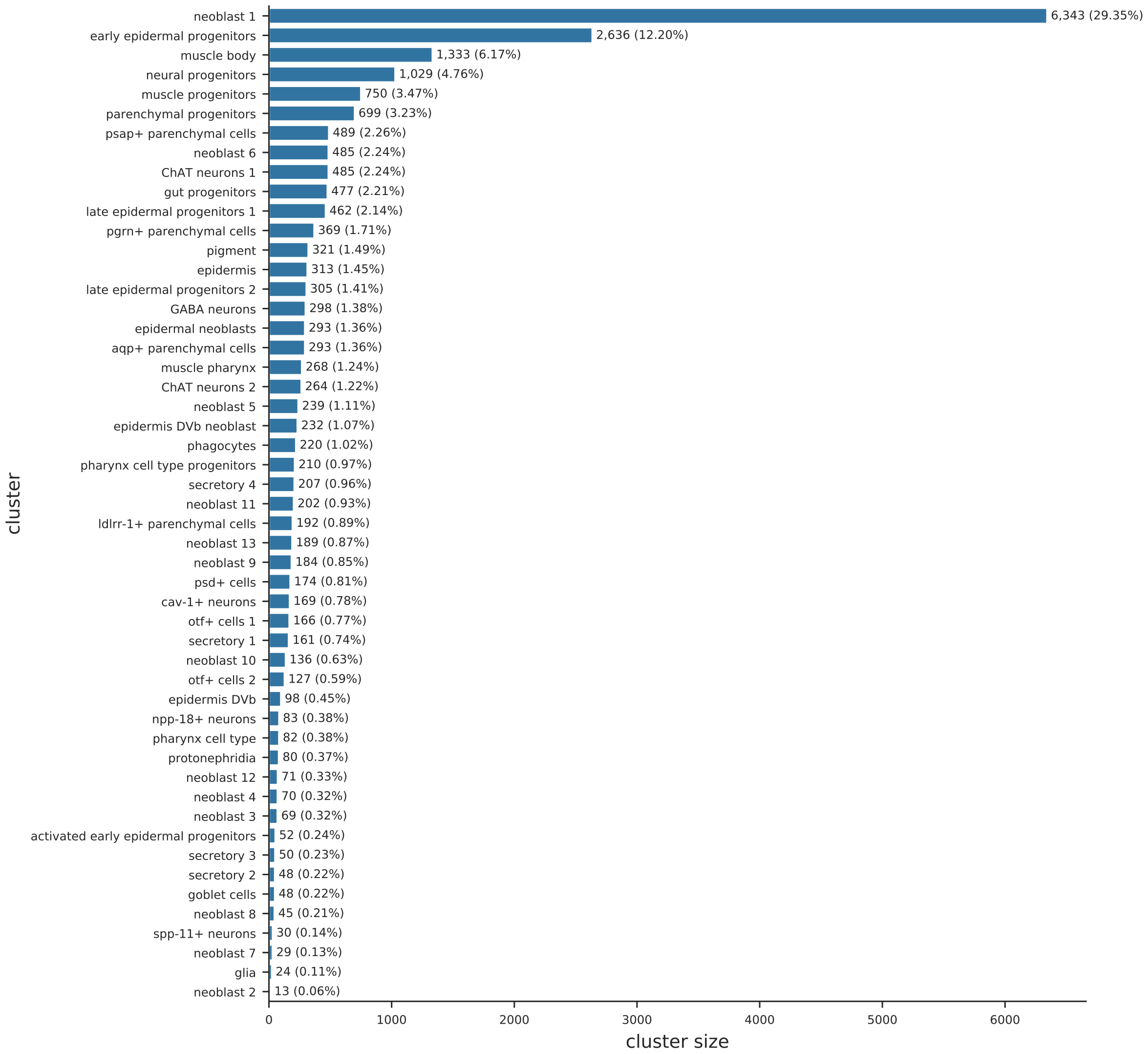

Figure 3: Distribution of cell types (Plass2018).

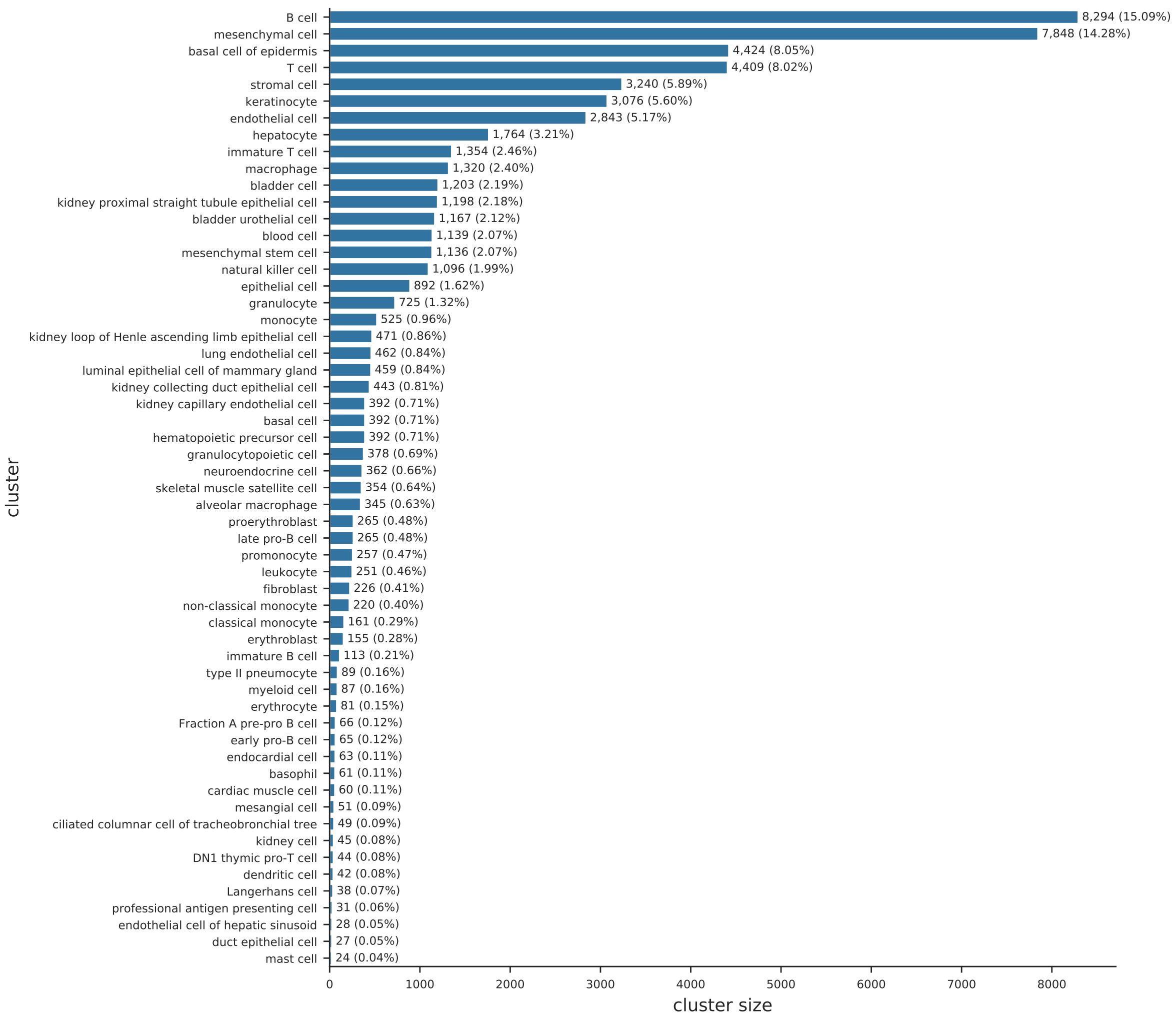

Figure 4: Distribution of cell types (TabulaMuris, Chromium).

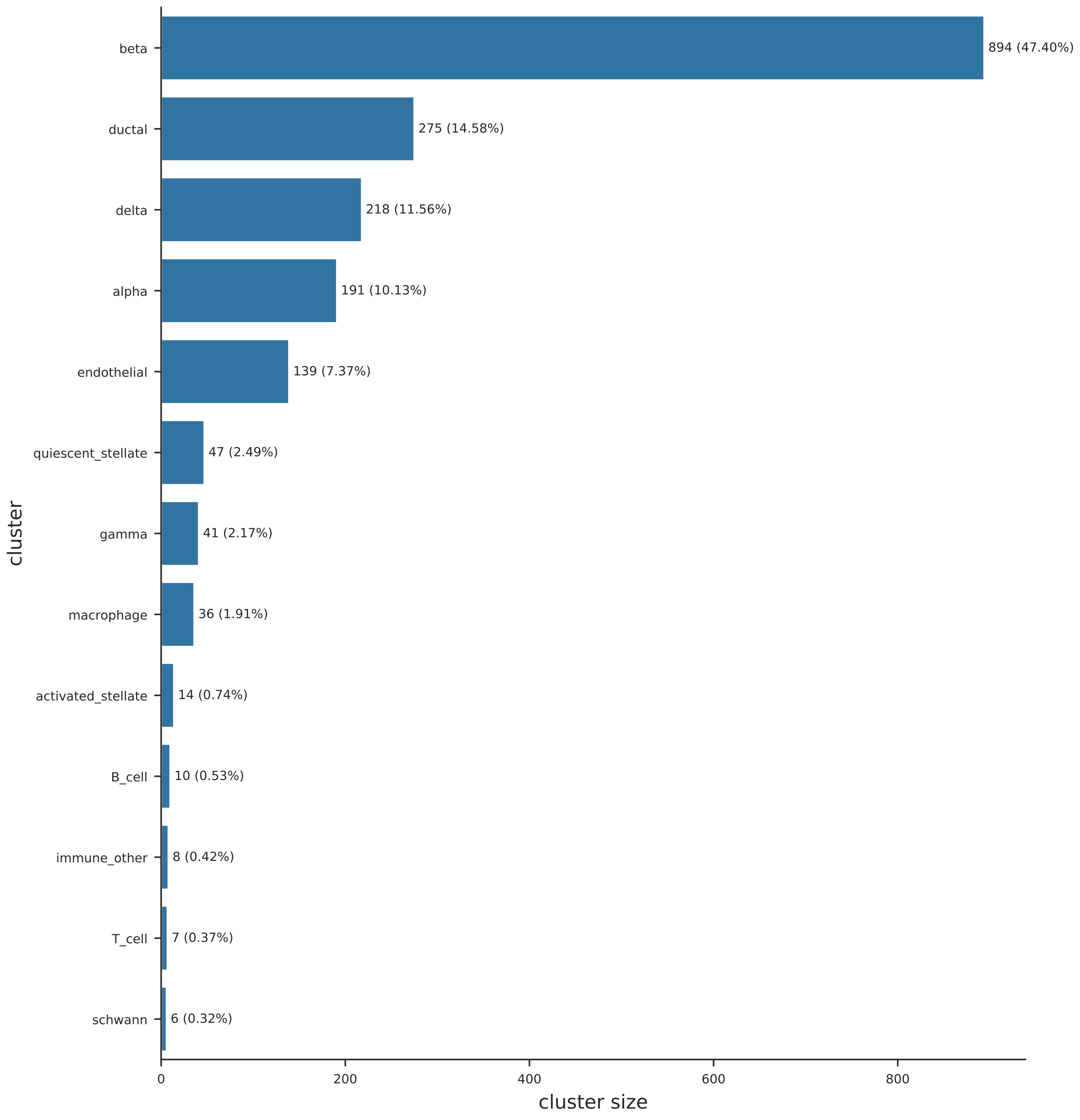

Figure 5: Distribution of cell types (Baron2016, mouse).

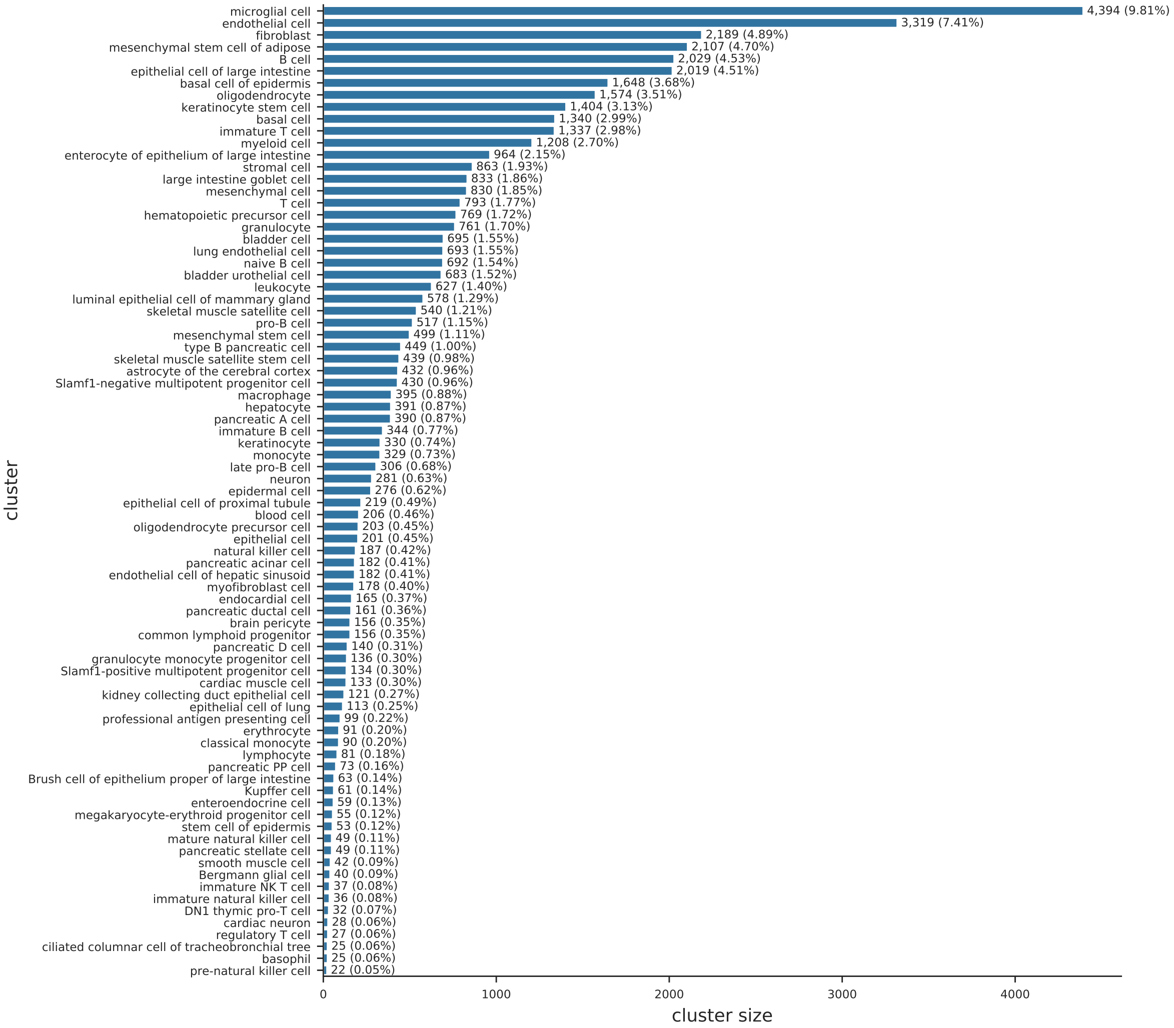

Figure 6: Distribution of cell types (TabulaMuris, Smart-Seq2).

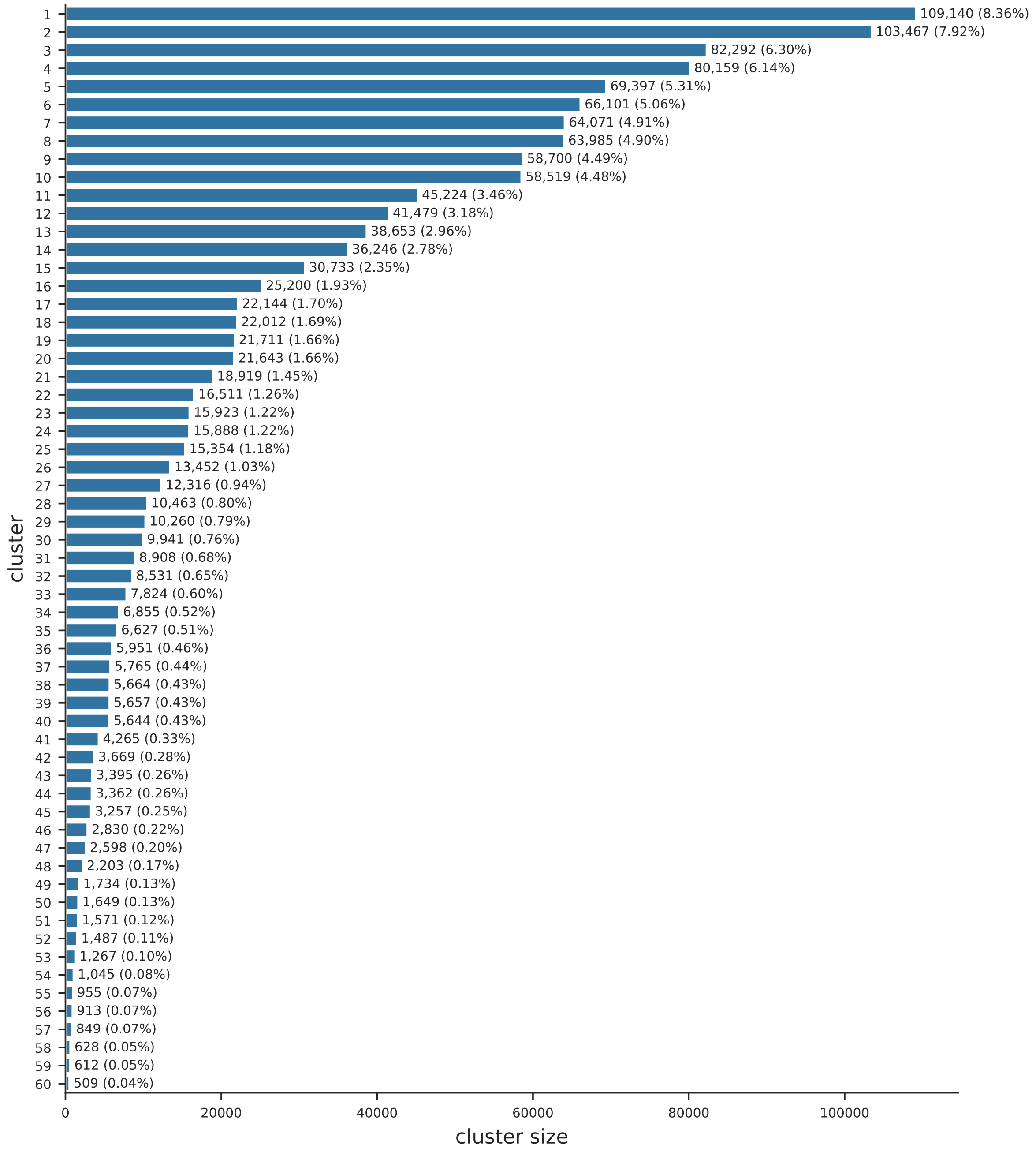

Figure 7: Distribution of cell types (1M_neurons).

### Randomized SVD

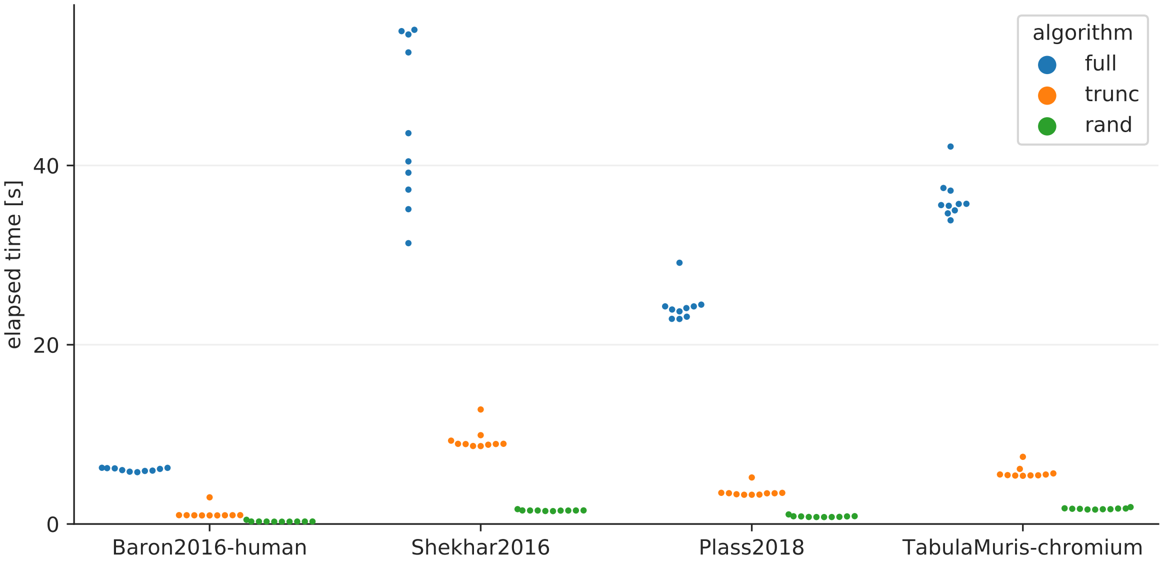

Figure 8: Elapsed time of different SVD algorithms. Please refer to the main manuscript for the detailed caption.

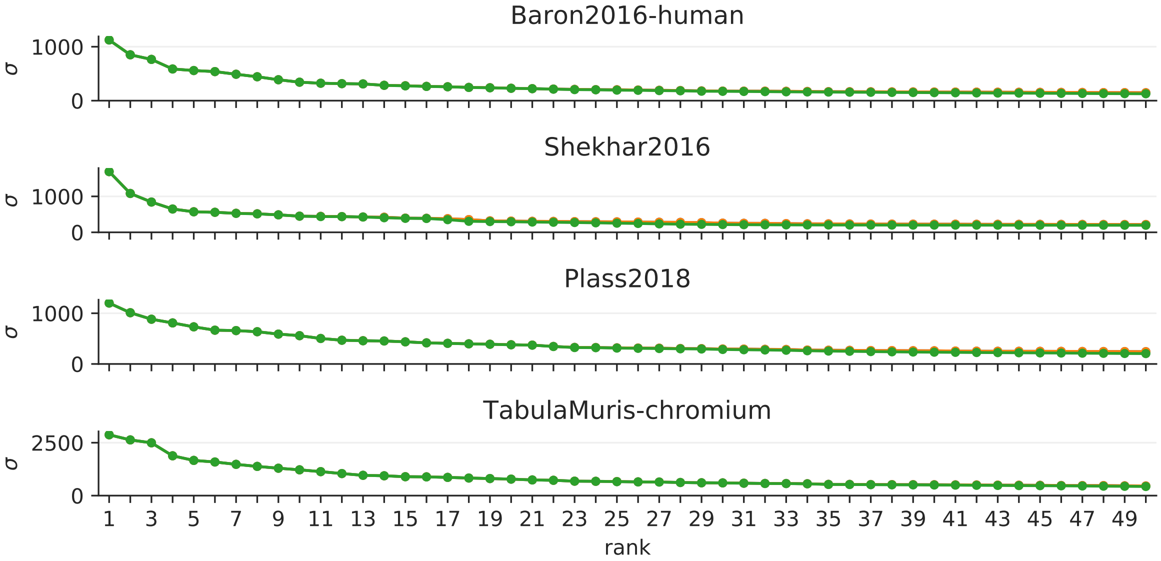

Figure 9: Singular values of the randomized SVD. Please refer to the main manuscript for the detailed caption.

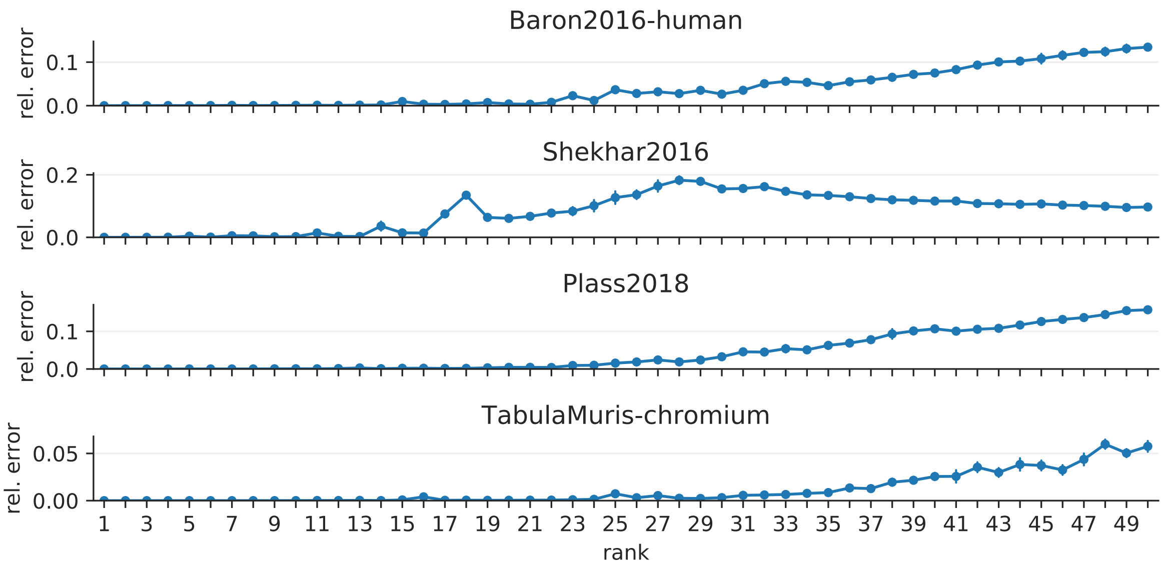

Figure 10: Relative errors of the randomized SVD. Please refer to the main manuscript for the detailed caption.

#### Similarity estimation via hashing

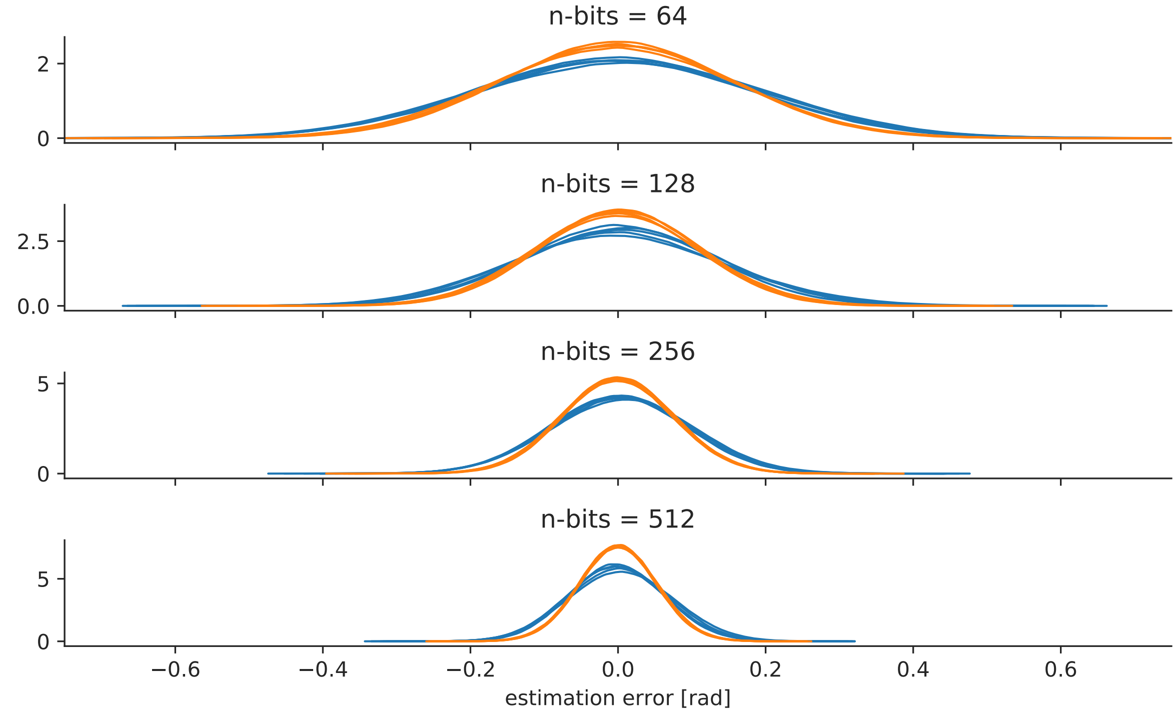

Figure 11: Distributions of estimation errors for angles (Baron2016, human). Please refer to the main manuscript for the detailed caption.

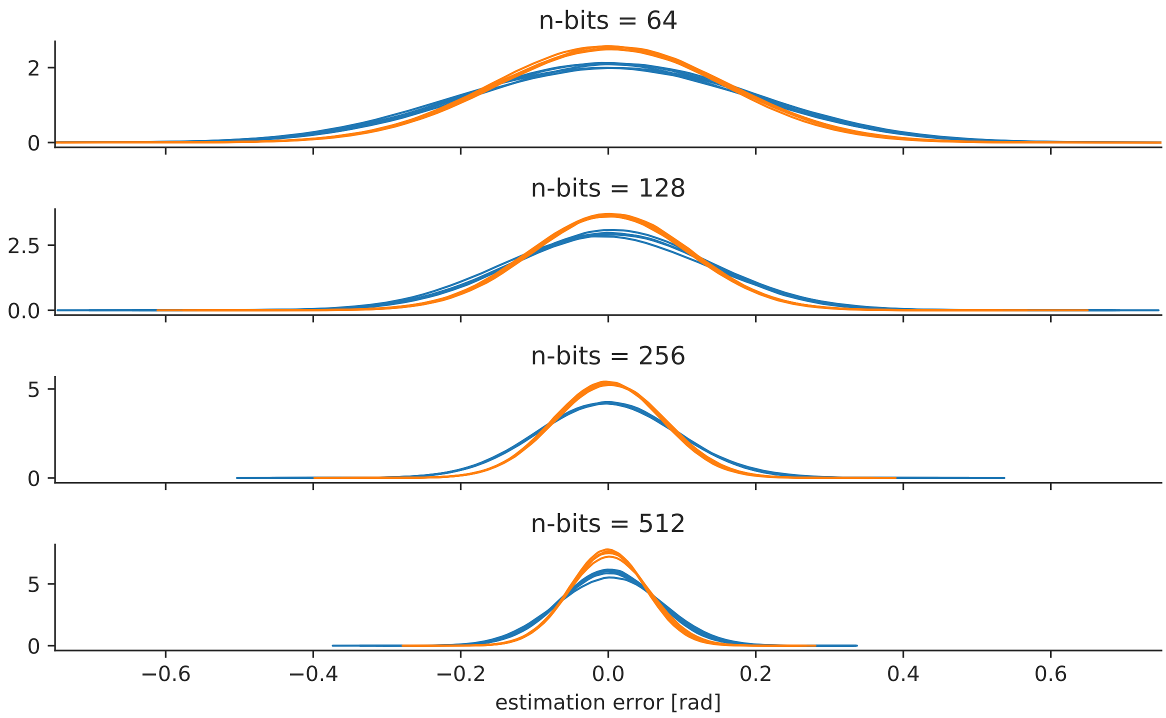

Figure 12: Distributions of estimation errors for angles (Shekhar2016). Please refer to the main manuscript for the detailed caption.

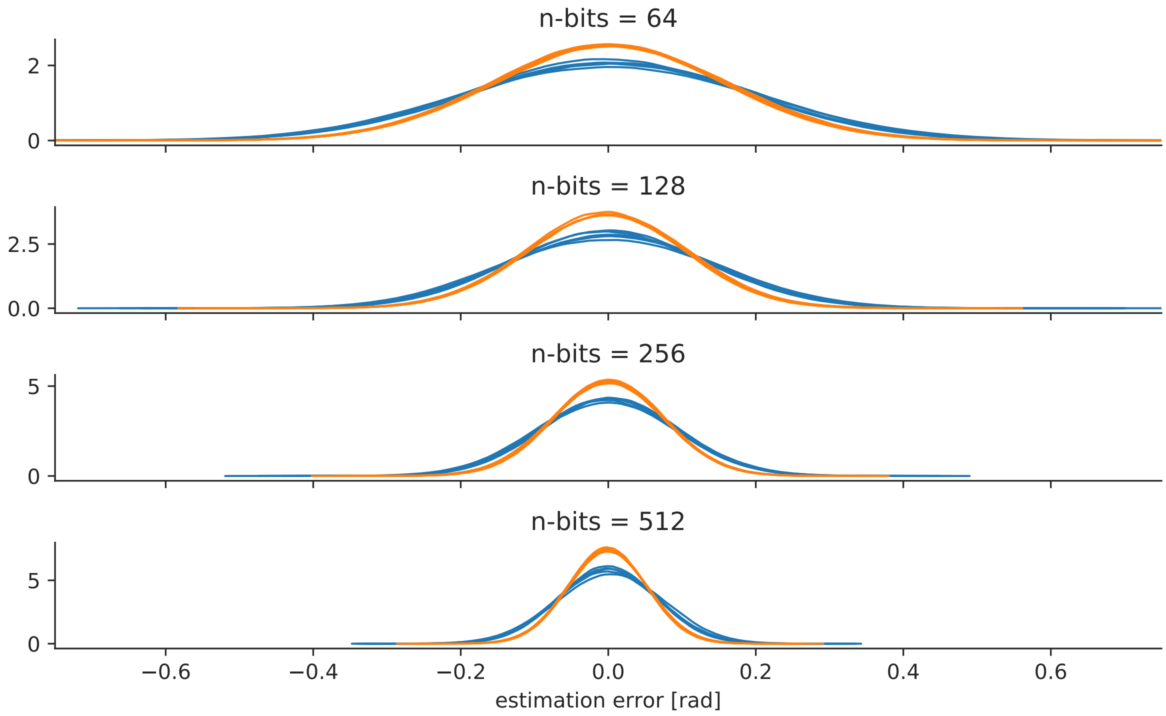

Figure 13: Distributions of estimation errors for angles (Plass2018). Please refer to the main manuscript for the detailed caption.

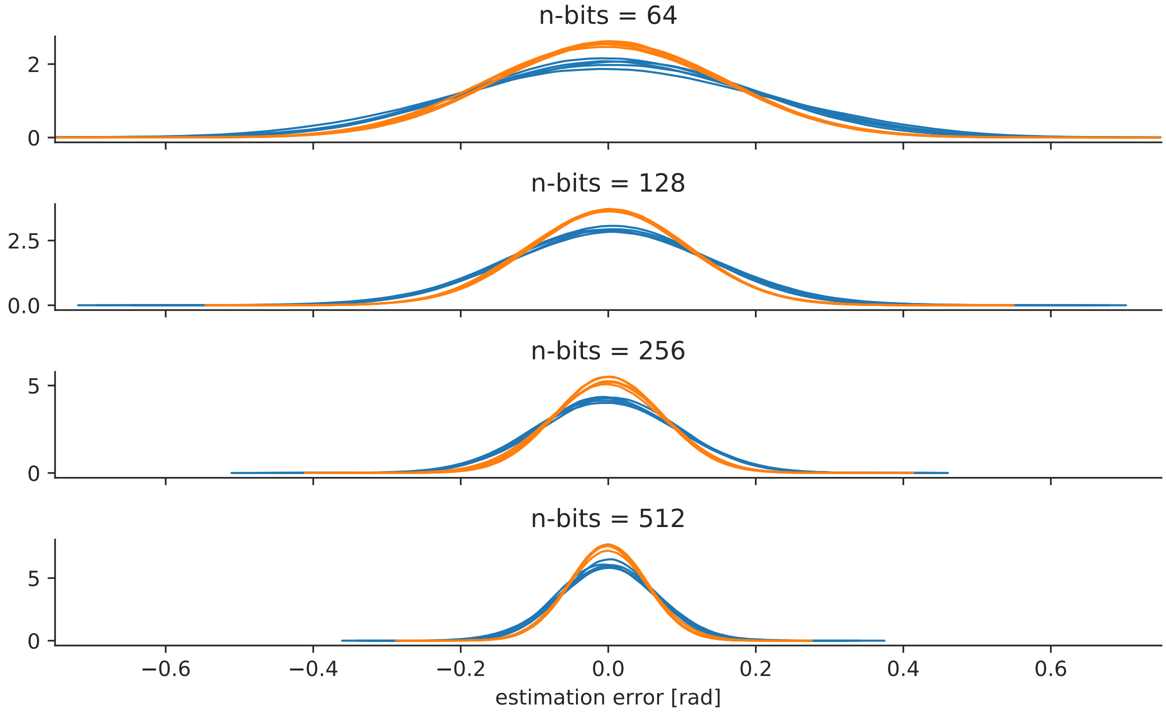

Figure 14: Distributions of estimation errors for angles (TabulaMuris, Chromium). Please refer to the main manuscript for the detailed caption.

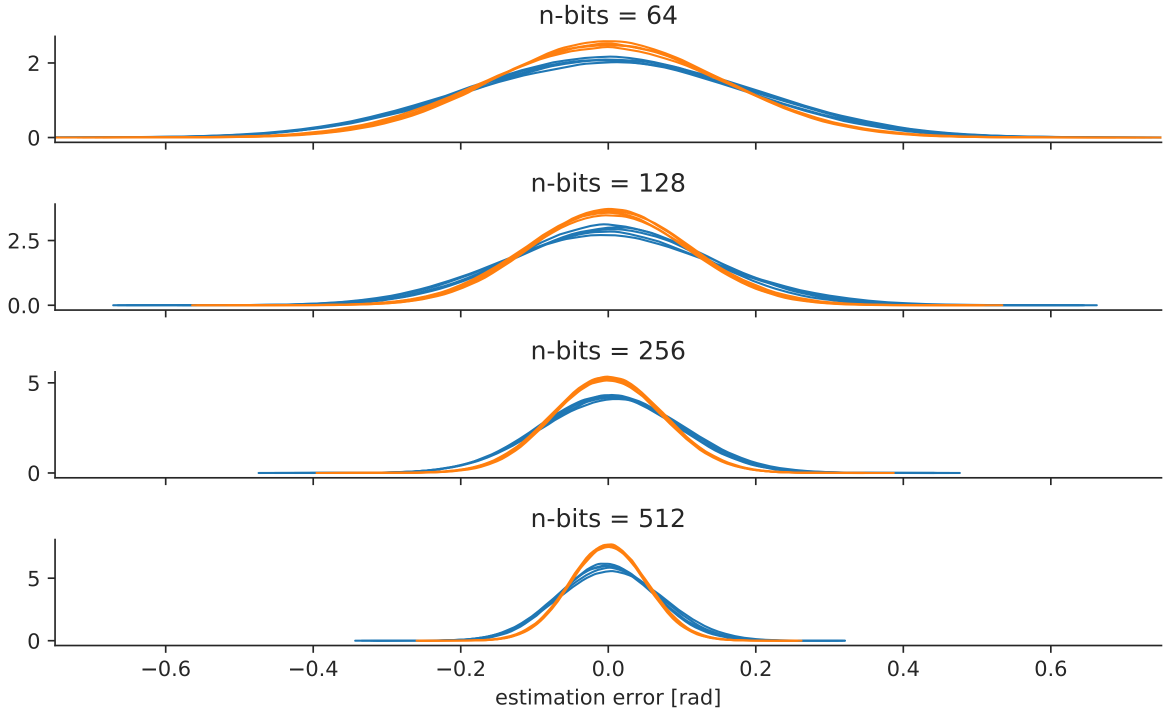

Figure 15: Scatter plots of the Hamming distance versus the cosine distance (Baron2016, human). Please refer to the main manuscript for the detailed caption.

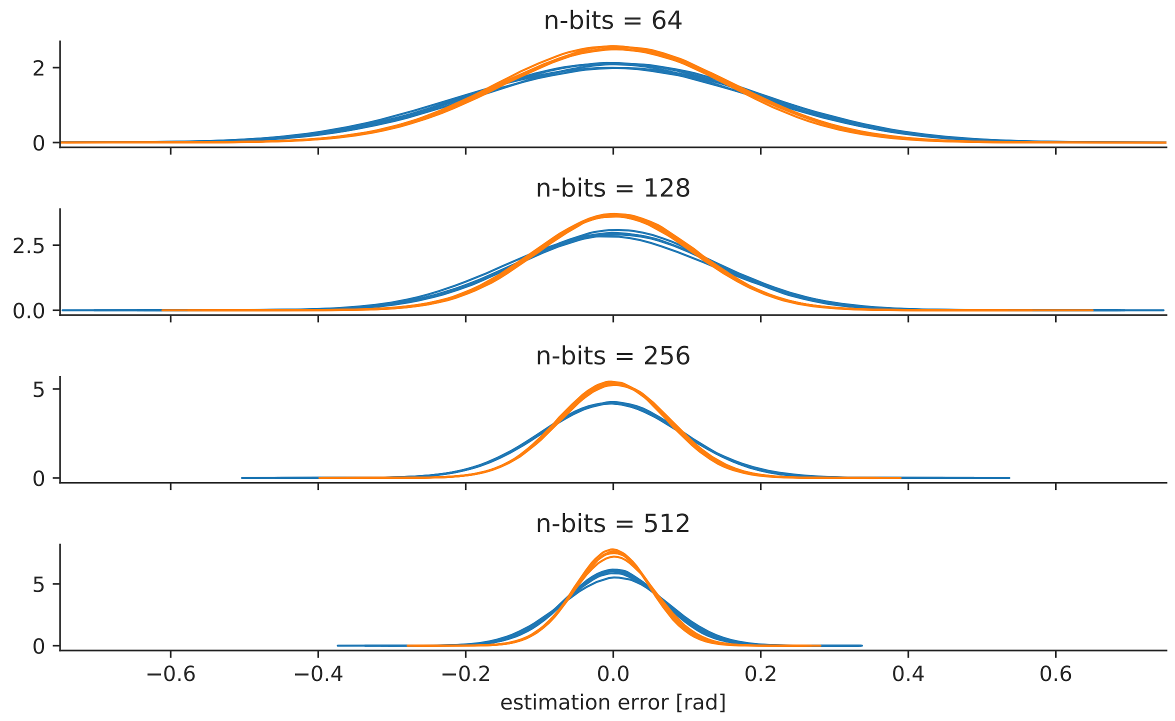

Figure 16: Scatter plots of the Hamming distance versus the cosine distance (Shekhar2016). Please refer to the main manuscript for the detailed caption.

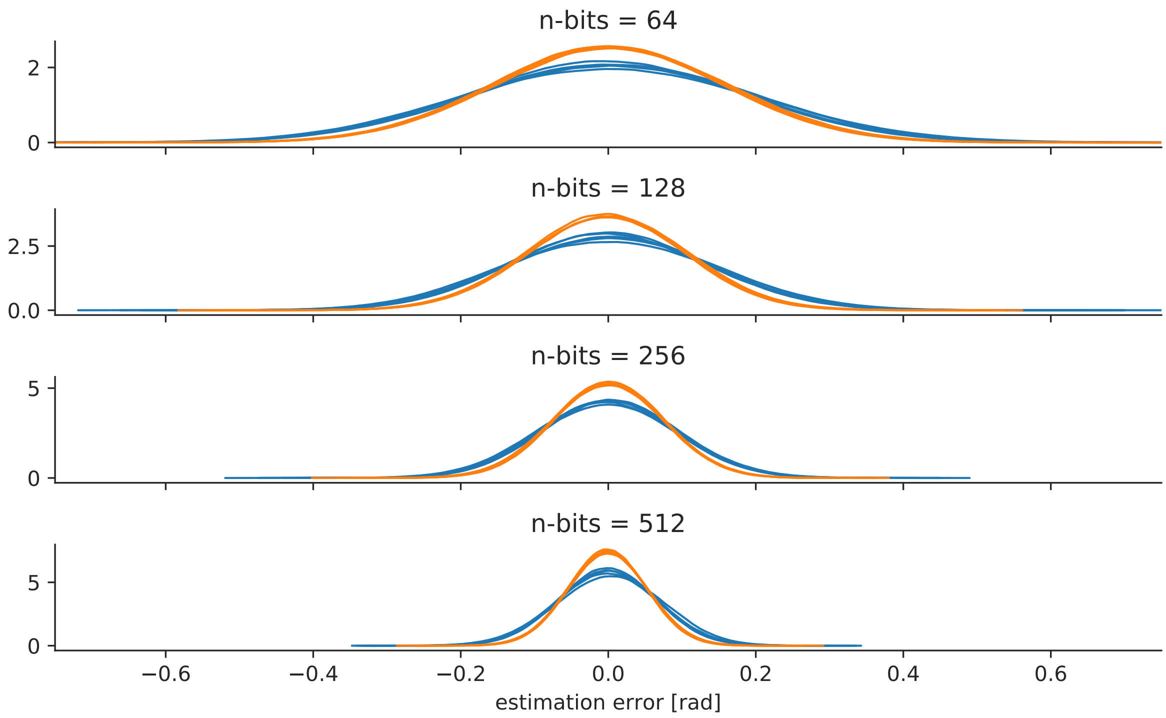

Figure 17: Scatter plots of the Hamming distance versus the cosine distance (Plass2018). Please refer to the main manuscript for the detailed caption.

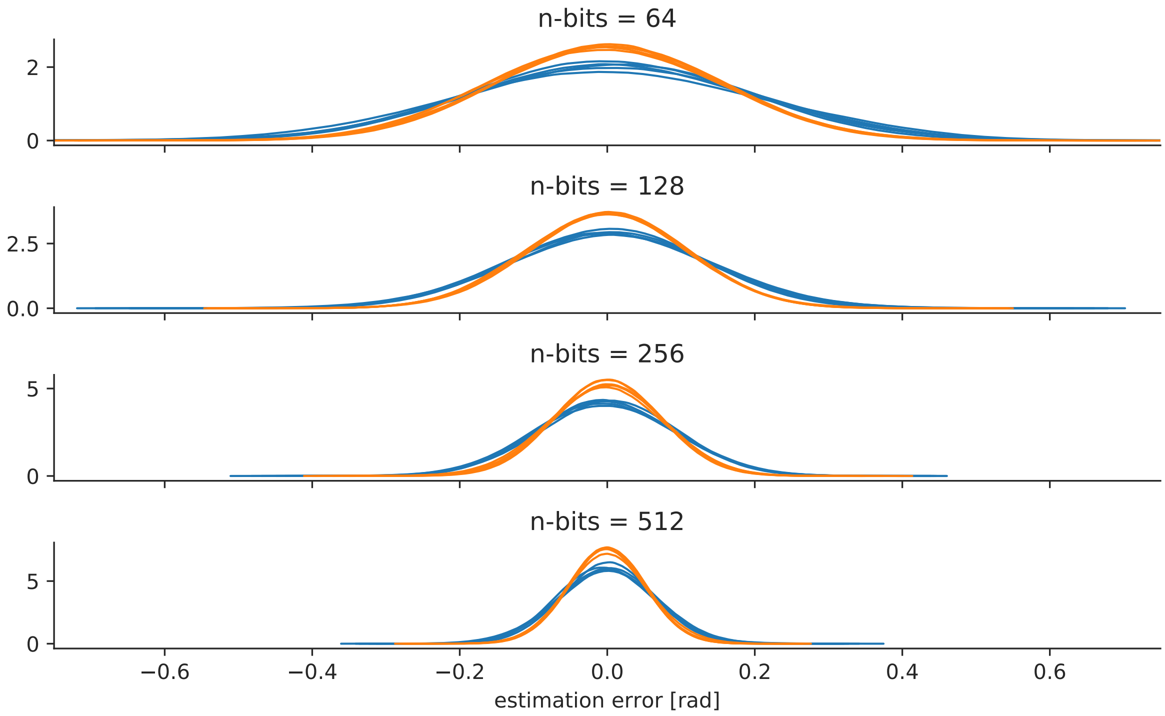

Figure 18: Scatter plots of the Hamming distance versus the cosine distance (TabulaMuris, Chromium). Please refer to the main manuscript for the detailed caption.

#### Visualization of expression profiles

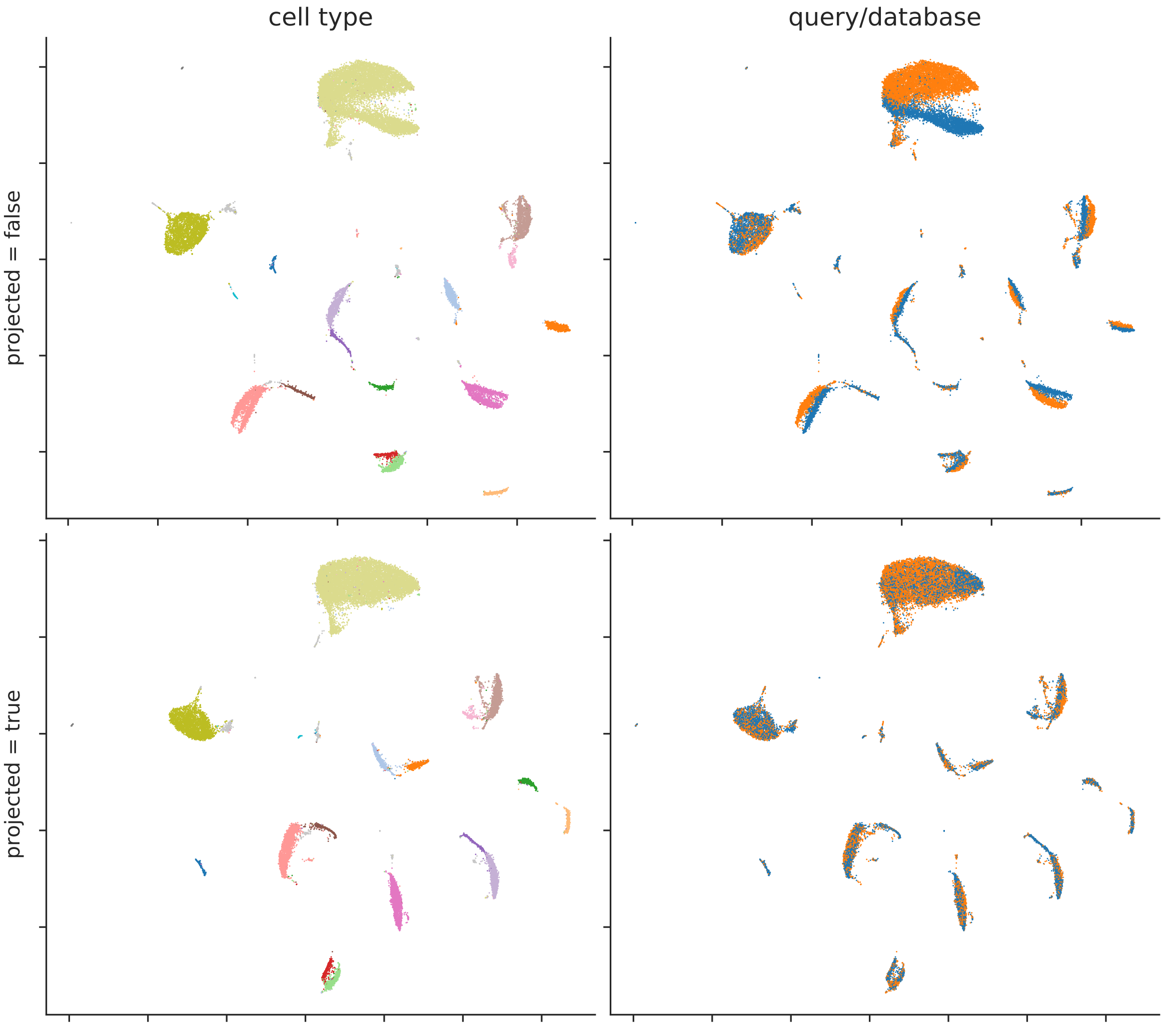

Figure 19: Two-dimensional embedding of expression profiles with UMAP (Shekhar2016, query batch = 1, Euclidean distance). In the left and right panels, points (cells) are colored by cell types and batches, respectively. In the upper panels, the query and database cells are mixed before dimensionality reduction. In the lower panels, the query cells are projected onto the space derived from the database cells.

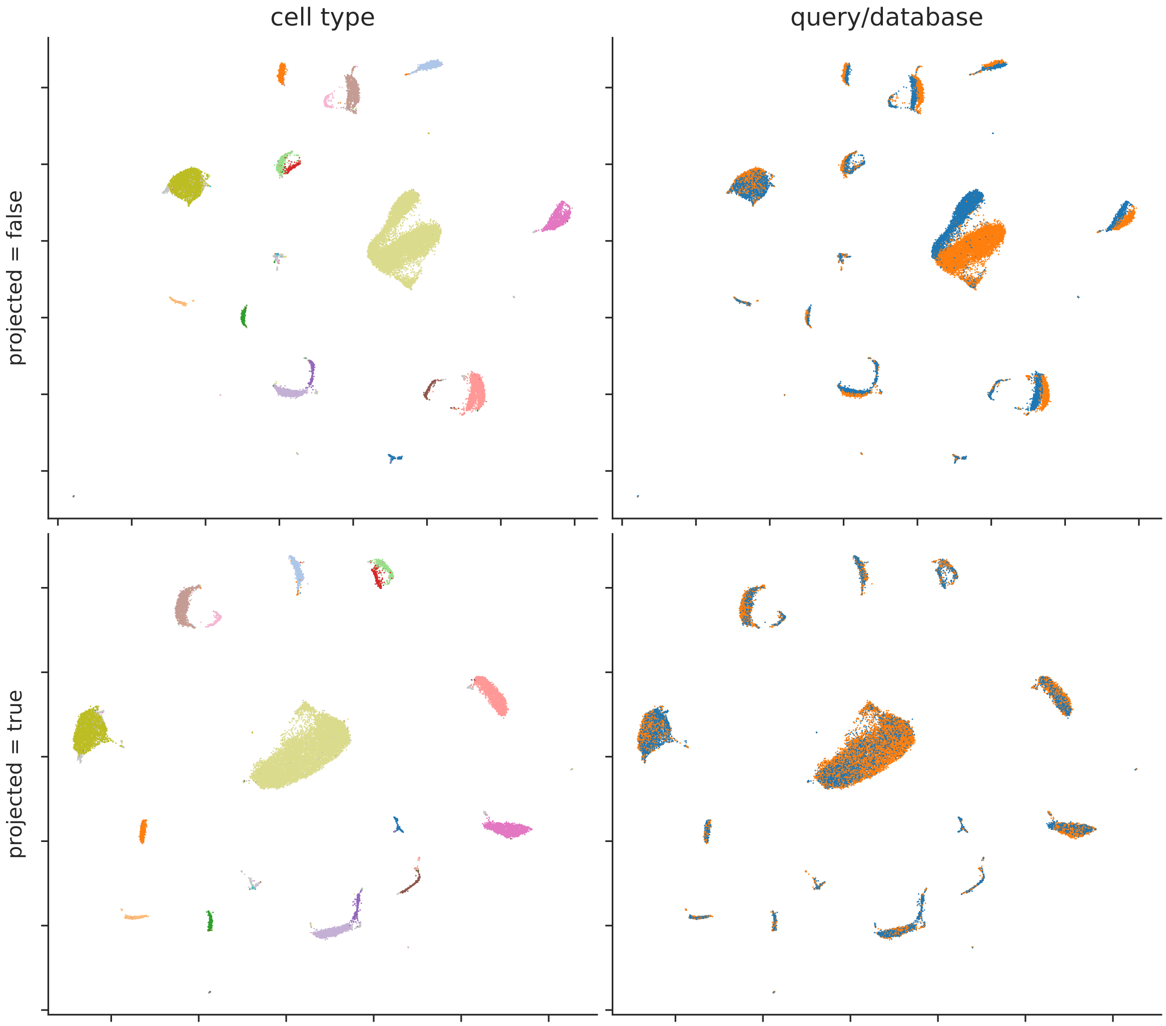

Figure 20: Two-dimensional embedding of expression profiles with UMAP (Shekhar2016, query batch = 1, cosine distance). In the left and right panels, points (cells) are colored by cell types and batches, respectively. In the upper panels, the query and database cells are mixed before dimensionality reduction. In the lower panels, the query cells are projected onto the space derived from the database cells.

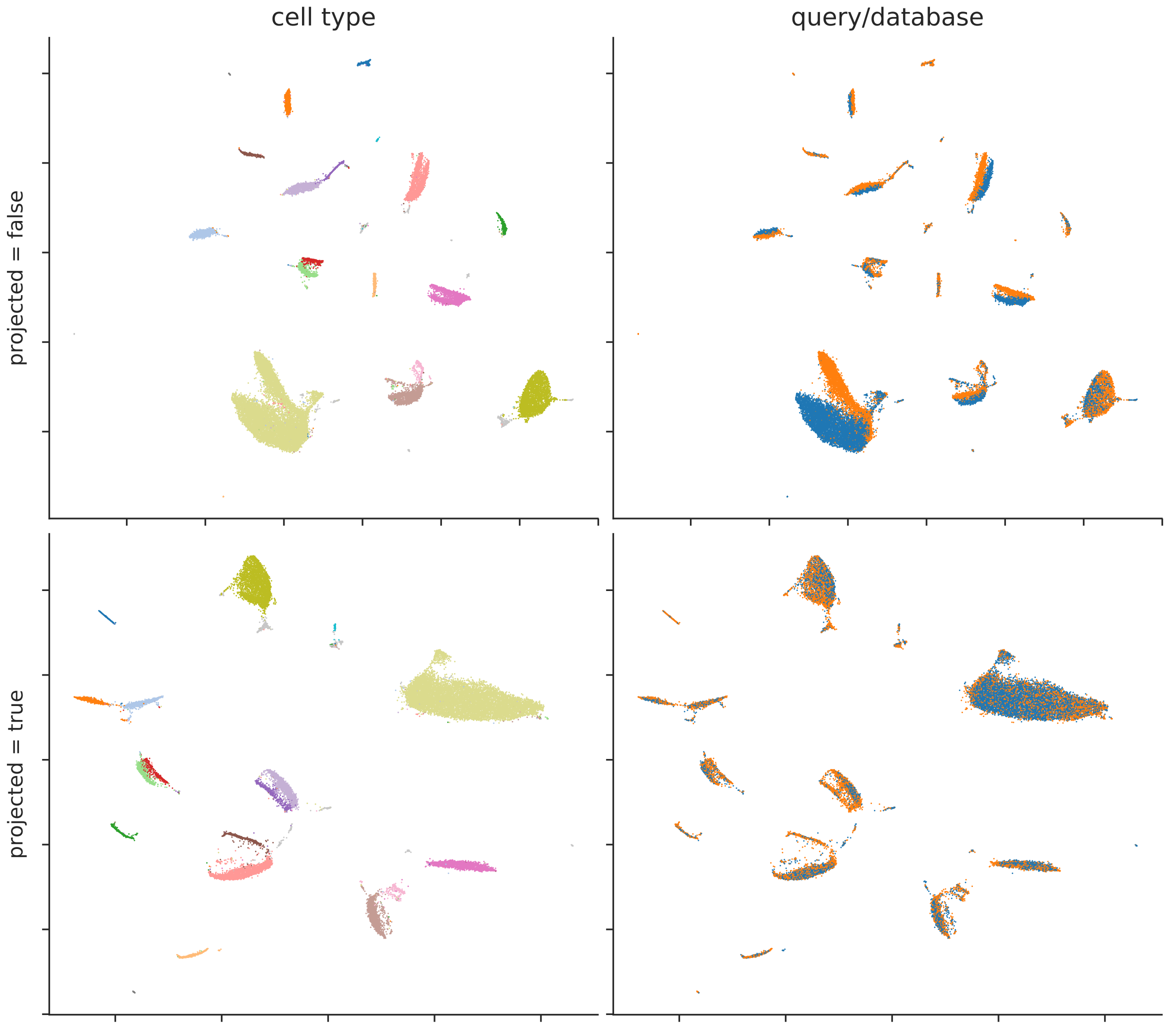

Figure 21: Two-dimensional embedding of expression profiles with UMAP (Shekhar2016, query batch = 2, Euclidean distance). In the left and right panels, points (cells) are colored by cell types and batches, respectively. In the upper panels, the query and database cells are mixed before dimensionality reduction. In the lower panels, the query cells are projected onto the space derived from the database cells.

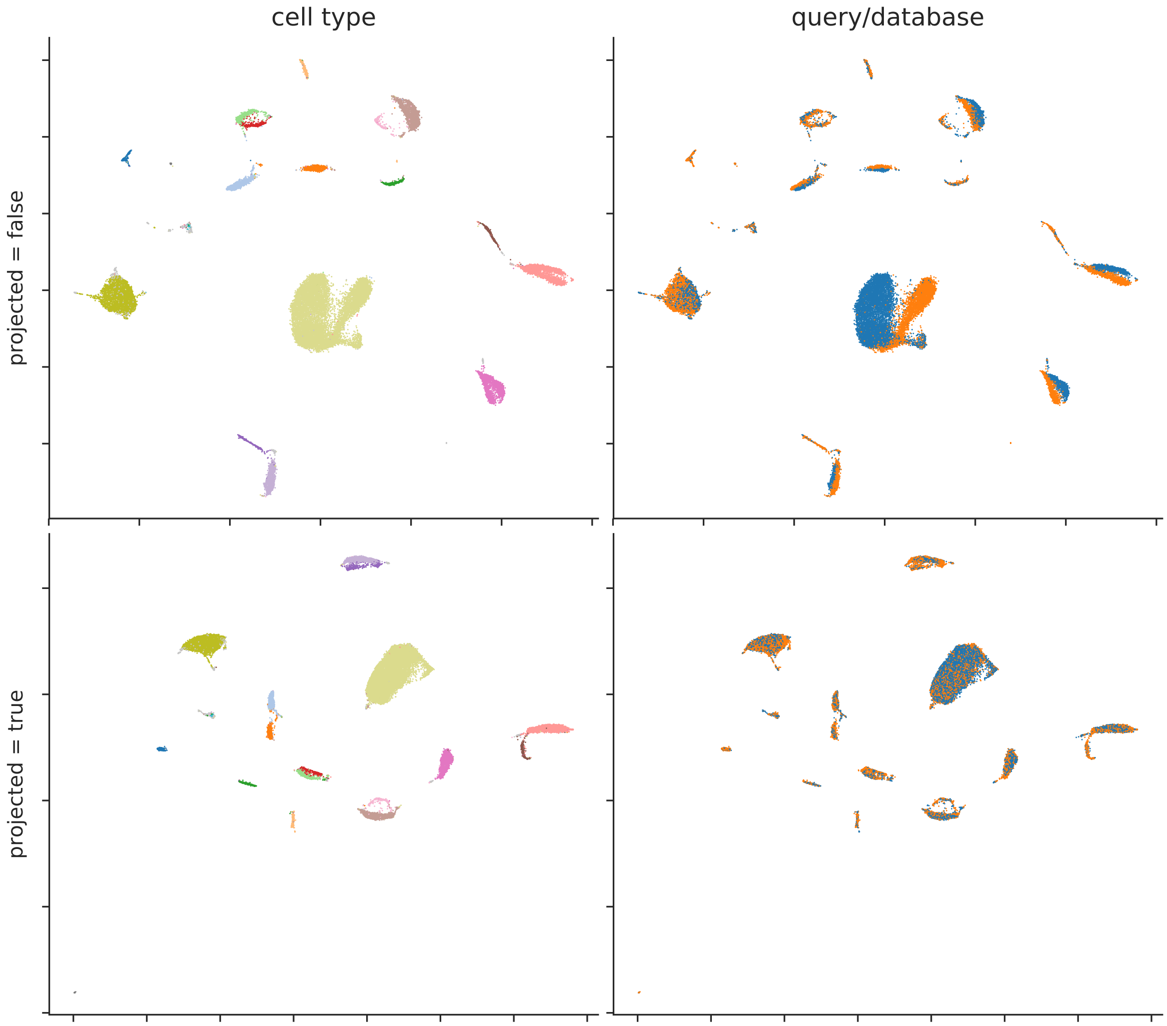

Figure 22: Two-dimensional embedding of expression profiles with UMAP (Shekhar2016, query batch = 2, cosine distance). In the left and right panels, points (cells) are colored by cell types and batches, respectively. In the upper panels, the query and database cells are mixed before dimensionality reduction. In the lower panels, the query cells are projected onto the space derived from the database cells.

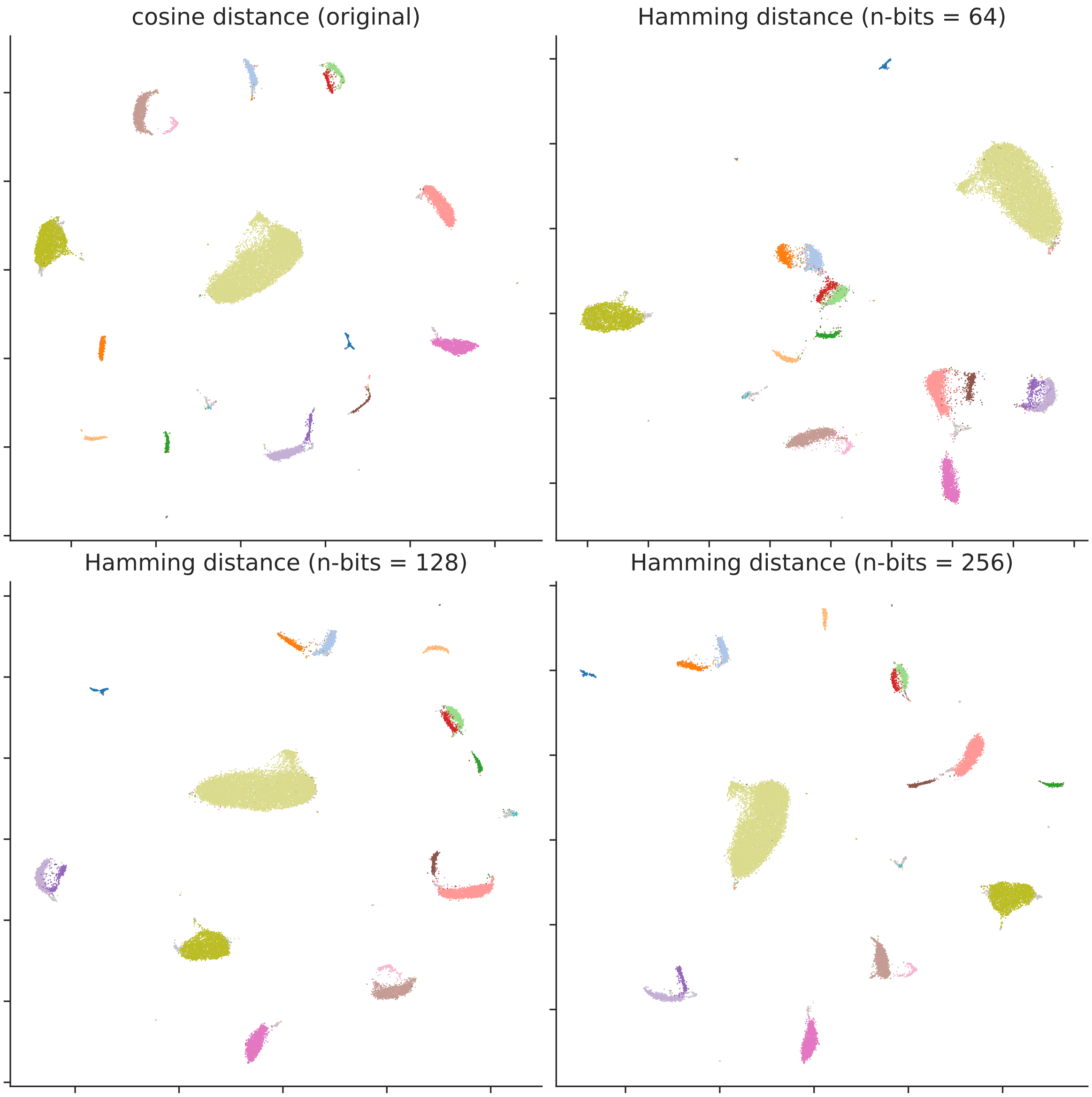

Figure 23: Two-dimensional embedding of expression profiles with UMAP (Shekhar2016, query batch = 1). Please refer to the main manuscript for the detailed caption.

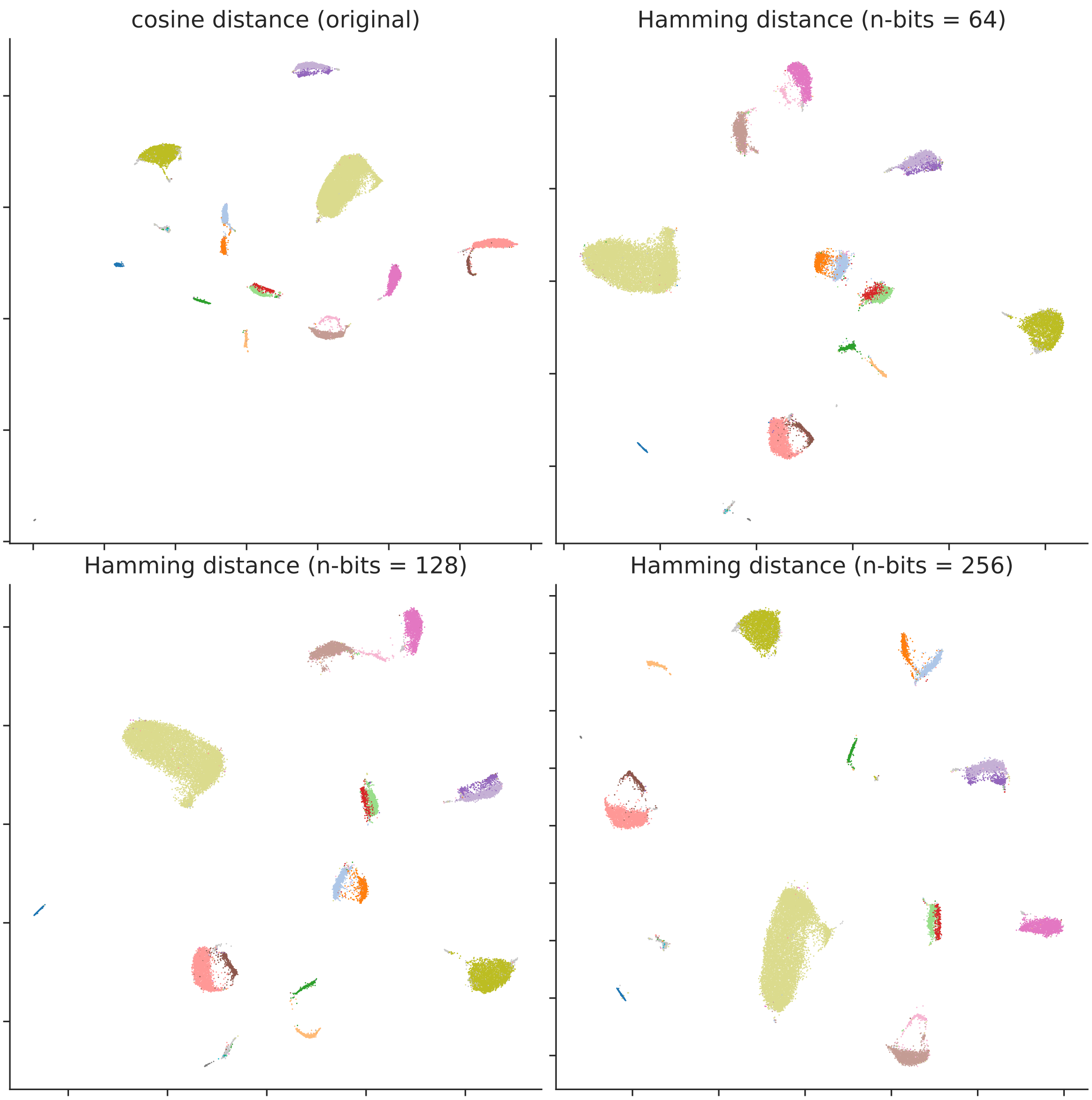

Figure 24: Two-dimensional embedding of expression profiles with UMAP (Shekhar2016, query batch = 2). Please refer to the main manuscript for the detailed caption.

#### Self-mapping experiments

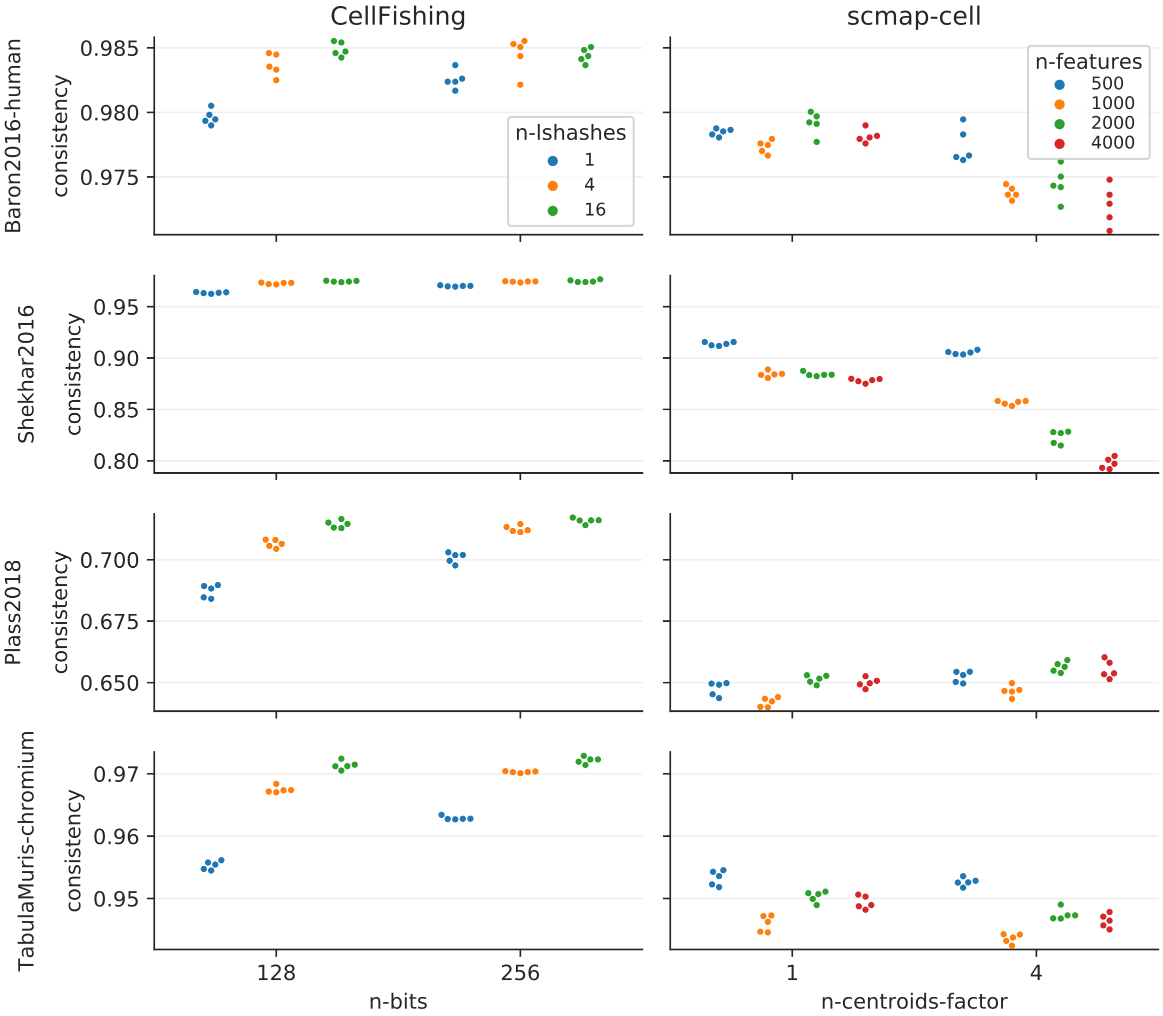

Figure 25: Comparison of consistency scores. Please refer to the main manuscript for the detailed caption.

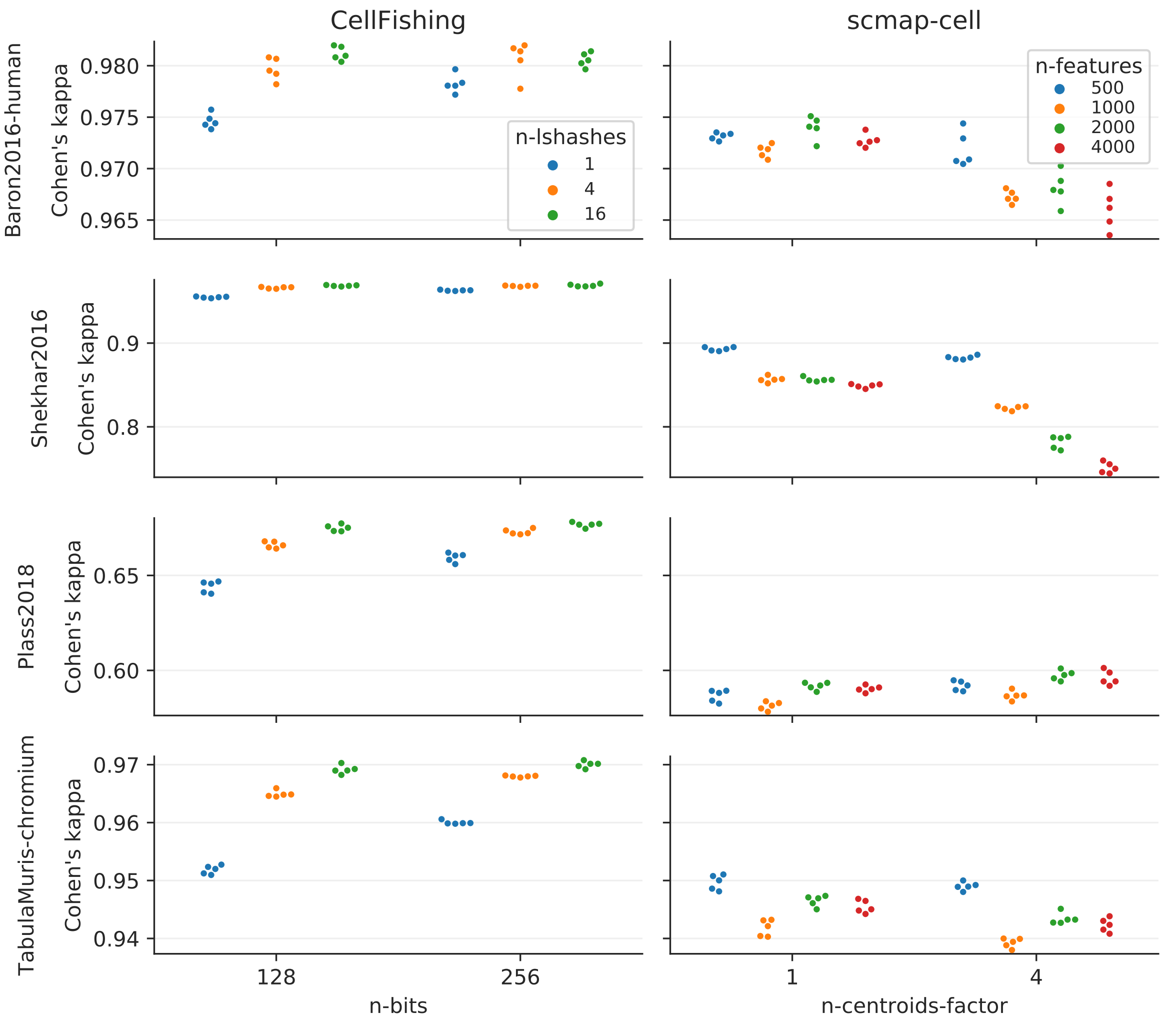

Figure 26: Comparison of Cohen's kappa scores. Please refer to the main manuscript for the detailed caption.

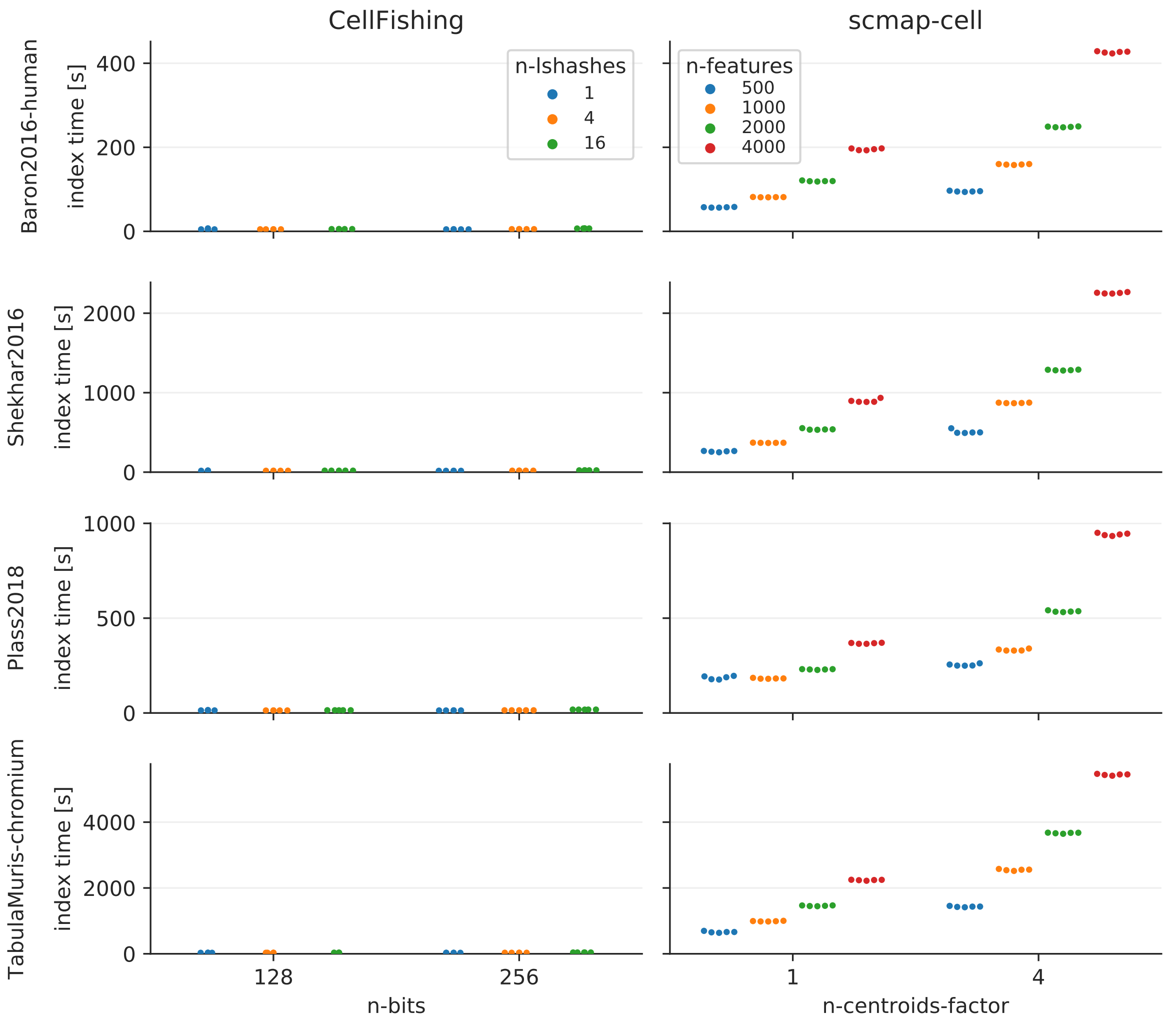

Figure 27: Comparison of index times. Please refer to the main manuscript for the detailed caption.

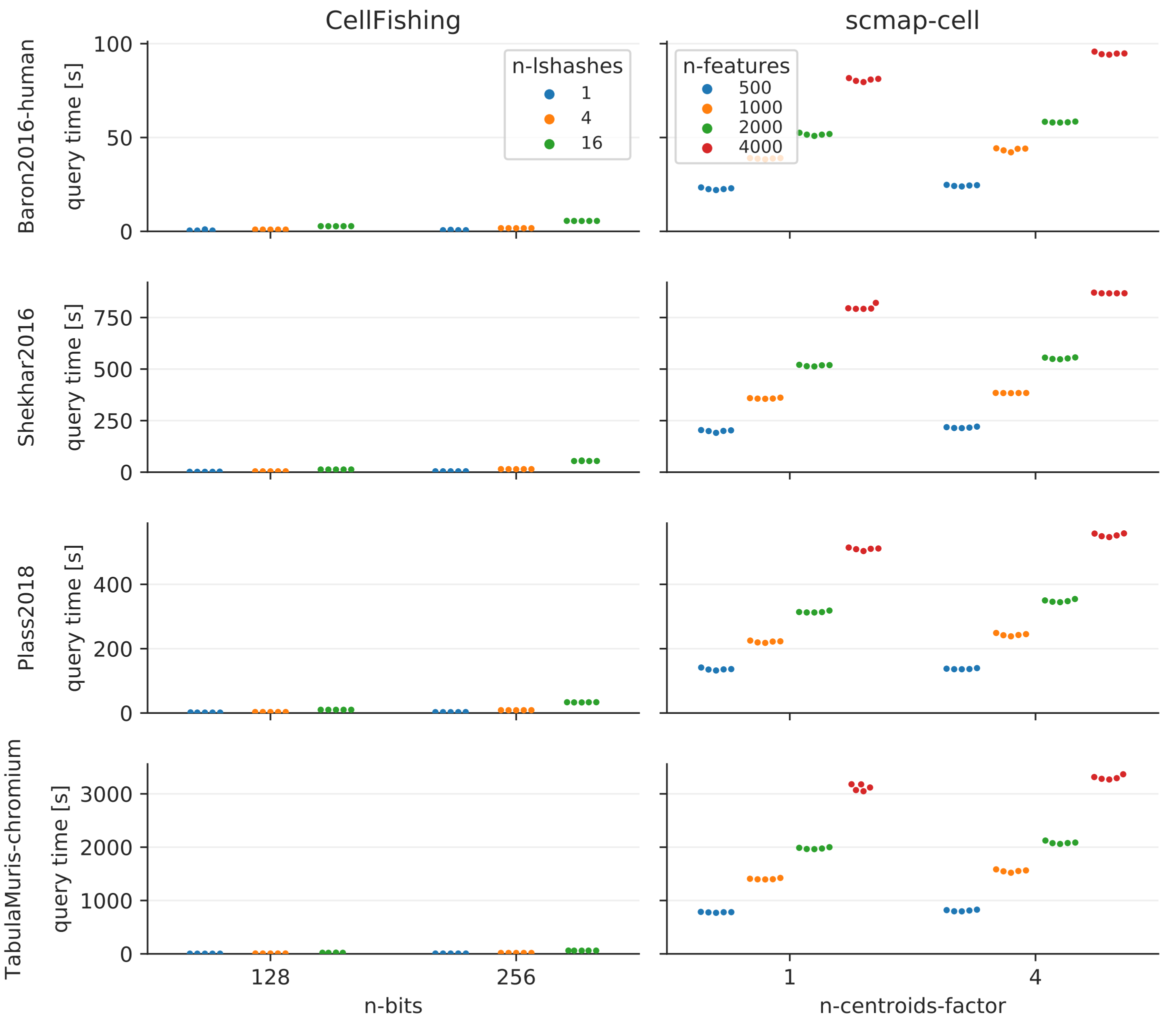

Figure 28: Comparison of query times. Please refer to the main manuscript for the detailed caption.

##### Cluster-specific scores

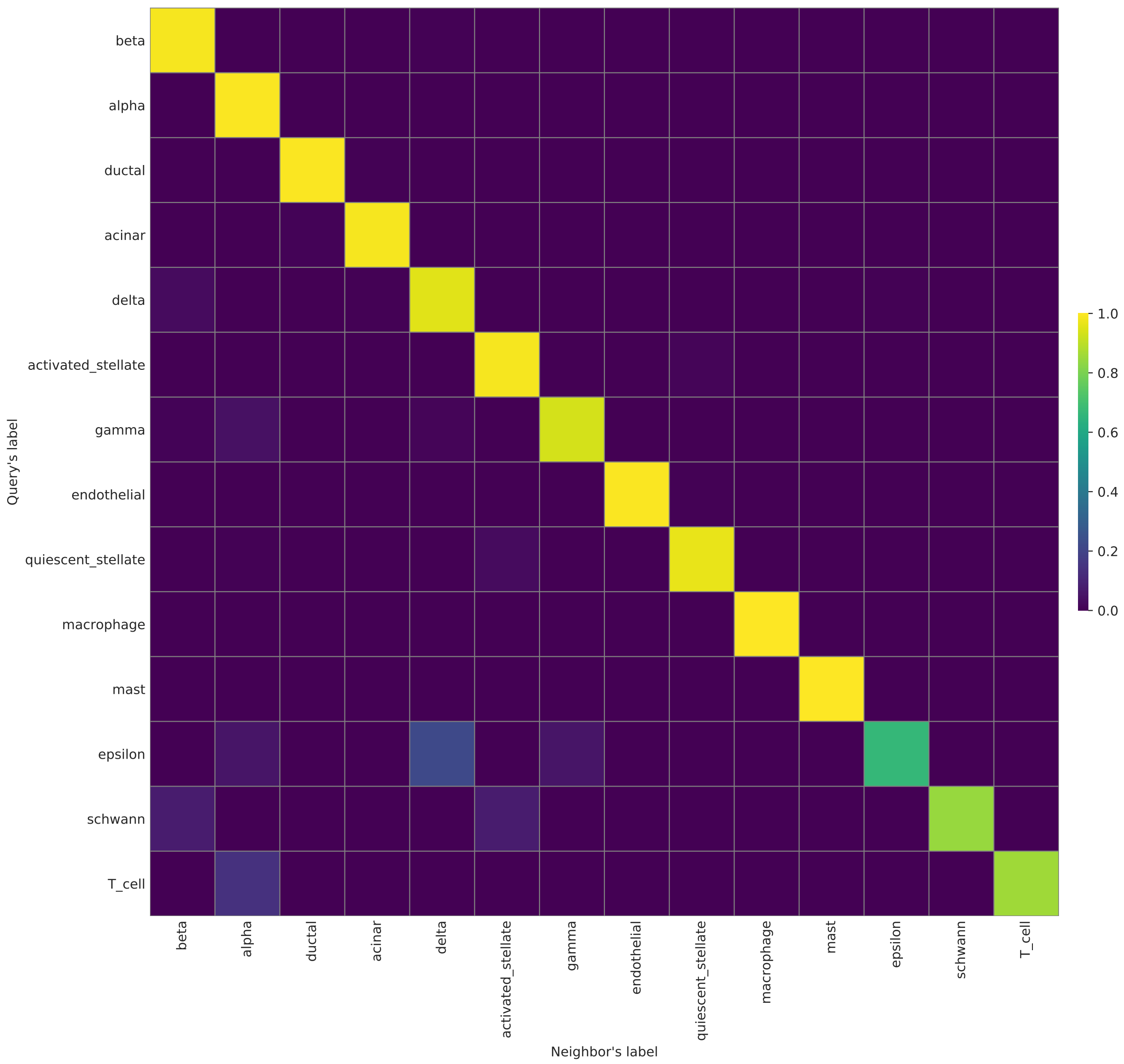

Figure 29: Proportions of cluster assignments (Baron2016, human, CellFishing). Please refer to the main manuscript for the detailed caption.

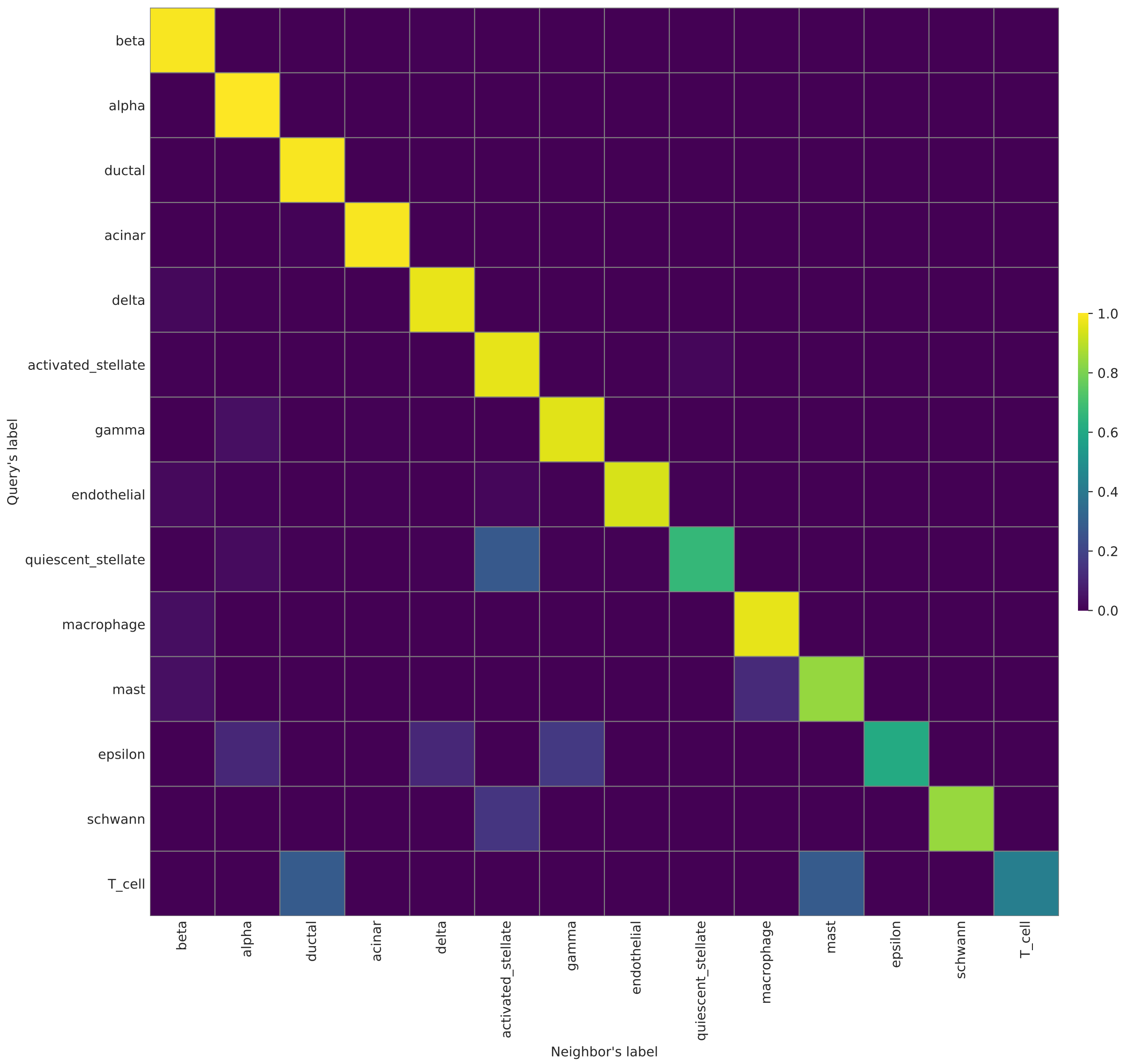

Figure 30: Proportions of cluster assignments (Baron2016, human, scmap-cell). Please refer to the main manuscript for the detailed caption.

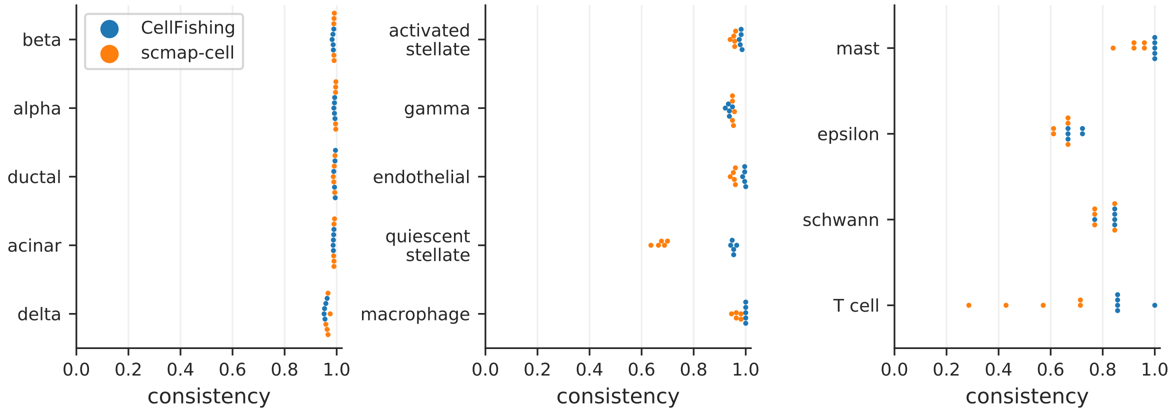

Figure 31: Cluster-specific consistency scores (Baron2016, human). Please refer to the main manuscript for the detailed caption.

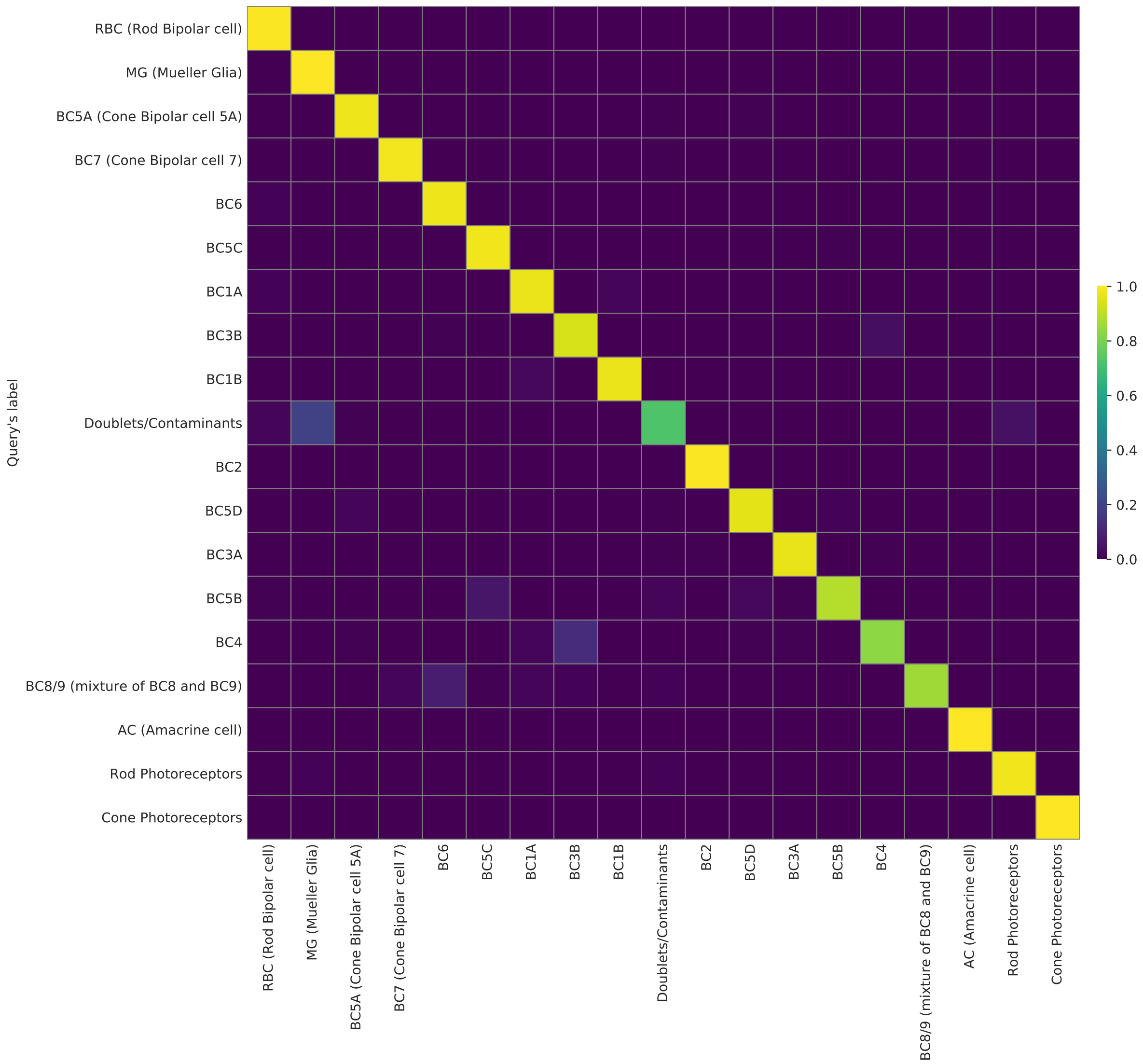

Figure 32: Proportions of cluster assignments (Shekhar2016, CellFishing). Please refer to the main manuscript for the detailed caption.

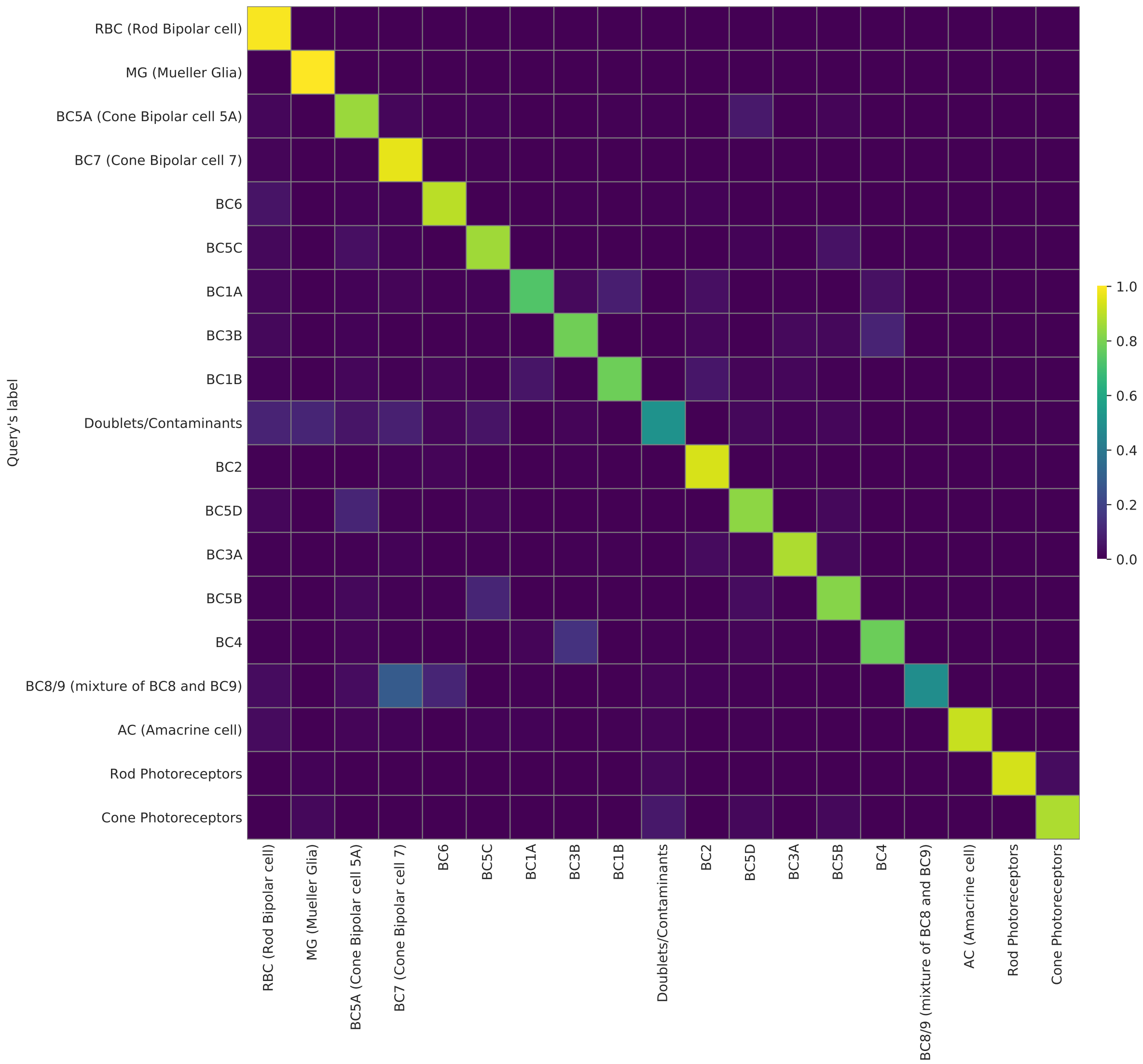

Figure 33: Proportions of cluster assignments (Shekhar2016, scmap-cell). Please refer to the main manuscript for the detailed caption.

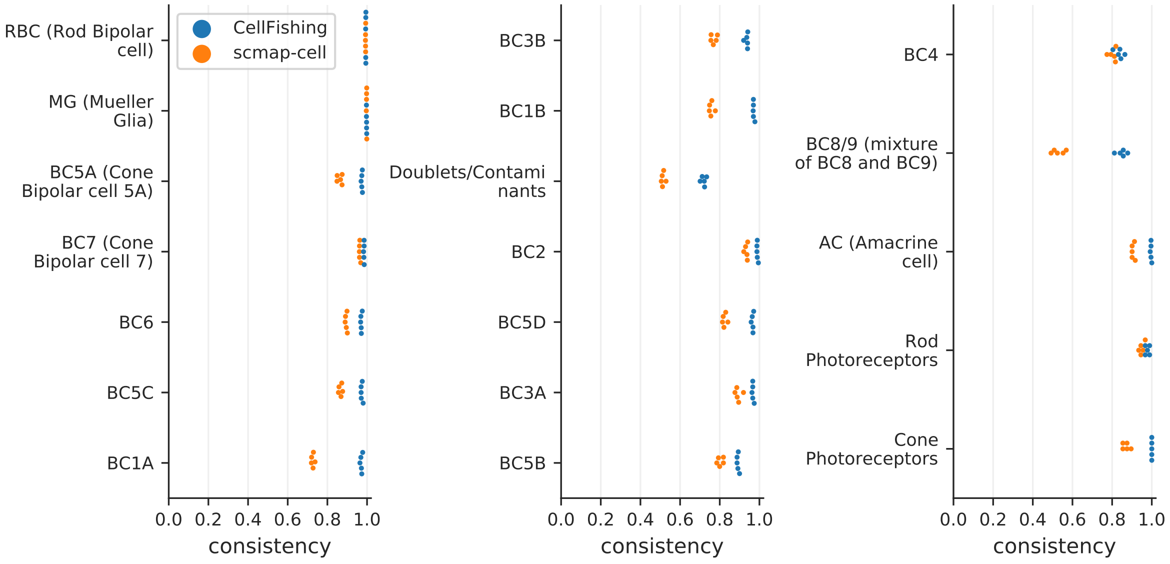

Figure 34: Cluster-specific consistency scores (Shekhar2016). Please refer to the main manuscript for the detailed caption.

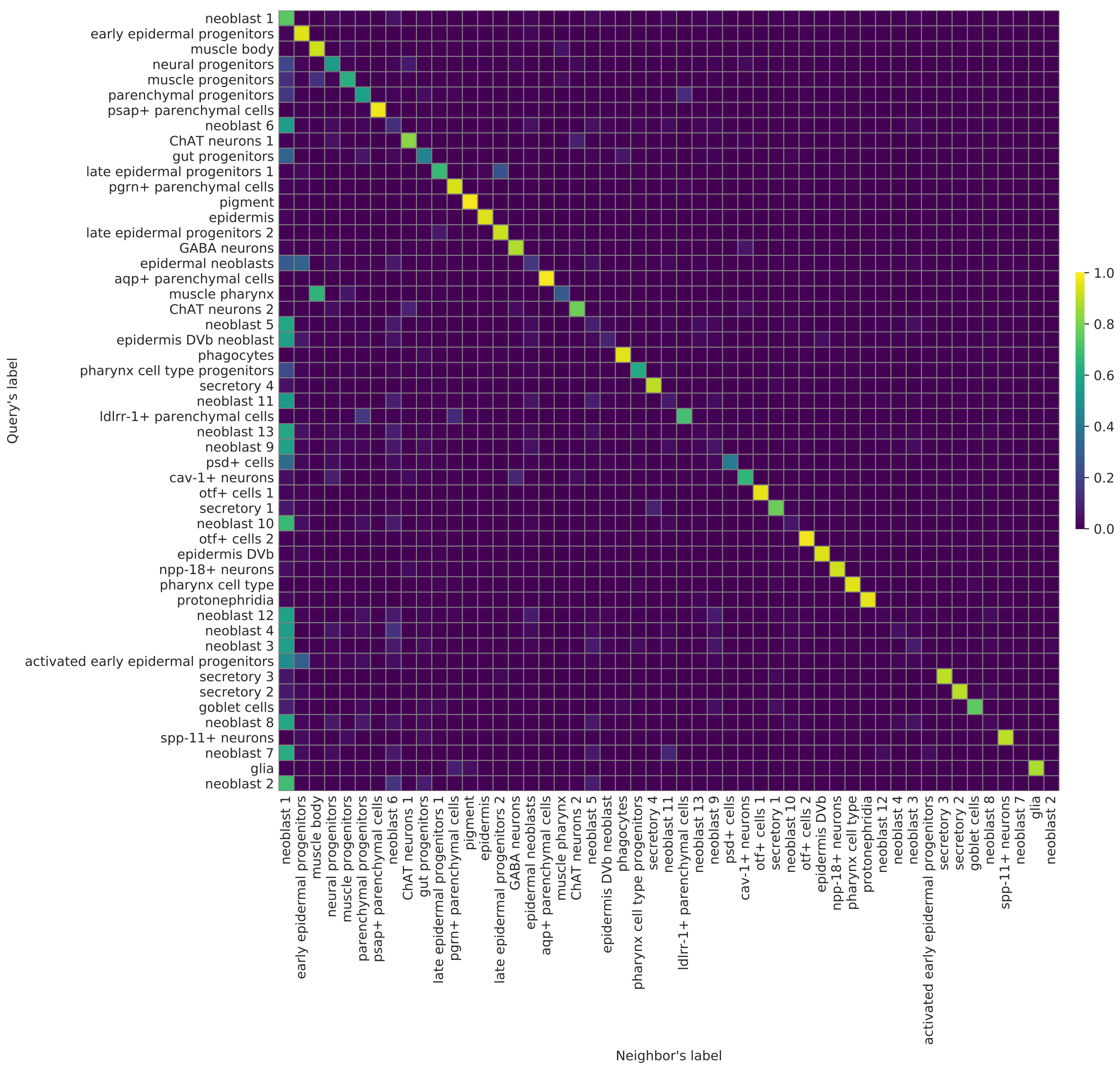

Figure 35: Proportions of cluster assignments (Plass2018, CellFishing). Please refer to the main manuscript for the detailed caption.

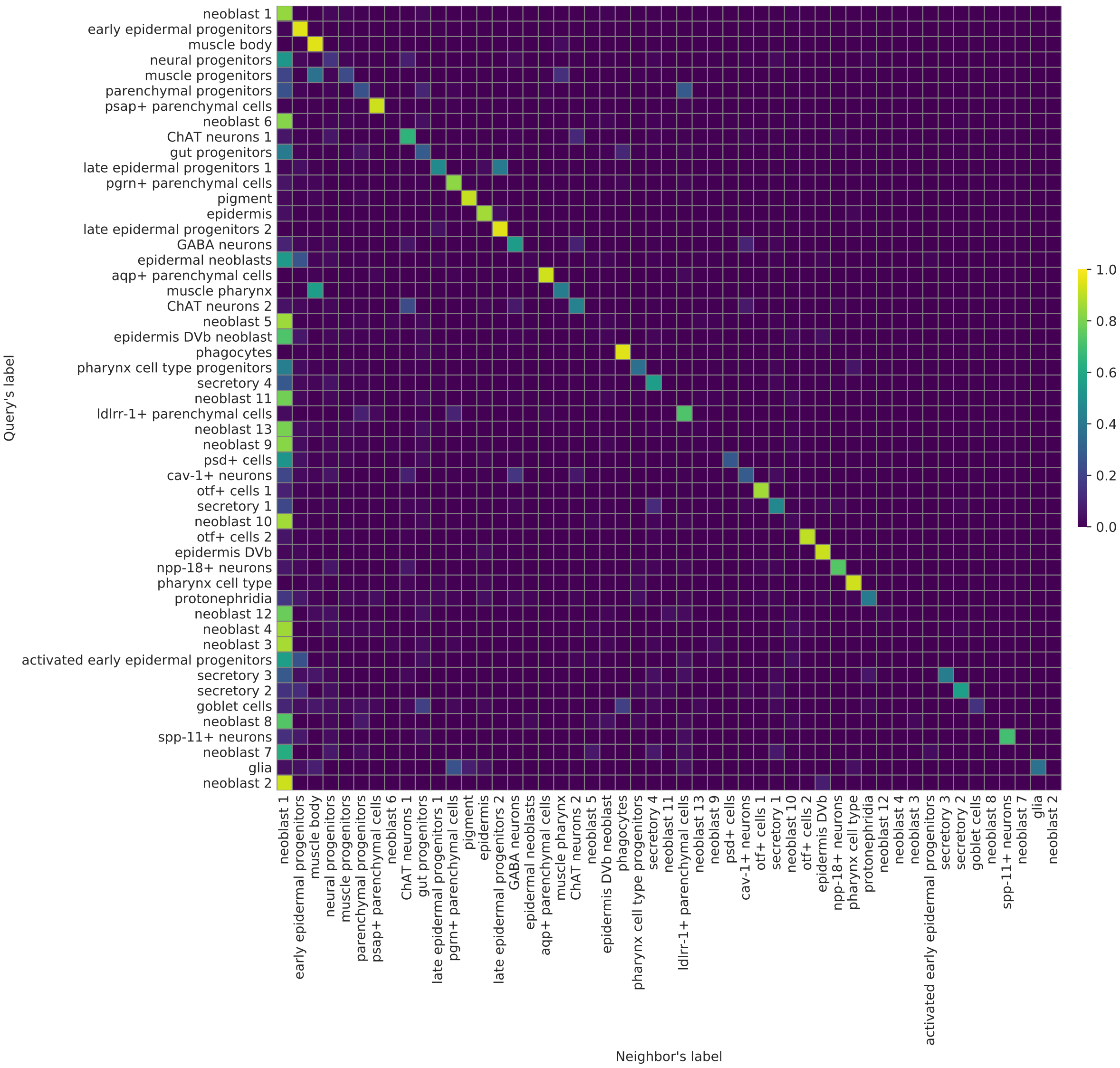

Figure 36: Proportions of cluster assignments (Plass2018, scmap-cell). Please refer to the main manuscript for the detailed caption.

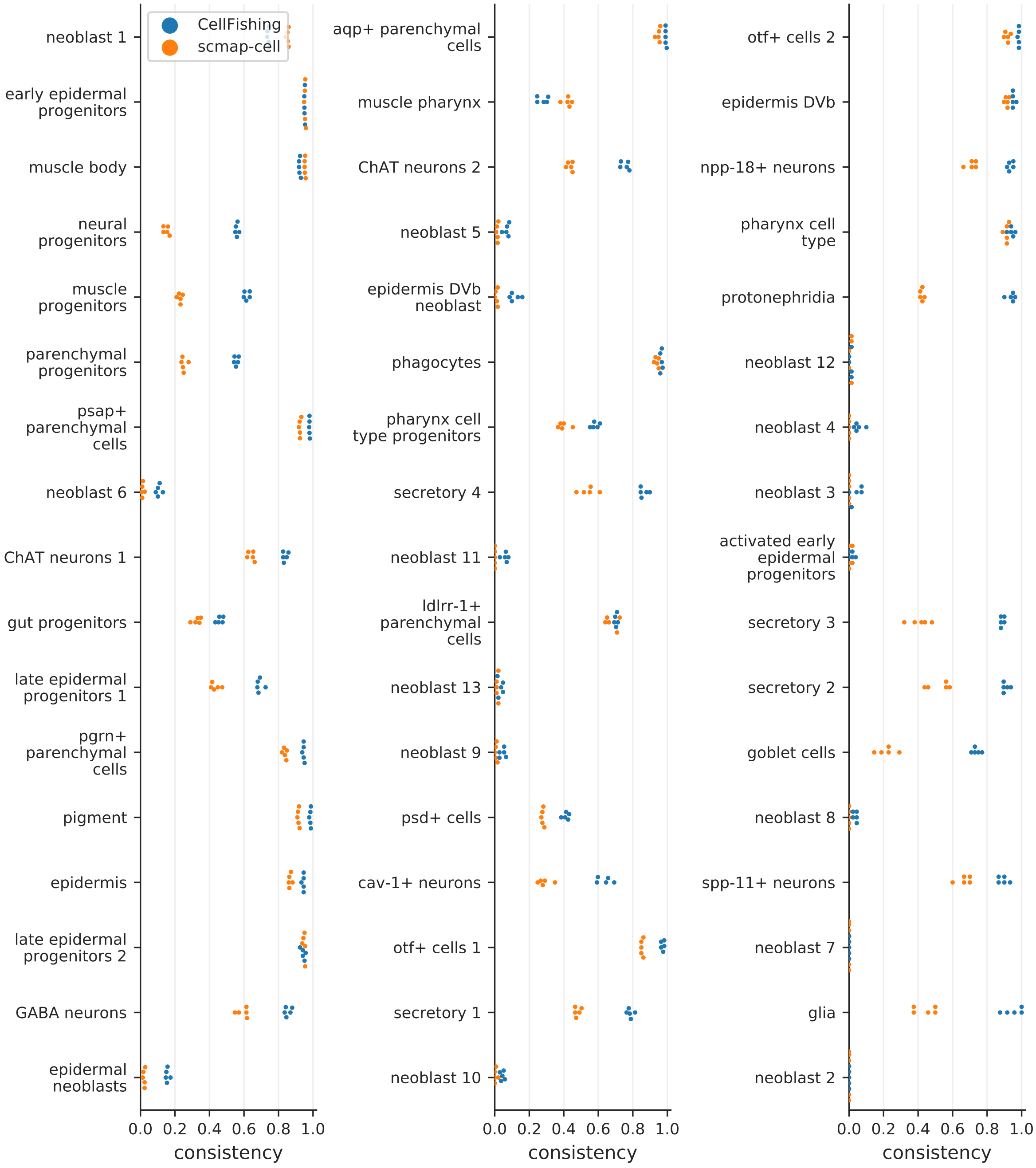

Figure 37: Cluster-specific consistency scores (Plass2018). Please refer to the main manuscript for the detailed caption.

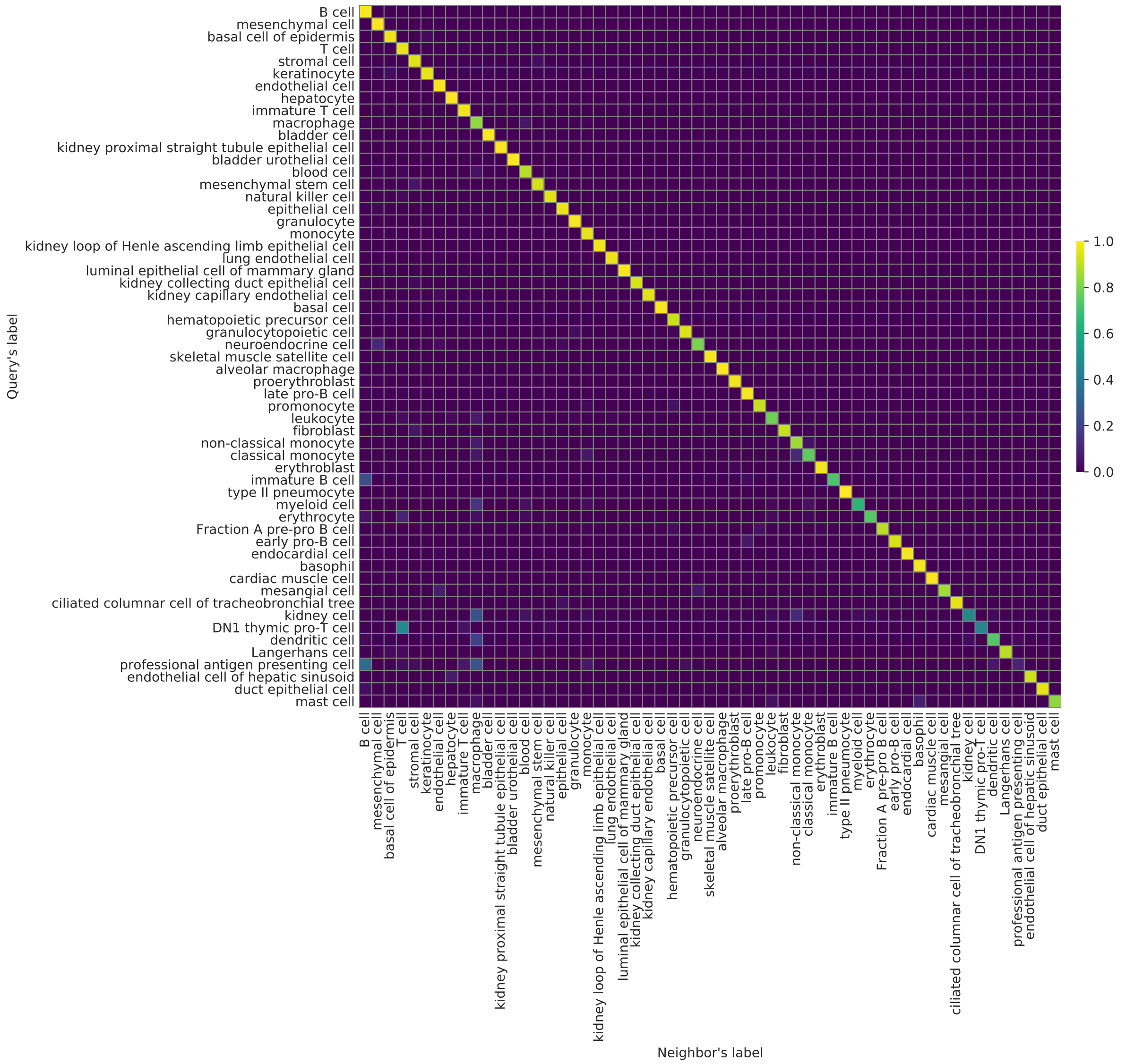

Figure 38: Proportions of cluster assignments (TabulaMuris, Chromium, Cell-Fishing). Please refer to the main manuscript for the detailed caption.

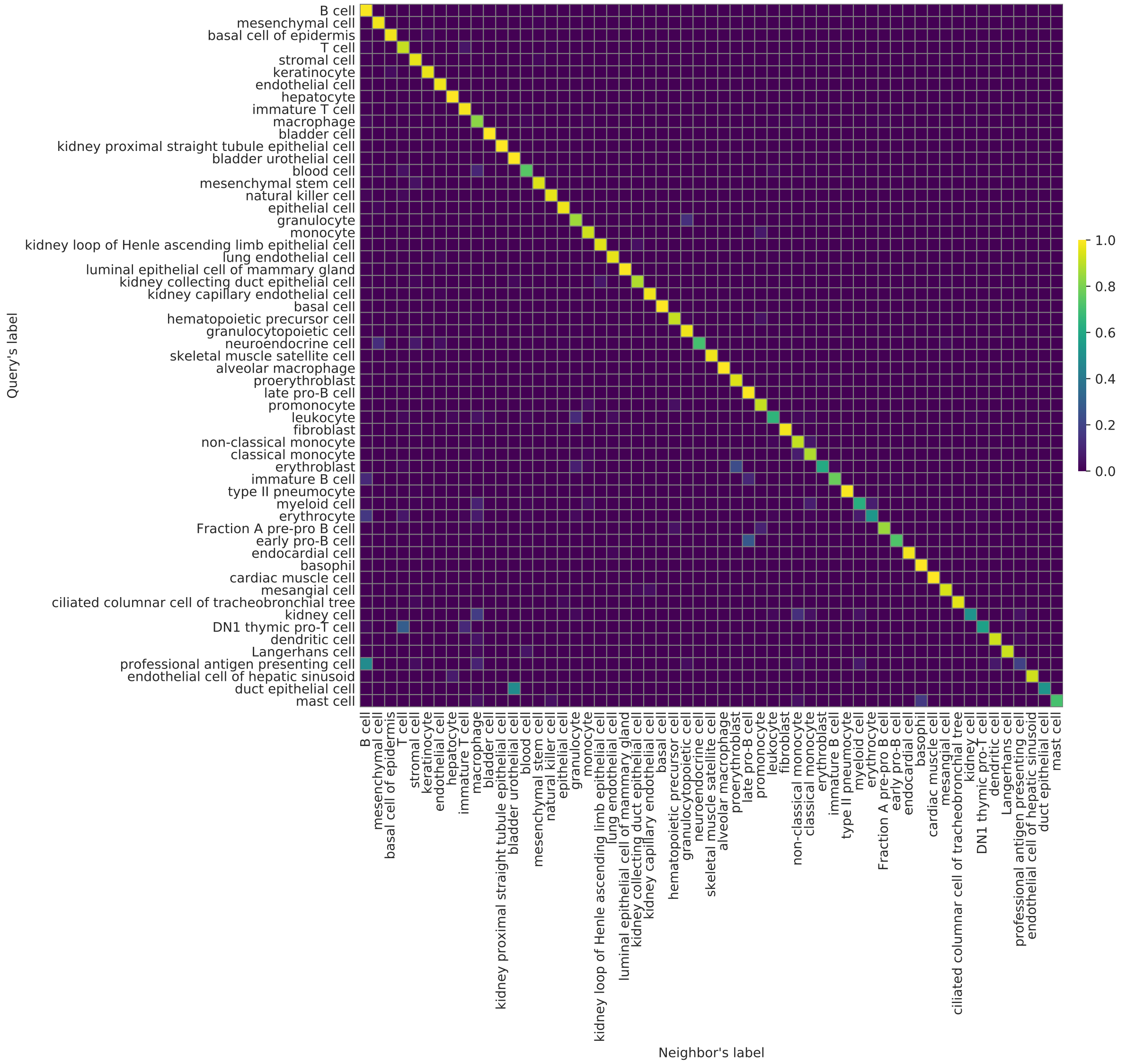

Figure 39: Proportions of cluster assignments (TabulaMuris, Chromium, scmap-cell). Please refer to the main manuscript for the detailed caption.

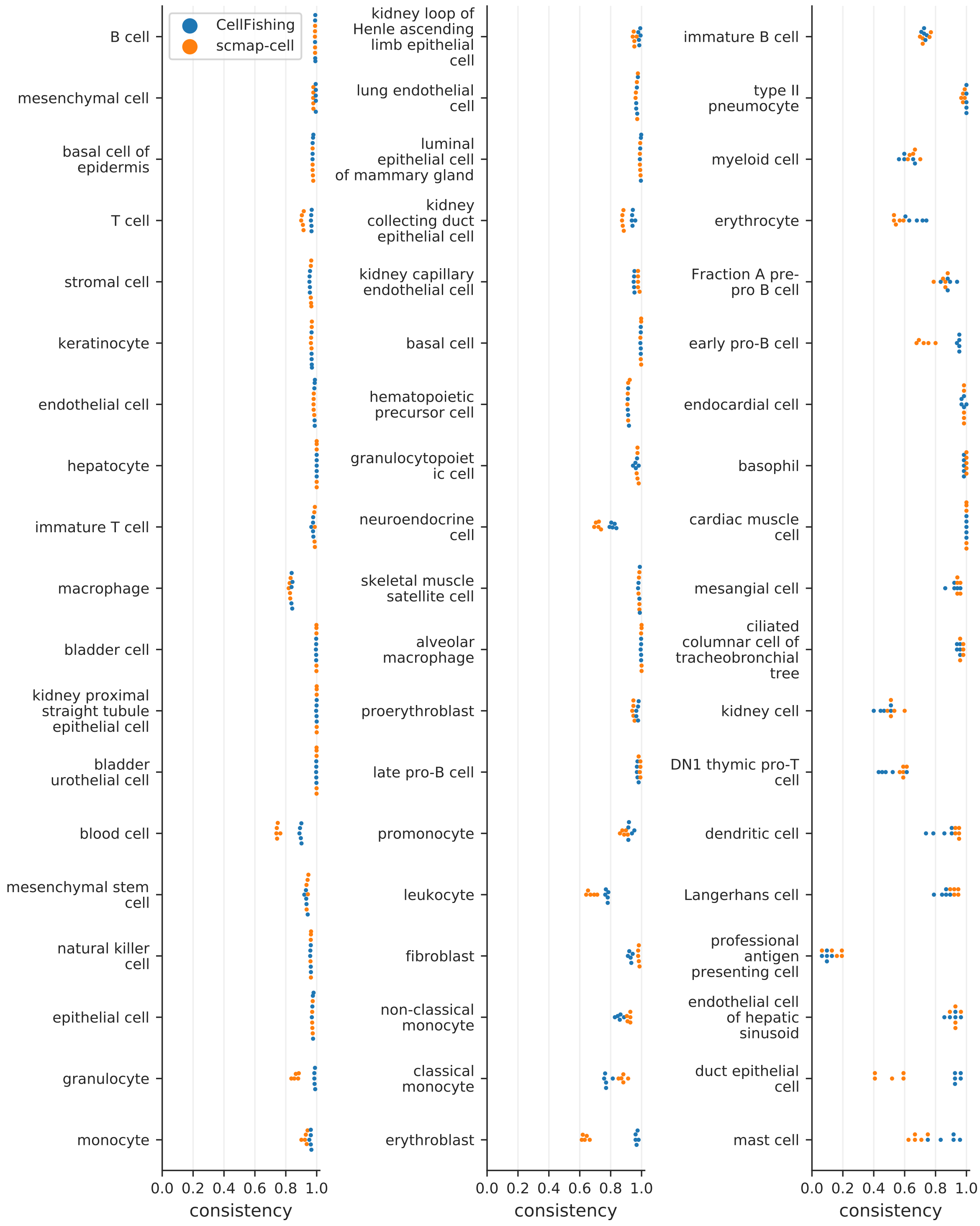

Figure 40: Cluster-specific consistency scores (TabulaMuris, Chromium). Please refer to the main manuscript for the detailed caption.

#### Similarities from nearest neighbors

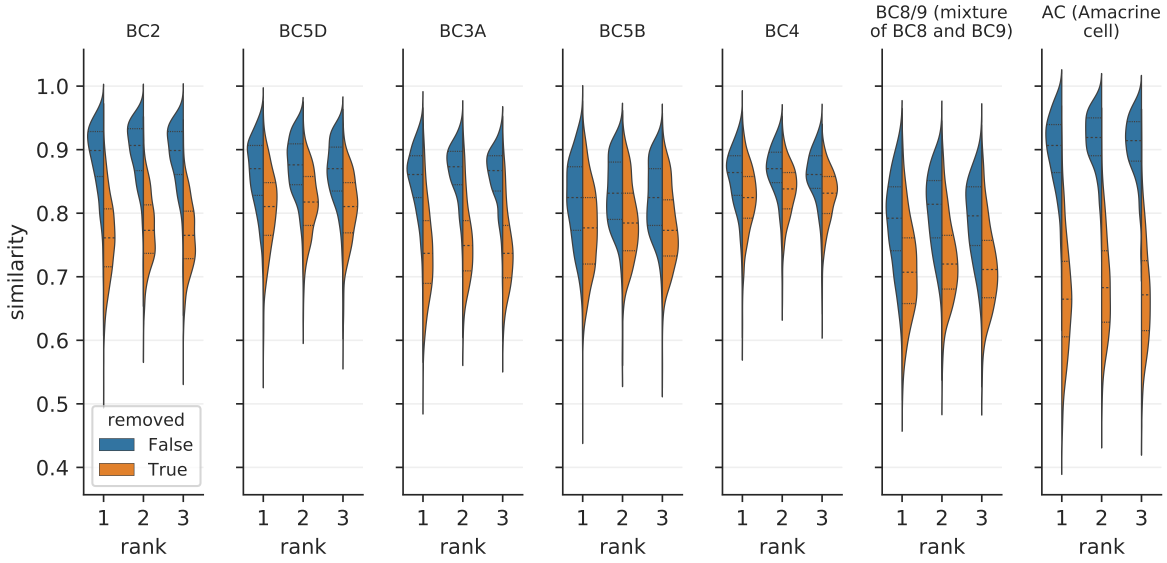

Figure 41: Distributions of similarities with or without a specific cell type (Shekhar2016). Please refer to the main manuscript for the detailed caption.

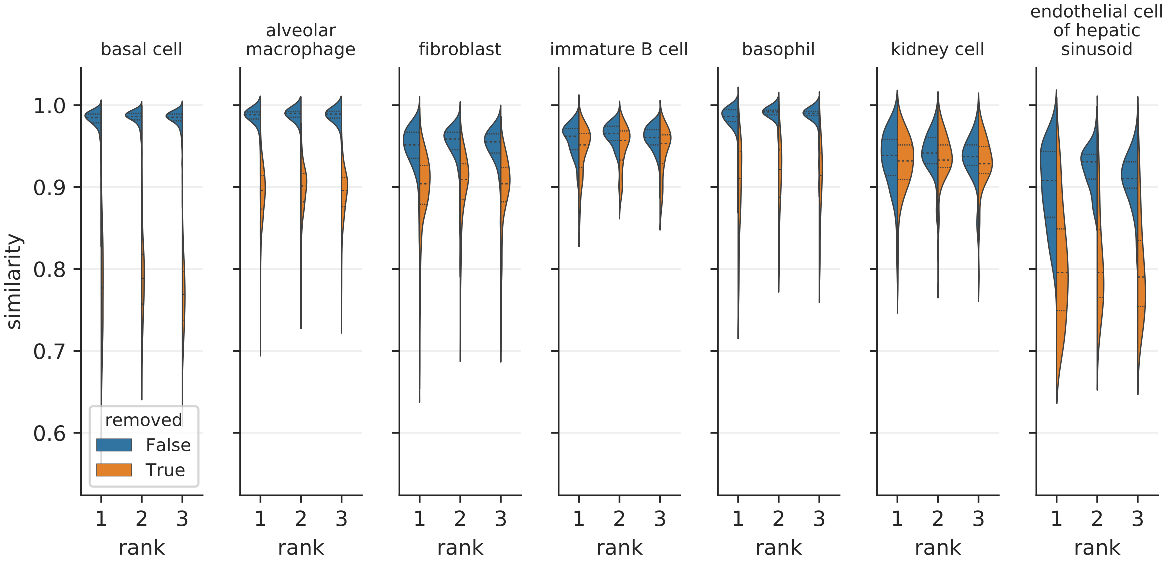

Figure 42: Distributions of similarities with or without a specific cell type (TabulaMuris, Chromium). Please refer to the main manuscript for the detailed caption.

#### Mapping across batches

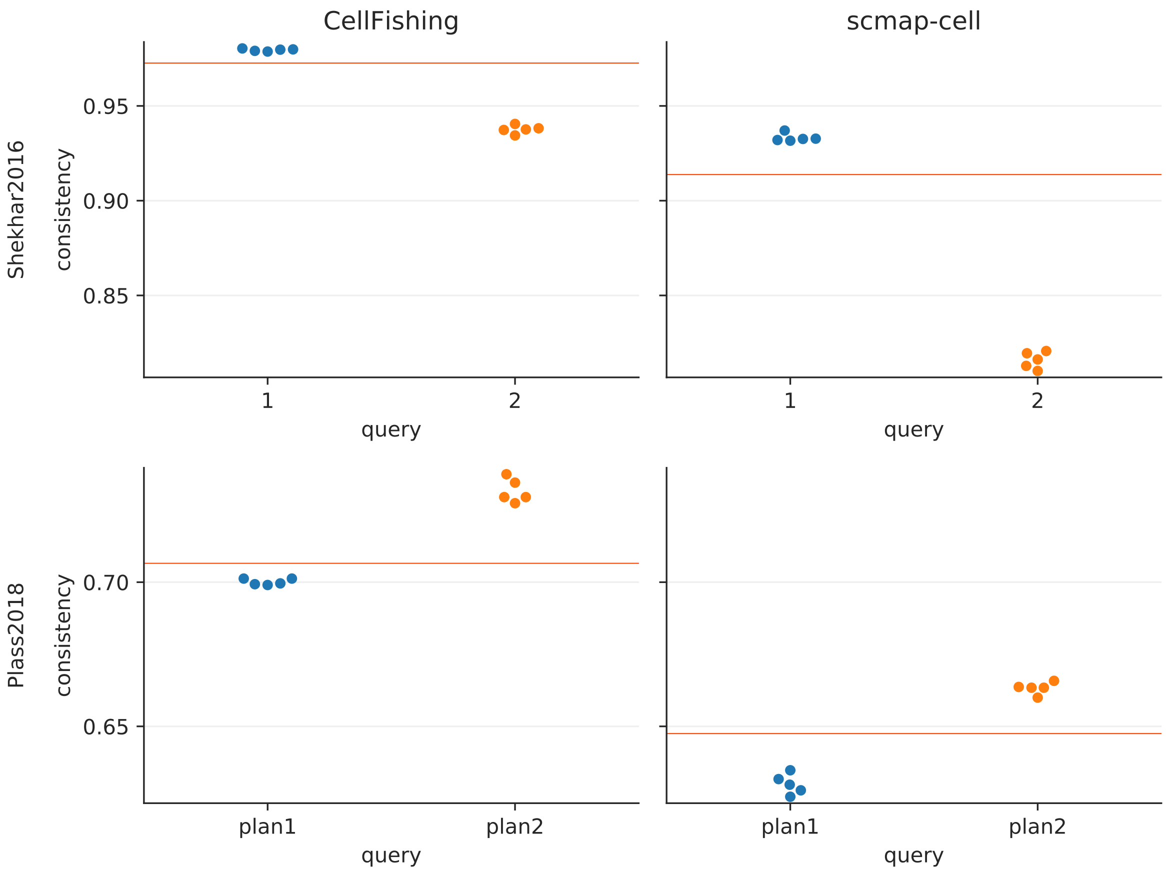

Figure 43: Cell mapping across batches (consistency). Please refer to the main manuscript for the detailed caption.

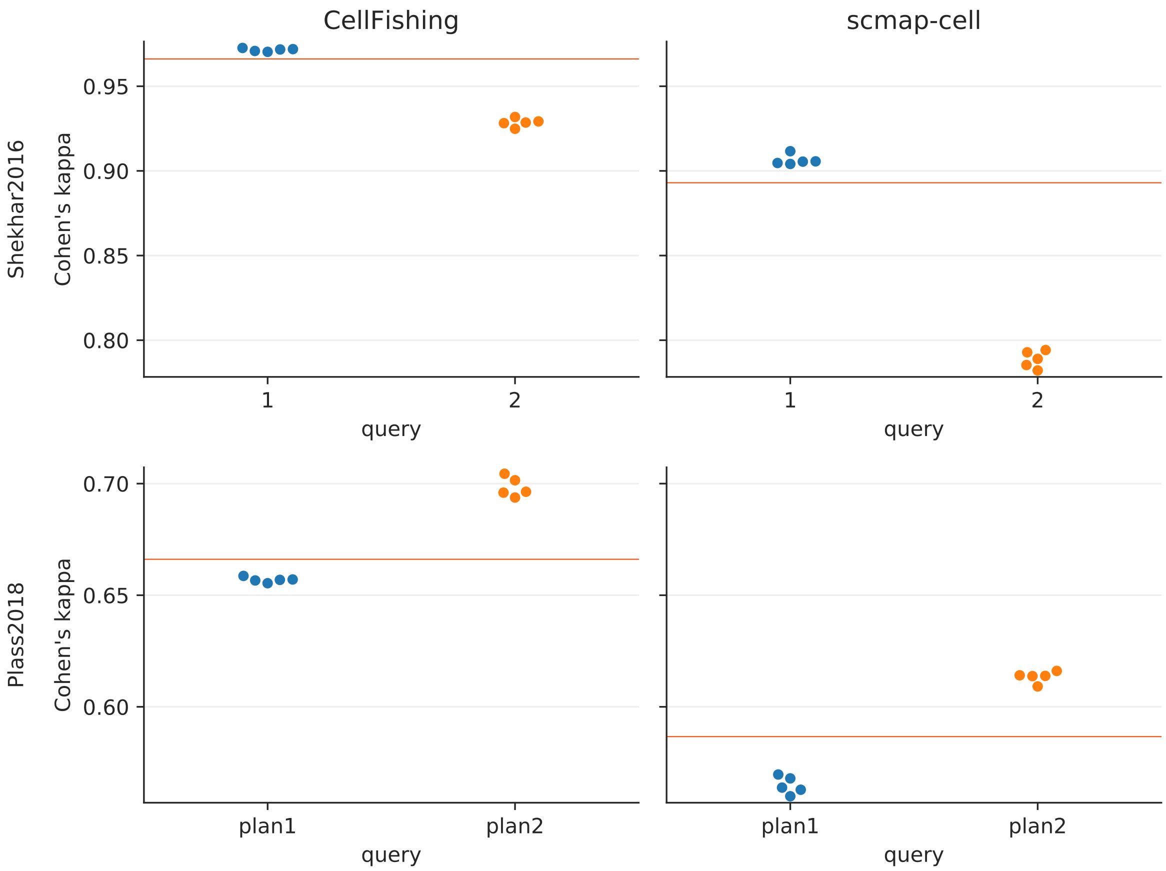

Figure 44: Cell mapping across batches (Cohen's kappa). Please refer to the main manuscript for the detailed caption.

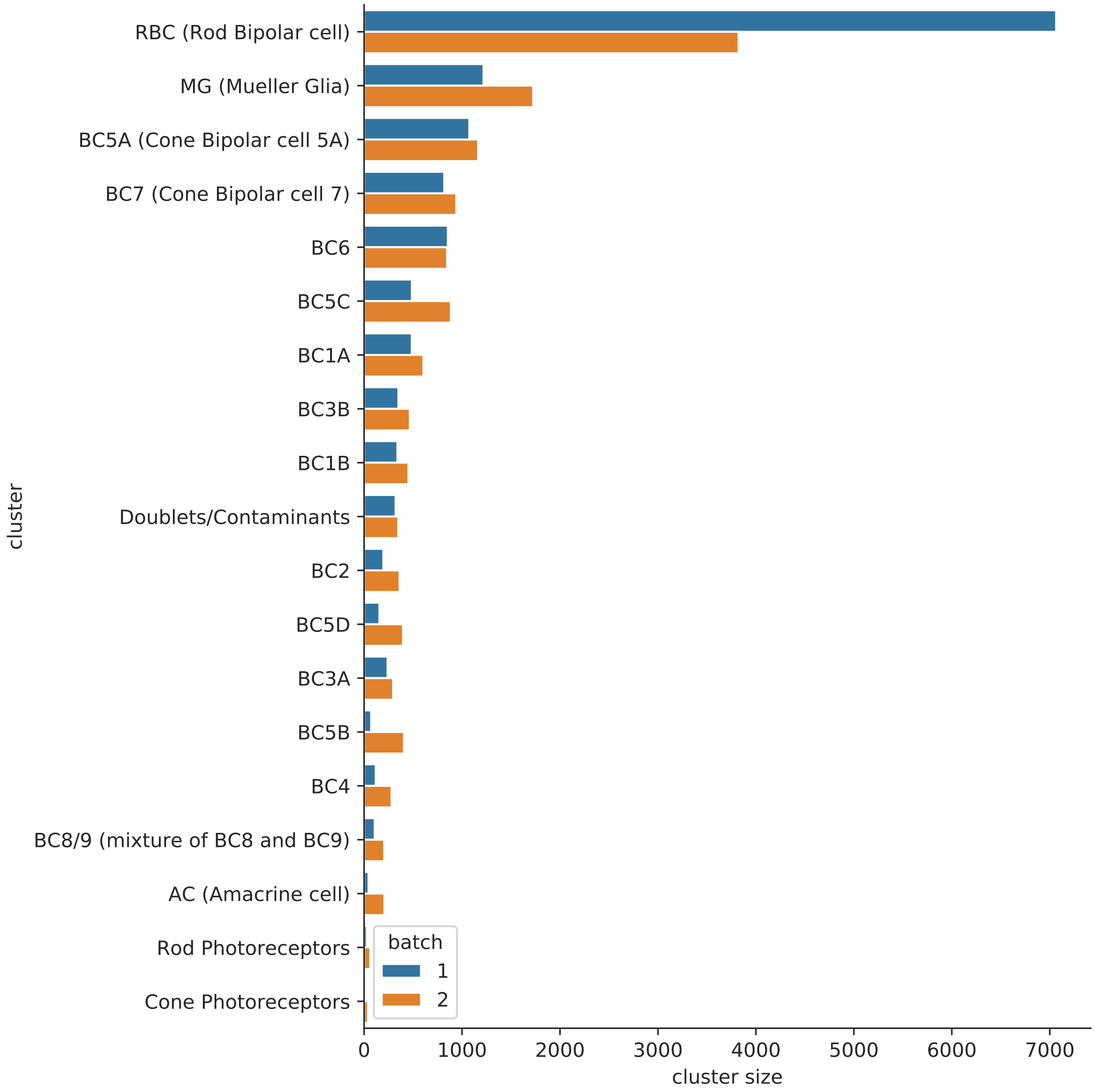

Figure 45: Distribution of cluster sizes (Shekhar2016). Please refer to the main manuscript for the detailed caption.

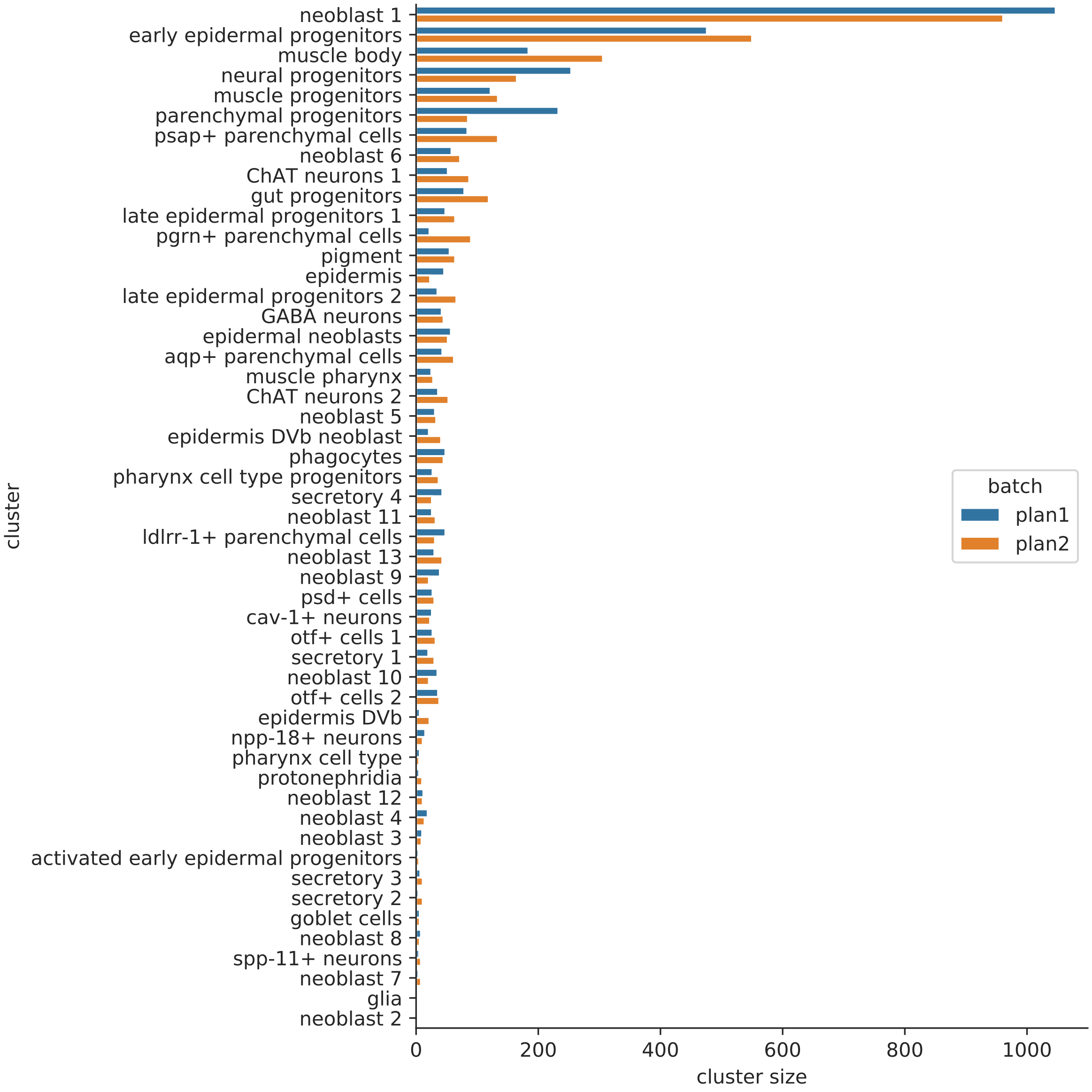

Figure 46: Distribution of cluster sizes (Plass2018). Please refer to the main manuscript for the detailed caption.

#### Mapping across protocols

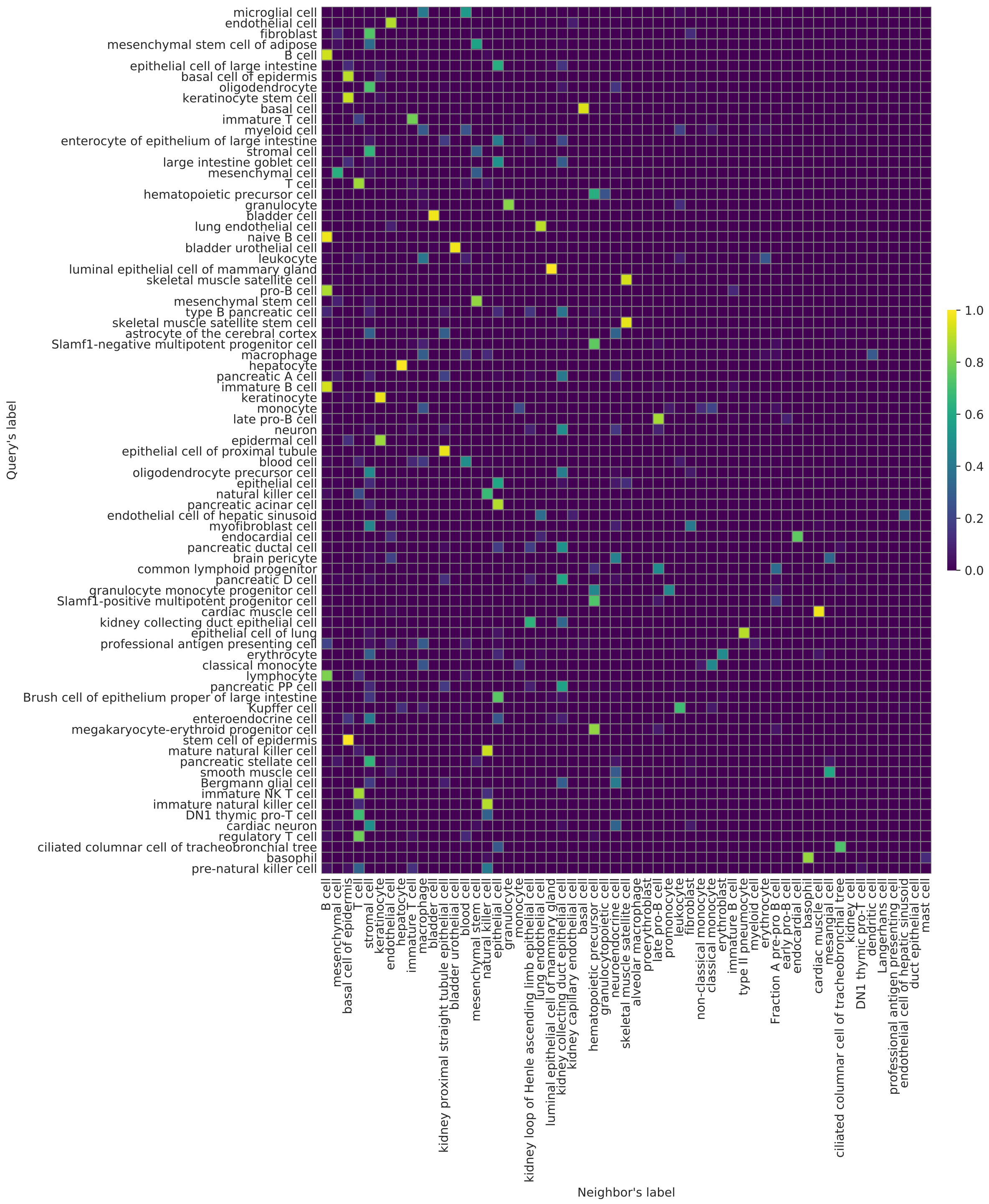

Figure 47: Cluster assignments across different protocol (TabulaMuris, Chromium, CellFishing, All). Please refer to the main manuscript for the detailed caption.

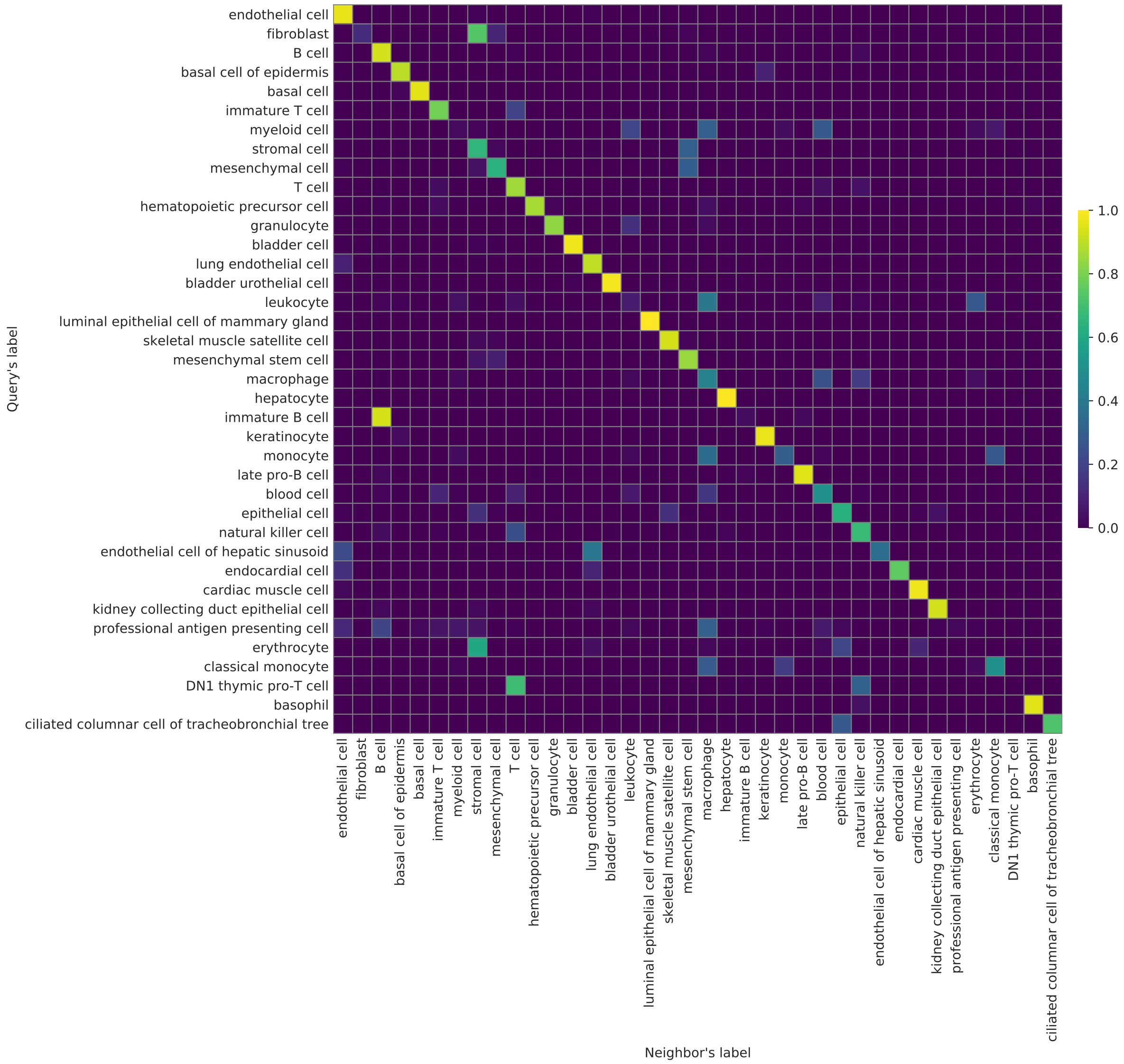

Figure 48: Cluster assignments across different protocol (TabulaMuris, Chromium, CellFishing, Overlapping cell types). Please refer to the main manuscript for the detailed caption.

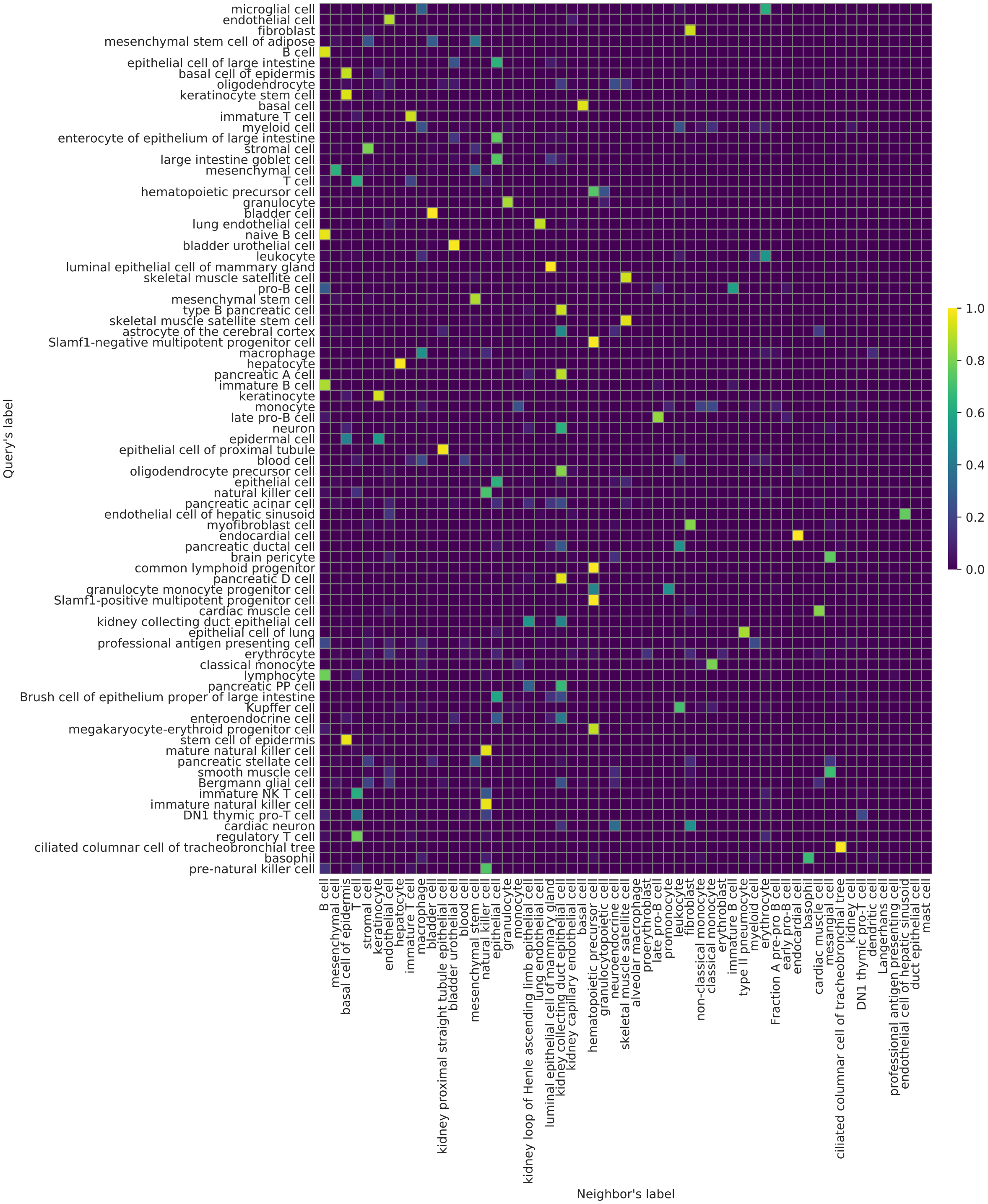

Figure 49: Cluster assignments across different protocol (TabulaMuris, Chromium, scmap-cell, All). Please refer to the main manuscript for the detailed caption.

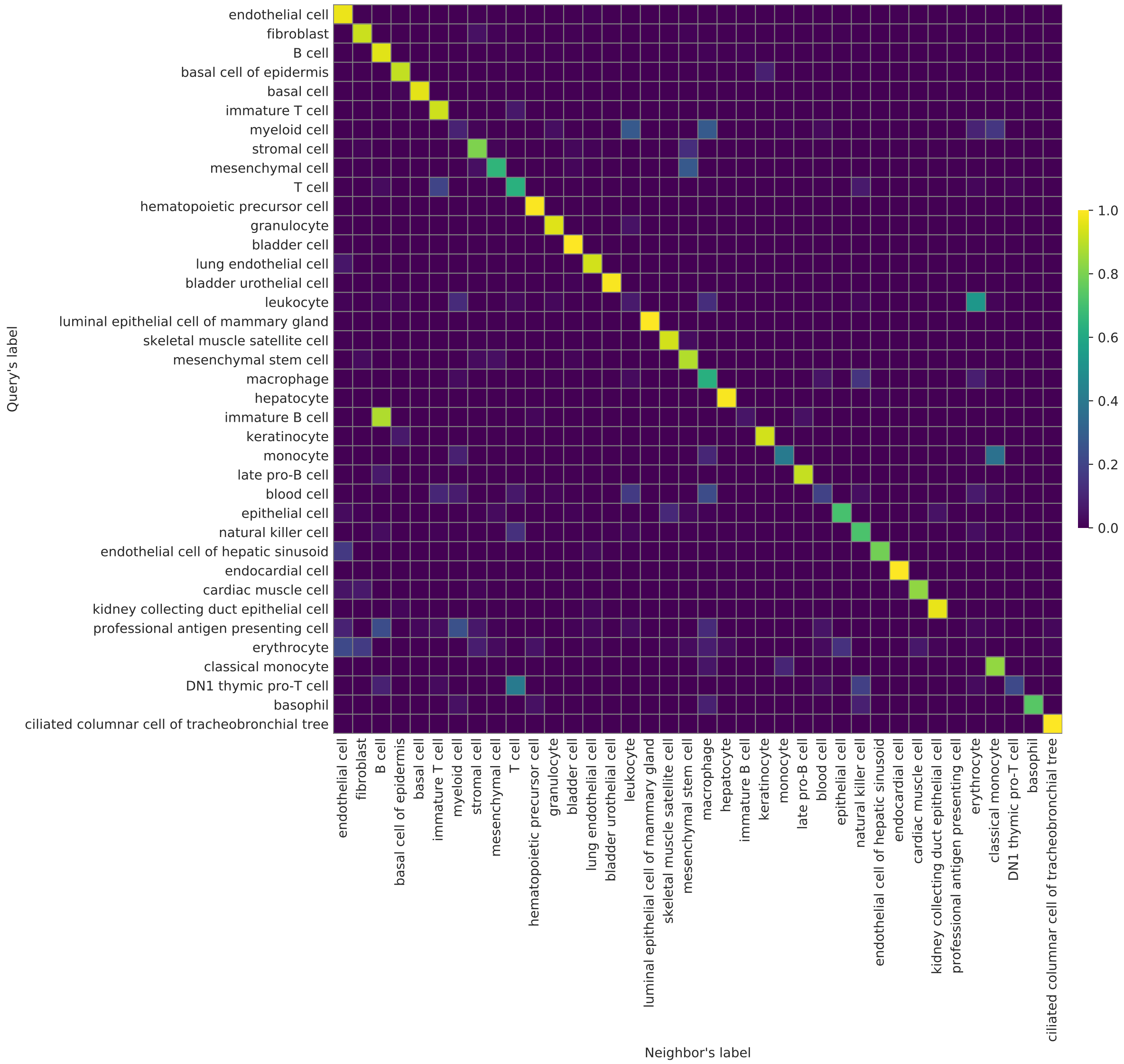

Figure 50: Cluster assignments across different protocol (TabulaMuris, Chromium, scmap-cell, Overlapping cell types). Please refer to the main manuscript for the detailed caption.

### Benchmarks of saving and loading

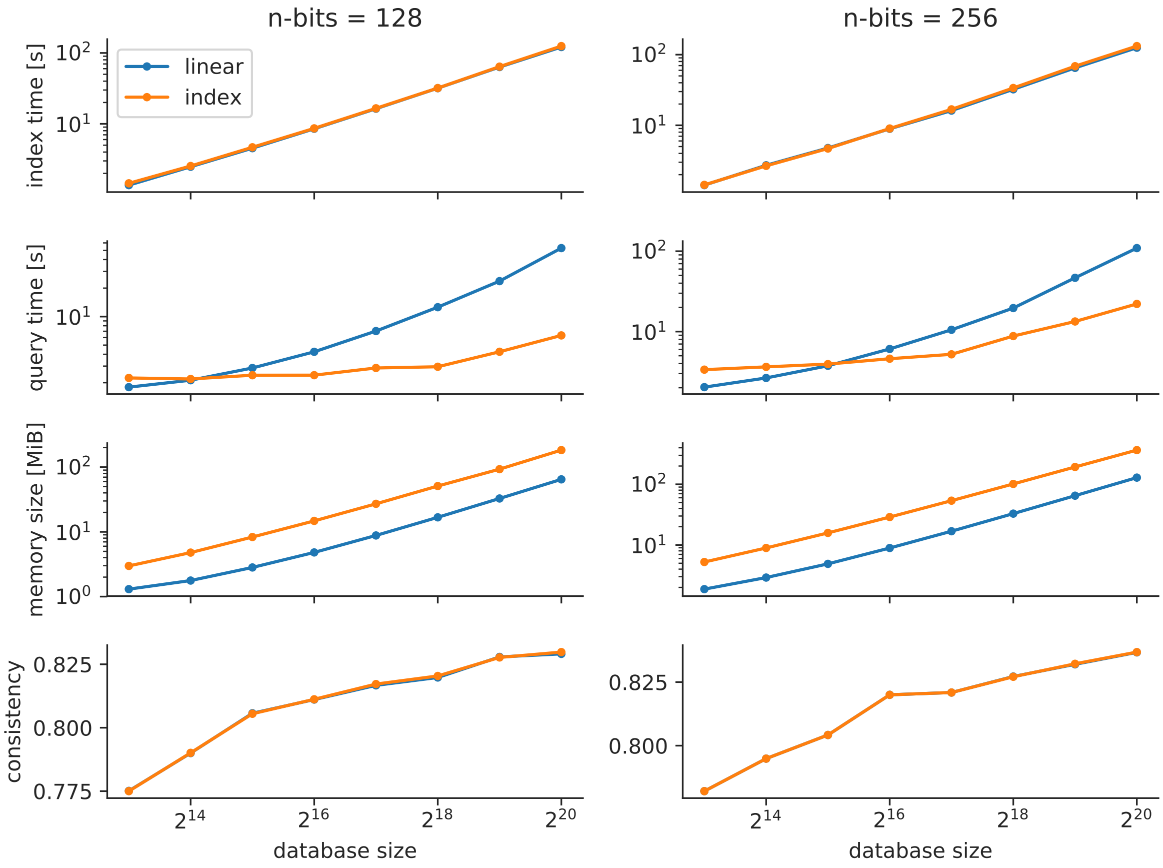

Figure 51: Scalability (1M_neurons).

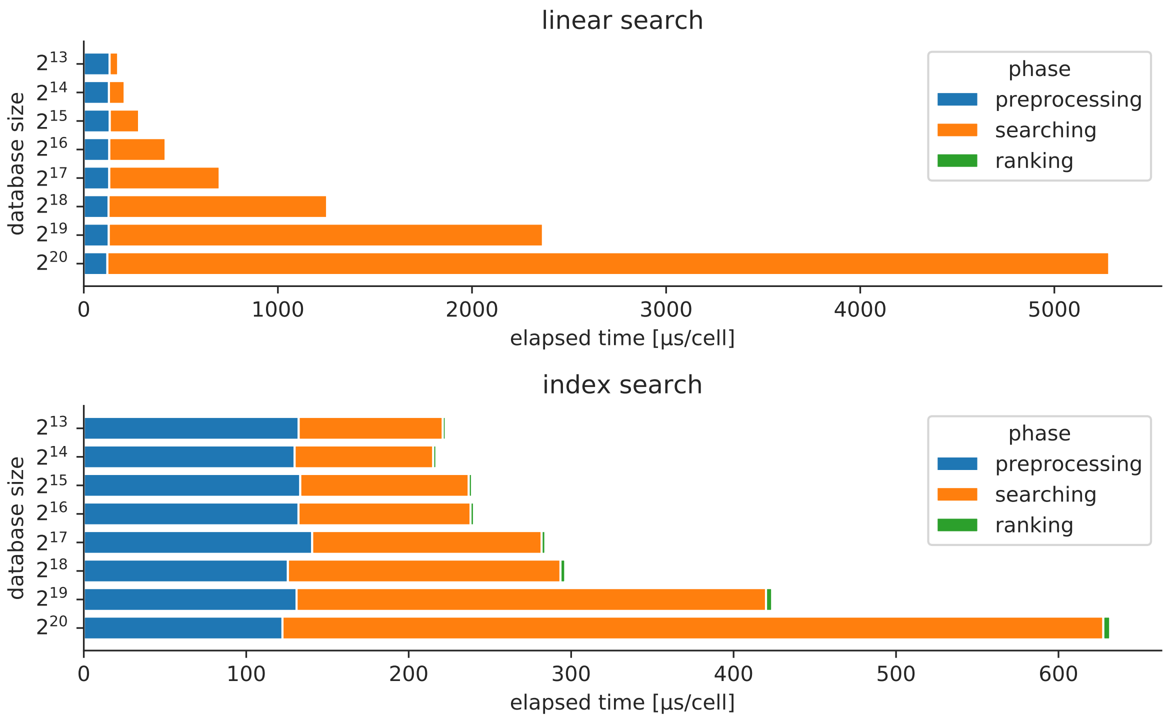

Figure 52: Computational costs of different phases (1M_neurons). Please refer to the main manuscript for the detailed caption.
