## Supplementary material for "CellFishing.jl: an ultrafast and scalable cell search method for single-cell RNA-sequencing"

Analysis


### Performance Analysis of CellFishing.jl¶

#### Table of contents¶

- Table of contents
- Cell-type distributions
- Randomized SVD
- Similarity estimation via hashing
- Visualization of expression profiles
- Self-mapping experiments
- Feature selection
- Cluster-specific scores
- Similarities from nearest neighbors
- Detecting DEGs
- Mapping across batches
- Mapping across species
- Mapping across protocols
- Benchmarks of saving and loading
- Scalability

In [1]:

```
import os
import re
import math
import textwrap
import toml
import numpy
import pandas
import scipy.stats
import sklearn.metrics
from matplotlib.pyplot import subplots
import matplotlib.pyplot
import matplotlib_venn
import seaborn
import umap
```

In [2]:

```
pkgname = "CellFishing"
```

In [3]:

```
ext = "pdf"
dpi = 200
seaborn.set_style("ticks", {"image.cmap": "viridis"})

def savefig(name):
    matplotlib.pyplot.savefig(os.path.join("figures", f"{name}.{ext}"), dpi="figure", bbox_inches="tight")
    
!mkdir -p figures
```

#### Cell-type distributions¶

In [4]:

```
datasets = ["Baron2016-human", "Shekhar2016", "Plass2018", "TabulaMuris-chromium"]
```

Load cell-type annotations:

In [5]:

```
clusters = dict()
for dataset in datasets:
    clusters[dataset] = pandas.read_table(f"data/{dataset}.cluster.tsv", index_col="cell")["cluster"]
clusters["Baron2016-mouse"] = pandas.read_table("data/Baron2016-mouse.cluster.tsv", index_col="cell")["cluster"]
clusters["Baron2016-human"][clusters["Baron2016-human"] == "t_cell"] = "T_cell"  # fix the name to match that of mouse
clusters["TabulaMuris-smart"] = pandas.read_table("data/TabulaMuris-smart.cluster.tsv", index_col="cell")["cluster"]
clusters["1M_neurons"] = pandas.read_table("data/1M_neurons.cluster.tsv", index_col="cell")["cluster"]
```

Summary of datasets:

In [6]:

```
print("{:20s}  {:>9}  {:>9s}".format("dataset", "#cells", "#clusters"))
print("------------------------------------------")
for dataset, cluster in sorted(clusters.items()):
    print("{:20s}  {:>9,d}  {:>9,d}".format(dataset, len(cluster), len(cluster.unique())))
```

```
dataset                  #cells  #clusters
------------------------------------------
1M_neurons            1,306,127         60
Baron2016-human           8,569         14
Baron2016-mouse           1,886         13
Plass2018                21,612         51
Shekhar2016              27,499         19
TabulaMuris-chromium     54,967         57
TabulaMuris-smart        44,807         81
```

Cluster (cell type) distributions:

In [7]:

```
for dataset, cluster in clusters.items():
    print(dataset)
    fig, ax = subplots(dpi=dpi, figsize=(8, 10))
    counts = cluster.value_counts()
    seaborn.countplot(
        y="cluster",
        data=pandas.DataFrame(cluster),
        order=counts.index.values,
        color="C0",
        ax=ax)
    fontsize = 6.5
    for i, c in enumerate(counts.index.values):
        ax.text(counts[c], i, " {:,} ({:.2f}%)".format(counts[c], counts[c] / len(cluster) * 100),
                verticalalignment="center",
                fontsize=fontsize)
    ax.tick_params(labelsize=fontsize)
    ax.set_xlabel("cluster size")
    seaborn.despine(fig=fig)
    savefig(f"{dataset}-clusters")
```

```
Baron2016-human
Shekhar2016
Plass2018
TabulaMuris-chromium
Baron2016-mouse
TabulaMuris-smart
1M_neurons
```

#### Randomized SVD¶

Load benchmark results:

In [8]:

```
def expand_singular_values(data):
    data = pandas.concat([
        pandas.DataFrame({"k": numpy.arange(1, r["n-dims"]+1), **dict(r)}) for ix, r in data.iterrows()], ignore_index=True)
    return data.rename(columns={"singular-values": "singular-value"})

svd = pandas.DataFrame()
for dataset in datasets:
    print(dataset)
    out = toml.load(f"results/{dataset}.svd.toml")
    print(out["date-time"])
    print(out["version-info"])
    expanded = expand_singular_values(pandas.DataFrame(out["experiment"]))
    expanded["dataset"] = dataset
    svd = svd.append(expanded, ignore_index=True)
```

```
Baron2016-human
2018-11-23 21:37:08.833000
Julia Version 1.0.1
Commit 0d713926f8 (2018-09-29 19:05 UTC)
Platform Info:
  OS: Linux (x86_64-pc-linux-gnu)
  CPU: Intel(R) Xeon(R) Gold 6126 CPU @ 2.60GHz
  WORD_SIZE: 64
  LIBM: libopenlibm
  LLVM: libLLVM-6.0.0 (ORCJIT, skylake)
Environment:
  JULIA_PROJECT = @.

Shekhar2016
2018-11-23 19:12:13.099000
Julia Version 1.0.1
Commit 0d713926f8 (2018-09-29 19:05 UTC)
Platform Info:
  OS: Linux (x86_64-pc-linux-gnu)
  CPU: Intel(R) Xeon(R) Gold 6126 CPU @ 2.60GHz
  WORD_SIZE: 64
  LIBM: libopenlibm
  LLVM: libLLVM-6.0.0 (ORCJIT, skylake)
Environment:
  JULIA_PROJECT = @.

Plass2018
2018-11-23 20:57:13.127000
Julia Version 1.0.1
Commit 0d713926f8 (2018-09-29 19:05 UTC)
Platform Info:
  OS: Linux (x86_64-pc-linux-gnu)
  CPU: Intel(R) Xeon(R) Gold 6126 CPU @ 2.60GHz
  WORD_SIZE: 64
  LIBM: libopenlibm
  LLVM: libLLVM-6.0.0 (ORCJIT, skylake)
Environment:
  JULIA_PROJECT = @.

TabulaMuris-chromium
2018-11-23 14:25:35.621000
Julia Version 1.0.1
Commit 0d713926f8 (2018-09-29 19:05 UTC)
Platform Info:
  OS: Linux (x86_64-pc-linux-gnu)
  CPU: Intel(R) Xeon(R) Gold 6126 CPU @ 2.60GHz
  WORD_SIZE: 64
  LIBM: libopenlibm
  LLVM: libLLVM-6.0.0 (ORCJIT, skylake)
Environment:
  JULIA_PROJECT = @.
```

Matrix sizes:

In [9]:

```
svd.groupby(["dataset"]).first()[["n-rows", "n-columns"]].loc[datasets,:]
```

Out[9]:

|  | n-rows | n-columns |
| --- | --- | --- |
| dataset |  |  |
| Baron2016-human | 2190 | 8569 |
| Shekhar2016 | 3270 | 27499 |
| Plass2018 | 3099 | 21612 |
| TabulaMuris-chromium | 2363 | 54967 |

Summary of benchmarks:

In [10]:

```
svd[svd["k"]==1].groupby(["dataset", "algorithm"]).describe()["time-svd"].loc[datasets,:]
```

Out[10]:

|  |  | count | mean | std | min | 25% | 50% | 75% | max |
| --- | --- | --- | --- | --- | --- | --- | --- | --- | --- |
| dataset | algorithm |  |  |  |  |  |  |  |  |
| Baron2016-human | full | 10.0 | 6.063345 | 0.183665 | 5.784003 | 5.929089 | 6.079253 | 6.225874 | 6.272859 |
|  | rand | 10.0 | 0.292286 | 0.062891 | 0.258365 | 0.268265 | 0.277413 | 0.278533 | 0.469917 |
|  | trunc | 10.0 | 1.173770 | 0.633607 | 0.958953 | 0.963400 | 0.976522 | 0.986208 | 2.976769 |
| Plass2018 | full | 10.0 | 24.281695 | 1.807840 | 22.876864 | 23.276733 | 24.009289 | 24.280506 | 29.146024 |
|  | rand | 10.0 | 0.841342 | 0.089431 | 0.773922 | 0.780410 | 0.820806 | 0.859711 | 1.070076 |
|  | trunc | 10.0 | 3.556973 | 0.580719 | 3.264094 | 3.290043 | 3.427950 | 3.466069 | 5.191450 |
| Shekhar2016 | full | 10.0 | 44.436990 | 9.126165 | 31.332782 | 37.777880 | 42.029988 | 54.116107 | 55.135777 |
|  | rand | 10.0 | 1.509863 | 0.058816 | 1.439421 | 1.496598 | 1.507051 | 1.512574 | 1.658688 |
|  | trunc | 10.0 | 9.396431 | 1.240010 | 8.693120 | 8.863809 | 8.934437 | 9.211806 | 12.777309 |
| TabulaMuris-chromium | full | 10.0 | 36.283684 | 2.311451 | 33.881575 | 35.119018 | 35.643392 | 36.825752 | 42.112005 |
|  | rand | 10.0 | 1.701845 | 0.080212 | 1.610664 | 1.649955 | 1.696022 | 1.729453 | 1.885414 |
|  | trunc | 10.0 | 5.742533 | 0.655307 | 5.380400 | 5.424111 | 5.487502 | 5.614463 | 7.494786 |

Plot the elapsed time:

In [11]:

```
fig, ax = subplots(dpi=dpi, figsize=(8, 4))
seaborn.swarmplot(x="dataset", y="time-svd", hue="algorithm", data=svd[svd["k"]==1], dodge=True, size=3)
ax.set_ylabel("elapsed time [s]")
ax.set_xlabel("")
ax.set_ylim(0, None)
ax.locator_params(nbins=5)
ax.grid(axis="y", alpha=0.3)
seaborn.despine(fig=fig)
fig.tight_layout()
savefig(f"svd-benchmark")
```

Plot the singular values:

In [12]:

```
fig, axes = subplots(4, sharex=True, dpi=dpi, figsize=(8, 4))
for i, dataset in enumerate(datasets):
    ax = axes[i]
    seaborn.pointplot(x="k", y="singular-value", hue="algorithm", data=svd[svd["dataset"]==dataset], scale=0.5, ci="sd", ax=ax)
    if i == 3:
        ax.set_xlabel("rank")
    else:
        ax.set_xlabel("")
    ax.set_ylim(0, None)
    for k, label in enumerate(ax.xaxis.get_ticklabels()):
        if k % 2 == 1:
            label.set_visible(False)
    ax.legend_.remove()
    ax.locator_params(axis="y", nbins=1)
    ax.set_ylabel("$\sigma$")
    ax.set_title(dataset)
    ax.grid(axis="y", alpha=0.3)
seaborn.despine(fig=fig)
fig.tight_layout()
savefig(f"svd-singular-values");
```

Compute the relative errors of the singular values:

In [13]:

```
svals = svd[svd["algorithm"]=="full"]["singular-value"].values
relerror = svd[svd["algorithm"]=="rand"]\
    .assign(**{"relative-error": lambda x: numpy.abs(1 - x["singular-value"] / svals)})
```

Plot the relative errors:

In [14]:

```
fig, axes = subplots(4, sharex=True, dpi=dpi, figsize=(8, 4))
for i, dataset in enumerate(datasets):
    ax = axes[i]
    seaborn.pointplot(x="k", y="relative-error", data=relerror[relerror["dataset"]==dataset], scale=0.5, errwidth=1.0, ci="sd", ax=ax)
    if i == 3:
        ax.set_xlabel("rank")
    else:
        ax.set_xlabel("")
    ax.set_ylim(0, None)
    for k, label in enumerate(ax.xaxis.get_ticklabels()):
        if k % 2 == 1:
            label.set_visible(False)
    ax.set_ylabel("rel. error")
    ax.set_title(dataset)
    ax.grid(axis="y", alpha=0.3)
seaborn.despine(fig=fig)
fig.tight_layout()
savefig(f"svd-relative-errors");
```

#### Similarity estimation via hashing¶

In [15]:

```
print("{:20s}  {:>6s}  {:>8s}  {:>7s}  {:>6s}".format("dataset", "n-bits", "superbit", "mean", "std"))
for dataset in datasets:
    fig, axes = subplots(4, dpi=dpi, figsize=(8, 5), sharex=True)
    for i, n_bits in enumerate([64, 128, 256, 512]):
        ax = axes[i]
        ax.set_title(f"n-bits = {n_bits}")
        ax.set_xlim(-0.75, +0.75)
        for j, superbit in enumerate([1, 50]):
            for r in range(1, 6):
                cosdist = numpy.loadtxt(f"results/{dataset}.estimator/n_bits-{n_bits}.superbit-{superbit}.{r}.cos.tsv.gz")
                angle = numpy.arccos(-cosdist+1)
                hamdist = numpy.loadtxt(f"results/{dataset}.estimator/n_bits-{n_bits}.superbit-{superbit}.{r}.ham.tsv.gz")
                approx = hamdist * math.pi / n_bits
                values = (approx - angle).ravel()
                print("{:20s}  {:>6d}  {:>8d}  {:>+.4f}  {:>.4f}".format(dataset, n_bits, superbit, numpy.mean(values), numpy.std(values)))
                kde = scipy.stats.gaussian_kde(values)
                x = numpy.linspace(values.min(), values.max(), num=200)
                ax.plot(x, kde(x), lw=1, color=f"C{j}")
    axes[-1].set_xlabel("estimation error [rad]")
    seaborn.despine(fig=fig)
    fig.tight_layout()
    savefig(f"{dataset}-estimator")
```

```
dataset               n-bits  superbit     mean     std
Baron2016-human           64         1  +0.0029  0.1907
Baron2016-human           64         1  -0.0000  0.1905
Baron2016-human           64         1  +0.0046  0.1941
Baron2016-human           64         1  -0.0084  0.1922
Baron2016-human           64         1  -0.0089  0.1849
Baron2016-human           64        50  -0.0007  0.1541
Baron2016-human           64        50  -0.0060  0.1644
Baron2016-human           64        50  -0.0033  0.1600
Baron2016-human           64        50  -0.0009  0.1571
Baron2016-human           64        50  -0.0040  0.1599
Baron2016-human          128         1  +0.0064  0.1327
Baron2016-human          128         1  -0.0020  0.1293
Baron2016-human          128         1  +0.0040  0.1368
Baron2016-human          128         1  +0.0000  0.1478
Baron2016-human          128         1  -0.0018  0.1430
Baron2016-human          128        50  +0.0030  0.1078
Baron2016-human          128        50  -0.0055  0.1109
Baron2016-human          128        50  -0.0004  0.1065
Baron2016-human          128        50  -0.0018  0.1106
Baron2016-human          128        50  +0.0002  0.1143
Baron2016-human          256         1  +0.0042  0.0949
Baron2016-human          256         1  +0.0013  0.0923
Baron2016-human          256         1  +0.0079  0.0967
Baron2016-human          256         1  -0.0012  0.0923
Baron2016-human          256         1  +0.0013  0.0947
Baron2016-human          256        50  +0.0018  0.0745
Baron2016-human          256        50  -0.0022  0.0753
Baron2016-human          256        50  +0.0008  0.0756
Baron2016-human          256        50  -0.0016  0.0763
Baron2016-human          256        50  -0.0005  0.0776
Baron2016-human          512         1  +0.0043  0.0710
Baron2016-human          512         1  -0.0038  0.0660
Baron2016-human          512         1  -0.0002  0.0647
Baron2016-human          512         1  +0.0015  0.0680
Baron2016-human          512         1  -0.0028  0.0668
Baron2016-human          512        50  +0.0002  0.0518
Baron2016-human          512        50  -0.0010  0.0523
Baron2016-human          512        50  +0.0016  0.0529
Baron2016-human          512        50  -0.0007  0.0535
Baron2016-human          512        50  +0.0005  0.0530
Shekhar2016               64         1  +0.0046  0.1977
Shekhar2016               64         1  -0.0031  0.1908
Shekhar2016               64         1  +0.0021  0.1853
Shekhar2016               64         1  -0.0054  0.1965
Shekhar2016               64         1  +0.0093  0.1889
Shekhar2016               64        50  +0.0054  0.1544
Shekhar2016               64        50  -0.0037  0.1542
Shekhar2016               64        50  +0.0048  0.1601
Shekhar2016               64        50  +0.0014  0.1583
Shekhar2016               64        50  +0.0074  0.1576
Shekhar2016              128         1  +0.0034  0.1288
Shekhar2016              128         1  +0.0030  0.1348
Shekhar2016              128         1  +0.0043  0.1372
Shekhar2016              128         1  -0.0049  0.1403
Shekhar2016              128         1  +0.0015  0.1355
Shekhar2016              128        50  +0.0027  0.1089
Shekhar2016              128        50  +0.0018  0.1069
Shekhar2016              128        50  -0.0002  0.1103
Shekhar2016              128        50  +0.0011  0.1104
Shekhar2016              128        50  +0.0058  0.1082
Shekhar2016              256         1  -0.0019  0.0944
Shekhar2016              256         1  -0.0008  0.0931
Shekhar2016              256         1  -0.0026  0.0948
Shekhar2016              256         1  -0.0020  0.0953
Shekhar2016              256         1  +0.0005  0.0963
Shekhar2016              256        50  +0.0023  0.0760
Shekhar2016              256        50  +0.0017  0.0749
Shekhar2016              256        50  -0.0018  0.0734
Shekhar2016              256        50  +0.0031  0.0752
Shekhar2016              256        50  +0.0004  0.0747
Shekhar2016              512         1  +0.0013  0.0681
Shekhar2016              512         1  +0.0017  0.0655
Shekhar2016              512         1  -0.0008  0.0644
Shekhar2016              512         1  +0.0014  0.0713
Shekhar2016              512         1  +0.0003  0.0663
Shekhar2016              512        50  +0.0016  0.0549
Shekhar2016              512        50  +0.0004  0.0528
Shekhar2016              512        50  -0.0017  0.0512
Shekhar2016              512        50  +0.0008  0.0525
Shekhar2016              512        50  +0.0010  0.0532
Plass2018                 64         1  -0.0036  0.1834
Plass2018                 64         1  -0.0031  0.1910
Plass2018                 64         1  +0.0049  0.1922
Plass2018                 64         1  +0.0091  0.1912
Plass2018                 64         1  +0.0015  0.2011
Plass2018                 64        50  +0.0070  0.1581
Plass2018                 64        50  +0.0036  0.1558
Plass2018                 64        50  +0.0007  0.1546
Plass2018                 64        50  -0.0022  0.1556
Plass2018                 64        50  -0.0017  0.1577
Plass2018                128         1  -0.0014  0.1336
Plass2018                128         1  +0.0020  0.1316
Plass2018                128         1  -0.0004  0.1391
Plass2018                128         1  +0.0005  0.1462
Plass2018                128         1  +0.0059  0.1403
Plass2018                128        50  +0.0009  0.1088
Plass2018                128        50  -0.0054  0.1066
Plass2018                128        50  -0.0031  0.1112
Plass2018                128        50  -0.0013  0.1091
Plass2018                128        50  -0.0012  0.1096
Plass2018                256         1  -0.0046  0.0941
Plass2018                256         1  +0.0021  0.0929
Plass2018                256         1  -0.0007  0.0975
Plass2018                256         1  +0.0040  0.0917
Plass2018                256         1  -0.0014  0.0920
Plass2018                256        50  -0.0002  0.0773
Plass2018                256        50  +0.0018  0.0744
Plass2018                256        50  -0.0010  0.0768
Plass2018                256        50  -0.0008  0.0747
Plass2018                256        50  -0.0007  0.0764
Plass2018                512         1  -0.0026  0.0701
Plass2018                512         1  +0.0032  0.0720
Plass2018                512         1  +0.0001  0.0671
Plass2018                512         1  -0.0014  0.0650
Plass2018                512         1  -0.0023  0.0673
Plass2018                512        50  -0.0001  0.0544
Plass2018                512        50  -0.0007  0.0550
Plass2018                512        50  -0.0016  0.0533
Plass2018                512        50  -0.0015  0.0524
Plass2018                512        50  -0.0017  0.0540
TabulaMuris-chromium      64         1  -0.0004  0.1809
TabulaMuris-chromium      64         1  +0.0010  0.2118
TabulaMuris-chromium      64         1  +0.0035  0.1931
TabulaMuris-chromium      64         1  +0.0052  0.1957
TabulaMuris-chromium      64         1  +0.0082  0.1945
TabulaMuris-chromium      64        50  -0.0038  0.1595
TabulaMuris-chromium      64        50  -0.0048  0.1534
TabulaMuris-chromium      64        50  +0.0022  0.1546
TabulaMuris-chromium      64        50  +0.0056  0.1529
TabulaMuris-chromium      64        50  -0.0003  0.1551
TabulaMuris-chromium     128         1  -0.0014  0.1289
TabulaMuris-chromium     128         1  -0.0007  0.1356
TabulaMuris-chromium     128         1  +0.0031  0.1434
TabulaMuris-chromium     128         1  -0.0033  0.1362
TabulaMuris-chromium     128         1  +0.0052  0.1385
TabulaMuris-chromium     128        50  -0.0017  0.1089
TabulaMuris-chromium     128        50  -0.0013  0.1085
TabulaMuris-chromium     128        50  -0.0026  0.1089
TabulaMuris-chromium     128        50  +0.0017  0.1097
TabulaMuris-chromium     128        50  +0.0021  0.1075
TabulaMuris-chromium     256         1  -0.0013  0.0936
TabulaMuris-chromium     256         1  -0.0012  0.0929
TabulaMuris-chromium     256         1  -0.0036  0.0951
TabulaMuris-chromium     256         1  -0.0024  0.0974
TabulaMuris-chromium     256         1  +0.0005  0.0940
TabulaMuris-chromium     256        50  +0.0002  0.0770
TabulaMuris-chromium     256        50  +0.0018  0.0724
TabulaMuris-chromium     256        50  +0.0007  0.0773
TabulaMuris-chromium     256        50  -0.0029  0.0790
TabulaMuris-chromium     256        50  +0.0002  0.0726
TabulaMuris-chromium     512         1  -0.0014  0.0683
TabulaMuris-chromium     512         1  +0.0010  0.0660
TabulaMuris-chromium     512         1  -0.0025  0.0655
TabulaMuris-chromium     512         1  +0.0010  0.0621
TabulaMuris-chromium     512         1  -0.0003  0.0699
TabulaMuris-chromium     512        50  -0.0013  0.0535
TabulaMuris-chromium     512        50  +0.0001  0.0523
TabulaMuris-chromium     512        50  -0.0009  0.0521
TabulaMuris-chromium     512        50  -0.0012  0.0525
TabulaMuris-chromium     512        50  +0.0009  0.0563
```

In [16]:

```
def n_positives(hamdist, cosdist, k):
    hamord = numpy.argsort(hamdist, axis=1)[:,0:k+1]
    cosord = numpy.argsort(cosdist, axis=1)[:,1]
    n = 0
    for i in range(hamord.shape[0]):
        n += numpy.sum(numpy.isin(cosord[i], hamord[i]))
    return n

n_cells = 6
x_max = 0.3  # upper limit of cosine similarity
print("{:20s}  {:>6s}  {:>2s}  {:>11s}".format("dataset", "n-bits", "k", "n-positives"))
for dataset in datasets:
    fig, axes = subplots(4, n_cells, dpi=dpi, figsize=(8, 5), sharex=True, sharey=True)
    for i, n_bits in enumerate([64, 128, 256, 512]):
        cosdist = numpy.loadtxt(f"results/{dataset}.estimator/n_bits-{n_bits}.superbit-50.1.cos.tsv.gz")
        hamdist = numpy.loadtxt(f"results/{dataset}.estimator/n_bits-{n_bits}.superbit-50.1.ham.tsv.gz")
        for k in [10, 30, 50]:
            print("{:20s}  {:>6d}  {:>2d}  {:>11d}".format(dataset, n_bits, k, n_positives(hamdist, cosdist, k)))
        axes[i,0].set_ylabel(f"n-bits = {n_bits}\n\nHamming dist.")
        for j in range(n_cells):
            ax = axes[i,j]
            x = cosdist[j,:]
            y = hamdist[j,:] / n_bits
            ax.scatter(x[x<x_max], y[x<x_max], s=0.3, lw=0, c="C1")
            ax.set_xlim(0, x_max)
            ax.set_ylim(0, None)
    for j in range(n_cells):    
        axes[3,j].set_xlabel("cosine dist.")
    seaborn.despine(fig=fig)
    fig.tight_layout(pad=0.5)
    savefig(f"{dataset}-correlation")
```

```
dataset               n-bits   k  n-positives
Baron2016-human           64  10           47
Baron2016-human           64  30           67
Baron2016-human           64  50           81
Baron2016-human          128  10           76
Baron2016-human          128  30           90
Baron2016-human          128  50           95
Baron2016-human          256  10           86
Baron2016-human          256  30           99
Baron2016-human          256  50          100
Baron2016-human          512  10           93
Baron2016-human          512  30          100
Baron2016-human          512  50          100
Shekhar2016               64  10           28
Shekhar2016               64  30           44
Shekhar2016               64  50           52
Shekhar2016              128  10           42
Shekhar2016              128  30           62
Shekhar2016              128  50           76
Shekhar2016              256  10           72
Shekhar2016              256  30           86
Shekhar2016              256  50           90
Shekhar2016              512  10           85
Shekhar2016              512  30           96
Shekhar2016              512  50           99
Plass2018                 64  10           36
Plass2018                 64  30           58
Plass2018                 64  50           67
Plass2018                128  10           62
Plass2018                128  30           78
Plass2018                128  50           86
Plass2018                256  10           79
Plass2018                256  30           91
Plass2018                256  50           94
Plass2018                512  10           87
Plass2018                512  30           98
Plass2018                512  50          100
TabulaMuris-chromium      64  10           33
TabulaMuris-chromium      64  30           46
TabulaMuris-chromium      64  50           50
TabulaMuris-chromium     128  10           47
TabulaMuris-chromium     128  30           66
TabulaMuris-chromium     128  50           74
TabulaMuris-chromium     256  10           65
TabulaMuris-chromium     256  30           84
TabulaMuris-chromium     256  50           89
TabulaMuris-chromium     512  10           82
TabulaMuris-chromium     512  30           95
TabulaMuris-chromium     512  50           98
```

#### Visualization of expression profiles¶

cell types and query/database plots

In [17]:

```
def readlines(filename):
    with open(filename) as f:
        return f.read().splitlines()

cluster = clusters["Shekhar2016"]
tab20 = matplotlib.cm.get_cmap("tab20")
for query in ["1", "2"]:
    print(f"query batch: {query}")
    log = toml.load(f"results/Shekhar2016.hash.batch-{query}.toml")
    print(log["date-time"])
    print(log["version-info"])
    outdir = f"results/Shekhar2016.hash.batch-{query}"
    for metric in ["euclidean", "cosine"]:
        fig, axes = subplots(2, 2, dpi=dpi, figsize=(8, 7))
        for ax in axes.ravel():
            ax.xaxis.set_ticklabels([])
            ax.yaxis.set_ticklabels([])
        for i, project in enumerate(["false", "true"]):
            cells_query = readlines(f"{outdir}/cells.query.project-{project}.txt")
            X_query = numpy.loadtxt(f"{outdir}/X.query.project-{project}.tsv").T
            cells_database = readlines(f"{outdir}/cells.database.project-{project}.txt")
            X_database = numpy.loadtxt(f"{outdir}/X.database.project-{project}.tsv").T
            X = numpy.vstack([X_query, X_database])
            X_umap = umap.UMAP(metric=metric, random_state=1234).fit_transform(X)
            shuf = numpy.arange(X_umap.shape[0])
            numpy.random.RandomState(1234).shuffle(shuf)
            labels = cluster[cells_query + cells_database]
            label2color = {label: tab20(k) for k, label in enumerate(numpy.sort(labels.unique()))}
            color = numpy.array([label2color[label] for label in labels])
            axes[i,0].scatter(X_umap[shuf,0], X_umap[shuf,1], s=0.5, lw=0, c=color[shuf])
            color = numpy.array(["C1"] * len(cells_query) + ["C0"] * len(cells_database))
            axes[i,1].scatter(X_umap[shuf,0], X_umap[shuf,1], s=0.5, lw=0, c=color[shuf])
        axes[0,0].set_title("cell type")
        axes[0,1].set_title("query/database")
        axes[0,0].set_ylabel("projected = false")
        axes[1,0].set_ylabel("projected = true")
        #fig.suptitle(f"query = {query}", va="bottom")
        seaborn.despine(fig=fig)
        fig.tight_layout(pad=0.5)
        savefig(f"Shekhar2016-hash-query-{query}-metric-{metric}")
```

```
query batch: 1
2018-11-23 16:45:49.993000
Julia Version 1.0.1
Commit 0d713926f8 (2018-09-29 19:05 UTC)
Platform Info:
  OS: Linux (x86_64-pc-linux-gnu)
  CPU: Intel(R) Xeon(R) Gold 6126 CPU @ 2.60GHz
  WORD_SIZE: 64
  LIBM: libopenlibm
  LLVM: libLLVM-6.0.0 (ORCJIT, skylake)
Environment:
  JULIA_PROJECT = @.
```

```
/home/ksato/.local/share/virtualenvs/cellfishing-experiments-YKL5dw-d/lib/python3.6/site-packages/umap/spectral.py:229: UserWarning: Embedding a total of 3 separate connected components using meta-embedding (experimental)
  n_components
/home/ksato/.local/share/virtualenvs/cellfishing-experiments-YKL5dw-d/lib/python3.6/site-packages/umap/spectral.py:229: UserWarning: Embedding a total of 2 separate connected components using meta-embedding (experimental)
  n_components
```

```
query batch: 2
2018-11-23 16:30:31.296000
Julia Version 1.0.1
Commit 0d713926f8 (2018-09-29 19:05 UTC)
Platform Info:
  OS: Linux (x86_64-pc-linux-gnu)
  CPU: Intel(R) Xeon(R) Gold 6126 CPU @ 2.60GHz
  WORD_SIZE: 64
  LIBM: libopenlibm
  LLVM: libLLVM-6.0.0 (ORCJIT, skylake)
Environment:
  JULIA_PROJECT = @.
```

```
/home/ksato/.local/share/virtualenvs/cellfishing-experiments-YKL5dw-d/lib/python3.6/site-packages/umap/spectral.py:229: UserWarning: Embedding a total of 2 separate connected components using meta-embedding (experimental)
  n_components
```

In [18]:

```
%load_ext Cython
```

In [19]:

```
%%cython

import numpy
cimport numpy
cimport cython

cdef extern int __builtin_popcount(unsigned int)

@cython.boundscheck(False)
@cython.wraparound(False)
cdef distance_matrix(numpy.ndarray[numpy.uint8_t,ndim=1] rows,
                     numpy.ndarray[numpy.uint8_t,ndim=1] cols,
                     int nbytes,
                     numpy.ndarray[numpy.uint16_t,ndim=2] output):
    cdef int i, j, k
    cdef int d
    cdef int m = rows.shape[0] // nbytes
    cdef int n = cols.shape[0] // nbytes
    for i in range(m):
        for j in range(n):
            d = 0
            for k in range(nbytes):
                d += __builtin_popcount(rows[i*nbytes+k] ^ cols[j*nbytes+k])
            output[i,j] = d

            
def hamming_distance_matrix(rows, cols, nbytes):
    assert rows.shape[0] % nbytes == 0
    assert cols.shape[0] % nbytes == 0
    m = rows.shape[0] // nbytes
    n = cols.shape[0] // nbytes
    output = numpy.zeros((m, n), dtype=numpy.uint16)
    distance_matrix(rows, cols, nbytes, output)
    return output
```

In [20]:

```
nbytes = 64 // 8
cluster = clusters["Shekhar2016"]
tab20 = matplotlib.cm.get_cmap("tab20")
n = len(cluster)
shuf = numpy.arange(n)
numpy.random.RandomState(1234).shuffle(shuf)
for query in ["1", "2"]:
    fig, axes = subplots(2, 2, dpi=dpi, figsize=(8, 8))
    axes = [axes[0,0], axes[0,1], axes[1,0], axes[1,1]]
    for ax in axes:
        ax.xaxis.set_ticklabels([])
        ax.yaxis.set_ticklabels([])
    outdir = f"results/Shekhar2016.hash.batch-{query}"
    cells_query = readlines(f"{outdir}/cells.query.project-true.txt")
    X_query = numpy.loadtxt(f"{outdir}/X.query.project-true.tsv").T
    cells_database = readlines(f"{outdir}/cells.database.project-true.txt")
    X_database = numpy.loadtxt(f"{outdir}/X.database.project-true.tsv").T
    X = numpy.vstack([X_query, X_database])
    X_umap = umap.UMAP(random_state=1234, metric="cosine").fit_transform(X)
    # plot
    labels = cluster[cells_query + cells_database]
    label2color = {label: tab20(k) for k, label in enumerate(numpy.sort(labels.unique()))}
    color = numpy.array([label2color[label] for label in labels])
    ax = axes.pop(0)
    ax.scatter(X_umap[shuf,0], X_umap[shuf,1], s=0.5, lw=0, c=color[shuf])
    ax.set_title("cosine distance (original)")
    # hashed profiles
    D = numpy.zeros((n, n), dtype=numpy.uint16)
    for l in range(4):
        # compute distance matrix
        Z_query = numpy.fromfile(f"{outdir}/Z.query.project-true.{l+1}.bin", dtype=numpy.uint8)
        Z_database = numpy.fromfile(f"{outdir}/Z.database.project-true.{l+1}.bin", dtype=numpy.uint8)
        Z = numpy.hstack([Z_query, Z_database])
        D += hamming_distance_matrix(Z, Z, nbytes)
        nbits = nbytes * (l+1) * 8
        if nbits in (64, 128, 256):
            D_umap = umap.UMAP(random_state=1234, metric="precomputed").fit_transform(D)         
            ax = axes.pop(0)
            ax.scatter(D_umap[shuf,0], D_umap[shuf,1], s=0.5, lw=0, c=color[shuf])
            ax.set_title(f"Hamming distance (n-bits = {nbits})")
    seaborn.despine(fig=fig)
    fig.tight_layout(pad=0.5)
    savefig(f"Shekhar2016-hash-query-{query}")
```

```
/home/ksato/.local/share/virtualenvs/cellfishing-experiments-YKL5dw-d/lib/python3.6/site-packages/umap/umap_.py:1429: UserWarning: Using precomputed metric; transform will be unavailable for new data
  warn('Using precomputed metric; transform will be unavailable for new data')
/home/ksato/.local/share/virtualenvs/cellfishing-experiments-YKL5dw-d/lib/python3.6/site-packages/umap/umap_.py:1429: UserWarning: Using precomputed metric; transform will be unavailable for new data
  warn('Using precomputed metric; transform will be unavailable for new data')
/home/ksato/.local/share/virtualenvs/cellfishing-experiments-YKL5dw-d/lib/python3.6/site-packages/umap/umap_.py:1429: UserWarning: Using precomputed metric; transform will be unavailable for new data
  warn('Using precomputed metric; transform will be unavailable for new data')
/home/ksato/.local/share/virtualenvs/cellfishing-experiments-YKL5dw-d/lib/python3.6/site-packages/umap/umap_.py:1429: UserWarning: Using precomputed metric; transform will be unavailable for new data
  warn('Using precomputed metric; transform will be unavailable for new data')
/home/ksato/.local/share/virtualenvs/cellfishing-experiments-YKL5dw-d/lib/python3.6/site-packages/umap/umap_.py:1429: UserWarning: Using precomputed metric; transform will be unavailable for new data
  warn('Using precomputed metric; transform will be unavailable for new data')
/home/ksato/.local/share/virtualenvs/cellfishing-experiments-YKL5dw-d/lib/python3.6/site-packages/umap/umap_.py:1429: UserWarning: Using precomputed metric; transform will be unavailable for new data
  warn('Using precomputed metric; transform will be unavailable for new data')
```

#### Self-mapping experiments¶

In [21]:

```
def load_knn(filename):
    return pandas.read_table(filename, index_col="cell")

# Compute the consistency, Cohen's kappa, and the adjusted Rand scores.
def cluster_metrics(filename, cluster, k=1, cluster_nn=None):
    if cluster_nn is None:
        cluster_nn = cluster
    knn = load_knn(filename)
    cell = cluster[knn.index].values
    nn = cluster_nn[knn[f"N{k}"]].values
    return dict(
        consistency=sklearn.metrics.accuracy_score(cell, nn),
        cohen_kappa=sklearn.metrics.cohen_kappa_score(cell, nn),
        adjusted_rand=sklearn.metrics.adjusted_rand_score(cell, nn),)
```

In [22]:

```
knn_cv = pandas.DataFrame()
knn_cv_scmap = pandas.DataFrame()
for dataset in datasets:
    print(dataset)
    # CellFishing
    log = toml.load(f"results/{dataset}.knn-cv.toml")
    print(log["date-time"])
    print(log["version-info"])
    exp = pandas.DataFrame(log["experiment"])
    exp["dataset"] = dataset
    exp = pandas.concat(
        [exp, pandas.DataFrame([cluster_metrics(x, clusters[dataset], k=1) for x in exp["filename"]])],
        axis=1)
    knn_cv = knn_cv.append(exp, ignore_index=True)
    # scmap
    log = toml.load(f"results/{dataset}.knn-cv.scmap.toml")
    print(log["date-time"])
    print(log["session-info"])
    exp = pandas.DataFrame(log["experiment"])
    exp["dataset"] = dataset
    exp = pandas.concat(
        [exp, pandas.DataFrame([cluster_metrics(x, clusters[dataset], k=1) for x in exp["filename"]])],
        axis=1)
    knn_cv_scmap = knn_cv_scmap.append(exp, ignore_index=True)
```

```
Baron2016-human
2018-11-23 21:30:48.455000
Julia Version 1.0.1
Commit 0d713926f8 (2018-09-29 19:05 UTC)
Platform Info:
  OS: Linux (x86_64-pc-linux-gnu)
  CPU: Intel(R) Xeon(R) Gold 6126 CPU @ 2.60GHz
  WORD_SIZE: 64
  LIBM: libopenlibm
  LLVM: libLLVM-6.0.0 (ORCJIT, skylake)
Environment:
  JULIA_PROJECT = @.

2018-07-09 06:09:06
R version 3.5.0 (2018-04-23)
Platform: x86_64-pc-linux-gnu (64-bit)
Running under: Debian GNU/Linux 9 (stretch)

Matrix products: default
BLAS: /usr/lib/openblas-base/libblas.so.3
LAPACK: /usr/lib/libopenblasp-r0.2.19.so

locale:
 [1] LC_CTYPE=en_US.UTF-8       LC_NUMERIC=C              
 [3] LC_TIME=en_US.UTF-8        LC_COLLATE=en_US.UTF-8    
 [5] LC_MONETARY=en_US.UTF-8    LC_MESSAGES=C             
 [7] LC_PAPER=en_US.UTF-8       LC_NAME=C                 
 [9] LC_ADDRESS=C               LC_TELEPHONE=C            
[11] LC_MEASUREMENT=en_US.UTF-8 LC_IDENTIFICATION=C       

attached base packages:
[1] parallel  stats4    stats     graphics  grDevices utils     datasets 
[8] methods   base     

other attached packages:
 [1] h5_0.9.9                    scmap_1.2.0                
 [3] scater_1.8.0                ggplot2_2.2.1              
 [5] SingleCellExperiment_1.2.0  SummarizedExperiment_1.10.1
 [7] DelayedArray_0.6.1          BiocParallel_1.14.1        
 [9] matrixStats_0.53.1          Biobase_2.40.0             
[11] GenomicRanges_1.32.3        GenomeInfoDb_1.16.0        
[13] IRanges_2.14.10             S4Vectors_0.18.3           
[15] BiocGenerics_0.26.0         readr_1.1.1                

loaded via a namespace (and not attached):
 [1] viridis_0.5.1            edgeR_3.22.2             jsonlite_1.5            
 [4] viridisLite_0.3.0        DelayedMatrixStats_1.2.0 shiny_1.1.0             
 [7] assertthat_0.2.0         GenomeInfoDbData_1.1.0   vipor_0.4.5             
[10] pillar_1.2.2             lattice_0.20-35          glue_1.2.0              
[13] limma_3.36.1             digest_0.6.15            promises_1.0.1          
[16] XVector_0.20.0           randomForest_4.6-14      colorspace_1.3-2        
[19] htmltools_0.3.6          httpuv_1.4.3             Matrix_1.2-14           
[22] plyr_1.8.4               pkgconfig_2.0.1          zlibbioc_1.26.0         
[25] purrr_0.2.5              xtable_1.8-2             scales_0.5.0            
[28] later_0.7.3              proxy_0.4-22             tibble_1.4.2            
[31] lazyeval_0.2.1           magrittr_1.5             mime_0.5                
[34] class_7.3-14             beeswarm_0.2.3           shinydashboard_0.7.0    
[37] tools_3.5.0              data.table_1.11.4        hms_0.4.2               
[40] stringr_1.3.0            googleVis_0.6.2          Rhdf5lib_1.2.1          
[43] munsell_0.5.0            locfit_1.5-9.1           bindrcpp_0.2.2          
[46] compiler_3.5.0           e1071_1.6-8              rlang_0.2.0             
[49] rhdf5_2.24.0             grid_3.5.0               RCurl_1.95-4.10         
[52] tximport_1.8.0           rjson_0.2.20             bitops_1.0-6            
[55] gtable_0.2.0             reshape2_1.4.3           R6_2.2.2                
[58] gridExtra_2.3            dplyr_0.7.5              bindr_0.1.1             
[61] stringi_1.2.2            ggbeeswarm_0.6.0         Rcpp_0.12.16            
[64] tidyselect_0.2.4        

Shekhar2016
2018-11-23 19:21:54.315000
Julia Version 1.0.1
Commit 0d713926f8 (2018-09-29 19:05 UTC)
Platform Info:
  OS: Linux (x86_64-pc-linux-gnu)
  CPU: Intel(R) Xeon(R) Gold 6126 CPU @ 2.60GHz
  WORD_SIZE: 64
  LIBM: libopenlibm
  LLVM: libLLVM-6.0.0 (ORCJIT, skylake)
Environment:
  JULIA_PROJECT = @.

2018-07-09 00:24:09
R version 3.5.0 (2018-04-23)
Platform: x86_64-pc-linux-gnu (64-bit)
Running under: Debian GNU/Linux 9 (stretch)

Matrix products: default
BLAS: /usr/lib/openblas-base/libblas.so.3
LAPACK: /usr/lib/libopenblasp-r0.2.19.so

locale:
 [1] LC_CTYPE=en_US.UTF-8       LC_NUMERIC=C              
 [3] LC_TIME=en_US.UTF-8        LC_COLLATE=en_US.UTF-8    
 [5] LC_MONETARY=en_US.UTF-8    LC_MESSAGES=C             
 [7] LC_PAPER=en_US.UTF-8       LC_NAME=C                 
 [9] LC_ADDRESS=C               LC_TELEPHONE=C            
[11] LC_MEASUREMENT=en_US.UTF-8 LC_IDENTIFICATION=C       

attached base packages:
[1] parallel  stats4    stats     graphics  grDevices utils     datasets 
[8] methods   base     

other attached packages:
 [1] h5_0.9.9                    scmap_1.2.0                
 [3] scater_1.8.0                ggplot2_2.2.1              
 [5] SingleCellExperiment_1.2.0  SummarizedExperiment_1.10.1
 [7] DelayedArray_0.6.1          BiocParallel_1.14.1        
 [9] matrixStats_0.53.1          Biobase_2.40.0             
[11] GenomicRanges_1.32.3        GenomeInfoDb_1.16.0        
[13] IRanges_2.14.10             S4Vectors_0.18.3           
[15] BiocGenerics_0.26.0         readr_1.1.1                

loaded via a namespace (and not attached):
 [1] viridis_0.5.1            edgeR_3.22.2             jsonlite_1.5            
 [4] viridisLite_0.3.0        DelayedMatrixStats_1.2.0 shiny_1.1.0             
 [7] assertthat_0.2.0         GenomeInfoDbData_1.1.0   vipor_0.4.5             
[10] pillar_1.2.2             lattice_0.20-35          glue_1.2.0              
[13] limma_3.36.1             digest_0.6.15            promises_1.0.1          
[16] XVector_0.20.0           randomForest_4.6-14      colorspace_1.3-2        
[19] htmltools_0.3.6          httpuv_1.4.3             Matrix_1.2-14           
[22] plyr_1.8.4               pkgconfig_2.0.1          zlibbioc_1.26.0         
[25] purrr_0.2.5              xtable_1.8-2             scales_0.5.0            
[28] later_0.7.3              proxy_0.4-22             tibble_1.4.2            
[31] lazyeval_0.2.1           magrittr_1.5             mime_0.5                
[34] class_7.3-14             beeswarm_0.2.3           shinydashboard_0.7.0    
[37] tools_3.5.0              data.table_1.11.4        hms_0.4.2               
[40] stringr_1.3.0            googleVis_0.6.2          Rhdf5lib_1.2.1          
[43] munsell_0.5.0            locfit_1.5-9.1           bindrcpp_0.2.2          
[46] compiler_3.5.0           e1071_1.6-8              rlang_0.2.0             
[49] rhdf5_2.24.0             grid_3.5.0               RCurl_1.95-4.10         
[52] tximport_1.8.0           rjson_0.2.20             bitops_1.0-6            
[55] gtable_0.2.0             reshape2_1.4.3           R6_2.2.2                
[58] gridExtra_2.3            dplyr_0.7.5              bindr_0.1.1             
[61] stringi_1.2.2            ggbeeswarm_0.6.0         Rcpp_0.12.16            
[64] tidyselect_0.2.4        

Plass2018
2018-11-23 20:32:08.025000
Julia Version 1.0.1
Commit 0d713926f8 (2018-09-29 19:05 UTC)
Platform Info:
  OS: Linux (x86_64-pc-linux-gnu)
  CPU: Intel(R) Xeon(R) Gold 6126 CPU @ 2.60GHz
  WORD_SIZE: 64
  LIBM: libopenlibm
  LLVM: libLLVM-6.0.0 (ORCJIT, skylake)
Environment:
  JULIA_PROJECT = @.

2018-07-09 04:38:07
R version 3.5.0 (2018-04-23)
Platform: x86_64-pc-linux-gnu (64-bit)
Running under: Debian GNU/Linux 9 (stretch)

Matrix products: default
BLAS: /usr/lib/openblas-base/libblas.so.3
LAPACK: /usr/lib/libopenblasp-r0.2.19.so

locale:
 [1] LC_CTYPE=en_US.UTF-8       LC_NUMERIC=C              
 [3] LC_TIME=en_US.UTF-8        LC_COLLATE=en_US.UTF-8    
 [5] LC_MONETARY=en_US.UTF-8    LC_MESSAGES=C             
 [7] LC_PAPER=en_US.UTF-8       LC_NAME=C                 
 [9] LC_ADDRESS=C               LC_TELEPHONE=C            
[11] LC_MEASUREMENT=en_US.UTF-8 LC_IDENTIFICATION=C       

attached base packages:
[1] parallel  stats4    stats     graphics  grDevices utils     datasets 
[8] methods   base     

other attached packages:
 [1] h5_0.9.9                    scmap_1.2.0                
 [3] scater_1.8.0                ggplot2_2.2.1              
 [5] SingleCellExperiment_1.2.0  SummarizedExperiment_1.10.1
 [7] DelayedArray_0.6.1          BiocParallel_1.14.1        
 [9] matrixStats_0.53.1          Biobase_2.40.0             
[11] GenomicRanges_1.32.3        GenomeInfoDb_1.16.0        
[13] IRanges_2.14.10             S4Vectors_0.18.3           
[15] BiocGenerics_0.26.0         readr_1.1.1                

loaded via a namespace (and not attached):
 [1] viridis_0.5.1            edgeR_3.22.2             jsonlite_1.5            
 [4] viridisLite_0.3.0        DelayedMatrixStats_1.2.0 shiny_1.1.0             
 [7] assertthat_0.2.0         GenomeInfoDbData_1.1.0   vipor_0.4.5             
[10] pillar_1.2.2             lattice_0.20-35          glue_1.2.0              
[13] limma_3.36.1             digest_0.6.15            promises_1.0.1          
[16] XVector_0.20.0           randomForest_4.6-14      colorspace_1.3-2        
[19] htmltools_0.3.6          httpuv_1.4.3             Matrix_1.2-14           
[22] plyr_1.8.4               pkgconfig_2.0.1          zlibbioc_1.26.0         
[25] purrr_0.2.5              xtable_1.8-2             scales_0.5.0            
[28] later_0.7.3              proxy_0.4-22             tibble_1.4.2            
[31] lazyeval_0.2.1           magrittr_1.5             mime_0.5                
[34] class_7.3-14             beeswarm_0.2.3           shinydashboard_0.7.0    
[37] tools_3.5.0              data.table_1.11.4        hms_0.4.2               
[40] stringr_1.3.0            googleVis_0.6.2          Rhdf5lib_1.2.1          
[43] munsell_0.5.0            locfit_1.5-9.1           bindrcpp_0.2.2          
[46] compiler_3.5.0           e1071_1.6-8              rlang_0.2.0             
[49] rhdf5_2.24.0             grid_3.5.0               RCurl_1.95-4.10         
[52] tximport_1.8.0           rjson_0.2.20             bitops_1.0-6            
[55] gtable_0.2.0             reshape2_1.4.3           R6_2.2.2                
[58] gridExtra_2.3            dplyr_0.7.5              bindr_0.1.1             
[61] stringi_1.2.2            ggbeeswarm_0.6.0         Rcpp_0.12.16            
[64] tidyselect_0.2.4        

TabulaMuris-chromium
2018-11-23 14:26:00.378000
Julia Version 1.0.1
Commit 0d713926f8 (2018-09-29 19:05 UTC)
Platform Info:
  OS: Linux (x86_64-pc-linux-gnu)
  CPU: Intel(R) Xeon(R) Gold 6126 CPU @ 2.60GHz
  WORD_SIZE: 64
  LIBM: libopenlibm
  LLVM: libLLVM-6.0.0 (ORCJIT, skylake)
Environment:
  JULIA_PROJECT = @.

2018-07-08 16:03:39
R version 3.5.0 (2018-04-23)
Platform: x86_64-pc-linux-gnu (64-bit)
Running under: Debian GNU/Linux 9 (stretch)

Matrix products: default
BLAS: /usr/lib/openblas-base/libblas.so.3
LAPACK: /usr/lib/libopenblasp-r0.2.19.so

locale:
 [1] LC_CTYPE=en_US.UTF-8       LC_NUMERIC=C              
 [3] LC_TIME=en_US.UTF-8        LC_COLLATE=en_US.UTF-8    
 [5] LC_MONETARY=en_US.UTF-8    LC_MESSAGES=C             
 [7] LC_PAPER=en_US.UTF-8       LC_NAME=C                 
 [9] LC_ADDRESS=C               LC_TELEPHONE=C            
[11] LC_MEASUREMENT=en_US.UTF-8 LC_IDENTIFICATION=C       

attached base packages:
[1] parallel  stats4    stats     graphics  grDevices utils     datasets 
[8] methods   base     

other attached packages:
 [1] h5_0.9.9                    scmap_1.2.0                
 [3] scater_1.8.0                ggplot2_2.2.1              
 [5] SingleCellExperiment_1.2.0  SummarizedExperiment_1.10.1
 [7] DelayedArray_0.6.1          BiocParallel_1.14.1        
 [9] matrixStats_0.53.1          Biobase_2.40.0             
[11] GenomicRanges_1.32.3        GenomeInfoDb_1.16.0        
[13] IRanges_2.14.10             S4Vectors_0.18.3           
[15] BiocGenerics_0.26.0         readr_1.1.1                

loaded via a namespace (and not attached):
 [1] viridis_0.5.1            edgeR_3.22.2             jsonlite_1.5            
 [4] viridisLite_0.3.0        DelayedMatrixStats_1.2.0 shiny_1.1.0             
 [7] assertthat_0.2.0         GenomeInfoDbData_1.1.0   vipor_0.4.5             
[10] pillar_1.2.2             lattice_0.20-35          glue_1.2.0              
[13] limma_3.36.1             digest_0.6.15            promises_1.0.1          
[16] XVector_0.20.0           randomForest_4.6-14      colorspace_1.3-2        
[19] htmltools_0.3.6          httpuv_1.4.3             Matrix_1.2-14           
[22] plyr_1.8.4               pkgconfig_2.0.1          zlibbioc_1.26.0         
[25] purrr_0.2.5              xtable_1.8-2             scales_0.5.0            
[28] later_0.7.3              proxy_0.4-22             tibble_1.4.2            
[31] lazyeval_0.2.1           magrittr_1.5             mime_0.5                
[34] class_7.3-14             beeswarm_0.2.3           shinydashboard_0.7.0    
[37] tools_3.5.0              data.table_1.11.4        hms_0.4.2               
[40] stringr_1.3.0            googleVis_0.6.2          Rhdf5lib_1.2.1          
[43] munsell_0.5.0            locfit_1.5-9.1           bindrcpp_0.2.2          
[46] compiler_3.5.0           e1071_1.6-8              rlang_0.2.0             
[49] rhdf5_2.24.0             grid_3.5.0               RCurl_1.95-4.10         
[52] tximport_1.8.0           rjson_0.2.20             bitops_1.0-6            
[55] gtable_0.2.0             reshape2_1.4.3           R6_2.2.2                
[58] gridExtra_2.3            dplyr_0.7.5              bindr_0.1.1             
[61] stringi_1.2.2            ggbeeswarm_0.6.0         Rcpp_0.12.16            
[64] tidyselect_0.2.4
```

Select columns:

In [23]:

```
knn_cv = knn_cv[["dataset", "n-bits", "n-lshashes", "transformer", "filename", "consistency", "cohen_kappa", "adjusted_rand", "index-time", "query-time"]]
knn_cv_scmap = knn_cv_scmap[["dataset", "n-features", "n-centroids-factor", "filename", "consistency", "cohen_kappa", "adjusted_rand", "index-time", "query-time"]]
```

Summary:

In [24]:

```
pandas.options.display.max_columns = 99
knn_cv.groupby(["dataset", "n-bits", "n-lshashes", "transformer"]).describe()
```

Out[24]:

|  |  |  |  | adjusted\_rand | | | | | | | | cohen\_kappa | | | | | | | | consistency | | | | | | | | index-time | | | | | | | | query-time | | | | | | | |
| --- | --- | --- | --- | --- | --- | --- | --- | --- | --- | --- | --- | --- | --- | --- | --- | --- | --- | --- | --- | --- | --- | --- | --- | --- | --- | --- | --- | --- | --- | --- | --- | --- | --- | --- | --- | --- | --- | --- | --- | --- | --- | --- | --- |
|  |  |  |  | count | mean | std | min | 25% | 50% | 75% | max | count | mean | std | min | 25% | 50% | 75% | max | count | mean | std | min | 25% | 50% | 75% | max | count | mean | std | min | 25% | 50% | 75% | max | count | mean | std | min | 25% | 50% | 75% | max |
| dataset | n-bits | n-lshashes | transformer |  |  |  |  |  |  |  |  |  |  |  |  |  |  |  |  |  |  |  |  |  |  |  |  |  |  |  |  |  |  |  |  |  |  |  |  |  |  |  |  |
| Baron2016-human | 128 | 1 | ftt | 5.0 | 0.964905 | 0.002258 | 0.961941 | 0.963399 | 0.965059 | 0.966987 | 0.967137 | 5.0 | 0.976453 | 0.001231 | 0.974998 | 0.975579 | 0.976305 | 0.977474 | 0.977908 | 5.0 | 0.981095 | 0.000987 | 0.979928 | 0.980394 | 0.980978 | 0.981912 | 0.982262 | 5.0 | 3.717403 | 0.045814 | 3.673193 | 3.675623 | 3.714009 | 3.743534 | 3.780658 | 5.0 | 0.298764 | 0.003785 | 0.295250 | 0.296407 | 0.296659 | 0.301495 | 0.304009 |
|  |  |  | log1p | 5.0 | 0.962464 | 0.001806 | 0.960971 | 0.961148 | 0.961333 | 0.964298 | 0.964572 | 5.0 | 0.974616 | 0.000720 | 0.973827 | 0.974266 | 0.974412 | 0.974846 | 0.975728 | 5.0 | 0.979624 | 0.000575 | 0.978994 | 0.979344 | 0.979461 | 0.979811 | 0.980511 | 5.0 | 5.232839 | 1.040440 | 4.716414 | 4.759946 | 4.764084 | 4.831166 | 7.092587 | 5.0 | 0.554049 | 0.272270 | 0.428423 | 0.430749 | 0.431977 | 0.438038 | 1.041060 |
|  |  | 4 | ftt | 5.0 | 0.971342 | 0.002018 | 0.968797 | 0.970296 | 0.970879 | 0.972983 | 0.973754 | 5.0 | 0.981135 | 0.000714 | 0.980524 | 0.980526 | 0.980815 | 0.981831 | 0.981978 | 5.0 | 0.984852 | 0.000574 | 0.984362 | 0.984362 | 0.984596 | 0.985413 | 0.985529 | 5.0 | 3.838014 | 0.035338 | 3.793426 | 3.823056 | 3.827917 | 3.861518 | 3.884155 | 5.0 | 0.778373 | 0.009219 | 0.763083 | 0.778702 | 0.780838 | 0.781311 | 0.787931 |
|  |  |  | log1p | 5.0 | 0.970294 | 0.002036 | 0.967999 | 0.968820 | 0.969931 | 0.971877 | 0.972843 | 5.0 | 0.979683 | 0.001083 | 0.978199 | 0.979217 | 0.979520 | 0.980670 | 0.980811 | 5.0 | 0.983685 | 0.000871 | 0.982495 | 0.983312 | 0.983545 | 0.984479 | 0.984596 | 5.0 | 4.867224 | 0.045700 | 4.823732 | 4.839330 | 4.841887 | 4.902086 | 4.929086 | 5.0 | 0.938610 | 0.005173 | 0.932786 | 0.936911 | 0.937307 | 0.939178 | 0.946868 |
|  |  | 16 | ftt | 5.0 | 0.972772 | 0.001556 | 0.971432 | 0.971629 | 0.972173 | 0.973455 | 0.975171 | 5.0 | 0.981660 | 0.000895 | 0.980962 | 0.981103 | 0.981254 | 0.981836 | 0.983146 | 5.0 | 0.985272 | 0.000717 | 0.984712 | 0.984829 | 0.984946 | 0.985413 | 0.986463 | 5.0 | 4.381316 | 0.043398 | 4.321394 | 4.361337 | 4.380621 | 4.409460 | 4.433768 | 5.0 | 2.610159 | 0.013708 | 2.595047 | 2.602713 | 2.606744 | 2.615437 | 2.630856 |
|  |  |  | log1p | 5.0 | 0.972885 | 0.001108 | 0.972035 | 0.972038 | 0.972350 | 0.973412 | 0.974590 | 5.0 | 0.981196 | 0.000687 | 0.980383 | 0.980814 | 0.980964 | 0.981837 | 0.981982 | 5.0 | 0.984899 | 0.000551 | 0.984246 | 0.984596 | 0.984712 | 0.985413 | 0.985529 | 5.0 | 5.386537 | 0.048024 | 5.328442 | 5.353350 | 5.392048 | 5.406621 | 5.452226 | 5.0 | 2.754696 | 0.013501 | 2.737475 | 2.744004 | 2.757912 | 2.765826 | 2.768264 |
|  | 256 | 1 | ftt | 5.0 | 0.968953 | 0.002446 | 0.965639 | 0.967725 | 0.969178 | 0.970098 | 0.972126 | 5.0 | 0.979277 | 0.001594 | 0.977190 | 0.978638 | 0.979217 | 0.979797 | 0.981542 | 5.0 | 0.983359 | 0.001283 | 0.981678 | 0.982845 | 0.983312 | 0.983779 | 0.985179 | 5.0 | 3.852656 | 0.154701 | 3.761866 | 3.782331 | 3.794249 | 3.796534 | 4.128300 | 5.0 | 0.472831 | 0.009288 | 0.459319 | 0.469158 | 0.475285 | 0.476047 | 0.484346 |
|  |  |  | log1p | 5.0 | 0.968391 | 0.002934 | 0.965174 | 0.966144 | 0.968186 | 0.970017 | 0.972433 | 5.0 | 0.978260 | 0.000892 | 0.977187 | 0.978057 | 0.978061 | 0.978340 | 0.979655 | 5.0 | 0.982542 | 0.000717 | 0.981678 | 0.982378 | 0.982378 | 0.982612 | 0.983662 | 5.0 | 4.842319 | 0.146106 | 4.757145 | 4.778473 | 4.783163 | 4.790062 | 5.102754 | 5.0 | 0.644718 | 0.096471 | 0.594669 | 0.601928 | 0.604552 | 0.605314 | 0.817127 |
|  |  | 4 | ftt | 5.0 | 0.972556 | 0.001324 | 0.971452 | 0.971940 | 0.972005 | 0.972569 | 0.974815 | 5.0 | 0.981515 | 0.000883 | 0.980816 | 0.980960 | 0.980961 | 0.981982 | 0.982857 | 5.0 | 0.985156 | 0.000707 | 0.984596 | 0.984712 | 0.984712 | 0.985529 | 0.986229 | 5.0 | 4.267599 | 0.104073 | 4.167924 | 4.222111 | 4.246383 | 4.258511 | 4.443064 | 5.0 | 1.524339 | 0.024527 | 1.486813 | 1.516939 | 1.529372 | 1.536470 | 1.552102 |
|  |  |  | log1p | 5.0 | 0.971797 | 0.002296 | 0.968319 | 0.970655 | 0.972747 | 0.973467 | 0.973796 | 5.0 | 0.980673 | 0.001715 | 0.977762 | 0.980530 | 0.981400 | 0.981694 | 0.981979 | 5.0 | 0.984479 | 0.001376 | 0.982145 | 0.984362 | 0.985062 | 0.985296 | 0.985529 | 5.0 | 5.240249 | 0.061523 | 5.190932 | 5.214184 | 5.214382 | 5.235044 | 5.346703 | 5.0 | 1.687732 | 0.010995 | 1.674595 | 1.679866 | 1.687934 | 1.694087 | 1.702178 |
|  |  | 16 | ftt | 5.0 | 0.972762 | 0.001626 | 0.970542 | 0.971968 | 0.972624 | 0.974220 | 0.974458 | 5.0 | 0.981486 | 0.000787 | 0.980380 | 0.980963 | 0.981831 | 0.981979 | 0.982274 | 5.0 | 0.985132 | 0.000632 | 0.984246 | 0.984712 | 0.985413 | 0.985529 | 0.985763 | 5.0 | 5.747153 | 0.071613 | 5.671821 | 5.691220 | 5.731632 | 5.801884 | 5.839208 | 5.0 | 5.218440 | 0.098497 | 5.096778 | 5.130996 | 5.253675 | 5.304616 | 5.306137 |
|  |  |  | log1p | 5.0 | 0.972384 | 0.001141 | 0.970722 | 0.971701 | 0.973004 | 0.973024 | 0.973471 | 5.0 | 0.980587 | 0.000695 | 0.979654 | 0.980241 | 0.980527 | 0.981108 | 0.981404 | 5.0 | 0.984409 | 0.000557 | 0.983662 | 0.984129 | 0.984362 | 0.984829 | 0.985062 | 5.0 | 6.871915 | 0.129302 | 6.763939 | 6.772803 | 6.841066 | 6.901060 | 7.080705 | 5.0 | 5.550288 | 0.027956 | 5.523888 | 5.537288 | 5.542704 | 5.550338 | 5.597222 |
| Plass2018 | 128 | 1 | ftt | 5.0 | 0.488277 | 0.005154 | 0.479141 | 0.489866 | 0.489889 | 0.491106 | 0.491382 | 5.0 | 0.641042 | 0.002116 | 0.638822 | 0.639921 | 0.640018 | 0.642467 | 0.643982 | 5.0 | 0.684610 | 0.001893 | 0.682676 | 0.683694 | 0.683741 | 0.685453 | 0.687488 | 5.0 | 11.186684 | 0.227652 | 11.004069 | 11.023573 | 11.055866 | 11.333606 | 11.516304 | 5.0 | 1.041303 | 0.011294 | 1.027123 | 1.031795 | 1.044609 | 1.051232 | 1.051756 |
|  |  |  | log1p | 5.0 | 0.493321 | 0.003513 | 0.487260 | 0.493264 | 0.495132 | 0.495288 | 0.495662 | 5.0 | 0.644047 | 0.003057 | 0.640376 | 0.641103 | 0.645670 | 0.646271 | 0.646815 | 5.0 | 0.687202 | 0.002625 | 0.684111 | 0.684666 | 0.688275 | 0.689293 | 0.689663 | 5.0 | 14.012541 | 0.971486 | 13.452991 | 13.534116 | 13.546827 | 13.793375 | 15.735397 | 5.0 | 1.477204 | 0.265845 | 1.343184 | 1.352351 | 1.366603 | 1.371547 | 1.952334 |
|  |  | 4 | ftt | 5.0 | 0.511670 | 0.002733 | 0.508854 | 0.508966 | 0.511783 | 0.514299 | 0.514448 | 5.0 | 0.663049 | 0.002213 | 0.661236 | 0.661295 | 0.661916 | 0.664734 | 0.666065 | 5.0 | 0.704007 | 0.002040 | 0.702202 | 0.702526 | 0.702989 | 0.705488 | 0.706830 | 5.0 | 11.167397 | 0.057440 | 11.115927 | 11.117341 | 11.173386 | 11.173672 | 11.256658 | 5.0 | 2.663776 | 0.014969 | 2.644864 | 2.656031 | 2.660126 | 2.677177 | 2.680681 |
|  |  |  | log1p | 5.0 | 0.513204 | 0.002403 | 0.509443 | 0.512632 | 0.513548 | 0.514678 | 0.515720 | 5.0 | 0.666099 | 0.001723 | 0.664139 | 0.664784 | 0.665847 | 0.667776 | 0.667949 | 5.0 | 0.706552 | 0.001582 | 0.704470 | 0.705627 | 0.706459 | 0.708033 | 0.708171 | 5.0 | 13.379228 | 0.085155 | 13.283675 | 13.313503 | 13.392609 | 13.406064 | 13.500291 | 5.0 | 3.126654 | 0.008152 | 3.114442 | 3.124210 | 3.126437 | 3.133970 | 3.134211 |
|  |  | 16 | ftt | 5.0 | 0.514289 | 0.003791 | 0.508337 | 0.513147 | 0.515109 | 0.516989 | 0.517863 | 5.0 | 0.668645 | 0.003464 | 0.663096 | 0.668266 | 0.668742 | 0.671376 | 0.671743 | 5.0 | 0.709125 | 0.003068 | 0.704192 | 0.708773 | 0.709282 | 0.711549 | 0.711827 | 5.0 | 12.427545 | 0.172742 | 12.279990 | 12.320841 | 12.397969 | 12.419389 | 12.719537 | 5.0 | 8.986059 | 0.073835 | 8.861482 | 8.982837 | 9.005108 | 9.039877 | 9.040989 |
|  |  |  | log1p | 5.0 | 0.524498 | 0.003716 | 0.518386 | 0.523854 | 0.525754 | 0.526637 | 0.527861 | 5.0 | 0.674999 | 0.001729 | 0.673252 | 0.673402 | 0.675127 | 0.675837 | 0.677375 | 5.0 | 0.714427 | 0.001525 | 0.712845 | 0.713076 | 0.714557 | 0.715112 | 0.716546 | 5.0 | 14.401086 | 0.077121 | 14.264553 | 14.427118 | 14.427139 | 14.432649 | 14.453970 | 5.0 | 10.028343 | 0.043434 | 9.966609 | 10.001759 | 10.042457 | 10.059282 | 10.071608 |
|  | 256 | 1 | ftt | 5.0 | 0.500564 | 0.003089 | 0.495286 | 0.500639 | 0.501608 | 0.502119 | 0.503170 | 5.0 | 0.654842 | 0.002247 | 0.652302 | 0.653634 | 0.654307 | 0.655798 | 0.658172 | 5.0 | 0.696974 | 0.002022 | 0.694614 | 0.695817 | 0.696743 | 0.697761 | 0.699935 | 5.0 | 11.245044 | 0.069221 | 11.150262 | 11.205262 | 11.264111 | 11.274807 | 11.330779 | 5.0 | 2.295233 | 0.031600 | 2.271256 | 2.271808 | 2.282618 | 2.303903 | 2.346578 |
|  |  |  | log1p | 5.0 | 0.508408 | 0.004170 | 0.501160 | 0.508535 | 0.510525 | 0.510889 | 0.510929 | 5.0 | 0.659474 | 0.002388 | 0.655972 | 0.658214 | 0.660471 | 0.660700 | 0.662014 | 5.0 | 0.700805 | 0.002142 | 0.697668 | 0.699611 | 0.701832 | 0.701925 | 0.702989 | 5.0 | 13.405713 | 0.301640 | 13.232788 | 13.244649 | 13.286452 | 13.323177 | 13.941499 | 5.0 | 2.746654 | 0.131074 | 2.672666 | 2.676561 | 2.676590 | 2.730243 | 2.977211 |
|  |  | 4 | ftt | 5.0 | 0.512536 | 0.003025 | 0.509524 | 0.510319 | 0.511473 | 0.514764 | 0.516600 | 5.0 | 0.666116 | 0.001333 | 0.664619 | 0.665458 | 0.665880 | 0.666442 | 0.668184 | 5.0 | 0.706811 | 0.001326 | 0.705210 | 0.706367 | 0.706644 | 0.706968 | 0.708865 | 5.0 | 12.237894 | 0.141680 | 12.104066 | 12.142734 | 12.165352 | 12.352571 | 12.424747 | 5.0 | 7.725106 | 0.122473 | 7.572591 | 7.689962 | 7.702010 | 7.750606 | 7.910360 |
|  |  |  | log1p | 5.0 | 0.523387 | 0.003864 | 0.518706 | 0.521807 | 0.521941 | 0.525912 | 0.528568 | 5.0 | 0.672946 | 0.001381 | 0.671644 | 0.672136 | 0.672251 | 0.673695 | 0.675004 | 5.0 | 0.712539 | 0.001347 | 0.711225 | 0.711688 | 0.711966 | 0.713307 | 0.714510 | 5.0 | 14.266884 | 0.042874 | 14.203213 | 14.243253 | 14.289430 | 14.291044 | 14.307481 | 5.0 | 8.709664 | 0.093072 | 8.598445 | 8.631647 | 8.721340 | 8.786349 | 8.810540 |
|  |  | 16 | ftt | 5.0 | 0.518644 | 0.003892 | 0.515792 | 0.516394 | 0.517784 | 0.517827 | 0.525425 | 5.0 | 0.670337 | 0.001812 | 0.668023 | 0.669866 | 0.670311 | 0.670400 | 0.673084 | 5.0 | 0.710651 | 0.001499 | 0.708634 | 0.710346 | 0.710670 | 0.710763 | 0.712845 | 5.0 | 15.865600 | 0.171886 | 15.685573 | 15.777681 | 15.790226 | 15.955337 | 16.119186 | 5.0 | 29.800970 | 0.330864 | 29.398653 | 29.602232 | 29.773648 | 29.976680 | 30.253639 |
|  |  |  | log1p | 5.0 | 0.526225 | 0.003547 | 0.520755 | 0.525165 | 0.526587 | 0.529057 | 0.529562 | 5.0 | 0.676698 | 0.001314 | 0.674602 | 0.676747 | 0.676750 | 0.677189 | 0.678202 | 5.0 | 0.715834 | 0.001118 | 0.714048 | 0.715945 | 0.715991 | 0.716037 | 0.717148 | 5.0 | 17.932459 | 0.089311 | 17.792226 | 17.910026 | 17.962139 | 17.967223 | 18.030683 | 5.0 | 33.265980 | 0.307553 | 32.887434 | 32.995569 | 33.376455 | 33.486485 | 33.583957 |
| Shekhar2016 | 128 | 1 | ftt | 5.0 | 0.971874 | 0.001026 | 0.970961 | 0.971004 | 0.971453 | 0.972900 | 0.973053 | 5.0 | 0.959141 | 0.000762 | 0.958113 | 0.958702 | 0.959155 | 0.959785 | 0.959951 | 5.0 | 0.966973 | 0.000616 | 0.966144 | 0.966617 | 0.966981 | 0.967490 | 0.967635 | 5.0 | 13.618053 | 0.160447 | 13.459037 | 13.535362 | 13.571757 | 13.645362 | 13.878745 | 5.0 | 1.442711 | 0.010182 | 1.427798 | 1.436681 | 1.447474 | 1.449717 | 1.451886 |
|  |  |  | log1p | 5.0 | 0.969727 | 0.001313 | 0.967392 | 0.970134 | 0.970263 | 0.970318 | 0.970526 | 5.0 | 0.954780 | 0.000849 | 0.953575 | 0.954375 | 0.954842 | 0.955366 | 0.955739 | 5.0 | 0.963446 | 0.000685 | 0.962471 | 0.963126 | 0.963490 | 0.963926 | 0.964217 | 5.0 | 18.246435 | 1.626557 | 17.180856 | 17.310832 | 17.458374 | 18.216107 | 21.066006 | 5.0 | 2.078371 | 0.341820 | 1.860708 | 1.892171 | 1.916162 | 2.046142 | 2.676673 |
|  |  | 4 | ftt | 5.0 | 0.976714 | 0.000628 | 0.976099 | 0.976210 | 0.976580 | 0.977072 | 0.977607 | 5.0 | 0.968542 | 0.001092 | 0.966953 | 0.968251 | 0.968618 | 0.968926 | 0.969960 | 5.0 | 0.974559 | 0.000885 | 0.973272 | 0.974326 | 0.974617 | 0.974872 | 0.975708 | 5.0 | 14.025118 | 0.277764 | 13.768480 | 13.813523 | 13.946738 | 14.152756 | 14.444095 | 5.0 | 3.813869 | 0.028507 | 3.785304 | 3.791465 | 3.804204 | 3.843478 | 3.844893 |
|  |  |  | log1p | 5.0 | 0.975487 | 0.000867 | 0.974376 | 0.975136 | 0.975258 | 0.976043 | 0.976621 | 5.0 | 0.966141 | 0.001024 | 0.964881 | 0.965198 | 0.966730 | 0.966768 | 0.967129 | 5.0 | 0.972617 | 0.000828 | 0.971599 | 0.971854 | 0.973090 | 0.973126 | 0.973417 | 5.0 | 17.605542 | 0.113194 | 17.484403 | 17.539235 | 17.587253 | 17.635326 | 17.781492 | 5.0 | 4.207394 | 0.082947 | 4.116006 | 4.125312 | 4.225589 | 4.280239 | 4.289821 |
|  |  | 16 | ftt | 5.0 | 0.977610 | 0.000800 | 0.976413 | 0.977244 | 0.977866 | 0.978071 | 0.978457 | 5.0 | 0.970722 | 0.000549 | 0.970147 | 0.970150 | 0.970905 | 0.971048 | 0.971361 | 5.0 | 0.976319 | 0.000445 | 0.975854 | 0.975854 | 0.976472 | 0.976581 | 0.976836 | 5.0 | 14.974952 | 0.255579 | 14.620645 | 14.794288 | 15.081576 | 15.188787 | 15.189463 | 5.0 | 12.916230 | 0.050769 | 12.857961 | 12.886367 | 12.898621 | 12.961282 | 12.976920 |
|  |  |  | log1p | 5.0 | 0.976576 | 0.000686 | 0.975578 | 0.976297 | 0.976659 | 0.976958 | 0.977386 | 5.0 | 0.968555 | 0.000721 | 0.967578 | 0.968299 | 0.968398 | 0.969069 | 0.969432 | 5.0 | 0.974566 | 0.000580 | 0.973781 | 0.974363 | 0.974435 | 0.974981 | 0.975272 | 5.0 | 18.848179 | 0.080930 | 18.742135 | 18.780478 | 18.891775 | 18.913043 | 18.913464 | 5.0 | 13.041363 | 0.058419 | 12.979966 | 13.003295 | 13.016472 | 13.097194 | 13.109889 |
|  | 256 | 1 | ftt | 5.0 | 0.975532 | 0.000369 | 0.975152 | 0.975182 | 0.975628 | 0.975662 | 0.976034 | 5.0 | 0.965821 | 0.000451 | 0.965328 | 0.965549 | 0.965646 | 0.966178 | 0.966403 | 5.0 | 0.972363 | 0.000366 | 0.971963 | 0.972144 | 0.972217 | 0.972654 | 0.972835 | 5.0 | 13.485717 | 0.158160 | 13.299532 | 13.333406 | 13.560745 | 13.586973 | 13.647929 | 5.0 | 4.048799 | 0.081130 | 3.935916 | 3.995296 | 4.072911 | 4.116695 | 4.123177 |
|  |  |  | log1p | 5.0 | 0.973334 | 0.000342 | 0.972855 | 0.973230 | 0.973326 | 0.973466 | 0.973793 | 5.0 | 0.962967 | 0.000624 | 0.962262 | 0.962537 | 0.963025 | 0.963125 | 0.963886 | 5.0 | 0.970057 | 0.000503 | 0.969490 | 0.969708 | 0.970108 | 0.970181 | 0.970799 | 5.0 | 16.980569 | 0.155286 | 16.872210 | 16.872483 | 16.946043 | 16.964025 | 17.248086 | 5.0 | 4.310925 | 0.099190 | 4.248474 | 4.255421 | 4.267161 | 4.298562 | 4.485009 |
|  |  | 4 | ftt | 5.0 | 0.977855 | 0.000349 | 0.977470 | 0.977674 | 0.977738 | 0.978023 | 0.978369 | 5.0 | 0.970899 | 0.000384 | 0.970548 | 0.970593 | 0.970732 | 0.971269 | 0.971355 | 5.0 | 0.976465 | 0.000311 | 0.976181 | 0.976217 | 0.976326 | 0.976763 | 0.976836 | 5.0 | 15.560010 | 0.566545 | 15.076371 | 15.122633 | 15.249947 | 16.126695 | 16.224402 | 5.0 | 15.540619 | 0.767196 | 15.079377 | 15.122242 | 15.130403 | 15.490388 | 16.880684 |
|  |  |  | log1p | 5.0 | 0.976774 | 0.000676 | 0.976129 | 0.976428 | 0.976486 | 0.976969 | 0.977856 | 5.0 | 0.968248 | 0.000597 | 0.967228 | 0.968255 | 0.968478 | 0.968532 | 0.968749 | 5.0 | 0.974319 | 0.000485 | 0.973490 | 0.974326 | 0.974508 | 0.974545 | 0.974726 | 5.0 | 18.566400 | 0.414539 | 18.274084 | 18.389627 | 18.412482 | 18.457838 | 19.297971 | 5.0 | 14.730118 | 0.100629 | 14.574827 | 14.703320 | 14.740536 | 14.795621 | 14.836288 |
|  |  | 16 | ftt | 5.0 | 0.978420 | 0.000435 | 0.977720 | 0.978410 | 0.978417 | 0.978686 | 0.978864 | 5.0 | 0.971369 | 0.000705 | 0.970553 | 0.970772 | 0.971399 | 0.971992 | 0.972127 | 5.0 | 0.976843 | 0.000569 | 0.976181 | 0.976363 | 0.976872 | 0.977345 | 0.977454 | 5.0 | 19.132078 | 0.078730 | 19.026862 | 19.108600 | 19.132436 | 19.146561 | 19.245931 | 5.0 | 57.701285 | 0.463692 | 57.015978 | 57.684865 | 57.720659 | 57.762328 | 58.322593 |
|  |  |  | log1p | 5.0 | 0.976824 | 0.001122 | 0.975220 | 0.976305 | 0.977016 | 0.977407 | 0.978173 | 5.0 | 0.968961 | 0.001444 | 0.967772 | 0.967811 | 0.968342 | 0.969786 | 0.971094 | 5.0 | 0.974894 | 0.001169 | 0.973926 | 0.973963 | 0.974399 | 0.975563 | 0.976617 | 5.0 | 22.902277 | 0.496196 | 22.503588 | 22.695394 | 22.726402 | 22.820334 | 23.765667 | 5.0 | 54.607733 | 1.184124 | 53.864850 | 54.043531 | 54.125221 | 54.297491 | 56.707573 |
| TabulaMuris-chromium | 128 | 1 | ftt | 5.0 | 0.938496 | 0.000655 | 0.937860 | 0.938181 | 0.938355 | 0.938495 | 0.939588 | 5.0 | 0.944558 | 0.001020 | 0.943648 | 0.943672 | 0.944178 | 0.945500 | 0.945790 | 5.0 | 0.948536 | 0.000944 | 0.947696 | 0.947714 | 0.948187 | 0.949406 | 0.949679 | 5.0 | 23.285202 | 0.697160 | 22.662000 | 22.670226 | 23.108707 | 23.722580 | 24.262496 | 5.0 | 2.339003 | 0.178397 | 2.167775 | 2.235568 | 2.316676 | 2.341411 | 2.633588 |
|  |  |  | log1p | 5.0 | 0.945836 | 0.000968 | 0.944295 | 0.945705 | 0.946043 | 0.946223 | 0.946913 | 5.0 | 0.951859 | 0.000748 | 0.950965 | 0.951233 | 0.952000 | 0.952355 | 0.952743 | 5.0 | 0.955315 | 0.000695 | 0.954482 | 0.954736 | 0.955446 | 0.955773 | 0.956137 | 5.0 | 30.627154 | 1.373100 | 29.541582 | 30.028411 | 30.161255 | 30.383953 | 33.020570 | 5.0 | 3.321298 | 0.288716 | 2.963337 | 3.159204 | 3.299935 | 3.465528 | 3.718485 |
|  |  | 4 | ftt | 5.0 | 0.953169 | 0.000502 | 0.952368 | 0.953152 | 0.953231 | 0.953353 | 0.953743 | 5.0 | 0.958590 | 0.000373 | 0.958134 | 0.958372 | 0.958508 | 0.958901 | 0.959036 | 5.0 | 0.961562 | 0.000346 | 0.961140 | 0.961359 | 0.961486 | 0.961850 | 0.961977 | 5.0 | 23.531871 | 0.820726 | 22.282308 | 23.381934 | 23.684855 | 23.761318 | 24.548939 | 5.0 | 5.537701 | 0.396202 | 5.077076 | 5.249611 | 5.490380 | 5.853092 | 6.018348 |
|  |  |  | log1p | 5.0 | 0.959821 | 0.000586 | 0.959458 | 0.959520 | 0.959619 | 0.959644 | 0.960861 | 5.0 | 0.964943 | 0.000577 | 0.964488 | 0.964606 | 0.964820 | 0.964860 | 0.965938 | 5.0 | 0.967457 | 0.000536 | 0.967035 | 0.967144 | 0.967344 | 0.967380 | 0.968381 | 5.0 | 30.469801 | 0.507090 | 29.791848 | 30.289590 | 30.343434 | 30.828193 | 31.095942 | 5.0 | 6.491028 | 0.325697 | 6.077408 | 6.226313 | 6.577190 | 6.772971 | 6.801256 |
|  |  | 16 | ftt | 5.0 | 0.957378 | 0.000788 | 0.956263 | 0.956980 | 0.957607 | 0.957697 | 0.958340 | 5.0 | 0.962748 | 0.000431 | 0.962270 | 0.962408 | 0.962777 | 0.962938 | 0.963348 | 5.0 | 0.965423 | 0.000400 | 0.964979 | 0.965106 | 0.965452 | 0.965598 | 0.965980 | 5.0 | 24.876913 | 0.527322 | 24.271999 | 24.690138 | 24.761884 | 24.951346 | 25.709197 | 5.0 | 18.753942 | 0.347634 | 18.416846 | 18.446048 | 18.733341 | 18.923895 | 19.249580 |
|  |  |  | log1p | 5.0 | 0.964153 | 0.001059 | 0.962358 | 0.964165 | 0.964394 | 0.964830 | 0.965019 | 5.0 | 0.969149 | 0.000742 | 0.968232 | 0.968976 | 0.968996 | 0.969250 | 0.970289 | 5.0 | 0.971361 | 0.000689 | 0.970510 | 0.971201 | 0.971219 | 0.971456 | 0.972420 | 5.0 | 32.785235 | 1.032009 | 31.418146 | 32.058392 | 33.001005 | 33.562084 | 33.886546 | 5.0 | 19.817969 | 1.139513 | 18.582210 | 19.026082 | 19.604493 | 20.447167 | 21.429895 |
|  | 256 | 1 | ftt | 5.0 | 0.948172 | 0.001039 | 0.947322 | 0.947390 | 0.948088 | 0.948161 | 0.949897 | 5.0 | 0.953617 | 0.000965 | 0.952315 | 0.953255 | 0.953570 | 0.954003 | 0.954940 | 5.0 | 0.956945 | 0.000896 | 0.955737 | 0.956610 | 0.956901 | 0.957302 | 0.958175 | 5.0 | 22.574396 | 0.095080 | 22.414768 | 22.578393 | 22.586486 | 22.635772 | 22.656560 | 5.0 | 4.600895 | 0.079647 | 4.488142 | 4.561152 | 4.610813 | 4.651951 | 4.692418 |
|  |  |  | log1p | 5.0 | 0.954683 | 0.000694 | 0.953838 | 0.954349 | 0.954476 | 0.955161 | 0.955592 | 5.0 | 0.960015 | 0.000323 | 0.959805 | 0.959864 | 0.959899 | 0.959921 | 0.960588 | 5.0 | 0.962883 | 0.000300 | 0.962687 | 0.962741 | 0.962778 | 0.962796 | 0.963414 | 5.0 | 29.836975 | 0.287995 | 29.478672 | 29.589898 | 29.948021 | 30.021613 | 30.146673 | 5.0 | 5.704838 | 0.095189 | 5.564199 | 5.659533 | 5.724919 | 5.776501 | 5.799036 |
|  |  | 4 | ftt | 5.0 | 0.955995 | 0.000708 | 0.954918 | 0.955627 | 0.956389 | 0.956514 | 0.956529 | 5.0 | 0.961491 | 0.000427 | 0.960782 | 0.961430 | 0.961624 | 0.961781 | 0.961840 | 5.0 | 0.964255 | 0.000397 | 0.963596 | 0.964197 | 0.964379 | 0.964524 | 0.964579 | 5.0 | 24.672266 | 0.075017 | 24.589160 | 24.616490 | 24.655986 | 24.740115 | 24.759580 | 5.0 | 15.270638 | 0.210636 | 14.898989 | 15.340062 | 15.342508 | 15.348193 | 15.423438 |
|  |  |  | log1p | 5.0 | 0.963191 | 0.000441 | 0.962724 | 0.963064 | 0.963081 | 0.963169 | 0.963919 | 5.0 | 0.967984 | 0.000135 | 0.967780 | 0.967956 | 0.967977 | 0.968074 | 0.968132 | 5.0 | 0.970280 | 0.000125 | 0.970091 | 0.970255 | 0.970273 | 0.970364 | 0.970419 | 5.0 | 31.821701 | 0.173331 | 31.654699 | 31.718679 | 31.743647 | 31.909134 | 32.082344 | 5.0 | 16.957797 | 0.094327 | 16.809100 | 16.919385 | 17.007452 | 17.019737 | 17.033310 |
|  |  | 16 | ftt | 5.0 | 0.958032 | 0.000522 | 0.957568 | 0.957597 | 0.957843 | 0.958408 | 0.958746 | 5.0 | 0.963537 | 0.000342 | 0.963112 | 0.963230 | 0.963702 | 0.963761 | 0.963879 | 5.0 | 0.966154 | 0.000317 | 0.965761 | 0.965870 | 0.966307 | 0.966362 | 0.966471 | 5.0 | 32.487938 | 0.370952 | 32.066628 | 32.230699 | 32.494404 | 32.622909 | 33.025049 | 5.0 | 57.240301 | 1.374670 | 56.020702 | 56.114898 | 56.616706 | 58.654733 | 58.794467 |
|  |  |  | log1p | 5.0 | 0.965015 | 0.000469 | 0.964463 | 0.964769 | 0.964836 | 0.965467 | 0.965542 | 5.0 | 0.970011 | 0.000577 | 0.969212 | 0.969761 | 0.970151 | 0.970153 | 0.970779 | 5.0 | 0.972161 | 0.000536 | 0.971419 | 0.971929 | 0.972292 | 0.972292 | 0.972875 | 5.0 | 39.850823 | 0.207958 | 39.586302 | 39.715568 | 39.839600 | 40.048084 | 40.064560 | 5.0 | 60.332995 | 0.728268 | 59.463947 | 59.806399 | 60.515863 | 60.539320 | 61.339445 |

In [25]:

```
knn_cv_scmap.groupby(["dataset", "n-features", "n-centroids-factor"]).describe()
```

Out[25]:

|  |  |  | adjusted\_rand | | | | | | | | cohen\_kappa | | | | | | | | consistency | | | | | | | | index-time | | | | | | | | query-time | | | | | | | |
| --- | --- | --- | --- | --- | --- | --- | --- | --- | --- | --- | --- | --- | --- | --- | --- | --- | --- | --- | --- | --- | --- | --- | --- | --- | --- | --- | --- | --- | --- | --- | --- | --- | --- | --- | --- | --- | --- | --- | --- | --- | --- | --- |
|  |  |  | count | mean | std | min | 25% | 50% | 75% | max | count | mean | std | min | 25% | 50% | 75% | max | count | mean | std | min | 25% | 50% | 75% | max | count | mean | std | min | 25% | 50% | 75% | max | count | mean | std | min | 25% | 50% | 75% | max |
| dataset | n-features | n-centroids-factor |  |  |  |  |  |  |  |  |  |  |  |  |  |  |  |  |  |  |  |  |  |  |  |  |  |  |  |  |  |  |  |  |  |  |  |  |  |  |  |  |
| Baron2016-human | 500 | 1 | 5.0 | 0.970908 | 0.000561 | 0.970293 | 0.970363 | 0.971055 | 0.971264 | 0.971567 | 5.0 | 0.973140 | 0.000352 | 0.972639 | 0.972939 | 0.973226 | 0.973375 | 0.973518 | 5.0 | 0.978457 | 0.000281 | 0.978060 | 0.978294 | 0.978527 | 0.978644 | 0.978761 | 5.0 | 57.155702 | 0.672800 | 56.45500 | 56.50400 | 57.34592 | 57.43396 | 58.03963 | 5.0 | 22.670936 | 0.533049 | 21.99949 | 22.48407 | 22.51334 | 22.94444 | 23.41334 |
|  |  | 4 | 5.0 | 0.970981 | 0.001681 | 0.969446 | 0.970304 | 0.970612 | 0.970686 | 0.973857 | 5.0 | 0.971881 | 0.001710 | 0.970454 | 0.970742 | 0.970889 | 0.972933 | 0.974387 | 5.0 | 0.977454 | 0.001369 | 0.976310 | 0.976543 | 0.976660 | 0.978294 | 0.979461 | 5.0 | 95.141148 | 1.116597 | 93.74152 | 94.77946 | 94.83448 | 95.59849 | 96.75179 | 5.0 | 24.382970 | 0.331719 | 23.93020 | 24.18853 | 24.43464 | 24.58763 | 24.77385 |
|  | 1000 | 1 | 5.0 | 0.967196 | 0.000924 | 0.966209 | 0.966478 | 0.967037 | 0.967841 | 0.968414 | 5.0 | 0.971715 | 0.000636 | 0.970864 | 0.971303 | 0.971887 | 0.972038 | 0.972481 | 5.0 | 0.977337 | 0.000505 | 0.976660 | 0.977010 | 0.977477 | 0.977594 | 0.977944 | 5.0 | 81.341326 | 0.301259 | 81.01097 | 81.08971 | 81.38747 | 81.45941 | 81.75907 | 5.0 | 38.807398 | 0.268174 | 38.39115 | 38.72099 | 38.85217 | 39.02899 | 39.04369 |
|  |  | 4 | 5.0 | 0.963776 | 0.001471 | 0.961932 | 0.962661 | 0.964298 | 0.964373 | 0.965617 | 5.0 | 0.967270 | 0.000621 | 0.966471 | 0.967063 | 0.967069 | 0.967658 | 0.968089 | 5.0 | 0.973789 | 0.000492 | 0.973159 | 0.973626 | 0.973626 | 0.974093 | 0.974443 | 5.0 | 159.205600 | 0.956523 | 157.86260 | 158.80880 | 159.10750 | 160.11820 | 160.13090 | 5.0 | 43.516568 | 0.885182 | 42.12374 | 43.15609 | 43.96796 | 44.08680 | 44.24825 |
|  | 2000 | 1 | 5.0 | 0.968000 | 0.001855 | 0.966451 | 0.966673 | 0.967111 | 0.968958 | 0.970809 | 5.0 | 0.973988 | 0.001114 | 0.972181 | 0.973927 | 0.974072 | 0.974665 | 0.975095 | 5.0 | 0.979157 | 0.000891 | 0.977710 | 0.979111 | 0.979227 | 0.979694 | 0.980044 | 5.0 | 119.627180 | 0.966773 | 118.49830 | 119.33040 | 119.51540 | 119.62720 | 121.16460 | 5.0 | 51.661450 | 0.605673 | 50.85426 | 51.53165 | 51.53473 | 51.86379 | 52.52282 |
|  |  | 4 | 5.0 | 0.961701 | 0.002914 | 0.957308 | 0.960917 | 0.962042 | 0.963058 | 0.965180 | 5.0 | 0.968133 | 0.001603 | 0.965876 | 0.967783 | 0.967927 | 0.968805 | 0.970272 | 5.0 | 0.974489 | 0.001277 | 0.972692 | 0.974209 | 0.974326 | 0.975026 | 0.976193 | 5.0 | 248.541400 | 0.914132 | 247.55280 | 247.76700 | 248.49920 | 249.18480 | 249.70320 | 5.0 | 58.198086 | 0.230564 | 57.97407 | 58.02217 | 58.11415 | 58.37919 | 58.50085 |
|  | 4000 | 1 | 5.0 | 0.967021 | 0.001092 | 0.965322 | 0.966676 | 0.967426 | 0.967482 | 0.968197 | 5.0 | 0.972729 | 0.000649 | 0.972025 | 0.972466 | 0.972614 | 0.972761 | 0.973780 | 5.0 | 0.978154 | 0.000518 | 0.977594 | 0.977944 | 0.978060 | 0.978177 | 0.978994 | 5.0 | 195.233280 | 2.145728 | 192.91330 | 193.14830 | 195.48870 | 197.22880 | 197.38730 | 5.0 | 80.696716 | 0.847593 | 79.53714 | 80.17671 | 80.86205 | 81.24790 | 81.65978 |
|  |  | 4 | 5.0 | 0.960666 | 0.002621 | 0.957472 | 0.958479 | 0.961466 | 0.962126 | 0.963785 | 5.0 | 0.966029 | 0.001928 | 0.963531 | 0.964857 | 0.966188 | 0.967056 | 0.968515 | 5.0 | 0.972809 | 0.001535 | 0.970825 | 0.971875 | 0.972926 | 0.973626 | 0.974793 | 5.0 | 426.395840 | 2.039697 | 423.38040 | 425.46810 | 426.94540 | 427.53250 | 428.65280 | 5.0 | 94.758720 | 0.620577 | 94.12244 | 94.39889 | 94.71934 | 94.79249 | 95.76044 |
| Plass2018 | 500 | 1 | 5.0 | 0.464535 | 0.005307 | 0.456000 | 0.463199 | 0.465906 | 0.468675 | 0.468893 | 5.0 | 0.586654 | 0.003140 | 0.582532 | 0.584073 | 0.588155 | 0.589212 | 0.589296 | 5.0 | 0.647501 | 0.002819 | 0.643716 | 0.645243 | 0.649176 | 0.649593 | 0.649778 | 5.0 | 186.276880 | 8.732591 | 176.26420 | 178.07850 | 188.40800 | 192.94910 | 195.68460 | 5.0 | 136.193440 | 3.362967 | 132.18960 | 135.14070 | 135.60760 | 136.58180 | 141.44750 |
|  |  | 4 | 5.0 | 0.469142 | 0.002737 | 0.466781 | 0.467489 | 0.468248 | 0.469495 | 0.473696 | 5.0 | 0.591928 | 0.002577 | 0.589008 | 0.589658 | 0.592091 | 0.594091 | 0.594790 | 5.0 | 0.652369 | 0.002278 | 0.649639 | 0.650287 | 0.653063 | 0.654405 | 0.654451 | 5.0 | 253.478940 | 5.354989 | 249.51870 | 249.89530 | 250.45030 | 255.51590 | 262.01450 | 5.0 | 137.277120 | 1.448147 | 136.08620 | 136.22210 | 136.71620 | 137.78520 | 139.57590 |
|  | 1000 | 1 | 5.0 | 0.469017 | 0.005154 | 0.464364 | 0.465524 | 0.466352 | 0.472566 | 0.476278 | 5.0 | 0.581250 | 0.002206 | 0.578248 | 0.579968 | 0.581452 | 0.582822 | 0.583760 | 5.0 | 0.642004 | 0.001903 | 0.639922 | 0.640154 | 0.642375 | 0.643439 | 0.644133 | 5.0 | 182.098240 | 1.968912 | 180.19660 | 180.83070 | 181.90740 | 182.26130 | 185.29520 | 5.0 | 221.522520 | 2.843169 | 218.00120 | 219.42850 | 222.11950 | 222.89120 | 225.17220 |
|  |  | 4 | 5.0 | 0.478252 | 0.002835 | 0.475809 | 0.476580 | 0.476595 | 0.479724 | 0.482551 | 5.0 | 0.586794 | 0.002388 | 0.583673 | 0.586360 | 0.586730 | 0.586827 | 0.590381 | 5.0 | 0.646641 | 0.002272 | 0.643393 | 0.646354 | 0.646632 | 0.647048 | 0.649778 | 5.0 | 332.476520 | 4.735158 | 329.14180 | 329.23290 | 329.53820 | 334.57700 | 339.89270 | 5.0 | 243.358100 | 3.909889 | 238.44280 | 241.79680 | 242.50000 | 245.16590 | 248.88500 |
|  | 2000 | 1 | 5.0 | 0.468279 | 0.001570 | 0.466708 | 0.466721 | 0.468668 | 0.468927 | 0.470374 | 5.0 | 0.591757 | 0.001978 | 0.588712 | 0.591087 | 0.592051 | 0.593441 | 0.593493 | 5.0 | 0.651342 | 0.001722 | 0.648899 | 0.650379 | 0.651629 | 0.652785 | 0.653017 | 5.0 | 229.652940 | 1.697726 | 227.00070 | 229.48640 | 229.49160 | 231.12900 | 231.15700 | 5.0 | 314.252680 | 2.454708 | 312.53770 | 312.56300 | 313.74800 | 313.92590 | 318.48880 |
|  |  | 4 | 5.0 | 0.478969 | 0.003239 | 0.475391 | 0.477027 | 0.478392 | 0.480218 | 0.483817 | 5.0 | 0.597433 | 0.002594 | 0.594214 | 0.595829 | 0.597567 | 0.598558 | 0.600998 | 5.0 | 0.656385 | 0.002079 | 0.653942 | 0.654868 | 0.656441 | 0.657505 | 0.659171 | 5.0 | 535.830200 | 3.664045 | 531.94000 | 534.08210 | 534.76140 | 536.74690 | 541.62060 | 5.0 | 348.530380 | 3.870894 | 344.57180 | 345.93490 | 347.72560 | 349.99570 | 354.42390 |
|  | 4000 | 1 | 5.0 | 0.467713 | 0.002892 | 0.464631 | 0.465812 | 0.466854 | 0.469503 | 0.471764 | 5.0 | 0.590322 | 0.001685 | 0.587958 | 0.589917 | 0.590147 | 0.591008 | 0.592582 | 5.0 | 0.649935 | 0.001940 | 0.647326 | 0.649269 | 0.649732 | 0.650750 | 0.652600 | 5.0 | 367.464780 | 2.543230 | 364.72770 | 364.88710 | 368.12750 | 369.30290 | 370.27870 | 5.0 | 509.813020 | 4.014794 | 503.47230 | 509.30560 | 510.44620 | 511.45450 | 514.38650 |
|  |  | 4 | 5.0 | 0.474737 | 0.005549 | 0.466985 | 0.473072 | 0.474778 | 0.476568 | 0.482284 | 5.0 | 0.596088 | 0.003854 | 0.591881 | 0.594203 | 0.594214 | 0.598866 | 0.601277 | 5.0 | 0.655386 | 0.003672 | 0.651397 | 0.653387 | 0.653757 | 0.658107 | 0.660281 | 5.0 | 941.772060 | 6.538889 | 933.43130 | 937.98240 | 941.46370 | 945.79580 | 950.18710 | 5.0 | 552.939880 | 5.073921 | 546.81590 | 549.59920 | 552.06890 | 557.83930 | 558.37610 |
| Shekhar2016 | 500 | 1 | 5.0 | 0.926637 | 0.002478 | 0.924078 | 0.924311 | 0.927263 | 0.927514 | 0.930019 | 5.0 | 0.892993 | 0.002233 | 0.890389 | 0.891216 | 0.892905 | 0.895196 | 0.895261 | 5.0 | 0.913786 | 0.001788 | 0.911706 | 0.912360 | 0.913706 | 0.915561 | 0.915597 | 5.0 | 260.558480 | 6.946313 | 250.20490 | 257.12200 | 262.53370 | 266.07710 | 266.85470 | 5.0 | 199.466940 | 5.235952 | 190.76980 | 199.38670 | 200.15250 | 202.87830 | 204.14740 |
|  |  | 4 | 5.0 | 0.923893 | 0.001373 | 0.922393 | 0.923314 | 0.923419 | 0.924314 | 0.926027 | 5.0 | 0.882686 | 0.002260 | 0.880413 | 0.880906 | 0.882734 | 0.883257 | 0.886117 | 5.0 | 0.905335 | 0.001809 | 0.903524 | 0.903887 | 0.905378 | 0.905815 | 0.908069 | 5.0 | 508.008760 | 24.636501 | 493.32590 | 495.29360 | 499.24060 | 500.40440 | 551.77930 | 5.0 | 216.691260 | 2.975053 | 213.72340 | 214.30200 | 216.20450 | 218.27630 | 220.95010 |
|  | 1000 | 1 | 5.0 | 0.909582 | 0.003618 | 0.906697 | 0.908047 | 0.908329 | 0.908955 | 0.915883 | 5.0 | 0.856641 | 0.003630 | 0.851942 | 0.855766 | 0.856312 | 0.857105 | 0.862077 | 5.0 | 0.884323 | 0.002985 | 0.880505 | 0.883596 | 0.884032 | 0.884650 | 0.888832 | 5.0 | 369.210020 | 1.405052 | 367.31540 | 368.88370 | 368.93860 | 369.74170 | 371.17070 | 5.0 | 358.011100 | 2.221996 | 355.81090 | 356.74670 | 357.10480 | 358.98500 | 361.40810 |
|  |  | 4 | 5.0 | 0.898594 | 0.001638 | 0.896618 | 0.897713 | 0.898346 | 0.899388 | 0.900908 | 5.0 | 0.822569 | 0.002539 | 0.818646 | 0.821415 | 0.823702 | 0.824502 | 0.824580 | 5.0 | 0.856569 | 0.002048 | 0.853413 | 0.855631 | 0.857449 | 0.858177 | 0.858177 | 5.0 | 871.111380 | 3.167970 | 867.77910 | 868.67480 | 870.22080 | 874.18040 | 874.70180 | 5.0 | 383.892060 | 0.443237 | 383.30560 | 383.61720 | 383.97730 | 384.10990 | 384.45030 |
|  | 2000 | 1 | 5.0 | 0.908230 | 0.002625 | 0.906222 | 0.906279 | 0.906448 | 0.910907 | 0.911292 | 5.0 | 0.856459 | 0.002461 | 0.854143 | 0.855460 | 0.855896 | 0.856158 | 0.860636 | 5.0 | 0.884119 | 0.001993 | 0.882287 | 0.883341 | 0.883632 | 0.883814 | 0.887523 | 5.0 | 539.567120 | 8.193210 | 533.00800 | 534.80080 | 537.47530 | 538.91410 | 553.63740 | 5.0 | 517.021940 | 3.442053 | 512.77810 | 514.02680 | 518.34960 | 519.21100 | 520.74420 |
|  |  | 4 | 5.0 | 0.875560 | 0.006957 | 0.866467 | 0.870184 | 0.877661 | 0.880983 | 0.882504 | 5.0 | 0.781743 | 0.007697 | 0.771873 | 0.774971 | 0.786411 | 0.787418 | 0.788043 | 5.0 | 0.823077 | 0.006417 | 0.814866 | 0.817412 | 0.826939 | 0.827848 | 0.828321 | 5.0 | 1284.425200 | 4.672343 | 1278.79900 | 1281.73200 | 1283.19200 | 1288.40500 | 1289.99800 | 5.0 | 552.178320 | 3.941707 | 547.59550 | 549.23210 | 551.73720 | 555.65870 | 556.66810 |
|  | 4000 | 1 | 5.0 | 0.905724 | 0.003270 | 0.902046 | 0.902459 | 0.907158 | 0.907653 | 0.909302 | 5.0 | 0.848924 | 0.002351 | 0.845259 | 0.848166 | 0.849393 | 0.850727 | 0.851077 | 5.0 | 0.878032 | 0.001948 | 0.875014 | 0.877377 | 0.878396 | 0.879559 | 0.879814 | 5.0 | 897.083060 | 21.552047 | 883.53150 | 885.41200 | 885.51140 | 896.41070 | 934.54970 | 5.0 | 798.755160 | 12.535108 | 792.02530 | 792.14270 | 793.63120 | 794.89740 | 821.07920 |
|  |  | 4 | 5.0 | 0.854074 | 0.007194 | 0.846234 | 0.849008 | 0.855039 | 0.855164 | 0.864925 | 5.0 | 0.751021 | 0.006464 | 0.744290 | 0.745865 | 0.749938 | 0.755271 | 0.759739 | 5.0 | 0.797607 | 0.005358 | 0.791883 | 0.793192 | 0.797193 | 0.801047 | 0.804720 | 5.0 | 2254.789000 | 7.682993 | 2247.17900 | 2248.22300 | 2255.14000 | 2257.26300 | 2266.14000 | 5.0 | 868.164740 | 1.518618 | 867.24010 | 867.51660 | 867.54910 | 867.65020 | 870.86770 |
| TabulaMuris-chromium | 500 | 1 | 5.0 | 0.943124 | 0.001750 | 0.940424 | 0.942352 | 0.943861 | 0.944300 | 0.944683 | 5.0 | 0.949715 | 0.001299 | 0.948128 | 0.948600 | 0.950029 | 0.950772 | 0.951046 | 5.0 | 0.953299 | 0.001207 | 0.951826 | 0.952262 | 0.953590 | 0.954282 | 0.954536 | 5.0 | 662.124680 | 21.851509 | 637.20970 | 652.41150 | 661.88220 | 662.43090 | 696.68910 | 5.0 | 779.215860 | 6.422588 | 769.43630 | 777.38540 | 780.57090 | 781.93610 | 786.75060 |
|  |  | 4 | 5.0 | 0.942044 | 0.001004 | 0.940709 | 0.941589 | 0.941937 | 0.942650 | 0.943333 | 5.0 | 0.949026 | 0.000708 | 0.948034 | 0.948911 | 0.948953 | 0.949224 | 0.950008 | 5.0 | 0.952659 | 0.000659 | 0.951735 | 0.952553 | 0.952590 | 0.952844 | 0.953572 | 5.0 | 1432.738000 | 15.125488 | 1414.84100 | 1424.51700 | 1433.44600 | 1435.34600 | 1455.54000 | 5.0 | 811.140960 | 13.600745 | 796.83920 | 798.31250 | 812.48480 | 819.76000 | 828.30830 |
|  | 1000 | 1 | 5.0 | 0.931309 | 0.001560 | 0.929553 | 0.929776 | 0.931919 | 0.932229 | 0.933066 | 5.0 | 0.941841 | 0.001416 | 0.940303 | 0.940423 | 0.942130 | 0.943126 | 0.943223 | 5.0 | 0.945993 | 0.001315 | 0.944567 | 0.944676 | 0.946259 | 0.947186 | 0.947277 | 5.0 | 990.544820 | 8.339468 | 982.06360 | 983.26210 | 990.53830 | 994.63110 | 1002.22900 | 5.0 | 1404.769200 | 11.513616 | 1395.39400 | 1397.63700 | 1399.32600 | 1407.96800 | 1423.52100 |
|  |  | 4 | 5.0 | 0.926436 | 0.001341 | 0.924404 | 0.925866 | 0.926739 | 0.927559 | 0.927612 | 5.0 | 0.939234 | 0.000828 | 0.938008 | 0.938831 | 0.939416 | 0.939926 | 0.939986 | 5.0 | 0.943577 | 0.000770 | 0.942438 | 0.943202 | 0.943748 | 0.944221 | 0.944276 | 5.0 | 2549.600600 | 22.071396 | 2518.45000 | 2540.44000 | 2553.78900 | 2557.07100 | 2578.25300 | 5.0 | 1553.903600 | 23.205745 | 1520.64500 | 1547.51400 | 1553.24900 | 1564.19400 | 1583.91600 |
|  | 2000 | 1 | 5.0 | 0.938039 | 0.001290 | 0.935939 | 0.937788 | 0.938342 | 0.939044 | 0.939081 | 5.0 | 0.946501 | 0.000946 | 0.945036 | 0.946089 | 0.946937 | 0.947095 | 0.947348 | 5.0 | 0.950312 | 0.000877 | 0.948951 | 0.949934 | 0.950716 | 0.950861 | 0.951098 | 5.0 | 1458.824000 | 10.558727 | 1446.07400 | 1451.37900 | 1457.82100 | 1468.35700 | 1470.48900 | 5.0 | 1978.130400 | 15.452936 | 1962.99800 | 1965.68400 | 1974.55400 | 1987.66000 | 1999.75600 |
|  |  | 4 | 5.0 | 0.933155 | 0.001730 | 0.931910 | 0.931956 | 0.932577 | 0.933240 | 0.936094 | 5.0 | 0.943414 | 0.000987 | 0.942703 | 0.942744 | 0.943253 | 0.943255 | 0.945113 | 5.0 | 0.947445 | 0.000916 | 0.946786 | 0.946823 | 0.947296 | 0.947296 | 0.949024 | 5.0 | 3668.239000 | 14.086305 | 3646.34100 | 3662.10100 | 3674.77200 | 3677.84400 | 3680.13700 | 5.0 | 2084.666200 | 23.940830 | 2060.25700 | 2075.19700 | 2077.46900 | 2086.33900 | 2124.06900 |
|  | 4000 | 1 | 5.0 | 0.936648 | 0.001129 | 0.935631 | 0.935826 | 0.936149 | 0.937362 | 0.938270 | 5.0 | 0.945479 | 0.001121 | 0.944230 | 0.944819 | 0.945033 | 0.946481 | 0.946833 | 5.0 | 0.949366 | 0.001043 | 0.948205 | 0.948751 | 0.948951 | 0.950297 | 0.950625 | 5.0 | 2238.827800 | 12.062036 | 2219.46400 | 2236.09000 | 2241.40200 | 2247.53500 | 2249.64800 | 5.0 | 3119.246200 | 59.969039 | 3049.24500 | 3070.42400 | 3118.91700 | 3177.68500 | 3179.96000 |
|  |  | 4 | 5.0 | 0.930650 | 0.001571 | 0.928746 | 0.929403 | 0.931088 | 0.931368 | 0.932646 | 5.0 | 0.942313 | 0.001193 | 0.940818 | 0.941527 | 0.942351 | 0.943036 | 0.943835 | 5.0 | 0.946426 | 0.001108 | 0.945040 | 0.945695 | 0.946459 | 0.947096 | 0.947841 | 5.0 | 5442.206800 | 21.376667 | 5410.58700 | 5433.78700 | 5448.71800 | 5450.23400 | 5467.70800 | 5.0 | 3305.307000 | 38.370046 | 3268.89500 | 3281.71700 | 3294.01000 | 3315.09800 | 3366.81500 |

Compare the elapsed time with the default parameters:

In [26]:

```
knn_cv_elapsed = knn_cv[(knn_cv["n-bits"]==128)&(knn_cv["n-lshashes"]==4)&(knn_cv["transformer"]=="log1p")][["dataset", "index-time", "query-time"]].groupby("dataset").median()
knn_cv_elapsed
```

Out[26]:

|  | index-time | query-time |
| --- | --- | --- |
| dataset |  |  |
| Baron2016-human | 4.841887 | 0.937307 |
| Plass2018 | 13.392609 | 3.126437 |
| Shekhar2016 | 17.587253 | 4.225589 |
| TabulaMuris-chromium | 30.343434 | 6.577190 |

In [27]:

```
knn_cv_scmap_elapsed = knn_cv_scmap[(knn_cv_scmap["n-features"]==500)&(knn_cv_scmap["n-centroids-factor"]==1)][["dataset", "index-time", "query-time"]].groupby("dataset").median()
knn_cv_scmap_elapsed
```

Out[27]:

|  | index-time | query-time |
| --- | --- | --- |
| dataset |  |  |
| Baron2016-human | 57.34592 | 22.51334 |
| Plass2018 | 188.40800 | 135.60760 |
| Shekhar2016 | 262.53370 | 200.15250 |
| TabulaMuris-chromium | 661.88220 | 780.57090 |

In [28]:

```
knn_cv_scmap_elapsed / knn_cv_elapsed
```

Out[28]:

|  | index-time | query-time |
| --- | --- | --- |
| dataset |  |  |
| Baron2016-human | 11.843713 | 24.019179 |
| Plass2018 | 14.068058 | 43.374489 |
| Shekhar2016 | 14.927499 | 47.366769 |
| TabulaMuris-chromium | 21.813028 | 118.678473 |

Plot consistency scores, Cohen's kappa coefficients, index times, and query times:

In [29]:

```
metric_labels = {
    "consistency": "consistency",
    "cohen_kappa": "Cohen's kappa",
    "index-time": "index time [s]",
    "query-time": "query time [s]"}
transformer = "log1p"
for metric in ["consistency", "cohen_kappa", "index-time", "query-time"]:
    fig, axes = subplots(len(datasets), 2, dpi=dpi, figsize=(8, 7))
    for i, dataset in enumerate(datasets):
        if i > 0:
            axes[i,0].get_shared_x_axes().join(axes[0,0], axes[i,0])
            axes[i,1].get_shared_x_axes().join(axes[0,1], axes[i,1])
        axes[i,0].get_shared_y_axes().join(axes[i,0], axes[i,1])
        data = knn_cv[(knn_cv["dataset"]==dataset)&(knn_cv["transformer"]==transformer)]
        data_scmap = knn_cv_scmap[knn_cv_scmap["dataset"]==dataset]
        ymin = min(data[metric].min(), data_scmap[metric].min())
        ymax = max(data[metric].max(), data_scmap[metric].max())
        ydiff = ymax - ymin
        markerscale = 0.5
        labelspacing = 0.2
        if metric.endswith("-time"):
            seaborn.swarmplot(x="n-bits", hue="n-lshashes", y=metric, data=data, ax=axes[i,0], size=3, dodge=True)
            seaborn.swarmplot(x="n-centroids-factor", hue="n-features", y=metric, data=data_scmap, ax=axes[i,1], size=3, dodge=True)
            axes[i,0].set_ylim(0, None)
            axes[i,0].legend(title="n-lshashes", fontsize="small", markerscale=markerscale)
            axes[i,1].legend(title="n-features", fontsize="small", markerscale=markerscale, loc="upper left", labelspacing=labelspacing)
        else:
            axes[i,0].set_ylim(ymin - ydiff * 0.02, ymax + ydiff * 0.02)  # need to set ylim before plotting
            seaborn.swarmplot(x="n-bits", hue="n-lshashes", y=metric, data=data, ax=axes[i,0], size=3, dodge=True)
            seaborn.swarmplot(x="n-centroids-factor", hue="n-features", y=metric, data=data_scmap, ax=axes[i,1], size=3, dodge=True)
            axes[i,0].legend(title="n-lshashes", fontsize="small", markerscale=markerscale, loc="lower right")
            axes[i,1].legend(title="n-features", fontsize="small", markerscale=markerscale, labelspacing=labelspacing)
        if i > 0:
            axes[i,0].legend_.remove()
            axes[i,1].legend_.remove()
        axes[i,0].set_ylabel(dataset + "\n\n" + metric_labels[metric])
        axes[i,1].set_ylabel("")
        axes[i,1].set_yticklabels([])
        for j in range(2):
            axes[i,j].locator_params(axis="y", nbins=4)
            axes[i,j].grid(axis="y", alpha=0.3)
        if i < len(datasets) - 1:
            for j in range(2):
                axes[i,j].set_xlabel("")
                axes[i,j].set_xticklabels([])
    axes[0,0].set_title("CellFishing")
    axes[0,1].set_title("scmap-cell")
    seaborn.despine(fig=fig)
    fig.tight_layout(h_pad=0.2)
    savefig(f"selfmapping-{metric}")
```

Summarize the adjusted Rand scores from SC3.

In [30]:

```
data = []
for dataset in datasets:
    # The TOML files were broken due to the noisy outputs from SC3.
    cluster = clusters[dataset]
    k = len(cluster.unique())
    for r in range(5):
        sc3 = pandas.read_table(f"results/{dataset}.clustering-sc3/k-{k}.{r+1}.tsv.gz", index_col="cell")["cluster"]
        assert sc3.index.equals(cluster.index)
        score = sklearn.metrics.adjusted_rand_score(cluster, sc3)
        data.append((dataset, score))
sc3 = pandas.DataFrame(data, columns=["dataset", "adjusted_rand"])
sc3.groupby("dataset").describe()
```

Out[30]:

|  | adjusted\_rand | | | | | | | |
| --- | --- | --- | --- | --- | --- | --- | --- | --- |
|  | count | mean | std | min | 25% | 50% | 75% | max |
| dataset |  |  |  |  |  |  |  |  |
| Baron2016-human | 5.0 | 0.316196 | 0.037334 | 0.251702 | 0.317039 | 0.334107 | 0.334157 | 0.343975 |
| Plass2018 | 5.0 | 0.094676 | 0.012361 | 0.080149 | 0.084944 | 0.094701 | 0.104365 | 0.109220 |
| Shekhar2016 | 5.0 | 0.501870 | 0.029370 | 0.480690 | 0.484287 | 0.493581 | 0.497854 | 0.552937 |
| TabulaMuris-chromium | 5.0 | 0.377031 | 0.016006 | 0.359022 | 0.368149 | 0.371357 | 0.387877 | 0.398750 |

#### Feature selection¶

Compare default features:

In [31]:

```
def readlines(filename):
    with open(filename) as file:
        return [line.rstrip() for line in file.readlines()]

fig, axes = subplots(2, 2, dpi=dpi)
for i, dataset in enumerate(datasets):
    features = readlines(f"results/{dataset}.select-features.n_features-default.txt")
    features_scmap = readlines(f"results/{dataset}.select-features.scmap.n_features-500.txt")
    ax = axes[i//2,i%2]
    matplotlib_venn.venn2([set(features), set(features_scmap)], set_labels=("", ""), ax=ax)
    ax.set_title(dataset)
```

In [32]:

```
for dataset in datasets:
    fig, axes = subplots(2, 2, dpi=dpi)
    for i, n_features in enumerate([500, 1000, 2000, 4000]):
        features = readlines(f"results/{dataset}.select-features.n_features-{n_features}.txt")
        features_scmap = readlines(f"results/{dataset}.select-features.scmap.n_features-{n_features}.txt")
        ax = axes[i//2,i%2]
        matplotlib_venn.venn2([set(features), set(features_scmap)], set_labels=("", ""), ax=ax)
        ax.set_title(f"n-features = {n_features}")
    fig.suptitle(dataset)
```

In [33]:

```
knn_cv_features = pandas.DataFrame({"dataset": [], "consistency": [], "cohen_kappa": []})
for dataset in datasets:
    for n_features in [500, 1000, 2000, 4000]:
        filename = f"results/{dataset}.knn-cv-features.n_features-{n_features}.toml"
        out = toml.load(filename)
        for exp in out["experiment"]:
            row = cluster_metrics(exp["filename"], clusters[dataset])
            row["dataset"] = dataset
            row["n-features"] = n_features
            knn_cv_features = knn_cv_features.append(row, ignore_index=True)
```

In [34]:

```
knn_cv_features.groupby(["dataset", "n-features"]).describe()
```

Out[34]:

|  |  | adjusted\_rand | | | | | | | | cohen\_kappa | | | | | | | | consistency | | | | | | | |
| --- | --- | --- | --- | --- | --- | --- | --- | --- | --- | --- | --- | --- | --- | --- | --- | --- | --- | --- | --- | --- | --- | --- | --- | --- | --- |
|  |  | count | mean | std | min | 25% | 50% | 75% | max | count | mean | std | min | 25% | 50% | 75% | max | count | mean | std | min | 25% | 50% | 75% | max |
| dataset | n-features |  |  |  |  |  |  |  |  |  |  |  |  |  |  |  |  |  |  |  |  |  |  |  |  |
| Baron2016-human | 500.0 | 5.0 | 0.970701 | 0.001690 | 0.968915 | 0.969693 | 0.970013 | 0.971858 | 0.973027 | 5.0 | 0.976511 | 0.000546 | 0.975586 | 0.976453 | 0.976746 | 0.976883 | 0.976887 | 5.0 | 0.981141 | 0.000441 | 0.980394 | 0.981095 | 0.981328 | 0.981445 | 0.981445 |
|  | 1000.0 | 5.0 | 0.972843 | 0.002114 | 0.970722 | 0.971688 | 0.971703 | 0.974349 | 0.975754 | 5.0 | 0.979218 | 0.001112 | 0.977763 | 0.978637 | 0.979072 | 0.980093 | 0.980524 | 5.0 | 0.983312 | 0.000893 | 0.982145 | 0.982845 | 0.983195 | 0.984012 | 0.984362 |
|  | 2000.0 | 5.0 | 0.973687 | 0.001099 | 0.972155 | 0.973192 | 0.973648 | 0.974487 | 0.974951 | 5.0 | 0.979972 | 0.000692 | 0.979071 | 0.979509 | 0.980085 | 0.980379 | 0.980815 | 5.0 | 0.983919 | 0.000556 | 0.983195 | 0.983545 | 0.984012 | 0.984246 | 0.984596 |
|  | 4000.0 | 5.0 | 0.972447 | 0.001231 | 0.970676 | 0.972010 | 0.972565 | 0.972973 | 0.974009 | 5.0 | 0.979475 | 0.000474 | 0.978780 | 0.979356 | 0.979511 | 0.979646 | 0.980085 | 5.0 | 0.983522 | 0.000382 | 0.982962 | 0.983429 | 0.983545 | 0.983662 | 0.984012 |
| Plass2018 | 500.0 | 5.0 | 0.509908 | 0.003136 | 0.507406 | 0.508246 | 0.509128 | 0.509426 | 0.515336 | 5.0 | 0.644824 | 0.001258 | 0.642780 | 0.644489 | 0.645327 | 0.645655 | 0.645869 | 5.0 | 0.686480 | 0.001147 | 0.684758 | 0.686008 | 0.686702 | 0.687257 | 0.687674 |
|  | 1000.0 | 5.0 | 0.536415 | 0.003290 | 0.531521 | 0.535502 | 0.536471 | 0.538295 | 0.540285 | 5.0 | 0.680006 | 0.002802 | 0.675042 | 0.680732 | 0.681201 | 0.681236 | 0.681819 | 5.0 | 0.717102 | 0.002683 | 0.712382 | 0.717703 | 0.718120 | 0.718258 | 0.719045 |
|  | 2000.0 | 5.0 | 0.541364 | 0.005121 | 0.533522 | 0.539454 | 0.542508 | 0.544829 | 0.546507 | 5.0 | 0.690329 | 0.003282 | 0.685216 | 0.689019 | 0.691670 | 0.692364 | 0.693377 | 5.0 | 0.725689 | 0.002984 | 0.720988 | 0.724597 | 0.726957 | 0.727466 | 0.728438 |
|  | 4000.0 | 5.0 | 0.527451 | 0.003520 | 0.523325 | 0.525867 | 0.526280 | 0.529313 | 0.532470 | 5.0 | 0.677304 | 0.002755 | 0.673438 | 0.675538 | 0.678420 | 0.678901 | 0.680221 | 5.0 | 0.714196 | 0.002446 | 0.710809 | 0.712567 | 0.715112 | 0.715713 | 0.716778 |
| Shekhar2016 | 500.0 | 5.0 | 0.955886 | 0.001001 | 0.954821 | 0.955339 | 0.955763 | 0.956030 | 0.957479 | 5.0 | 0.921114 | 0.001961 | 0.919589 | 0.919719 | 0.920130 | 0.921947 | 0.924187 | 5.0 | 0.936281 | 0.001580 | 0.935052 | 0.935161 | 0.935489 | 0.936943 | 0.938761 |
|  | 1000.0 | 5.0 | 0.962997 | 0.000487 | 0.962261 | 0.962992 | 0.963041 | 0.963061 | 0.963630 | 5.0 | 0.937810 | 0.000950 | 0.936760 | 0.937087 | 0.937625 | 0.938646 | 0.938930 | 5.0 | 0.949751 | 0.000769 | 0.948907 | 0.949162 | 0.949598 | 0.950435 | 0.950653 |
|  | 2000.0 | 5.0 | 0.970651 | 0.001038 | 0.968908 | 0.970584 | 0.971077 | 0.971086 | 0.971598 | 5.0 | 0.955282 | 0.001478 | 0.953349 | 0.954220 | 0.955560 | 0.956408 | 0.956872 | 5.0 | 0.963846 | 0.001194 | 0.962290 | 0.962980 | 0.964071 | 0.964762 | 0.965126 |
|  | 4000.0 | 5.0 | 0.973257 | 0.000699 | 0.972481 | 0.972585 | 0.973396 | 0.973807 | 0.974017 | 5.0 | 0.962310 | 0.000333 | 0.961814 | 0.962138 | 0.962443 | 0.962531 | 0.962626 | 5.0 | 0.969526 | 0.000270 | 0.969126 | 0.969381 | 0.969635 | 0.969708 | 0.969781 |
| TabulaMuris-chromium | 500.0 | 5.0 | 0.954765 | 0.000844 | 0.953565 | 0.954530 | 0.954725 | 0.955135 | 0.955868 | 5.0 | 0.958826 | 0.000998 | 0.957554 | 0.957962 | 0.959392 | 0.959437 | 0.959786 | 5.0 | 0.961777 | 0.000926 | 0.960595 | 0.960977 | 0.962305 | 0.962341 | 0.962669 |
|  | 1000.0 | 5.0 | 0.959798 | 0.000980 | 0.958726 | 0.958731 | 0.960441 | 0.960450 | 0.960643 | 5.0 | 0.963745 | 0.000963 | 0.962551 | 0.963058 | 0.963766 | 0.964469 | 0.964881 | 5.0 | 0.966343 | 0.000895 | 0.965234 | 0.965707 | 0.966362 | 0.967017 | 0.967399 |
|  | 2000.0 | 5.0 | 0.962337 | 0.000589 | 0.961599 | 0.961877 | 0.962454 | 0.962744 | 0.963009 | 5.0 | 0.966110 | 0.000364 | 0.965720 | 0.965957 | 0.966055 | 0.966114 | 0.966704 | 5.0 | 0.968541 | 0.000337 | 0.968181 | 0.968399 | 0.968490 | 0.968545 | 0.969091 |
|  | 4000.0 | 5.0 | 0.959375 | 0.000646 | 0.958366 | 0.959264 | 0.959444 | 0.959698 | 0.960105 | 5.0 | 0.963932 | 0.000433 | 0.963234 | 0.963881 | 0.963961 | 0.964273 | 0.964311 | 5.0 | 0.966518 | 0.000402 | 0.965870 | 0.966471 | 0.966544 | 0.966835 | 0.966871 |

In [35]:

```
knn_cv_features_scmap = pandas.DataFrame({"dataset": [], "consistency": [], "cohen_kappa": []})
for dataset in datasets:
    for n_min_features in ["default", 500, 1000, 2000, 4000]:
        filename = f"results/{dataset}.knn-cv-features.scmap.n_features-{n_min_features}.toml"
        out = toml.load(filename)
        for exp in out["experiment"]:
            row = cluster_metrics(exp["filename"], clusters[dataset])
            row["dataset"] = dataset
            row["n-min-features"] = n_min_features
            with open(exp["features-file"]) as f:
                row["n-features"] = len(f.readlines())
            knn_cv_features_scmap = knn_cv_features_scmap.append(row, ignore_index=True)
```

In [36]:

```
knn_cv_features_scmap.groupby(["dataset", "n-min-features"]).describe()
```

Out[36]:

|  |  | adjusted\_rand | | | | | | | | cohen\_kappa | | | | | | | | consistency | | | | | | | | n-features | | | | | | | |
| --- | --- | --- | --- | --- | --- | --- | --- | --- | --- | --- | --- | --- | --- | --- | --- | --- | --- | --- | --- | --- | --- | --- | --- | --- | --- | --- | --- | --- | --- | --- | --- | --- | --- |
|  |  | count | mean | std | min | 25% | 50% | 75% | max | count | mean | std | min | 25% | 50% | 75% | max | count | mean | std | min | 25% | 50% | 75% | max | count | mean | std | min | 25% | 50% | 75% | max |
| dataset | n-min-features |  |  |  |  |  |  |  |  |  |  |  |  |  |  |  |  |  |  |  |  |  |  |  |  |  |  |  |  |  |  |  |  |
| Baron2016-human | 500 | 5.0 | 0.966448 | 0.002043 | 0.964448 | 0.964617 | 0.966524 | 0.967261 | 0.969392 | 5.0 | 0.974598 | 0.001191 | 0.973229 | 0.973959 | 0.974680 | 0.974689 | 0.976435 | 5.0 | 0.979624 | 0.000954 | 0.978527 | 0.979111 | 0.979694 | 0.979694 | 0.981095 | 5.0 | 503.0 | 0.0 | 503.0 | 503.0 | 503.0 | 503.0 | 503.0 |
|  | 1000 | 5.0 | 0.967865 | 0.001112 | 0.966641 | 0.966814 | 0.968087 | 0.968586 | 0.969196 | 5.0 | 0.975097 | 0.000576 | 0.974656 | 0.974660 | 0.974805 | 0.975388 | 0.975976 | 5.0 | 0.980044 | 0.000459 | 0.979694 | 0.979694 | 0.979811 | 0.980278 | 0.980745 | 5.0 | 1022.0 | 0.0 | 1022.0 | 1022.0 | 1022.0 | 1022.0 | 1022.0 |
|  | 2000 | 5.0 | 0.969115 | 0.002536 | 0.967012 | 0.967212 | 0.967599 | 0.971609 | 0.972143 | 5.0 | 0.976440 | 0.001075 | 0.975241 | 0.975536 | 0.976407 | 0.977289 | 0.977723 | 5.0 | 0.981118 | 0.000859 | 0.980161 | 0.980394 | 0.981095 | 0.981795 | 0.982145 | 5.0 | 2190.0 | 0.0 | 2190.0 | 2190.0 | 2190.0 | 2190.0 | 2190.0 |
|  | 4000 | 5.0 | 0.967560 | 0.002239 | 0.964334 | 0.966264 | 0.968452 | 0.968838 | 0.969913 | 5.0 | 0.974130 | 0.001589 | 0.971583 | 0.974215 | 0.974365 | 0.974512 | 0.975975 | 5.0 | 0.979274 | 0.001267 | 0.977244 | 0.979344 | 0.979461 | 0.979578 | 0.980745 | 5.0 | 4961.0 | 0.0 | 4961.0 | 4961.0 | 4961.0 | 4961.0 | 4961.0 |
|  | default | 5.0 | 0.969115 | 0.002536 | 0.967012 | 0.967212 | 0.967599 | 0.971609 | 0.972143 | 5.0 | 0.976440 | 0.001075 | 0.975241 | 0.975536 | 0.976407 | 0.977289 | 0.977723 | 5.0 | 0.981118 | 0.000859 | 0.980161 | 0.980394 | 0.981095 | 0.981795 | 0.982145 | 5.0 | 2190.0 | 0.0 | 2190.0 | 2190.0 | 2190.0 | 2190.0 | 2190.0 |
| Plass2018 | 500 | 5.0 | 0.429908 | 0.002227 | 0.427958 | 0.428105 | 0.429497 | 0.430583 | 0.433397 | 5.0 | 0.566058 | 0.001016 | 0.565120 | 0.565383 | 0.565530 | 0.566802 | 0.567457 | 5.0 | 0.634629 | 0.000771 | 0.634046 | 0.634092 | 0.634092 | 0.635249 | 0.635665 | 5.0 | 518.0 | 0.0 | 518.0 | 518.0 | 518.0 | 518.0 | 518.0 |
|  | 1000 | 5.0 | 0.420238 | 0.002672 | 0.416388 | 0.418669 | 0.421238 | 0.421935 | 0.422959 | 5.0 | 0.552447 | 0.001153 | 0.550789 | 0.552152 | 0.552404 | 0.552955 | 0.553938 | 5.0 | 0.624597 | 0.000915 | 0.623126 | 0.624468 | 0.624838 | 0.624977 | 0.625578 | 5.0 | 1030.0 | 0.0 | 1030.0 | 1030.0 | 1030.0 | 1030.0 | 1030.0 |
|  | 2000 | 5.0 | 0.400499 | 0.002198 | 0.398727 | 0.399129 | 0.399313 | 0.401330 | 0.403994 | 5.0 | 0.543788 | 0.001158 | 0.542741 | 0.542953 | 0.543370 | 0.544324 | 0.545550 | 5.0 | 0.617277 | 0.001301 | 0.615954 | 0.616509 | 0.616972 | 0.617620 | 0.619332 | 5.0 | 2291.0 | 0.0 | 2291.0 | 2291.0 | 2291.0 | 2291.0 | 2291.0 |
|  | 4000 | 5.0 | 0.382820 | 0.005084 | 0.376402 | 0.380831 | 0.382151 | 0.384479 | 0.390238 | 5.0 | 0.531169 | 0.003332 | 0.527454 | 0.528384 | 0.531518 | 0.532863 | 0.535626 | 5.0 | 0.607209 | 0.003152 | 0.603415 | 0.604757 | 0.607857 | 0.608736 | 0.611281 | 5.0 | 4764.0 | 0.0 | 4764.0 | 4764.0 | 4764.0 | 4764.0 | 4764.0 |
|  | default | 5.0 | 0.404710 | 0.004892 | 0.399044 | 0.400603 | 0.405344 | 0.407766 | 0.410793 | 5.0 | 0.536900 | 0.002900 | 0.531904 | 0.537084 | 0.537780 | 0.538596 | 0.539136 | 5.0 | 0.609856 | 0.002663 | 0.605219 | 0.610078 | 0.611096 | 0.611096 | 0.611790 | 5.0 | 3099.0 | 0.0 | 3099.0 | 3099.0 | 3099.0 | 3099.0 | 3099.0 |
| Shekhar2016 | 500 | 5.0 | 0.871708 | 0.003303 | 0.866421 | 0.871391 | 0.871913 | 0.873709 | 0.875107 | 5.0 | 0.824797 | 0.002975 | 0.819752 | 0.824557 | 0.825998 | 0.826613 | 0.827066 | 5.0 | 0.858111 | 0.002445 | 0.853958 | 0.857922 | 0.859159 | 0.859522 | 0.859995 | 5.0 | 537.0 | 0.0 | 537.0 | 537.0 | 537.0 | 537.0 | 537.0 |
|  | 1000 | 5.0 | 0.856159 | 0.006317 | 0.847249 | 0.854563 | 0.855984 | 0.858291 | 0.864708 | 5.0 | 0.791235 | 0.005184 | 0.784635 | 0.788293 | 0.790355 | 0.796142 | 0.796752 | 5.0 | 0.830663 | 0.004314 | 0.825230 | 0.828103 | 0.829957 | 0.834867 | 0.835158 | 5.0 | 1100.0 | 0.0 | 1100.0 | 1100.0 | 1100.0 | 1100.0 | 1100.0 |
|  | 2000 | 5.0 | 0.868315 | 0.003433 | 0.863342 | 0.867012 | 0.868213 | 0.871087 | 0.871921 | 5.0 | 0.804126 | 0.003338 | 0.799672 | 0.803118 | 0.803263 | 0.806066 | 0.808509 | 5.0 | 0.841209 | 0.002790 | 0.837485 | 0.840285 | 0.840576 | 0.842831 | 0.844867 | 5.0 | 2297.0 | 0.0 | 2297.0 | 2297.0 | 2297.0 | 2297.0 | 2297.0 |
|  | 4000 | 5.0 | 0.870023 | 0.007672 | 0.859545 | 0.866312 | 0.870752 | 0.873533 | 0.879975 | 5.0 | 0.809950 | 0.004928 | 0.804935 | 0.805345 | 0.810752 | 0.811990 | 0.816730 | 5.0 | 0.845776 | 0.004184 | 0.841594 | 0.841740 | 0.846540 | 0.847522 | 0.851486 | 5.0 | 4956.0 | 0.0 | 4956.0 | 4956.0 | 4956.0 | 4956.0 | 4956.0 |
|  | default | 5.0 | 0.880858 | 0.004425 | 0.875418 | 0.877652 | 0.881789 | 0.882738 | 0.886693 | 5.0 | 0.812287 | 0.004174 | 0.805515 | 0.811846 | 0.812776 | 0.815024 | 0.816275 | 5.0 | 0.847951 | 0.003412 | 0.842394 | 0.847558 | 0.848504 | 0.850067 | 0.851231 | 5.0 | 3270.0 | 0.0 | 3270.0 | 3270.0 | 3270.0 | 3270.0 | 3270.0 |
| TabulaMuris-chromium | 500 | 5.0 | 0.943385 | 0.000954 | 0.942008 | 0.943039 | 0.943378 | 0.944002 | 0.944500 | 5.0 | 0.951033 | 0.000711 | 0.950389 | 0.950468 | 0.950748 | 0.951564 | 0.951995 | 5.0 | 0.954533 | 0.000661 | 0.953936 | 0.954009 | 0.954263 | 0.955028 | 0.955428 | 5.0 | 500.0 | 0.0 | 500.0 | 500.0 | 500.0 | 500.0 | 500.0 |
|  | 1000 | 5.0 | 0.927357 | 0.000727 | 0.926680 | 0.926800 | 0.927190 | 0.927648 | 0.928467 | 5.0 | 0.940003 | 0.000298 | 0.939696 | 0.939772 | 0.939931 | 0.940218 | 0.940399 | 5.0 | 0.944290 | 0.000278 | 0.944003 | 0.944076 | 0.944221 | 0.944494 | 0.944658 | 5.0 | 1011.0 | 0.0 | 1011.0 | 1011.0 | 1011.0 | 1011.0 | 1011.0 |
|  | 2000 | 5.0 | 0.926411 | 0.001029 | 0.925457 | 0.925561 | 0.926058 | 0.927215 | 0.927765 | 5.0 | 0.935693 | 0.000719 | 0.934881 | 0.935301 | 0.935393 | 0.936319 | 0.936571 | 5.0 | 0.940277 | 0.000666 | 0.939527 | 0.939909 | 0.940000 | 0.940855 | 0.941092 | 5.0 | 2026.0 | 0.0 | 2026.0 | 2026.0 | 2026.0 | 2026.0 | 2026.0 |
|  | 4000 | 5.0 | 0.931205 | 0.001206 | 0.929868 | 0.930049 | 0.931614 | 0.931829 | 0.932667 | 5.0 | 0.939276 | 0.001036 | 0.938337 | 0.938358 | 0.939216 | 0.939629 | 0.940838 | 5.0 | 0.943603 | 0.000965 | 0.942729 | 0.942747 | 0.943548 | 0.943930 | 0.945058 | 5.0 | 4019.0 | 0.0 | 4019.0 | 4019.0 | 4019.0 | 4019.0 | 4019.0 |
|  | default | 5.0 | 0.924817 | 0.000821 | 0.923693 | 0.924331 | 0.924966 | 0.925312 | 0.925782 | 5.0 | 0.933905 | 0.001024 | 0.932865 | 0.933101 | 0.934036 | 0.934059 | 0.935462 | 5.0 | 0.938603 | 0.000954 | 0.937635 | 0.937854 | 0.938727 | 0.938745 | 0.940055 | 5.0 | 2363.0 | 0.0 | 2363.0 | 2363.0 | 2363.0 | 2363.0 | 2363.0 |

#### Cluster-specific scores¶

In [37]:

```
import collections

def count_assignments(cell, nn, cluster):
    consistent = cluster[cell].values == cluster[nn].values
    cluster = cluster[cell]
    # make counters
    count_all = collections.Counter(cluster)
    count_consistent = collections.Counter({c: 0 for c in count_all.keys()})
    count_consistent.update(cluster[consistent])
    count_inconsistent = collections.Counter({c: 0 for c in count_all.keys()})
    count_inconsistent.update(cluster[~consistent])
    # join counts
    counts = pandas.DataFrame({"all": pandas.Series(count_all), "consistent": pandas.Series(count_consistent), "inconsistent": pandas.Series(count_inconsistent)})
    counts = counts.assign(consistency=lambda x: x["consistent"] / x["all"])[["all", "consistent", "inconsistent", "consistency"]]
    counts.sort_values(by="all", ascending=False, inplace=True)
    counts.index.name = "cluster"
    return counts


def plot_assignments_matrix(cell, nn, cluster, cluster2=None, overlapping=False, figsize=(12,12)):
    cluster1 = cluster
    if cluster2 is None:
        cluster2 = cluster1
    labels1 = cluster1.value_counts().index.values
    labels2 = cluster2.value_counts().index.values
    if overlapping:
        labels2 = labels1
    assignments = pandas.crosstab(
        pandas.Categorical(cluster1[cell].values, categories=labels1),
        pandas.Categorical(cluster2[nn].values, categories=labels2),
        rownames=["query"],
        colnames=["neighbor"],
        dropna=False,
        normalize="index")
    assignments = assignments.loc[labels1,labels2]
    fig, ax = subplots(dpi=dpi, figsize=figsize)
    im = ax.imshow(assignments, vmin=0, vmax=1)
    ax.set_ylabel("Query's label")
    ax.set_xlabel("Neighbor's label")
    ax.set_yticks(numpy.arange(assignments.shape[0]))
    ax.set_xticks(numpy.arange(assignments.shape[1]))
    ax.set_yticklabels(assignments.index)
    ax.set_xticklabels(assignments.columns, rotation=90)
    ax.set_yticks(numpy.arange(assignments.shape[0]+1)-.5, minor=True)
    ax.set_xticks(numpy.arange(assignments.shape[1]+1)-.5, minor=True)
    ax.grid(False)  # turn off seaborn's grid
    ax.grid(which="minor", color="gray", linestyle='-', linewidth=0.8)
    ax.tick_params(which="minor", bottom=False, left=False)
    ax.tick_params(length=0)
    cbar = fig.colorbar(im, fraction=0.046, pad=0.02, shrink=0.3, aspect=30)
    cbar.outline.set_visible(False)
    for x in ["left", "right", "top", "bottom"]:
        ax.spines[x].set_visible(False)
    fig.tight_layout()
    return fig, ax
```

In [38]:

```
# wrap long labels
def wraplabel(label, width=16):
    return "\n".join(textwrap.wrap(label.replace("_", " "), width=width))
```

In [39]:

```
for dataset in datasets:
    print(dataset)
    cluster = clusters[dataset]
    # CellFishing
    filenames = knn_cv[(knn_cv["dataset"]==dataset)&(knn_cv["n-bits"]==128)&(knn_cv["n-lshashes"]==4)&(knn_cv["transformer"]=="log1p")]["filename"]
    knn = [load_knn(x) for x in filenames]
    counts1 = [count_assignments(x.index, x["N1"], cluster) for x in knn]
    fig, ax = plot_assignments_matrix(knn[0].index, knn[0]["N1"], cluster)
    savefig(f"{dataset}-selfmapping-matrix")
    # scmap
    filenames = knn_cv_scmap[(knn_cv_scmap["dataset"]==dataset)&(knn_cv_scmap["n-features"]==500)&(knn_cv_scmap["n-centroids-factor"]==1)]["filename"]
    knn = [load_knn(x) for x in filenames]
    counts2 = [count_assignments(x.index, x["N1"], cluster) for x in knn]
    fig, ax = plot_assignments_matrix(knn[0].index, knn[0]["N1"], cluster)
    savefig(f"{dataset}-selfmapping-matrix-scmap")
    # merge
    x = pandas.concat([x["consistency"].reset_index() for x in counts1], ignore_index=True)
    x["method"] = "CellFishing"
    y = pandas.concat([x["consistency"].reset_index() for x in counts2], ignore_index=True)
    y["method"] = "scmap-cell";
    data = pandas.concat([x, y], ignore_index=True)
    cluster_unique = cluster.value_counts().index.values
    n_cols = 3
    y_max = math.ceil(len(cluster_unique) / n_cols)
    fig, axes = subplots(1, n_cols, dpi=dpi, figsize=(8, math.ceil(y_max / 2)))
    for j in range(n_cols):
        if n_cols == 1:
            ax = axes
        else:
            ax = axes[j]
        ax.set_xlim(0, 1.02)
        ax.set_xticks([0, 0.2, 0.4, 0.6, 0.8, 1.0])
        y = cluster_unique[j*y_max:min((j+1)*y_max, len(cluster_unique))]
        seaborn.swarmplot(x="consistency", y="cluster", hue="method", data=data[data["cluster"].isin(y)], order=y, orient="h", dodge=False, size=2.4, ax=ax)
        ax.tick_params(labelsize="small")
        ax.set_ylabel("")
        ax.set_yticklabels([wraplabel(x.get_text()) for x in ax.get_yticklabels()])
        ax.grid(axis="x", alpha=0.3)
        if j > 0:
            ax.legend_.remove()
        else:
            ax.legend(loc="upper left", fontsize="small").set_title("")
    fig.tight_layout(w_pad=0.5)
    seaborn.despine(fig=fig)
    savefig(f"{dataset}-selfmapping-clusters")
```

```
Baron2016-human
Shekhar2016
Plass2018
TabulaMuris-chromium
```

#### Similarities from nearest neighbors¶

In [40]:

```
def plot_similarities(dataset, cluster, targets):
    fig, axes = subplots(1, len(targets), dpi=dpi, figsize=(8, 4), sharey=True)
    for i, target in enumerate(targets):
        target_escaped = target.replace('/', ';')
        log = toml.load(f"results/{dataset}.knn-similarities.target-{target_escaped}.toml")
        similarities_with_target_cells = pandas.read_table(log["experiment"][0]["filename-similarities-with-target-cells"], index_col="cell")
        similarities_without_target_cells = pandas.read_table(log["experiment"][0]["filename-similarities-without-target-cells"], index_col="cell")
        cells_target = cluster[(cluster == target).values].index.values
        #cells_nontarget = cluster[(cluster != target).values].index.values
        # melt data
        df1 = similarities_with_target_cells.loc[cells_target].reset_index().melt(id_vars="cell", var_name="rank", value_name="similarity")
        df1["removed"] = False
        df2 = similarities_without_target_cells.loc[cells_target].reset_index().melt(id_vars="cell", var_name="rank", value_name="similarity")
        df2["removed"] = True
        df = df1.append(df2)
        df["rank"] = [int(x[1]) for x in df["rank"]]
        df = df[df["rank"]<=3]
        # plot
        ax = axes[i]
        seaborn.violinplot(x="rank", y="similarity", hue="removed", data=df, split=True, inner="quart", linewidth=0.6, ax=ax)
        #seaborn.boxplot(x="rank", y="similarity", hue="removed", data=df, linewidth=0.8, fliersize=0.5, ax=ax)
        ax.set_title(wraplabel(target), fontsize="small")
        ax.legend(loc="lower left")
        ax.legend_.remove()
        ax.grid(axis="y", alpha=0.3)
        if i > 0:
            ax.set_ylabel("")
    axes[0].legend(title="removed", loc="lower left", fontsize="small", title_fontsize="small")
    seaborn.despine(fig=fig)
    fig.tight_layout()
    savefig(f"{dataset}-selfmapping-remove")


def list_nearest_neighbors(dataset, cluster, targets):
    for target in targets:
        target_escaped = target.replace('/', ';')
        log = toml.load(f"results/{dataset}.knn-similarities.target-{target_escaped}.toml")
        #knn_with_target_cells = pandas.read_table(log["experiment"][0]["filename-knn-with-target-cells"], index_col="cell")
        knn_without_target_cells = pandas.read_table(log["experiment"][0]["filename-knn-without-target-cells"], index_col="cell")
        cells_target = cluster[(cluster == target).values].index.values
        value_counts = cluster.loc[knn_without_target_cells.loc[cells_target,"N1"]].value_counts()
        print(target)
        for i, (label, n) in enumerate(value_counts.items()):
            if i > 4:
                break
            print("    {}: {}".format(label, n))
```

In [41]:

```
dataset = "Shekhar2016"
targets = ["BC2", "BC5D", "BC3A", "BC5B", "BC4", "BC8/9 (mixture of BC8 and BC9)", "AC (Amacrine cell)"]
plot_similarities(dataset, clusters[dataset], targets)
```

In [42]:

```
list_nearest_neighbors(dataset, clusters[dataset], targets)
```

```
BC2
    BC3A: 227
    BC1B: 203
    BC6: 67
    BC1A: 41
    BC3B: 5
BC5D
    BC5A (Cone Bipolar cell 5A): 483
    BC5B: 45
    Doublets/Contaminants: 7
    BC1B: 4
    BC7 (Cone Bipolar cell 7): 3
BC3A
    BC2: 265
    BC3B: 86
    BC1B: 52
    BC1A: 40
    BC6: 26
BC5B
    BC5C: 282
    BC5D: 147
    BC5A (Cone Bipolar cell 5A): 18
    BC3B: 6
    Doublets/Contaminants: 6
BC4
    BC3B: 344
    BC1A: 37
    BC3A: 5
    BC5C: 3
    Doublets/Contaminants: 3
BC8/9 (mixture of BC8 and BC9)
    BC6: 106
    BC7 (Cone Bipolar cell 7): 96
    BC1A: 46
    Doublets/Contaminants: 12
    BC5C: 9
AC (Amacrine cell)
    BC5D: 76
    BC2: 44
    BC1A: 44
    BC8/9 (mixture of BC8 and BC9): 13
    BC4: 12
```

In [43]:

```
dataset = "TabulaMuris-chromium"
targets = ["basal cell", "alveolar macrophage", "fibroblast", "immature B cell", "basophil", "kidney cell", "endothelial cell of hepatic sinusoid"]
plot_similarities(dataset, clusters[dataset], targets)
```

In [44]:

```
list_nearest_neighbors(dataset, clusters[dataset], targets)
```

```
basal cell
    luminal epithelial cell of mammary gland: 342
    stromal cell: 36
    skeletal muscle satellite cell: 12
    endothelial cell: 1
    neuroendocrine cell: 1
alveolar macrophage
    non-classical monocyte: 117
    classical monocyte: 117
    blood cell: 63
    promonocyte: 22
    macrophage: 9
fibroblast
    stromal cell: 175
    mesenchymal cell: 20
    mesenchymal stem cell: 18
    kidney collecting duct epithelial cell: 8
    endothelial cell: 4
immature B cell
    B cell: 91
    late pro-B cell: 10
    early pro-B cell: 10
    macrophage: 1
    T cell: 1
basophil
    hematopoietic precursor cell: 31
    mast cell: 19
    macrophage: 4
    promonocyte: 3
    granulocytopoietic cell: 1
kidney cell
    macrophage: 28
    blood cell: 5
    non-classical monocyte: 5
    leukocyte: 3
    Fraction A pre-pro B cell: 1
endothelial cell of hepatic sinusoid
    leukocyte: 11
    lung endothelial cell: 7
    hepatocyte: 4
    endothelial cell: 3
    kidney capillary endothelial cell: 3
```

#### Detecting DEGs¶

In [45]:

```
def load_degs(cluster, target, k):
    knns = pandas.read_table(f"results/TabulaMuris-chromium.knn-degs.target-{target}/knn.k-{k}.tsv.gz", index_col="cell")
    degs = pandas.read_table(f"results/TabulaMuris-chromium.knn-degs.target-{target}/degs.k-{k}.tsv.gz", index_col=["cell", "gene"])
    return degs.join(cluster).join(knns["N1"]).join(cluster.rename_axis("N1"), on="N1", rsuffix="N1").drop("N1", axis=1)


def top_genes(by, n=10000, upper=-4):
    def fetch(df):
        x = df.sort_values(by).head(n)
        return x[x[by] <= upper].reset_index("cell", drop=True)
    return fetch


def make_crosstable(cluster, celltypes, target, k, by):
    degs = load_degs(cluster, target, k)
    x = degs.groupby("cell").apply(top_genes(by)).reset_index("gene")
    return pandas.crosstab(x["gene"], x["clusterN1"]).loc[:,celltypes]
```

In [46]:

```
celltype_query = "immature B cell"
celltypes = ['early pro-B cell', 'late pro-B cell', 'B cell',]
cluster = clusters["TabulaMuris-chromium"]
```

In [47]:

```
load_degs(cluster, celltype_query, 5).groupby("cell").first()["clusterN1"].value_counts()
```

Out[47]:

```
B cell                          91
late pro-B cell                 10
early pro-B cell                 9
hematopoietic precursor cell     1
dendritic cell                   1
natural killer cell              1
Name: clusterN1, dtype: int64
```

List top DEGs.

In [48]:

```
for celltype in celltypes:
    print(celltype)
    print("-"*len(celltype))
    for k in [5, 10, 20]:
        print(f"k = {k}")
        negative = make_crosstable(cluster, celltypes, celltype_query, k, "negative")
        positive = make_crosstable(cluster, celltypes, celltype_query, k, "positive")
        print("  negative: ", end="")
        degs = negative.sort_values(celltype, ascending=False)[celltype]
        for i in range(10):
            if degs.iloc[i] == 0:
                break
            print("{:s} {:d}".format(degs.index[i], degs.iloc[i]), end=", ")
        print()
        print("  positive: ", end="")
        degs = positive.sort_values(celltype, ascending=False)[celltype]
        for i in range(10):
            if degs.iloc[i] == 0:
                break
            print("{:s} {:d}".format(degs.index[i], degs.iloc[i]), end=", ")
        print()
    print()
```

```
early pro-B cell
----------------
k = 5
  negative: Vpreb1 9, Igll1 9, Dntt 6, Vpreb3 1, Ucp2 1, Crip1 1, Malat1 1, Rps24 1, Gmfg 1, 
  positive: Ly6d 6, Malat1 5, Dnajc7 5, Tmsb10 5, Klf2 4, Cd74 4, Btg1 3, Myl4 3, Herpud1 3, Tmsb4x 3, 
k = 10
  negative: Vpreb1 9, Igll1 9, Dntt 8, Crip1 3, Rps24 1, Ucp2 1, 
  positive: Dnajc7 8, Ly6d 6, Malat1 5, Tmsb10 5, Klf2 4, Cd74 4, Herpud1 4, Myl4 4, Btg1 3, Tmsb4x 3, 
k = 20
  negative: Vpreb1 9, Igll1 9, Dntt 9, Crip1 3, Ucp2 1, 
  positive: Dnajc7 8, Ly6d 6, Malat1 5, Tmsb10 5, Myl4 4, Klf2 4, Cd74 4, Rgs2 3, Cpm 3, Tmsb4x 3, 

late pro-B cell
---------------
k = 5
  negative: Hmgb2 5, Ptma 2, Hist1h2ao 2, Tuba1b 1, H2afz 1, Stmn1 1, 
  positive: Malat1 6, Tmsb4x 4, Dnajc7 3, Tmsb10 3, Actb 3, Ly6d 2, Rsph1 1, Hist1h1e 1, Hn1 1, Calm1 1, 
k = 10
  negative: Hmgb2 6, Ptma 2, H2afz 2, Tuba1b 1, Stmn1 1, Hist1h2ao 1, 
  positive: Malat1 7, Tmsb4x 4, Dnajc7 3, Tmsb10 3, Actb 3, Ly6d 2, Jun 2, Ets2 1, Hist1h1c 1, Hist1h1e 1, 
k = 20
  negative: Hmgb2 6, Ptma 2, H2afz 2, Hist1h2ao 2, 2810417H13Rik 1, Tuba1b 1, Stmn1 1, Rplp1 1, 
  positive: Malat1 8, Dnajc7 4, Tmsb4x 4, Actb 3, Btg2 3, Tmsb10 3, Ly6d 2, Ifi27l2a 2, Hist1h1c 1, Mef2c 1, 

B cell
------
k = 5
  negative: Cd74 36, H2-Ab1 20, H2-Eb1 14, Malat1 13, H2-Aa 13, Actb 11, Rps28 8, Tmsb4x 7, Tmsb10 5, Rn45s 5, 
  positive: Beta-s 81, S100a8 76, S100a9 73, Fos 30, Junb 24, Klf2 22, Xist 19, Jun 19, Malat1 16, Camp 15, 
k = 10
  negative: Cd74 41, H2-Ab1 19, H2-Aa 15, H2-Eb1 15, Malat1 15, Actb 10, Rps28 10, Tmsb10 9, Tmsb4x 7, Rn45s 7, 
  positive: S100a8 83, Beta-s 81, S100a9 76, Fos 36, Camp 30, Klf2 25, Junb 23, Jun 19, Ngp 19, Xist 18, 
k = 20
  negative: Cd74 45, H2-Ab1 22, H2-Eb1 16, H2-Aa 16, Rps28 13, Malat1 11, Tmsb10 8, Actb 8, Rn45s 7, Tmsb4x 7, 
  positive: S100a8 85, Beta-s 84, S100a9 80, Fos 41, Klf2 27, Camp 27, Jun 25, Junb 22, Ngp 21, Malat1 19,
```

Generate LaTeX tables.

In [49]:

```
n = 6  # the number of top DEGs

def capitalize(s):
    return s[0].upper() + s[1:]

for k in [5, 10, 20]:
    negative = make_crosstable(cluster, celltypes, celltype_query, k, "negative")
    positive = make_crosstable(cluster, celltypes, celltype_query, k, "positive")
    print(f"% k = {k}")
    print(r"\begin{tabular}{" + "lr" * (len(celltypes) * 2) + "}")
    print(r"\toprule")

    for j, celltype in enumerate(celltypes):
        print(r"\multicolumn{4}{c}{", capitalize(celltype), r"}", end=" & " if j < len(celltypes)-1 else r" \\")
    print()
    for j in range(len(celltypes)):
        print(r"\cmidrule(lr){", 4 * j + 1, "-", 4 * j + 4, r"}", end=" ")
    print()

    for j in range(len(celltypes)):
        print(r"\multicolumn{2}{c}{Negative} & \multicolumn{2}{c}{Positive}", end=" & " if j < len(celltypes)-1 else r" \\")
    print()
    for j in range(len(celltypes)):
        print(r"\cmidrule(lr){", 4 * j + 1, "-", 4 * j + 2, r"}", end=" ")
        print(r"\cmidrule(lr){", 4 * j + 3, "-", 4 * j + 4, r"}", end=" ")
    print()

    # Gene count
    for i in range(n):
        for j, celltype in enumerate(celltypes):
            df = negative.sort_values(celltype)
            row = df.iloc[-(i+1)]
            print("\\textit{{{:10s}}} & {:3d}".format(row.name if row[celltype] > 0 else "--", row[celltype]), end=" & ")
            df = positive.sort_values(celltype)
            row = df.iloc[-(i+1)]
            print("\\textit{{{:10s}}} & {:3d}".format(row.name if row[celltype] > 0 else "--", row[celltype]), end=" & " if j < len(celltypes)-1 else r" \\")
        print()

    # Total count
    print(r"\addlinespace")
    for j, celltype in enumerate(celltypes):
        print("Total & {:3d}".format(negative[celltype].sum()), end=" & ")
        print("Total & {:3d}".format(positive[celltype].sum()), end=" & " if j < len(celltypes)-1 else r" \\")
    
    print()
    print(r"\bottomrule")
    print(r"\end{tabular}")
    print()
```

```
% k = 5
\begin{tabular}{lrlrlrlrlrlr}
\toprule
\multicolumn{4}{c}{ Early pro-B cell } & \multicolumn{4}{c}{ Late pro-B cell } & \multicolumn{4}{c}{ B cell } \\
\cmidrule(lr){ 1 - 4 } \cmidrule(lr){ 5 - 8 } \cmidrule(lr){ 9 - 12 } 
\multicolumn{2}{c}{Negative} & \multicolumn{2}{c}{Positive} & \multicolumn{2}{c}{Negative} & \multicolumn{2}{c}{Positive} & \multicolumn{2}{c}{Negative} & \multicolumn{2}{c}{Positive} \\
\cmidrule(lr){ 1 - 2 } \cmidrule(lr){ 3 - 4 } \cmidrule(lr){ 5 - 6 } \cmidrule(lr){ 7 - 8 } \cmidrule(lr){ 9 - 10 } \cmidrule(lr){ 11 - 12 } 
\textit{Vpreb1    } &   9 & \textit{Ly6d      } &   6 & \textit{Hmgb2     } &   5 & \textit{Malat1    } &   6 & \textit{Cd74      } &  36 & \textit{Beta-s    } &  81 \\
\textit{Igll1     } &   9 & \textit{Malat1    } &   5 & \textit{Ptma      } &   2 & \textit{Tmsb4x    } &   4 & \textit{H2-Ab1    } &  20 & \textit{S100a8    } &  76 \\
\textit{Dntt      } &   6 & \textit{Dnajc7    } &   5 & \textit{Hist1h2ao } &   2 & \textit{Tmsb10    } &   3 & \textit{H2-Eb1    } &  14 & \textit{S100a9    } &  73 \\
\textit{Vpreb3    } &   1 & \textit{Tmsb10    } &   5 & \textit{Tuba1b    } &   1 & \textit{Dnajc7    } &   3 & \textit{H2-Aa     } &  13 & \textit{Fos       } &  30 \\
\textit{Crip1     } &   1 & \textit{Klf2      } &   4 & \textit{Stmn1     } &   1 & \textit{Actb      } &   3 & \textit{Malat1    } &  13 & \textit{Junb      } &  24 \\
\textit{Rps24     } &   1 & \textit{Cd74      } &   4 & \textit{H2afz     } &   1 & \textit{Ly6d      } &   2 & \textit{Actb      } &  11 & \textit{Klf2      } &  22 \\
\addlinespace
Total &  30 & Total &  74 & Total &  12 & Total &  38 & Total & 167 & Total & 661 \\
\bottomrule
\end{tabular}

% k = 10
\begin{tabular}{lrlrlrlrlrlr}
\toprule
\multicolumn{4}{c}{ Early pro-B cell } & \multicolumn{4}{c}{ Late pro-B cell } & \multicolumn{4}{c}{ B cell } \\
\cmidrule(lr){ 1 - 4 } \cmidrule(lr){ 5 - 8 } \cmidrule(lr){ 9 - 12 } 
\multicolumn{2}{c}{Negative} & \multicolumn{2}{c}{Positive} & \multicolumn{2}{c}{Negative} & \multicolumn{2}{c}{Positive} & \multicolumn{2}{c}{Negative} & \multicolumn{2}{c}{Positive} \\
\cmidrule(lr){ 1 - 2 } \cmidrule(lr){ 3 - 4 } \cmidrule(lr){ 5 - 6 } \cmidrule(lr){ 7 - 8 } \cmidrule(lr){ 9 - 10 } \cmidrule(lr){ 11 - 12 } 
\textit{Vpreb1    } &   9 & \textit{Dnajc7    } &   8 & \textit{Hmgb2     } &   6 & \textit{Malat1    } &   7 & \textit{Cd74      } &  41 & \textit{S100a8    } &  83 \\
\textit{Igll1     } &   9 & \textit{Ly6d      } &   6 & \textit{Ptma      } &   2 & \textit{Tmsb4x    } &   4 & \textit{H2-Ab1    } &  19 & \textit{Beta-s    } &  81 \\
\textit{Dntt      } &   8 & \textit{Malat1    } &   5 & \textit{H2afz     } &   2 & \textit{Actb      } &   3 & \textit{H2-Eb1    } &  15 & \textit{S100a9    } &  76 \\
\textit{Crip1     } &   3 & \textit{Tmsb10    } &   5 & \textit{Tuba1b    } &   1 & \textit{Dnajc7    } &   3 & \textit{Malat1    } &  15 & \textit{Fos       } &  36 \\
\textit{Rps24     } &   1 & \textit{Cd74      } &   4 & \textit{Stmn1     } &   1 & \textit{Tmsb10    } &   3 & \textit{H2-Aa     } &  15 & \textit{Camp      } &  30 \\
\textit{Ucp2      } &   1 & \textit{Herpud1   } &   4 & \textit{Hist1h2ao } &   1 & \textit{Ly6d      } &   2 & \textit{Rps28     } &  10 & \textit{Klf2      } &  25 \\
\addlinespace
Total &  31 & Total &  91 & Total &  13 & Total &  47 & Total & 182 & Total & 752 \\
\bottomrule
\end{tabular}

% k = 20
\begin{tabular}{lrlrlrlrlrlr}
\toprule
\multicolumn{4}{c}{ Early pro-B cell } & \multicolumn{4}{c}{ Late pro-B cell } & \multicolumn{4}{c}{ B cell } \\
\cmidrule(lr){ 1 - 4 } \cmidrule(lr){ 5 - 8 } \cmidrule(lr){ 9 - 12 } 
\multicolumn{2}{c}{Negative} & \multicolumn{2}{c}{Positive} & \multicolumn{2}{c}{Negative} & \multicolumn{2}{c}{Positive} & \multicolumn{2}{c}{Negative} & \multicolumn{2}{c}{Positive} \\
\cmidrule(lr){ 1 - 2 } \cmidrule(lr){ 3 - 4 } \cmidrule(lr){ 5 - 6 } \cmidrule(lr){ 7 - 8 } \cmidrule(lr){ 9 - 10 } \cmidrule(lr){ 11 - 12 } 
\textit{Vpreb1    } &   9 & \textit{Dnajc7    } &   8 & \textit{Hmgb2     } &   6 & \textit{Malat1    } &   8 & \textit{Cd74      } &  45 & \textit{S100a8    } &  85 \\
\textit{Igll1     } &   9 & \textit{Ly6d      } &   6 & \textit{Ptma      } &   2 & \textit{Dnajc7    } &   4 & \textit{H2-Ab1    } &  22 & \textit{Beta-s    } &  84 \\
\textit{Dntt      } &   9 & \textit{Tmsb10    } &   5 & \textit{Hist1h2ao } &   2 & \textit{Tmsb4x    } &   4 & \textit{H2-Eb1    } &  16 & \textit{S100a9    } &  80 \\
\textit{Crip1     } &   3 & \textit{Malat1    } &   5 & \textit{H2afz     } &   2 & \textit{Tmsb10    } &   3 & \textit{H2-Aa     } &  16 & \textit{Fos       } &  41 \\
\textit{Ucp2      } &   1 & \textit{Cd74      } &   4 & \textit{2810417H13Rik} &   1 & \textit{Actb      } &   3 & \textit{Rps28     } &  13 & \textit{Camp      } &  27 \\
\textit{--        } &   0 & \textit{Myl4      } &   4 & \textit{Stmn1     } &   1 & \textit{Btg2      } &   3 & \textit{Malat1    } &  11 & \textit{Klf2      } &  27 \\
\addlinespace
Total &  31 & Total &  94 & Total &  16 & Total &  57 & Total & 189 & Total & 772 \\
\bottomrule
\end{tabular}
```

#### Mapping across batches¶

In [50]:

```
knn_batch_shekhar2016 = pandas.DataFrame()
knn_batch_shekhar2016_scmap = pandas.DataFrame()
cluster = clusters["Shekhar2016"]
for query in ["1", "2"]:
    print(f"query = {query}")
    # CellFishing
    log = toml.load(f"results/Shekhar2016.knn-batch.batch-{query}.toml")
    print(log["date-time"])
    print(log["version-info"])
    exp = pandas.DataFrame(log["experiment"])
    exp["query"] = query
    exp = pandas.concat(
        [exp,
         pandas.DataFrame([cluster_metrics(x, cluster) for x in exp["filename"]])],
        axis=1)
    knn_batch_shekhar2016 = knn_batch_shekhar2016.append(exp, ignore_index=True)
    # scmap
    log = toml.load(f"results/Shekhar2016.knn-batch.scmap.batch-{query}.toml")
    print(log["date-time"])
    print(log["session-info"])
    exp = pandas.DataFrame(log["experiment"])
    exp["query"] = query
    exp = pandas.concat(
        [exp,
         pandas.DataFrame([cluster_metrics(x, cluster) for x in exp["filename"]])],
        axis=1)
    knn_batch_shekhar2016_scmap = knn_batch_shekhar2016_scmap.append(exp, ignore_index=True)
```

```
query = 1
2018-11-23 16:42:12.057000
Julia Version 1.0.1
Commit 0d713926f8 (2018-09-29 19:05 UTC)
Platform Info:
  OS: Linux (x86_64-pc-linux-gnu)
  CPU: Intel(R) Xeon(R) Gold 6126 CPU @ 2.60GHz
  WORD_SIZE: 64
  LIBM: libopenlibm
  LLVM: libLLVM-6.0.0 (ORCJIT, skylake)
Environment:
  JULIA_PROJECT = @.

2018-07-20 01:20:39
R version 3.5.0 (2018-04-23)
Platform: x86_64-pc-linux-gnu (64-bit)
Running under: Debian GNU/Linux 9 (stretch)

Matrix products: default
BLAS: /usr/lib/openblas-base/libblas.so.3
LAPACK: /usr/lib/libopenblasp-r0.2.19.so

locale:
 [1] LC_CTYPE=en_US.UTF-8       LC_NUMERIC=C              
 [3] LC_TIME=en_US.UTF-8        LC_COLLATE=en_US.UTF-8    
 [5] LC_MONETARY=en_US.UTF-8    LC_MESSAGES=C             
 [7] LC_PAPER=en_US.UTF-8       LC_NAME=C                 
 [9] LC_ADDRESS=C               LC_TELEPHONE=C            
[11] LC_MEASUREMENT=en_US.UTF-8 LC_IDENTIFICATION=C       

attached base packages:
[1] parallel  stats4    stats     graphics  grDevices utils     datasets 
[8] methods   base     

other attached packages:
 [1] h5_0.9.9                    scmap_1.2.0                
 [3] scater_1.8.0                ggplot2_2.2.1              
 [5] SingleCellExperiment_1.2.0  SummarizedExperiment_1.10.1
 [7] DelayedArray_0.6.1          BiocParallel_1.14.1        
 [9] matrixStats_0.53.1          Biobase_2.40.0             
[11] GenomicRanges_1.32.3        GenomeInfoDb_1.16.0        
[13] IRanges_2.14.10             S4Vectors_0.18.3           
[15] BiocGenerics_0.26.0         readr_1.1.1                

loaded via a namespace (and not attached):
 [1] viridis_0.5.1            edgeR_3.22.2             jsonlite_1.5            
 [4] viridisLite_0.3.0        DelayedMatrixStats_1.2.0 shiny_1.1.0             
 [7] assertthat_0.2.0         GenomeInfoDbData_1.1.0   vipor_0.4.5             
[10] pillar_1.2.2             lattice_0.20-35          glue_1.2.0              
[13] limma_3.36.1             digest_0.6.15            promises_1.0.1          
[16] XVector_0.20.0           randomForest_4.6-14      colorspace_1.3-2        
[19] htmltools_0.3.6          httpuv_1.4.3             Matrix_1.2-14           
[22] plyr_1.8.4               pkgconfig_2.0.1          zlibbioc_1.26.0         
[25] purrr_0.2.5              xtable_1.8-2             scales_0.5.0            
[28] later_0.7.3              proxy_0.4-22             tibble_1.4.2            
[31] lazyeval_0.2.1           magrittr_1.5             mime_0.5                
[34] class_7.3-14             beeswarm_0.2.3           shinydashboard_0.7.0    
[37] tools_3.5.0              data.table_1.11.4        hms_0.4.2               
[40] stringr_1.3.0            googleVis_0.6.2          Rhdf5lib_1.2.1          
[43] munsell_0.5.0            locfit_1.5-9.1           bindrcpp_0.2.2          
[46] compiler_3.5.0           e1071_1.6-8              rlang_0.2.0             
[49] rhdf5_2.24.0             grid_3.5.0               RCurl_1.95-4.10         
[52] tximport_1.8.0           rjson_0.2.20             bitops_1.0-6            
[55] gtable_0.2.0             reshape2_1.4.3           R6_2.2.2                
[58] gridExtra_2.3            dplyr_0.7.5              bindr_0.1.1             
[61] stringi_1.2.2            ggbeeswarm_0.6.0         Rcpp_0.12.16            
[64] tidyselect_0.2.4        

query = 2
2018-11-23 16:31:09.637000
Julia Version 1.0.1
Commit 0d713926f8 (2018-09-29 19:05 UTC)
Platform Info:
  OS: Linux (x86_64-pc-linux-gnu)
  CPU: Intel(R) Xeon(R) Gold 6126 CPU @ 2.60GHz
  WORD_SIZE: 64
  LIBM: libopenlibm
  LLVM: libLLVM-6.0.0 (ORCJIT, skylake)
Environment:
  JULIA_PROJECT = @.

2018-07-20 01:55:28
R version 3.5.0 (2018-04-23)
Platform: x86_64-pc-linux-gnu (64-bit)
Running under: Debian GNU/Linux 9 (stretch)

Matrix products: default
BLAS: /usr/lib/openblas-base/libblas.so.3
LAPACK: /usr/lib/libopenblasp-r0.2.19.so

locale:
 [1] LC_CTYPE=en_US.UTF-8       LC_NUMERIC=C              
 [3] LC_TIME=en_US.UTF-8        LC_COLLATE=en_US.UTF-8    
 [5] LC_MONETARY=en_US.UTF-8    LC_MESSAGES=C             
 [7] LC_PAPER=en_US.UTF-8       LC_NAME=C                 
 [9] LC_ADDRESS=C               LC_TELEPHONE=C            
[11] LC_MEASUREMENT=en_US.UTF-8 LC_IDENTIFICATION=C       

attached base packages:
[1] parallel  stats4    stats     graphics  grDevices utils     datasets 
[8] methods   base     

other attached packages:
 [1] h5_0.9.9                    scmap_1.2.0                
 [3] scater_1.8.0                ggplot2_2.2.1              
 [5] SingleCellExperiment_1.2.0  SummarizedExperiment_1.10.1
 [7] DelayedArray_0.6.1          BiocParallel_1.14.1        
 [9] matrixStats_0.53.1          Biobase_2.40.0             
[11] GenomicRanges_1.32.3        GenomeInfoDb_1.16.0        
[13] IRanges_2.14.10             S4Vectors_0.18.3           
[15] BiocGenerics_0.26.0         readr_1.1.1                

loaded via a namespace (and not attached):
 [1] viridis_0.5.1            edgeR_3.22.2             jsonlite_1.5            
 [4] viridisLite_0.3.0        DelayedMatrixStats_1.2.0 shiny_1.1.0             
 [7] assertthat_0.2.0         GenomeInfoDbData_1.1.0   vipor_0.4.5             
[10] pillar_1.2.2             lattice_0.20-35          glue_1.2.0              
[13] limma_3.36.1             digest_0.6.15            promises_1.0.1          
[16] XVector_0.20.0           randomForest_4.6-14      colorspace_1.3-2        
[19] htmltools_0.3.6          httpuv_1.4.3             Matrix_1.2-14           
[22] plyr_1.8.4               pkgconfig_2.0.1          zlibbioc_1.26.0         
[25] purrr_0.2.5              xtable_1.8-2             scales_0.5.0            
[28] later_0.7.3              proxy_0.4-22             tibble_1.4.2            
[31] lazyeval_0.2.1           magrittr_1.5             mime_0.5                
[34] class_7.3-14             beeswarm_0.2.3           shinydashboard_0.7.0    
[37] tools_3.5.0              data.table_1.11.4        hms_0.4.2               
[40] stringr_1.3.0            googleVis_0.6.2          Rhdf5lib_1.2.1          
[43] munsell_0.5.0            locfit_1.5-9.1           bindrcpp_0.2.2          
[46] compiler_3.5.0           e1071_1.6-8              rlang_0.2.0             
[49] rhdf5_2.24.0             grid_3.5.0               RCurl_1.95-4.10         
[52] tximport_1.8.0           rjson_0.2.20             bitops_1.0-6            
[55] gtable_0.2.0             reshape2_1.4.3           R6_2.2.2                
[58] gridExtra_2.3            dplyr_0.7.5              bindr_0.1.1             
[61] stringi_1.2.2            ggbeeswarm_0.6.0         Rcpp_0.12.16            
[64] tidyselect_0.2.4
```

In [51]:

```
knn_batch_plass2018 = pandas.DataFrame()
knn_batch_plass2018_scmap = pandas.DataFrame()
cluster = clusters["Plass2018"]
for query in ["plan1", "plan2"]:
    print(f"query = {query}")
    # CellScape
    log = toml.load(f"results/Plass2018.knn-batch.batch-{query}.toml")
    print(log["date-time"])
    print(log["version-info"])
    exp = pandas.DataFrame(log["experiment"])
    exp["query"] = query
    exp = pandas.concat(
        [exp,
         pandas.DataFrame([cluster_metrics(x, cluster) for x in exp["filename"]])],
        axis=1)
    knn_batch_plass2018 = knn_batch_plass2018.append(exp, ignore_index=True)
    # scmap
    log = toml.load(f"results/Plass2018.knn-batch.scmap.batch-{query}.toml")
    print(log["date-time"])
    print(log["session-info"])
    exp = pandas.DataFrame(log["experiment"])
    exp["query"] = query
    exp = pandas.concat(
        [exp,
         pandas.DataFrame([cluster_metrics(x, cluster) for x in exp["filename"]])],
        axis=1)
    knn_batch_plass2018_scmap = knn_batch_plass2018_scmap.append(exp, ignore_index=True)
```

```
query = plan1
2018-11-23 20:01:30.353000
Julia Version 1.0.1
Commit 0d713926f8 (2018-09-29 19:05 UTC)
Platform Info:
  OS: Linux (x86_64-pc-linux-gnu)
  CPU: Intel(R) Xeon(R) Gold 6126 CPU @ 2.60GHz
  WORD_SIZE: 64
  LIBM: libopenlibm
  LLVM: libLLVM-6.0.0 (ORCJIT, skylake)
Environment:
  JULIA_PROJECT = @.

2018-07-20 07:07:00
R version 3.5.0 (2018-04-23)
Platform: x86_64-pc-linux-gnu (64-bit)
Running under: Debian GNU/Linux 9 (stretch)

Matrix products: default
BLAS: /usr/lib/openblas-base/libblas.so.3
LAPACK: /usr/lib/libopenblasp-r0.2.19.so

locale:
 [1] LC_CTYPE=en_US.UTF-8       LC_NUMERIC=C              
 [3] LC_TIME=en_US.UTF-8        LC_COLLATE=en_US.UTF-8    
 [5] LC_MONETARY=en_US.UTF-8    LC_MESSAGES=C             
 [7] LC_PAPER=en_US.UTF-8       LC_NAME=C                 
 [9] LC_ADDRESS=C               LC_TELEPHONE=C            
[11] LC_MEASUREMENT=en_US.UTF-8 LC_IDENTIFICATION=C       

attached base packages:
[1] parallel  stats4    stats     graphics  grDevices utils     datasets 
[8] methods   base     

other attached packages:
 [1] h5_0.9.9                    scmap_1.2.0                
 [3] scater_1.8.0                ggplot2_2.2.1              
 [5] SingleCellExperiment_1.2.0  SummarizedExperiment_1.10.1
 [7] DelayedArray_0.6.1          BiocParallel_1.14.1        
 [9] matrixStats_0.53.1          Biobase_2.40.0             
[11] GenomicRanges_1.32.3        GenomeInfoDb_1.16.0        
[13] IRanges_2.14.10             S4Vectors_0.18.3           
[15] BiocGenerics_0.26.0         readr_1.1.1                

loaded via a namespace (and not attached):
 [1] viridis_0.5.1            edgeR_3.22.2             jsonlite_1.5            
 [4] viridisLite_0.3.0        DelayedMatrixStats_1.2.0 shiny_1.1.0             
 [7] assertthat_0.2.0         GenomeInfoDbData_1.1.0   vipor_0.4.5             
[10] pillar_1.2.2             lattice_0.20-35          glue_1.2.0              
[13] limma_3.36.1             digest_0.6.15            promises_1.0.1          
[16] XVector_0.20.0           randomForest_4.6-14      colorspace_1.3-2        
[19] htmltools_0.3.6          httpuv_1.4.3             Matrix_1.2-14           
[22] plyr_1.8.4               pkgconfig_2.0.1          zlibbioc_1.26.0         
[25] purrr_0.2.5              xtable_1.8-2             scales_0.5.0            
[28] later_0.7.3              proxy_0.4-22             tibble_1.4.2            
[31] lazyeval_0.2.1           magrittr_1.5             mime_0.5                
[34] class_7.3-14             beeswarm_0.2.3           shinydashboard_0.7.0    
[37] tools_3.5.0              data.table_1.11.4        hms_0.4.2               
[40] stringr_1.3.0            googleVis_0.6.2          Rhdf5lib_1.2.1          
[43] munsell_0.5.0            locfit_1.5-9.1           bindrcpp_0.2.2          
[46] compiler_3.5.0           e1071_1.6-8              rlang_0.2.0             
[49] rhdf5_2.24.0             grid_3.5.0               RCurl_1.95-4.10         
[52] tximport_1.8.0           rjson_0.2.20             bitops_1.0-6            
[55] gtable_0.2.0             reshape2_1.4.3           R6_2.2.2                
[58] gridExtra_2.3            dplyr_0.7.5              bindr_0.1.1             
[61] stringi_1.2.2            ggbeeswarm_0.6.0         Rcpp_0.12.16            
[64] tidyselect_0.2.4        

query = plan2
2018-11-23 20:00:57.589000
Julia Version 1.0.1
Commit 0d713926f8 (2018-09-29 19:05 UTC)
Platform Info:
  OS: Linux (x86_64-pc-linux-gnu)
  CPU: Intel(R) Xeon(R) Gold 6126 CPU @ 2.60GHz
  WORD_SIZE: 64
  LIBM: libopenlibm
  LLVM: libLLVM-6.0.0 (ORCJIT, skylake)
Environment:
  JULIA_PROJECT = @.

2018-07-20 06:38:58
R version 3.5.0 (2018-04-23)
Platform: x86_64-pc-linux-gnu (64-bit)
Running under: Debian GNU/Linux 9 (stretch)

Matrix products: default
BLAS: /usr/lib/openblas-base/libblas.so.3
LAPACK: /usr/lib/libopenblasp-r0.2.19.so

locale:
 [1] LC_CTYPE=en_US.UTF-8       LC_NUMERIC=C              
 [3] LC_TIME=en_US.UTF-8        LC_COLLATE=en_US.UTF-8    
 [5] LC_MONETARY=en_US.UTF-8    LC_MESSAGES=C             
 [7] LC_PAPER=en_US.UTF-8       LC_NAME=C                 
 [9] LC_ADDRESS=C               LC_TELEPHONE=C            
[11] LC_MEASUREMENT=en_US.UTF-8 LC_IDENTIFICATION=C       

attached base packages:
[1] parallel  stats4    stats     graphics  grDevices utils     datasets 
[8] methods   base     

other attached packages:
 [1] h5_0.9.9                    scmap_1.2.0                
 [3] scater_1.8.0                ggplot2_2.2.1              
 [5] SingleCellExperiment_1.2.0  SummarizedExperiment_1.10.1
 [7] DelayedArray_0.6.1          BiocParallel_1.14.1        
 [9] matrixStats_0.53.1          Biobase_2.40.0             
[11] GenomicRanges_1.32.3        GenomeInfoDb_1.16.0        
[13] IRanges_2.14.10             S4Vectors_0.18.3           
[15] BiocGenerics_0.26.0         readr_1.1.1                

loaded via a namespace (and not attached):
 [1] viridis_0.5.1            edgeR_3.22.2             jsonlite_1.5            
 [4] viridisLite_0.3.0        DelayedMatrixStats_1.2.0 shiny_1.1.0             
 [7] assertthat_0.2.0         GenomeInfoDbData_1.1.0   vipor_0.4.5             
[10] pillar_1.2.2             lattice_0.20-35          glue_1.2.0              
[13] limma_3.36.1             digest_0.6.15            promises_1.0.1          
[16] XVector_0.20.0           randomForest_4.6-14      colorspace_1.3-2        
[19] htmltools_0.3.6          httpuv_1.4.3             Matrix_1.2-14           
[22] plyr_1.8.4               pkgconfig_2.0.1          zlibbioc_1.26.0         
[25] purrr_0.2.5              xtable_1.8-2             scales_0.5.0            
[28] later_0.7.3              proxy_0.4-22             tibble_1.4.2            
[31] lazyeval_0.2.1           magrittr_1.5             mime_0.5                
[34] class_7.3-14             beeswarm_0.2.3           shinydashboard_0.7.0    
[37] tools_3.5.0              data.table_1.11.4        hms_0.4.2               
[40] stringr_1.3.0            googleVis_0.6.2          Rhdf5lib_1.2.1          
[43] munsell_0.5.0            locfit_1.5-9.1           bindrcpp_0.2.2          
[46] compiler_3.5.0           e1071_1.6-8              rlang_0.2.0             
[49] rhdf5_2.24.0             grid_3.5.0               RCurl_1.95-4.10         
[52] tximport_1.8.0           rjson_0.2.20             bitops_1.0-6            
[55] gtable_0.2.0             reshape2_1.4.3           R6_2.2.2                
[58] gridExtra_2.3            dplyr_0.7.5              bindr_0.1.1             
[61] stringi_1.2.2            ggbeeswarm_0.6.0         Rcpp_0.12.16            
[64] tidyselect_0.2.4
```

In [52]:

```
knn_batch_shekhar2016 = knn_batch_shekhar2016[["query", "n-bits", "n-lshashes", "transformer", "filename", "consistency", "cohen_kappa"]]
knn_batch_shekhar2016_scmap = knn_batch_shekhar2016_scmap[["query", "n-features", "n-centroids-factor", "filename", "consistency", "cohen_kappa"]]
```

In [53]:

```
knn_batch_plass2018 = knn_batch_plass2018[["query", "n-bits", "n-lshashes", "transformer", "filename", "consistency", "cohen_kappa"]]
knn_batch_plass2018_scmap = knn_batch_plass2018_scmap[["query", "n-features", "n-centroids-factor", "filename", "consistency", "cohen_kappa"]]
```

In [54]:

```
knn_batch_shekhar2016.groupby(["query", "n-bits", "n-lshashes", "transformer"]).describe()
```

Out[54]:

|  |  |  |  | cohen\_kappa | | | | | | | | consistency | | | | | | | |
| --- | --- | --- | --- | --- | --- | --- | --- | --- | --- | --- | --- | --- | --- | --- | --- | --- | --- | --- | --- |
|  |  |  |  | count | mean | std | min | 25% | 50% | 75% | max | count | mean | std | min | 25% | 50% | 75% | max |
| query | n-bits | n-lshashes | transformer |  |  |  |  |  |  |  |  |  |  |  |  |  |  |  |  |
| 1 | 128 | 1 | ftt | 5.0 | 0.967343 | 0.001597 | 0.965431 | 0.966473 | 0.966835 | 0.968746 | 0.969229 | 5.0 | 0.976550 | 0.001144 | 0.975191 | 0.975906 | 0.976192 | 0.977551 | 0.977908 |
|  |  |  | log1p | 5.0 | 0.961930 | 0.001039 | 0.960866 | 0.961100 | 0.961950 | 0.962256 | 0.963477 | 5.0 | 0.972660 | 0.000746 | 0.971902 | 0.972045 | 0.972689 | 0.972903 | 0.973761 |
|  |  | 4 | ftt | 5.0 | 0.974257 | 0.001073 | 0.973026 | 0.973521 | 0.974010 | 0.975216 | 0.975510 | 5.0 | 0.981511 | 0.000771 | 0.980625 | 0.980982 | 0.981340 | 0.982198 | 0.982412 |
|  |  |  | log1p | 5.0 | 0.971488 | 0.000911 | 0.970327 | 0.970832 | 0.971729 | 0.971933 | 0.972620 | 5.0 | 0.979524 | 0.000653 | 0.978695 | 0.979052 | 0.979695 | 0.979838 | 0.980339 |
|  |  | 16 | ftt | 5.0 | 0.977567 | 0.000510 | 0.976908 | 0.977214 | 0.977606 | 0.978002 | 0.978105 | 5.0 | 0.983885 | 0.000367 | 0.983413 | 0.983628 | 0.983914 | 0.984200 | 0.984271 |
|  |  |  | log1p | 5.0 | 0.973762 | 0.000964 | 0.972727 | 0.973030 | 0.973523 | 0.974611 | 0.974917 | 5.0 | 0.981154 | 0.000694 | 0.980410 | 0.980625 | 0.980982 | 0.981769 | 0.981983 |
|  | 256 | 1 | ftt | 5.0 | 0.972639 | 0.000692 | 0.971533 | 0.972413 | 0.972913 | 0.973113 | 0.973224 | 5.0 | 0.980353 | 0.000499 | 0.979552 | 0.980196 | 0.980553 | 0.980696 | 0.980768 |
|  |  |  | log1p | 5.0 | 0.969441 | 0.001844 | 0.966959 | 0.969035 | 0.969444 | 0.969639 | 0.972130 | 5.0 | 0.978051 | 0.001325 | 0.976264 | 0.977765 | 0.978051 | 0.978194 | 0.979981 |
|  |  | 4 | ftt | 5.0 | 0.977088 | 0.001253 | 0.974916 | 0.977096 | 0.977807 | 0.977808 | 0.977814 | 5.0 | 0.983542 | 0.000898 | 0.981983 | 0.983556 | 0.984057 | 0.984057 | 0.984057 |
|  |  |  | log1p | 5.0 | 0.974006 | 0.000960 | 0.972938 | 0.973030 | 0.974330 | 0.974815 | 0.974918 | 5.0 | 0.981326 | 0.000692 | 0.980553 | 0.980625 | 0.981554 | 0.981912 | 0.981983 |
|  |  | 16 | ftt | 5.0 | 0.977688 | 0.000896 | 0.976409 | 0.977309 | 0.977809 | 0.978110 | 0.978801 | 5.0 | 0.983971 | 0.000642 | 0.983056 | 0.983699 | 0.984057 | 0.984271 | 0.984772 |
|  |  |  | log1p | 5.0 | 0.975598 | 0.001578 | 0.973925 | 0.974320 | 0.975421 | 0.976616 | 0.977710 | 5.0 | 0.982469 | 0.001132 | 0.981268 | 0.981554 | 0.982341 | 0.983199 | 0.983985 |
| 2 | 128 | 1 | ftt | 5.0 | 0.917644 | 0.003515 | 0.913387 | 0.915785 | 0.917754 | 0.918432 | 0.922860 | 5.0 | 0.928123 | 0.003048 | 0.924438 | 0.926510 | 0.928212 | 0.928804 | 0.932652 |
|  |  |  | log1p | 5.0 | 0.913144 | 0.001221 | 0.912232 | 0.912589 | 0.912633 | 0.912991 | 0.915275 | 5.0 | 0.924201 | 0.001069 | 0.923401 | 0.923697 | 0.923771 | 0.924067 | 0.926066 |
|  |  | 4 | ftt | 5.0 | 0.933430 | 0.003395 | 0.929295 | 0.931085 | 0.933130 | 0.936350 | 0.937288 | 5.0 | 0.941874 | 0.002956 | 0.938277 | 0.939831 | 0.941607 | 0.944420 | 0.945234 |
|  |  |  | log1p | 5.0 | 0.928546 | 0.002495 | 0.924890 | 0.928201 | 0.928559 | 0.929227 | 0.931856 | 5.0 | 0.937611 | 0.002174 | 0.934429 | 0.937315 | 0.937611 | 0.938203 | 0.940497 |
|  |  | 16 | ftt | 5.0 | 0.937546 | 0.001687 | 0.935498 | 0.936791 | 0.937460 | 0.937885 | 0.940094 | 5.0 | 0.945456 | 0.001468 | 0.943680 | 0.944790 | 0.945382 | 0.945752 | 0.947676 |
|  |  |  | log1p | 5.0 | 0.934173 | 0.001788 | 0.932465 | 0.933553 | 0.933738 | 0.933898 | 0.937210 | 5.0 | 0.942510 | 0.001561 | 0.941015 | 0.941978 | 0.942126 | 0.942274 | 0.945160 |
|  | 256 | 1 | ftt | 5.0 | 0.928776 | 0.002921 | 0.923697 | 0.928956 | 0.929980 | 0.930575 | 0.930674 | 5.0 | 0.937818 | 0.002544 | 0.933393 | 0.937981 | 0.938869 | 0.939387 | 0.939461 |
|  |  |  | log1p | 5.0 | 0.922948 | 0.002586 | 0.920292 | 0.921486 | 0.922256 | 0.923691 | 0.927013 | 5.0 | 0.932741 | 0.002251 | 0.930432 | 0.931468 | 0.932134 | 0.933393 | 0.936279 |
|  |  | 4 | ftt | 5.0 | 0.935409 | 0.001970 | 0.933051 | 0.934573 | 0.934659 | 0.936700 | 0.938061 | 5.0 | 0.943591 | 0.001716 | 0.941533 | 0.942866 | 0.942940 | 0.944716 | 0.945900 |
|  |  |  | log1p | 5.0 | 0.931859 | 0.001522 | 0.930087 | 0.930417 | 0.932369 | 0.932960 | 0.933463 | 5.0 | 0.940497 | 0.001332 | 0.938943 | 0.939239 | 0.940941 | 0.941459 | 0.941903 |
|  |  | 16 | ftt | 5.0 | 0.938063 | 0.000659 | 0.937300 | 0.937551 | 0.938072 | 0.938482 | 0.938910 | 5.0 | 0.945900 | 0.000576 | 0.945234 | 0.945456 | 0.945900 | 0.946270 | 0.946640 |
|  |  |  | log1p | 5.0 | 0.933663 | 0.002473 | 0.931871 | 0.932285 | 0.932804 | 0.933386 | 0.937968 | 5.0 | 0.942066 | 0.002160 | 0.940497 | 0.940867 | 0.941311 | 0.941829 | 0.945826 |

In [55]:

```
knn_batch_shekhar2016_scmap.groupby(["query", "n-features", "n-centroids-factor"]).describe()
```

Out[55]:

|  |  |  | cohen\_kappa | | | | | | | | consistency | | | | | | | |
| --- | --- | --- | --- | --- | --- | --- | --- | --- | --- | --- | --- | --- | --- | --- | --- | --- | --- | --- |
|  |  |  | count | mean | std | min | 25% | 50% | 75% | max | count | mean | std | min | 25% | 50% | 75% | max |
| query | n-features | n-centroids-factor |  |  |  |  |  |  |  |  |  |  |  |  |  |  |  |  |
| 1 | 500 | 1 | 5.0 | 0.906302 | 0.003045 | 0.904100 | 0.904668 | 0.905469 | 0.905639 | 0.911634 | 5.0 | 0.933195 | 0.002178 | 0.931651 | 0.932008 | 0.932580 | 0.932723 | 0.937013 |
|  |  | 4 | 5.0 | 0.902334 | 0.004108 | 0.895238 | 0.903309 | 0.903560 | 0.903618 | 0.905946 | 5.0 | 0.930235 | 0.002988 | 0.925073 | 0.931007 | 0.931079 | 0.931150 | 0.932866 |
|  | 1000 | 1 | 5.0 | 0.889980 | 0.001959 | 0.887270 | 0.888738 | 0.890646 | 0.891067 | 0.892178 | 5.0 | 0.921541 | 0.001427 | 0.919568 | 0.920641 | 0.921999 | 0.922356 | 0.923143 |
|  |  | 4 | 5.0 | 0.871561 | 0.006414 | 0.864892 | 0.865593 | 0.871590 | 0.876290 | 0.879441 | 5.0 | 0.908243 | 0.004567 | 0.903482 | 0.903982 | 0.908272 | 0.911704 | 0.913777 |
|  | 2000 | 1 | 5.0 | 0.878273 | 0.006497 | 0.867889 | 0.877997 | 0.878764 | 0.881234 | 0.885480 | 5.0 | 0.913105 | 0.004702 | 0.905627 | 0.912776 | 0.913491 | 0.915278 | 0.918353 |
|  |  | 4 | 5.0 | 0.846589 | 0.005323 | 0.839447 | 0.843484 | 0.846639 | 0.851092 | 0.852284 | 5.0 | 0.890198 | 0.003896 | 0.884822 | 0.888039 | 0.890398 | 0.893544 | 0.894187 |
|  | 4000 | 1 | 5.0 | 0.873499 | 0.004293 | 0.866051 | 0.874054 | 0.874996 | 0.875438 | 0.876955 | 5.0 | 0.909702 | 0.003161 | 0.904197 | 0.910202 | 0.910846 | 0.911060 | 0.912204 |
|  |  | 4 | 5.0 | 0.834190 | 0.004355 | 0.828442 | 0.832742 | 0.833356 | 0.836229 | 0.840180 | 5.0 | 0.881061 | 0.003437 | 0.876457 | 0.880174 | 0.880246 | 0.882677 | 0.885751 |
| 2 | 500 | 1 | 5.0 | 0.788696 | 0.005047 | 0.782134 | 0.785335 | 0.788975 | 0.792834 | 0.794202 | 5.0 | 0.815882 | 0.004419 | 0.810169 | 0.812833 | 0.816237 | 0.819494 | 0.820678 |
|  |  | 4 | 5.0 | 0.702258 | 0.009858 | 0.687276 | 0.700306 | 0.703877 | 0.705414 | 0.714418 | 5.0 | 0.739624 | 0.008704 | 0.726391 | 0.737937 | 0.741045 | 0.742377 | 0.750370 |
|  | 1000 | 1 | 5.0 | 0.720922 | 0.011637 | 0.709187 | 0.713184 | 0.718718 | 0.724537 | 0.738982 | 5.0 | 0.755817 | 0.010399 | 0.745337 | 0.748964 | 0.753774 | 0.759029 | 0.771980 |
|  |  | 4 | 5.0 | 0.550824 | 0.010224 | 0.537429 | 0.543543 | 0.553679 | 0.556677 | 0.562793 | 5.0 | 0.603390 | 0.009375 | 0.590956 | 0.596951 | 0.605906 | 0.608792 | 0.614343 |
|  | 2000 | 1 | 5.0 | 0.730028 | 0.016452 | 0.703494 | 0.729786 | 0.731799 | 0.736931 | 0.748131 | 5.0 | 0.764032 | 0.014531 | 0.740601 | 0.763766 | 0.765690 | 0.770056 | 0.780047 |
|  |  | 4 | 5.0 | 0.527771 | 0.016518 | 0.511607 | 0.512860 | 0.527063 | 0.536518 | 0.550810 | 5.0 | 0.582371 | 0.015332 | 0.567274 | 0.568532 | 0.581853 | 0.590512 | 0.603686 |
|  | 4000 | 1 | 5.0 | 0.699650 | 0.011761 | 0.680889 | 0.695128 | 0.707004 | 0.707600 | 0.707626 | 5.0 | 0.736871 | 0.010450 | 0.720249 | 0.732756 | 0.743413 | 0.743931 | 0.744005 |
|  |  | 4 | 5.0 | 0.562187 | 0.012952 | 0.546283 | 0.555290 | 0.560579 | 0.568473 | 0.580311 | 5.0 | 0.613766 | 0.011582 | 0.599467 | 0.607534 | 0.612419 | 0.619597 | 0.629811 |

In [56]:

```
knn_batch_plass2018.groupby(["query", "n-bits", "n-lshashes", "transformer"]).describe()
```

Out[56]:

|  |  |  |  | cohen\_kappa | | | | | | | | consistency | | | | | | | |
| --- | --- | --- | --- | --- | --- | --- | --- | --- | --- | --- | --- | --- | --- | --- | --- | --- | --- | --- | --- |
|  |  |  |  | count | mean | std | min | 25% | 50% | 75% | max | count | mean | std | min | 25% | 50% | 75% | max |
| query | n-bits | n-lshashes | transformer |  |  |  |  |  |  |  |  |  |  |  |  |  |  |  |  |
| plan1 | 128 | 1 | ftt | 5.0 | 0.631777 | 0.004916 | 0.624646 | 0.629288 | 0.632919 | 0.634929 | 0.637105 | 5.0 | 0.677441 | 0.004812 | 0.670097 | 0.675661 | 0.678720 | 0.680111 | 0.682615 |
|  |  |  | log1p | 5.0 | 0.643424 | 0.002561 | 0.639112 | 0.643465 | 0.643889 | 0.645029 | 0.645626 | 5.0 | 0.688122 | 0.002229 | 0.684284 | 0.688178 | 0.689013 | 0.689291 | 0.689847 |
|  |  | 4 | ftt | 5.0 | 0.650583 | 0.004607 | 0.643232 | 0.649433 | 0.651804 | 0.653352 | 0.655095 | 5.0 | 0.693908 | 0.004423 | 0.687065 | 0.692629 | 0.694854 | 0.696245 | 0.698748 |
|  |  |  | log1p | 5.0 | 0.656917 | 0.001169 | 0.655381 | 0.656619 | 0.656881 | 0.657055 | 0.658649 | 5.0 | 0.700083 | 0.001084 | 0.699026 | 0.699305 | 0.699583 | 0.701252 | 0.701252 |
|  |  | 16 | ftt | 5.0 | 0.658275 | 0.003980 | 0.653278 | 0.656200 | 0.657694 | 0.660609 | 0.663594 | 5.0 | 0.700695 | 0.003600 | 0.695688 | 0.699305 | 0.700417 | 0.702921 | 0.705146 |
|  |  |  | log1p | 5.0 | 0.666558 | 0.002022 | 0.663337 | 0.665827 | 0.667485 | 0.668038 | 0.668105 | 5.0 | 0.708317 | 0.001453 | 0.705981 | 0.707928 | 0.708762 | 0.709318 | 0.709597 |
|  | 256 | 1 | ftt | 5.0 | 0.646580 | 0.008267 | 0.637798 | 0.639629 | 0.646903 | 0.650475 | 0.658097 | 5.0 | 0.690125 | 0.007676 | 0.682058 | 0.684006 | 0.690125 | 0.693185 | 0.701252 |
|  |  |  | log1p | 5.0 | 0.657549 | 0.003121 | 0.653767 | 0.655284 | 0.657987 | 0.659009 | 0.661698 | 5.0 | 0.700640 | 0.003537 | 0.696523 | 0.697636 | 0.701530 | 0.702364 | 0.705146 |
|  |  | 4 | ftt | 5.0 | 0.657944 | 0.003259 | 0.653933 | 0.655524 | 0.658035 | 0.660655 | 0.661575 | 5.0 | 0.700473 | 0.003011 | 0.696801 | 0.698470 | 0.700139 | 0.702921 | 0.704033 |
|  |  |  | log1p | 5.0 | 0.661615 | 0.006855 | 0.656504 | 0.657665 | 0.659449 | 0.660960 | 0.673496 | 5.0 | 0.704145 | 0.006472 | 0.699026 | 0.701530 | 0.701808 | 0.702921 | 0.715438 |
|  |  | 16 | ftt | 5.0 | 0.660677 | 0.002823 | 0.657563 | 0.657866 | 0.661344 | 0.663222 | 0.663388 | 5.0 | 0.702754 | 0.002373 | 0.700139 | 0.700695 | 0.702643 | 0.705146 | 0.705146 |
|  |  |  | log1p | 5.0 | 0.668463 | 0.004182 | 0.662497 | 0.666139 | 0.669222 | 0.672158 | 0.672301 | 5.0 | 0.709875 | 0.003551 | 0.705146 | 0.707928 | 0.709597 | 0.713213 | 0.713491 |
| plan2 | 128 | 1 | ftt | 5.0 | 0.671158 | 0.003761 | 0.665638 | 0.668844 | 0.673457 | 0.673639 | 0.674211 | 5.0 | 0.707827 | 0.003582 | 0.702538 | 0.705711 | 0.709677 | 0.710471 | 0.710735 |
|  |  |  | log1p | 5.0 | 0.672607 | 0.006777 | 0.661987 | 0.670130 | 0.675608 | 0.676105 | 0.679206 | 5.0 | 0.708673 | 0.006042 | 0.699101 | 0.706504 | 0.711793 | 0.711793 | 0.714172 |
|  |  | 4 | ftt | 5.0 | 0.689283 | 0.005419 | 0.682900 | 0.686182 | 0.688023 | 0.692677 | 0.696631 | 5.0 | 0.723903 | 0.005056 | 0.717874 | 0.721047 | 0.722898 | 0.726864 | 0.730830 |
|  |  |  | log1p | 5.0 | 0.698393 | 0.004403 | 0.693752 | 0.695955 | 0.696346 | 0.701497 | 0.704413 | 5.0 | 0.731676 | 0.004157 | 0.727393 | 0.729508 | 0.729508 | 0.734532 | 0.737441 |
|  |  | 16 | ftt | 5.0 | 0.697151 | 0.004885 | 0.690891 | 0.693706 | 0.697629 | 0.700871 | 0.702657 | 5.0 | 0.730989 | 0.004521 | 0.725013 | 0.727922 | 0.731623 | 0.734532 | 0.735854 |
|  |  |  | log1p | 5.0 | 0.704023 | 0.003164 | 0.700928 | 0.701194 | 0.703382 | 0.706794 | 0.707816 | 5.0 | 0.736542 | 0.003041 | 0.733474 | 0.734003 | 0.735854 | 0.739027 | 0.740349 |
|  | 256 | 1 | ftt | 5.0 | 0.680336 | 0.006121 | 0.673749 | 0.674068 | 0.681954 | 0.685117 | 0.686793 | 5.0 | 0.715865 | 0.005580 | 0.709942 | 0.710206 | 0.717081 | 0.719989 | 0.722105 |
|  |  |  | log1p | 5.0 | 0.685099 | 0.007544 | 0.676830 | 0.680510 | 0.682066 | 0.691875 | 0.694215 | 5.0 | 0.719619 | 0.006894 | 0.712057 | 0.715230 | 0.717081 | 0.725806 | 0.727922 |
|  |  | 4 | ftt | 5.0 | 0.693821 | 0.004347 | 0.688234 | 0.691417 | 0.693535 | 0.696500 | 0.699418 | 5.0 | 0.728027 | 0.003857 | 0.722898 | 0.726071 | 0.727922 | 0.730301 | 0.732946 |
|  |  |  | log1p | 5.0 | 0.700533 | 0.007146 | 0.688950 | 0.698548 | 0.703299 | 0.705628 | 0.706239 | 5.0 | 0.733421 | 0.006253 | 0.723162 | 0.731888 | 0.736118 | 0.737705 | 0.738234 |
|  |  | 16 | ftt | 5.0 | 0.698753 | 0.003741 | 0.695068 | 0.696387 | 0.698315 | 0.699210 | 0.704786 | 5.0 | 0.732417 | 0.003582 | 0.728715 | 0.730566 | 0.731888 | 0.732681 | 0.738234 |
|  |  |  | log1p | 5.0 | 0.707578 | 0.004233 | 0.702634 | 0.706240 | 0.706259 | 0.708653 | 0.714101 | 5.0 | 0.739714 | 0.003816 | 0.735325 | 0.738234 | 0.738763 | 0.740613 | 0.745637 |

In [57]:

```
knn_batch_plass2018_scmap.groupby(["query", "n-features", "n-centroids-factor"]).describe()
```

Out[57]:

|  |  |  | cohen\_kappa | | | | | | | | consistency | | | | | | | |
| --- | --- | --- | --- | --- | --- | --- | --- | --- | --- | --- | --- | --- | --- | --- | --- | --- | --- | --- |
|  |  |  | count | mean | std | min | 25% | 50% | 75% | max | count | mean | std | min | 25% | 50% | 75% | max |
| query | n-features | n-centroids-factor |  |  |  |  |  |  |  |  |  |  |  |  |  |  |  |  |
| plan1 | 500 | 1 | 5.0 | 0.564847 | 0.003940 | 0.559935 | 0.562857 | 0.563818 | 0.567943 | 0.569681 | 5.0 | 0.629930 | 0.003533 | 0.625591 | 0.627816 | 0.629764 | 0.631711 | 0.634771 |
|  |  | 4 | 5.0 | 0.571527 | 0.000749 | 0.570600 | 0.570901 | 0.571808 | 0.571937 | 0.572389 | 5.0 | 0.635828 | 0.000457 | 0.635327 | 0.635605 | 0.635605 | 0.636161 | 0.636439 |
|  | 1000 | 1 | 5.0 | 0.545385 | 0.001743 | 0.542856 | 0.545098 | 0.545123 | 0.546266 | 0.547582 | 5.0 | 0.610125 | 0.002222 | 0.606954 | 0.609458 | 0.610292 | 0.610848 | 0.613074 |
|  |  | 4 | 5.0 | 0.547384 | 0.003382 | 0.545366 | 0.545765 | 0.546113 | 0.546272 | 0.553402 | 5.0 | 0.611683 | 0.002957 | 0.610292 | 0.610292 | 0.610292 | 0.610570 | 0.616968 |
|  | 2000 | 1 | 5.0 | 0.561276 | 0.004049 | 0.558606 | 0.559457 | 0.559592 | 0.560283 | 0.568441 | 5.0 | 0.626092 | 0.003323 | 0.623922 | 0.624757 | 0.624757 | 0.625035 | 0.631989 |
|  |  | 4 | 5.0 | 0.563815 | 0.003546 | 0.559003 | 0.561223 | 0.565395 | 0.566034 | 0.567421 | 5.0 | 0.629040 | 0.003045 | 0.624757 | 0.626982 | 0.630598 | 0.630876 | 0.631989 |
|  | 4000 | 1 | 5.0 | 0.556327 | 0.003973 | 0.552442 | 0.553276 | 0.556222 | 0.557207 | 0.562488 | 5.0 | 0.621864 | 0.003797 | 0.618081 | 0.618915 | 0.621419 | 0.623366 | 0.627538 |
|  |  | 4 | 5.0 | 0.557978 | 0.002785 | 0.553798 | 0.557435 | 0.557882 | 0.559484 | 0.561290 | 5.0 | 0.623978 | 0.002339 | 0.620584 | 0.623644 | 0.623644 | 0.625035 | 0.626982 |
| plan2 | 500 | 1 | 5.0 | 0.613393 | 0.002563 | 0.609135 | 0.613764 | 0.613859 | 0.614121 | 0.616088 | 5.0 | 0.663247 | 0.002087 | 0.659968 | 0.663406 | 0.663406 | 0.663670 | 0.665785 |
|  |  | 4 | 5.0 | 0.620226 | 0.001119 | 0.618497 | 0.620159 | 0.620416 | 0.620441 | 0.621615 | 5.0 | 0.669276 | 0.001081 | 0.667636 | 0.668958 | 0.669487 | 0.669751 | 0.670545 |
|  | 1000 | 1 | 5.0 | 0.595551 | 0.003354 | 0.590917 | 0.594744 | 0.594882 | 0.597239 | 0.599975 | 5.0 | 0.647541 | 0.003084 | 0.643046 | 0.646748 | 0.647277 | 0.649392 | 0.651243 |
|  |  | 4 | 5.0 | 0.601025 | 0.004850 | 0.595862 | 0.597494 | 0.601572 | 0.601818 | 0.608377 | 5.0 | 0.651031 | 0.004446 | 0.646219 | 0.648070 | 0.651243 | 0.651772 | 0.657853 |
|  | 2000 | 1 | 5.0 | 0.615866 | 0.003010 | 0.612617 | 0.613119 | 0.615845 | 0.618734 | 0.619015 | 5.0 | 0.665415 | 0.002442 | 0.662348 | 0.663406 | 0.666050 | 0.667372 | 0.667901 |
|  |  | 4 | 5.0 | 0.616300 | 0.001055 | 0.615036 | 0.615853 | 0.616270 | 0.616416 | 0.617925 | 5.0 | 0.665838 | 0.000961 | 0.664728 | 0.665521 | 0.665785 | 0.665785 | 0.667372 |
|  | 4000 | 1 | 5.0 | 0.616679 | 0.004683 | 0.610578 | 0.614331 | 0.616111 | 0.619766 | 0.622607 | 5.0 | 0.665151 | 0.004474 | 0.659175 | 0.663141 | 0.664463 | 0.668429 | 0.670545 |
|  |  | 4 | 5.0 | 0.619582 | 0.005265 | 0.611036 | 0.618354 | 0.621143 | 0.623409 | 0.623970 | 5.0 | 0.668112 | 0.004615 | 0.660762 | 0.666843 | 0.669223 | 0.671602 | 0.672131 |

In [58]:

```
n_bits = 128
n_lshashes = 4
transformer = "log1p"
n_features = 500
n_centroids_factor = 1
for metric in ["consistency", "cohen_kappa"]:
    fig, axes = subplots(2, 2, dpi=dpi, figsize=(8, 6))
    for i, (dataset, data, data_scmap) in enumerate([
            ("Shekhar2016", knn_batch_shekhar2016, knn_batch_shekhar2016_scmap),
            ("Plass2018", knn_batch_plass2018, knn_batch_plass2018_scmap)]):
        if i > 0:
            axes[i,0].get_shared_x_axes().join(axes[0,0], axes[i,0])
            axes[i,1].get_shared_x_axes().join(axes[0,1], axes[i,1])
        axes[i,0].get_shared_y_axes().join(axes[i,0], axes[i,1])
        data = data[(data["n-bits"]==n_bits)&(data["n-lshashes"]==n_lshashes)&(data["transformer"]==transformer)]
        mean = knn_cv[(knn_cv["dataset"]==dataset)&(knn_cv["n-bits"]==n_bits)&(knn_cv["n-lshashes"]==n_lshashes)&(knn_cv["transformer"]==transformer)][metric].mean()
        data_scmap = data_scmap[(data_scmap["n-features"]==n_features)&(data_scmap["n-centroids-factor"]==n_centroids_factor)]
        mean_scmap = knn_cv_scmap[(knn_cv_scmap["dataset"]==dataset)&(knn_cv_scmap["n-features"]==n_features)&(knn_cv_scmap["n-centroids-factor"]==n_centroids_factor)][metric].mean()
        ymin = min(data[metric].min(), data_scmap[metric].min())
        ymax = max(data[metric].max(), data_scmap[metric].max())
        ydiff = ymax - ymin
        axes[i,0].set_ylim(ymin - ydiff * 0.02, ymax + ydiff * 0.02)  # need to set ylim before plotting
        seaborn.swarmplot(x="query", y=metric, data=data, ax=axes[i,0])
        axes[i,0].axhline(mean, color="orangered", lw=0.5)
        seaborn.swarmplot(x="query", y=metric, data=data_scmap, ax=axes[i,1])
        axes[i,1].axhline(mean_scmap, color="orangered", lw=0.5)
        axes[i,0].set_ylabel(dataset + "\n\n" + metric_labels[metric])
        axes[i,1].set_ylabel("")
        axes[i,1].set_yticklabels([])
        for j in range(2):
            axes[i,j].locator_params(axis="y", nbins=4)
            axes[i,j].grid(axis="y", alpha=0.3)
    axes[0,0].set_title("CellFishing")
    axes[0,1].set_title("scmap-cell")
    seaborn.despine(fig=fig)
    fig.tight_layout()
    savefig(f"mapping-{metric}")
```

In [59]:

```
batch1 = pandas.read_table("data/Shekhar2016.batch-1.txt", names=["cell"])["cell"]
batch2 = pandas.read_table("data/Shekhar2016.batch-2.txt", names=["cell"])["cell"]
cluster = clusters["Shekhar2016"]
fig, ax = subplots(dpi=dpi, figsize=(8, 8))
seaborn.countplot(y="cluster", hue="batch", data=pandas.concat([
    pandas.DataFrame(cluster[batch1]).assign(batch=1),
    pandas.DataFrame(cluster[batch2]).assign(batch=2),
]), order=cluster.value_counts().index.values, log=False, ax=ax)
ax.set_xlabel("cluster size")
seaborn.despine(fig=fig)
fig.tight_layout()
savefig(f"Shekhar2016-clusters-batches")
```

In [60]:

```
batch1 = pandas.read_table("data/Plass2018.batch-plan1.txt", names=["cell"])["cell"]
batch2 = pandas.read_table("data/Plass2018.batch-plan2.txt", names=["cell"])["cell"]
cluster = clusters["Plass2018"]
fig, ax = subplots(dpi=dpi, figsize=(8, 8))
seaborn.countplot(y="cluster", hue="batch", data=pandas.concat([
    pandas.DataFrame(cluster[batch1]).assign(batch="plan1"),
    pandas.DataFrame(cluster[batch2]).assign(batch="plan2"),
]), order=cluster.value_counts().index.values, log=False, ax=ax)
ax.set_xlabel("cluster size")
seaborn.despine(fig=fig)
fig.tight_layout()
savefig(f"Plass2018-clusters-batches")
```

#### Mapping across species¶

In [61]:

```
knn_species = toml.load("results/Baron2016.knn-species.toml")
print(knn_species["date-time"])
print(knn_species["version-info"])
knn_species = pandas.DataFrame(knn_species["experiment"])
```

```
2018-11-23 21:02:23.979000
Julia Version 1.0.1
Commit 0d713926f8 (2018-09-29 19:05 UTC)
Platform Info:
  OS: Linux (x86_64-pc-linux-gnu)
  CPU: Intel(R) Xeon(R) Gold 6126 CPU @ 2.60GHz
  WORD_SIZE: 64
  LIBM: libopenlibm
  LLVM: libLLVM-6.0.0 (ORCJIT, skylake)
Environment:
  JULIA_PROJECT = @.
```

In [62]:

```
items = []
for row in knn_species.itertuples():
    if row.query == "mouse":
        query = "Baron2016-mouse"
        reference = "Baron2016-human"
    else:
        query = "Baron2016-human"
        reference = "Baron2016-mouse"
    items.append(cluster_metrics(row.filename, clusters[query], cluster_nn=clusters[reference]))
knn_species = knn_species.join(pandas.DataFrame(items))
knn_species
```

Out[62]:

|  | filename | index-time | infer-stats | n-cells-database | n-cells-query | n-nns | preproc-time | query | query-time | rank-time | search-time | adjusted\_rand | cohen\_kappa | consistency |
| --- | --- | --- | --- | --- | --- | --- | --- | --- | --- | --- | --- | --- | --- | --- |
| 0 | results/Baron2016.knn-species/n\_nns-10.query-m... | 2.962004 | query | 8569 | 1886 | 10 | 110401684 | mouse | 1.013063 | 108799558 | 526099945 | 0.607250 | 0.706694 | 0.780488 |
| 1 | results/Baron2016.knn-species/n\_nns-10.query-m... | 0.779242 | query | 8569 | 1886 | 10 | 35827374 | mouse | 0.417084 | 3975116 | 374951135 | 0.643186 | 0.725126 | 0.795864 |
| 2 | results/Baron2016.knn-species/n\_nns-10.query-m... | 0.770178 | query | 8569 | 1886 | 10 | 35457902 | mouse | 0.394713 | 3934688 | 353003249 | 0.633194 | 0.716453 | 0.787381 |
| 3 | results/Baron2016.knn-species/n\_nns-10.query-m... | 0.769975 | query | 8569 | 1886 | 10 | 34882312 | mouse | 0.414550 | 3930024 | 373479224 | 0.543707 | 0.659480 | 0.742312 |
| 4 | results/Baron2016.knn-species/n\_nns-10.query-m... | 0.770069 | query | 8569 | 1886 | 10 | 34709885 | mouse | 0.414439 | 3957316 | 373495476 | 0.622179 | 0.714670 | 0.787381 |
| 5 | results/Baron2016.knn-species/n\_nns-10.query-m... | 0.770058 | database | 8569 | 1886 | 10 | 33223131 | mouse | 0.313570 | 3682028 | 274421359 | 0.453689 | 0.392828 | 0.487805 |
| 6 | results/Baron2016.knn-species/n\_nns-10.query-m... | 0.769583 | database | 8569 | 1886 | 10 | 33309111 | mouse | 0.352209 | 3713406 | 312968888 | 0.402568 | 0.467695 | 0.567338 |
| 7 | results/Baron2016.knn-species/n\_nns-10.query-m... | 0.773256 | database | 8569 | 1886 | 10 | 33441033 | mouse | 0.309970 | 3611276 | 270618281 | 0.406496 | 0.480516 | 0.582185 |
| 8 | results/Baron2016.knn-species/n\_nns-10.query-m... | 0.765204 | database | 8569 | 1886 | 10 | 33009988 | mouse | 0.329575 | 3630709 | 290771587 | 0.445199 | 0.414089 | 0.512725 |
| 9 | results/Baron2016.knn-species/n\_nns-10.query-m... | 0.801739 | database | 8569 | 1886 | 10 | 34376533 | mouse | 0.349429 | 3764286 | 309040595 | 0.397220 | 0.473617 | 0.573171 |
| 10 | results/Baron2016.knn-species/n\_nns-10.query-m... | 0.771486 | both | 8569 | 1886 | 10 | 37141193 | mouse | 0.327573 | 3813030 | 284417498 | 0.408034 | 0.526010 | 0.622481 |
| 11 | results/Baron2016.knn-species/n\_nns-10.query-m... | 0.771699 | both | 8569 | 1886 | 10 | 37634239 | mouse | 0.343803 | 3854522 | 299968601 | 0.449726 | 0.595460 | 0.687169 |
| 12 | results/Baron2016.knn-species/n\_nns-10.query-m... | 0.778782 | both | 8569 | 1886 | 10 | 36761681 | mouse | 0.328937 | 3814393 | 286108736 | 0.424304 | 0.548964 | 0.644751 |
| 13 | results/Baron2016.knn-species/n\_nns-10.query-m... | 0.787213 | both | 8569 | 1886 | 10 | 37775770 | mouse | 0.341566 | 3815668 | 297655097 | 0.419717 | 0.511584 | 0.608696 |
| 14 | results/Baron2016.knn-species/n\_nns-10.query-m... | 0.768953 | both | 8569 | 1886 | 10 | 37694533 | mouse | 0.339578 | 3782283 | 295797842 | 0.441854 | 0.585310 | 0.677625 |
| 15 | results/Baron2016.knn-species/n\_nns-10.query-h... | 0.172224 | query | 1886 | 8569 | 10 | 177457388 | human | 1.269630 | 15934453 | 1073915885 | 0.490496 | 0.599429 | 0.680943 |
| 16 | results/Baron2016.knn-species/n\_nns-10.query-h... | 0.166375 | query | 1886 | 8569 | 10 | 178760904 | human | 1.208241 | 16033892 | 1011063600 | 0.500296 | 0.595276 | 0.675341 |
| 17 | results/Baron2016.knn-species/n\_nns-10.query-h... | 0.176876 | query | 1886 | 8569 | 10 | 179259148 | human | 1.240794 | 15892804 | 1043222727 | 0.497955 | 0.608721 | 0.688178 |
| 18 | results/Baron2016.knn-species/n\_nns-10.query-h... | 0.165778 | query | 1886 | 8569 | 10 | 182565945 | human | 1.250201 | 15885618 | 1049124305 | 0.513354 | 0.609662 | 0.688178 |
| 19 | results/Baron2016.knn-species/n\_nns-10.query-h... | 0.166396 | query | 1886 | 8569 | 10 | 182017011 | human | 1.263320 | 16166594 | 1062681406 | 0.489869 | 0.595236 | 0.677209 |
| 20 | results/Baron2016.knn-species/n\_nns-10.query-h... | 0.179293 | database | 1886 | 8569 | 10 | 168662426 | human | 1.255497 | 15374887 | 1068851604 | 0.428792 | 0.531667 | 0.620959 |
| 21 | results/Baron2016.knn-species/n\_nns-10.query-h... | 0.178917 | database | 1886 | 8569 | 10 | 168786208 | human | 1.242249 | 15328302 | 1055613499 | 0.432310 | 0.531348 | 0.621309 |
| 22 | results/Baron2016.knn-species/n\_nns-10.query-h... | 0.163544 | database | 1886 | 8569 | 10 | 169421901 | human | 1.281383 | 15379457 | 1094039674 | 0.402185 | 0.496642 | 0.592718 |
| 23 | results/Baron2016.knn-species/n\_nns-10.query-h... | 0.174869 | database | 1886 | 8569 | 10 | 168348943 | human | 1.149910 | 15060105 | 963974172 | 0.397057 | 0.458730 | 0.558875 |
| 24 | results/Baron2016.knn-species/n\_nns-10.query-h... | 0.168301 | database | 1886 | 8569 | 10 | 170273685 | human | 1.229251 | 15146050 | 1041110074 | 0.414118 | 0.498335 | 0.594585 |
| 25 | results/Baron2016.knn-species/n\_nns-10.query-h... | 0.172814 | both | 1886 | 8569 | 10 | 192268175 | human | 1.229972 | 15921786 | 1019258242 | 0.574908 | 0.651010 | 0.720387 |
| 26 | results/Baron2016.knn-species/n\_nns-10.query-h... | 0.179699 | both | 1886 | 8569 | 10 | 194848506 | human | 1.186444 | 15711187 | 973004843 | 0.589214 | 0.646148 | 0.714552 |
| 27 | results/Baron2016.knn-species/n\_nns-10.query-h... | 0.163050 | both | 1886 | 8569 | 10 | 191819781 | human | 1.226071 | 15902218 | 1015811669 | 0.577940 | 0.655060 | 0.722838 |
| 28 | results/Baron2016.knn-species/n\_nns-10.query-h... | 0.164571 | both | 1886 | 8569 | 10 | 196346516 | human | 1.227923 | 15966670 | 1013128787 | 0.596315 | 0.657014 | 0.724239 |
| 29 | results/Baron2016.knn-species/n\_nns-10.query-h... | 0.174641 | both | 1886 | 8569 | 10 | 193702665 | human | 1.226928 | 16008870 | 1014675886 | 0.562397 | 0.643619 | 0.714669 |

In [63]:

```
knn_species_scmap = toml.load("results/Baron2016.knn-species.scmap.toml")
print(knn_species_scmap["date-time"])
print(knn_species_scmap["session-info"])
knn_species_scmap = pandas.DataFrame(knn_species_scmap["experiment"])
```

```
2018-07-09 05:43:12
R version 3.5.0 (2018-04-23)
Platform: x86_64-pc-linux-gnu (64-bit)
Running under: Debian GNU/Linux 9 (stretch)

Matrix products: default
BLAS: /usr/lib/openblas-base/libblas.so.3
LAPACK: /usr/lib/libopenblasp-r0.2.19.so

locale:
 [1] LC_CTYPE=en_US.UTF-8       LC_NUMERIC=C              
 [3] LC_TIME=en_US.UTF-8        LC_COLLATE=en_US.UTF-8    
 [5] LC_MONETARY=en_US.UTF-8    LC_MESSAGES=C             
 [7] LC_PAPER=en_US.UTF-8       LC_NAME=C                 
 [9] LC_ADDRESS=C               LC_TELEPHONE=C            
[11] LC_MEASUREMENT=en_US.UTF-8 LC_IDENTIFICATION=C       

attached base packages:
[1] parallel  stats4    stats     graphics  grDevices utils     datasets 
[8] methods   base     

other attached packages:
 [1] h5_0.9.9                    scmap_1.2.0                
 [3] scater_1.8.0                ggplot2_2.2.1              
 [5] SingleCellExperiment_1.2.0  SummarizedExperiment_1.10.1
 [7] DelayedArray_0.6.1          BiocParallel_1.14.1        
 [9] matrixStats_0.53.1          Biobase_2.40.0             
[11] GenomicRanges_1.32.3        GenomeInfoDb_1.16.0        
[13] IRanges_2.14.10             S4Vectors_0.18.3           
[15] BiocGenerics_0.26.0         readr_1.1.1                

loaded via a namespace (and not attached):
 [1] viridis_0.5.1            edgeR_3.22.2             jsonlite_1.5            
 [4] viridisLite_0.3.0        DelayedMatrixStats_1.2.0 shiny_1.1.0             
 [7] assertthat_0.2.0         GenomeInfoDbData_1.1.0   vipor_0.4.5             
[10] pillar_1.2.2             lattice_0.20-35          glue_1.2.0              
[13] limma_3.36.1             digest_0.6.15            promises_1.0.1          
[16] XVector_0.20.0           randomForest_4.6-14      colorspace_1.3-2        
[19] htmltools_0.3.6          httpuv_1.4.3             Matrix_1.2-14           
[22] plyr_1.8.4               pkgconfig_2.0.1          zlibbioc_1.26.0         
[25] purrr_0.2.5              xtable_1.8-2             scales_0.5.0            
[28] later_0.7.3              proxy_0.4-22             tibble_1.4.2            
[31] lazyeval_0.2.1           magrittr_1.5             mime_0.5                
[34] class_7.3-14             beeswarm_0.2.3           shinydashboard_0.7.0    
[37] tools_3.5.0              data.table_1.11.4        hms_0.4.2               
[40] stringr_1.3.0            googleVis_0.6.2          Rhdf5lib_1.2.1          
[43] munsell_0.5.0            locfit_1.5-9.1           bindrcpp_0.2.2          
[46] compiler_3.5.0           e1071_1.6-8              rlang_0.2.0             
[49] rhdf5_2.24.0             grid_3.5.0               RCurl_1.95-4.10         
[52] tximport_1.8.0           rjson_0.2.20             bitops_1.0-6            
[55] gtable_0.2.0             reshape2_1.4.3           R6_2.2.2                
[58] gridExtra_2.3            dplyr_0.7.5              bindr_0.1.1             
[61] stringi_1.2.2            ggbeeswarm_0.6.0         Rcpp_0.12.16            
[64] tidyselect_0.2.4
```

In [64]:

```
items = []
for row in knn_species_scmap.itertuples():
    if row.query == "mouse":
        query = "Baron2016-mouse"
        reference = "Baron2016-human"
    else:
        query = "Baron2016-human"
        reference = "Baron2016-mouse"
    items.append(cluster_metrics(row.filename, clusters[query], cluster_nn=clusters[reference]))
knn_species_scmap = knn_species_scmap.join(pandas.DataFrame(items))
knn_species_scmap
```

Out[64]:

|  | filename | index-time | n-cells-database | n-cells-query | n-genes | n-nns | query | query-time | adjusted\_rand | cohen\_kappa | consistency |
| --- | --- | --- | --- | --- | --- | --- | --- | --- | --- | --- | --- |
| 0 | results/Baron2016.knn-species.scmap//n\_nns-10.... | 14.029520 | 8569 | 1886 | 12413 | 10 | mouse | 5.021914 | 0.686938 | 0.753460 | 0.831919 |
| 1 | results/Baron2016.knn-species.scmap//n\_nns-10.... | 12.674630 | 8569 | 1886 | 12413 | 10 | mouse | 4.987169 | 0.689099 | 0.749580 | 0.828738 |
| 2 | results/Baron2016.knn-species.scmap//n\_nns-10.... | 12.126720 | 8569 | 1886 | 12413 | 10 | mouse | 4.987212 | 0.721139 | 0.775076 | 0.845175 |
| 3 | results/Baron2016.knn-species.scmap//n\_nns-10.... | 12.255130 | 8569 | 1886 | 12413 | 10 | mouse | 5.063193 | 0.673505 | 0.746800 | 0.828208 |
| 4 | results/Baron2016.knn-species.scmap//n\_nns-10.... | 12.812600 | 8569 | 1886 | 12413 | 10 | mouse | 4.988313 | 0.700171 | 0.762435 | 0.837222 |
| 5 | results/Baron2016.knn-species.scmap//n\_nns-10.... | 2.060711 | 1886 | 8569 | 12413 | 10 | human | 39.585630 | 0.444698 | 0.469996 | 0.574046 |
| 6 | results/Baron2016.knn-species.scmap//n\_nns-10.... | 1.676198 | 1886 | 8569 | 12413 | 10 | human | 39.781850 | 0.443234 | 0.454600 | 0.562143 |
| 7 | results/Baron2016.knn-species.scmap//n\_nns-10.... | 1.670642 | 1886 | 8569 | 12413 | 10 | human | 40.327050 | 0.444597 | 0.455020 | 0.562609 |
| 8 | results/Baron2016.knn-species.scmap//n\_nns-10.... | 1.764882 | 1886 | 8569 | 12413 | 10 | human | 39.352410 | 0.432666 | 0.448977 | 0.557708 |
| 9 | results/Baron2016.knn-species.scmap//n\_nns-10.... | 1.687775 | 1886 | 8569 | 12413 | 10 | human | 39.794070 | 0.437458 | 0.460456 | 0.567978 |

In [65]:

```
knn_species.groupby(["query", "infer-stats"]).describe()[["consistency","cohen_kappa"]]
```

Out[65]:

|  |  | consistency | | | | | | | | cohen\_kappa | | | | | | | |
| --- | --- | --- | --- | --- | --- | --- | --- | --- | --- | --- | --- | --- | --- | --- | --- | --- | --- |
|  |  | count | mean | std | min | 25% | 50% | 75% | max | count | mean | std | min | 25% | 50% | 75% | max |
| query | infer-stats |  |  |  |  |  |  |  |  |  |  |  |  |  |  |  |  |
| human | both | 5.0 | 0.719337 | 0.004530 | 0.714552 | 0.714669 | 0.720387 | 0.722838 | 0.724239 | 5.0 | 0.650570 | 0.005695 | 0.643619 | 0.646148 | 0.651010 | 0.655060 | 0.657014 |
|  | database | 5.0 | 0.597689 | 0.025692 | 0.558875 | 0.592718 | 0.594585 | 0.620959 | 0.621309 | 5.0 | 0.503345 | 0.030195 | 0.458730 | 0.496642 | 0.498335 | 0.531348 | 0.531667 |
|  | query | 5.0 | 0.681970 | 0.006016 | 0.675341 | 0.677209 | 0.680943 | 0.688178 | 0.688178 | 5.0 | 0.601665 | 0.007087 | 0.595236 | 0.595276 | 0.599429 | 0.608721 | 0.609662 |
| mouse | both | 5.0 | 0.648144 | 0.033979 | 0.608696 | 0.622481 | 0.644751 | 0.677625 | 0.687169 | 5.0 | 0.553465 | 0.036420 | 0.511584 | 0.526010 | 0.548964 | 0.585310 | 0.595460 |
|  | database | 5.0 | 0.544645 | 0.041796 | 0.487805 | 0.512725 | 0.567338 | 0.573171 | 0.582185 | 5.0 | 0.445749 | 0.039591 | 0.392828 | 0.414089 | 0.467695 | 0.473617 | 0.480516 |
|  | query | 5.0 | 0.778685 | 0.021051 | 0.742312 | 0.780488 | 0.787381 | 0.787381 | 0.795864 | 5.0 | 0.704485 | 0.025997 | 0.659480 | 0.706694 | 0.714670 | 0.716453 | 0.725126 |

In [66]:

```
knn_species_scmap.groupby(["query"]).describe()[["consistency","cohen_kappa"]]
```

Out[66]:

|  | consistency | | | | | | | | cohen\_kappa | | | | | | | |
| --- | --- | --- | --- | --- | --- | --- | --- | --- | --- | --- | --- | --- | --- | --- | --- | --- |
|  | count | mean | std | min | 25% | 50% | 75% | max | count | mean | std | min | 25% | 50% | 75% | max |
| query |  |  |  |  |  |  |  |  |  |  |  |  |  |  |  |  |
| human | 5.0 | 0.564897 | 0.006279 | 0.557708 | 0.562143 | 0.562609 | 0.567978 | 0.574046 | 5.0 | 0.45781 | 0.007931 | 0.448977 | 0.45460 | 0.45502 | 0.460456 | 0.469996 |
| mouse | 5.0 | 0.834252 | 0.007080 | 0.828208 | 0.828738 | 0.831919 | 0.837222 | 0.845175 | 5.0 | 0.75747 | 0.011476 | 0.746800 | 0.74958 | 0.75346 | 0.762435 | 0.775076 |

#### Mapping across protocols¶

In [67]:

```
len(clusters["TabulaMuris-chromium"].unique())
```

Out[67]:

```
57
```

In [68]:

```
len(clusters["TabulaMuris-smart"].unique())
```

Out[68]:

```
81
```

In [69]:

```
overlappings = set(clusters["TabulaMuris-chromium"].unique()) & set(clusters["TabulaMuris-smart"].unique())
len(overlappings)
```

Out[69]:

```
38
```

In [70]:

```
nns = load_knn("results/TabulaMuris.knn-protocol/n_nns-10.1.tsv.gz")
plot_assignments_matrix(nns.index.values, nns["N1"].values, clusters["TabulaMuris-smart"], clusters["TabulaMuris-chromium"], figsize=(12, 14))
savefig("TabulaMuris-smart-chromium-matrix")
```

In [71]:

```
plot_assignments_matrix(nns.index.values, nns["N1"].values,
                        clusters["TabulaMuris-smart"][clusters["TabulaMuris-smart"].isin(overlappings)],
                        clusters["TabulaMuris-chromium"][clusters["TabulaMuris-chromium"].isin(overlappings)],
                        overlapping=True, figsize=(12, 14))
savefig("TabulaMuris-smart-chromium-matrix-overlapping")
```

In [72]:

```
nns = load_knn("results/TabulaMuris.knn-protocol.scmap/n_nns-10.1.tsv.gz")
plot_assignments_matrix(nns.index.values, nns["N1"].values, clusters["TabulaMuris-smart"], clusters["TabulaMuris-chromium"], figsize=(12, 14))
savefig("TabulaMuris-smart-chromium-matrix-scmap")
```

In [73]:

```
plot_assignments_matrix(nns.index.values, nns["N1"].values,
                        clusters["TabulaMuris-smart"][clusters["TabulaMuris-smart"].isin(overlappings)],
                        clusters["TabulaMuris-chromium"][clusters["TabulaMuris-chromium"].isin(overlappings)],
                        overlapping=True, figsize=(12, 14))
savefig("TabulaMuris-smart-chromium-matrix-scmap-overlapping")
```

#### Benachmarks of saving and loading¶

In [74]:

```
saveload = pandas.DataFrame()
for dataset in datasets:
    out = toml.load(f"results/{dataset}.save-load.toml")
    print(out["date-time"])
    print(out["version-info"])
    exp = pandas.DataFrame(out["experiment"])
    exp["dataset"] = dataset
    for x in exp.columns:
        if x.endswith("-size"):
            exp[x] /= 1024**2  # B => MiB
        if x.endswith("-time"):
            exp[x] *= 1000  # s => ms
    saveload = saveload.append(exp)
```

```
2018-11-23 21:21:00.212000
Julia Version 1.0.1
Commit 0d713926f8 (2018-09-29 19:05 UTC)
Platform Info:
  OS: Linux (x86_64-pc-linux-gnu)
  CPU: Intel(R) Xeon(R) Gold 6126 CPU @ 2.60GHz
  WORD_SIZE: 64
  LIBM: libopenlibm
  LLVM: libLLVM-6.0.0 (ORCJIT, skylake)
Environment:
  JULIA_PROJECT = @.

2018-11-23 16:54:36.303000
Julia Version 1.0.1
Commit 0d713926f8 (2018-09-29 19:05 UTC)
Platform Info:
  OS: Linux (x86_64-pc-linux-gnu)
  CPU: Intel(R) Xeon(R) Gold 6126 CPU @ 2.60GHz
  WORD_SIZE: 64
  LIBM: libopenlibm
  LLVM: libLLVM-6.0.0 (ORCJIT, skylake)
Environment:
  JULIA_PROJECT = @.

2018-11-23 20:16:15.891000
Julia Version 1.0.1
Commit 0d713926f8 (2018-09-29 19:05 UTC)
Platform Info:
  OS: Linux (x86_64-pc-linux-gnu)
  CPU: Intel(R) Xeon(R) Gold 6126 CPU @ 2.60GHz
  WORD_SIZE: 64
  LIBM: libopenlibm
  LLVM: libLLVM-6.0.0 (ORCJIT, skylake)
Environment:
  JULIA_PROJECT = @.

2018-11-23 14:33:35.055000
Julia Version 1.0.1
Commit 0d713926f8 (2018-09-29 19:05 UTC)
Platform Info:
  OS: Linux (x86_64-pc-linux-gnu)
  CPU: Intel(R) Xeon(R) Gold 6126 CPU @ 2.60GHz
  WORD_SIZE: 64
  LIBM: libopenlibm
  LLVM: libLLVM-6.0.0 (ORCJIT, skylake)
Environment:
  JULIA_PROJECT = @.
```

In [75]:

```
saveload_median = saveload\
    .groupby(["dataset", "n-bits", "keep-counts"])\
    .median()\
    [["n-cells", "n-genes",
      "ram-size", "file-size",
      "save-time", "load-time",]]
saveload_median
```

Out[75]:

|  |  |  | n-cells | n-genes | ram-size | file-size | save-time | load-time |
| --- | --- | --- | --- | --- | --- | --- | --- | --- |
| dataset | n-bits | keep-counts |  |  |  |  |  |  |
| Baron2016-human | 128 | False | 8569 | 20125 | 3.082169 | 2.933306 | 2.333969 | 2.443834 |
|  |  | True | 8569 | 20125 | 10.856806 | 10.707905 | 6.450365 | 3.060793 |
|  | 256 | False | 8569 | 20125 | 5.390549 | 5.236307 | 3.677291 | 5.282122 |
|  |  | True | 8569 | 20125 | 13.165186 | 13.010906 | 7.958927 | 3.581396 |
| Plass2018 | 128 | False | 21612 | 28065 | 6.714789 | 6.379647 | 4.667075 | 4.981560 |
|  |  | True | 21612 | 28065 | 17.491735 | 17.156554 | 10.449806 | 6.199515 |
|  | 256 | False | 21612 | 28065 | 11.985328 | 11.645482 | 8.027909 | 9.599168 |
|  |  | True | 21612 | 28065 | 22.762273 | 22.422389 | 14.861338 | 9.230306 |
| Shekhar2016 | 128 | False | 27499 | 24904 | 8.194370 | 7.778868 | 5.614199 | 8.280028 |
|  |  | True | 27499 | 24904 | 29.709252 | 29.293712 | 16.287731 | 6.665521 |
|  | 256 | False | 27499 | 24904 | 14.564318 | 14.144112 | 9.140735 | 8.255374 |
|  |  | True | 27499 | 24904 | 36.079200 | 35.658956 | 20.740924 | 11.054664 |
| TabulaMuris-chromium | 128 | False | 54967 | 23337 | 13.814836 | 13.044705 | 8.584768 | 12.476224 |
|  |  | True | 54967 | 23337 | 67.231367 | 66.461199 | 35.257272 | 16.862708 |
|  | 256 | False | 54967 | 23337 | 24.930642 | 24.156331 | 13.796409 | 17.974066 |
|  |  | True | 54967 | 23337 | 78.347174 | 77.572824 | 42.397889 | 21.938694 |

Generate a LaTeX table.

In [76]:

```
print(r"""
\begin{tabular}{l cc rrrr}
\toprule
Data set & \#bits & Raw counts & \multicolumn{2}{c}{Size [MiB]} & \multicolumn{2}{c}{Time [ms]} \\
\cmidrule(lr){4-5} \cmidrule(lr){6-7}
         &        &            & Memory & File                    & Save & Load \\""")
for i, dataset in enumerate(datasets):
    if i == 0:
        print(r"\midrule")
    else:
        print(r"\addlinespace")
    print(r"\multirow{4}{*}{", dataset, "}")
    for n_bits in [128, 256]:
        print(r"  & \multirow{2}{*}{", n_bits, "} ", end="")
        for keep_counts in [False, True]:
            if keep_counts:
                print("  &                        ", end="")
            print(" &", r"\texttt{+}" if keep_counts else r"\texttt{-}", end="")
            for col in ["ram-size", "file-size", "save-time", "load-time"]:
                row = saveload_median.loc[(dataset, n_bits, keep_counts)]
                print(" & {:4.1f} ".format(row[col]), end="")
            print(r"\\")
print(r"""\bottomrule
\end{tabular}
""")
```

```
\begin{tabular}{l cc rrrr}
\toprule
Data set & \#bits & Raw counts & \multicolumn{2}{c}{Size [MiB]} & \multicolumn{2}{c}{Time [ms]} \\
\cmidrule(lr){4-5} \cmidrule(lr){6-7}
         &        &            & Memory & File                    & Save & Load \\
\midrule
\multirow{4}{*}{ Baron2016-human }
  & \multirow{2}{*}{ 128 }  & \texttt{-} &  3.1  &  2.9  &  2.3  &  2.4 \\
  &                         & \texttt{+} & 10.9  & 10.7  &  6.5  &  3.1 \\
  & \multirow{2}{*}{ 256 }  & \texttt{-} &  5.4  &  5.2  &  3.7  &  5.3 \\
  &                         & \texttt{+} & 13.2  & 13.0  &  8.0  &  3.6 \\
\addlinespace
\multirow{4}{*}{ Shekhar2016 }
  & \multirow{2}{*}{ 128 }  & \texttt{-} &  8.2  &  7.8  &  5.6  &  8.3 \\
  &                         & \texttt{+} & 29.7  & 29.3  & 16.3  &  6.7 \\
  & \multirow{2}{*}{ 256 }  & \texttt{-} & 14.6  & 14.1  &  9.1  &  8.3 \\
  &                         & \texttt{+} & 36.1  & 35.7  & 20.7  & 11.1 \\
\addlinespace
\multirow{4}{*}{ Plass2018 }
  & \multirow{2}{*}{ 128 }  & \texttt{-} &  6.7  &  6.4  &  4.7  &  5.0 \\
  &                         & \texttt{+} & 17.5  & 17.2  & 10.4  &  6.2 \\
  & \multirow{2}{*}{ 256 }  & \texttt{-} & 12.0  & 11.6  &  8.0  &  9.6 \\
  &                         & \texttt{+} & 22.8  & 22.4  & 14.9  &  9.2 \\
\addlinespace
\multirow{4}{*}{ TabulaMuris-chromium }
  & \multirow{2}{*}{ 128 }  & \texttt{-} & 13.8  & 13.0  &  8.6  & 12.5 \\
  &                         & \texttt{+} & 67.2  & 66.5  & 35.3  & 16.9 \\
  & \multirow{2}{*}{ 256 }  & \texttt{-} & 24.9  & 24.2  & 13.8  & 18.0 \\
  &                         & \texttt{+} & 78.3  & 77.6  & 42.4  & 21.9 \\
\bottomrule
\end{tabular}
```

#### Scalability¶

In [77]:

```
dataset = "1M_neurons"
```

In [78]:

```
out = toml.load(f"results/{dataset}.scalability.toml")
print(out["date-time"])
print(out["version-info"])
scalability = pandas.DataFrame(out["experiment"])
scalability["ram-size"] /= 1024**2  # byte => MiB
scalability = pandas.concat(
    [scalability, pandas.DataFrame([cluster_metrics(x, clusters[dataset], k=1) for x in scalability["filename"]])],
    axis=1)
scalability = scalability.drop(["n-cells-query", "n-genes", "n-nns"], axis=1)  # drop constant columns
```

```
2018-11-24 15:11:38.828000
Julia Version 1.0.1
Commit 0d713926f8 (2018-09-29 19:05 UTC)
Platform Info:
  OS: Linux (x86_64-pc-linux-gnu)
  CPU: Intel(R) Xeon(R) CPU E5-2637 v4 @ 3.50GHz
  WORD_SIZE: 64
  LIBM: libopenlibm
  LLVM: libLLVM-6.0.0 (ORCJIT, broadwell)
Environment:
  JULIA_PROJECT = @.
```

In [79]:

```
scalability.groupby(["n-bits", "n-cells-database", "indexed"]).median()
```

Out[79]:

|  |  |  | index-time | preproc-time | query-time | ram-size | rank-time | search-time | adjusted\_rand | cohen\_kappa | consistency |
| --- | --- | --- | --- | --- | --- | --- | --- | --- | --- | --- | --- |
| n-bits | n-cells-database | indexed |  |  |  |  |  |  |  |  |  |
| 128 | 8192 | False | 1.365656 | 1330290302 | 1.797388 | 1.303389 | 18220000 | 443179509 | 0.618090 | 0.765209 | 0.7751 |
|  |  | True | 1.448293 | 1322688601 | 2.248178 | 2.977102 | 18020163 | 887183066 | 0.617982 | 0.765105 | 0.7750 |
|  | 16384 | False | 2.482641 | 1296051170 | 2.127062 | 1.769693 | 19537110 | 813080198 | 0.638576 | 0.780841 | 0.7900 |
|  |  | True | 2.541771 | 1297755890 | 2.192187 | 4.783045 | 19238684 | 853255043 | 0.638778 | 0.780944 | 0.7901 |
|  | 32768 | False | 4.537608 | 1338261675 | 2.863432 | 2.819775 | 20510502 | 1508250510 | 0.658046 | 0.797111 | 0.8057 |
|  |  | True | 4.669582 | 1332525822 | 2.402167 | 8.321144 | 19956847 | 1037183573 | 0.657743 | 0.796963 | 0.8055 |
|  | 65536 | False | 8.526317 | 1314085930 | 4.243564 | 4.812159 | 21575883 | 2904284234 | 0.663753 | 0.802872 | 0.8111 |
|  |  | True | 8.634133 | 1321695737 | 2.405027 | 14.770630 | 21110887 | 1058982545 | 0.663429 | 0.802979 | 0.8112 |
|  | 131072 | False | 16.355783 | 1312527883 | 7.030339 | 8.798080 | 23946077 | 5692370445 | 0.672096 | 0.808708 | 0.8167 |
|  |  | True | 16.547326 | 1405899402 | 2.864753 | 27.101412 | 22902152 | 1412383485 | 0.672669 | 0.809172 | 0.8172 |
|  | 262144 | False | 31.853674 | 1275638550 | 12.566691 | 16.771540 | 31973860 | 11261905409 | 0.677865 | 0.811876 | 0.8198 |
|  |  | True | 31.965757 | 1256338321 | 2.950319 | 51.014181 | 29122782 | 1678035027 | 0.678635 | 0.812547 | 0.8204 |
|  | 524288 | False | 63.290781 | 1286824049 | 23.688397 | 32.781694 | 37505437 | 22368780480 | 0.688553 | 0.820310 | 0.8279 |
|  |  | True | 64.130570 | 1309795244 | 4.248334 | 93.018644 | 37578602 | 2889820480 | 0.688548 | 0.820106 | 0.8277 |
|  | 1048576 | False | 121.284624 | 1215232525 | 52.886248 | 64.746845 | 42698142 | 51616138013 | 0.692556 | 0.821616 | 0.8291 |
|  |  | True | 125.002472 | 1223074661 | 6.324869 | 183.274239 | 41975407 | 5052890613 | 0.694702 | 0.822339 | 0.8298 |
| 256 | 8192 | False | 1.422101 | 1294464879 | 2.034880 | 1.871273 | 19021191 | 717970487 | 0.625441 | 0.772651 | 0.7821 |
|  |  | True | 1.433398 | 1289761606 | 3.358104 | 5.237467 | 18772318 | 2028273109 | 0.625441 | 0.772651 | 0.7821 |
|  | 16384 | False | 2.715621 | 1344538676 | 2.646955 | 2.904737 | 20046113 | 1278591025 | 0.642372 | 0.785940 | 0.7949 |
|  |  | True | 2.664941 | 1343113695 | 3.637275 | 8.925112 | 19765457 | 2283677601 | 0.642372 | 0.785940 | 0.7949 |
|  | 32768 | False | 4.764645 | 1307671651 | 3.751781 | 4.883736 | 21273538 | 2422754799 | 0.658642 | 0.795639 | 0.8042 |
|  |  | True | 4.687539 | 1278897919 | 3.938566 | 15.807877 | 20946822 | 2633239003 | 0.658642 | 0.795639 | 0.8042 |
|  | 65536 | False | 8.920253 | 1338783335 | 6.076809 | 8.909584 | 24243157 | 4699550779 | 0.676966 | 0.812144 | 0.8200 |
|  |  | True | 9.001467 | 1313204186 | 4.594189 | 28.848030 | 22654988 | 3258006372 | 0.676966 | 0.812144 | 0.8200 |
|  | 131072 | False | 16.148443 | 1253268340 | 10.523046 | 16.854425 | 28620320 | 9231525437 | 0.677127 | 0.813099 | 0.8209 |
|  |  | True | 16.804400 | 1238547622 | 5.211698 | 53.696632 | 28027527 | 3941549035 | 0.677127 | 0.813099 | 0.8209 |
|  | 262144 | False | 32.312373 | 1271301274 | 19.611065 | 32.853964 | 37967189 | 18302813573 | 0.693016 | 0.819649 | 0.8272 |
|  |  | True | 33.727287 | 1285896064 | 8.789510 | 101.330185 | 48950697 | 7453217565 | 0.692863 | 0.819543 | 0.8271 |
|  | 524288 | False | 65.024346 | 1286554461 | 46.660451 | 64.882582 | 43336511 | 45352689828 | 0.695216 | 0.824645 | 0.8320 |
|  |  | True | 68.603339 | 1322193069 | 13.352726 | 192.900106 | 45510119 | 11981324216 | 0.695613 | 0.824854 | 0.8322 |
|  | 1048576 | False | 125.434165 | 1409368494 | 109.339366 | 128.848425 | 48053660 | 107885552313 | 0.700789 | 0.829543 | 0.8367 |
|  |  | True | 132.198818 | 1264752278 | 22.079090 | 365.941727 | 47175564 | 20606257565 | 0.700238 | 0.829677 | 0.8368 |

In [80]:

```
fig, axes = subplots(4, 2, dpi=dpi, figsize=(8, 6), sharex=True)
for i, metric in enumerate(["index-time", "query-time", "ram-size", "consistency"]):
    for j, n_bits in enumerate([128, 256]):
        ax = axes[i,j]
        for indexed in [False, True]:
            data = scalability[(scalability["n-bits"]==n_bits)&(scalability["indexed"]==indexed)].groupby("n-cells-database").median().reset_index()
            ax.plot(data["n-cells-database"], data[metric], ".-", label="index" if indexed else "linear")
        ax.set_xscale("log", basex=2)
        if metric != "consistency":
            ax.set_yscale("log")
        if i == 0:
            ax.set_title(f"n-bits = {n_bits}")
        if i == 3:
            ax.set_xlabel("database size")
        if j == 0:
            if metric == "index-time":
                ax.set_ylabel("index time [s]")
            elif metric == "query-time":
                ax.set_ylabel("query time [s]")
            elif metric == "ram-size":
                ax.set_ylabel("memory size [MiB]")
            else:
                ax.set_ylabel("consistency")
        if i == 0 and j == 0:
            ax.legend()
seaborn.despine(fig=fig)
fig.tight_layout()
savefig("scalability-metrics")
```

Generate a LaTeX table.

In [81]:

```
scalability_median = scalability.groupby(["n-bits", "n-cells-database", "indexed"]).median()
indexed = True
print(r"""
\begin{tabular}{l cc rrrrrrrr}
\toprule
& \#bits  & Index & \multicolumn{8}{c}{Database size $N$}\\
\cmidrule{4-11}
&         &       & $2^{13}$ (1.0) & $2^{14}$ (2.0) & $2^{15}$ (4.0) & $2^{16}$ (8.0) & $2^{17}$ (16.0) & $2^{18}$ (32.0) & $2^{19}$ (64.0) & $2^{20}$ (128.0)\\
""")
scales = numpy.arange(13, 21)
for i, metric in enumerate(["index-time", "query-time", "ram-size"]):
    if i == 0:
        print(r"\midrule")
    else:
        print(r"\addlinespace")
    print(r"\multirow{4}{*}{", end="")
    if metric == "index-time":
        print("Index time [s]", end="")
    elif metric == "query-time":
        print("Query time [s]", end="")
    else:
        print("Memory size [MiB]", end="")
    print(r"}")
    for n_bits in [128, 256]:
        for indexed in [False, True]:
            print(r"  &", n_bits, "&", r"\texttt{+}" if indexed else r"\texttt{-}", end="")
            base = scalability_median[metric].loc[n_bits,2**scales[0],indexed]
            for s in scales:
                n = 2**s
                val = scalability_median[metric].loc[n_bits,n,indexed]
                print(" & {:5.1f} ({:.1f})".format(val, val/base), end="")
            print(r"\\")
print(r"\bottomrule")
print(r"\end{tabular}")
```

```
\begin{tabular}{l cc rrrrrrrr}
\toprule
& \#bits  & Index & \multicolumn{8}{c}{Database size $N$}\\
\cmidrule{4-11}
&         &       & $2^{13}$ (1.0) & $2^{14}$ (2.0) & $2^{15}$ (4.0) & $2^{16}$ (8.0) & $2^{17}$ (16.0) & $2^{18}$ (32.0) & $2^{19}$ (64.0) & $2^{20}$ (128.0)\\

\midrule
\multirow{4}{*}{Index time [s]}
  & 128 & \texttt{-} &   1.4 (1.0) &   2.5 (1.8) &   4.5 (3.3) &   8.5 (6.2) &  16.4 (12.0) &  31.9 (23.3) &  63.3 (46.3) & 121.3 (88.8)\\
  & 128 & \texttt{+} &   1.4 (1.0) &   2.5 (1.8) &   4.7 (3.2) &   8.6 (6.0) &  16.5 (11.4) &  32.0 (22.1) &  64.1 (44.3) & 125.0 (86.3)\\
  & 256 & \texttt{-} &   1.4 (1.0) &   2.7 (1.9) &   4.8 (3.4) &   8.9 (6.3) &  16.1 (11.4) &  32.3 (22.7) &  65.0 (45.7) & 125.4 (88.2)\\
  & 256 & \texttt{+} &   1.4 (1.0) &   2.7 (1.9) &   4.7 (3.3) &   9.0 (6.3) &  16.8 (11.7) &  33.7 (23.5) &  68.6 (47.9) & 132.2 (92.2)\\
\addlinespace
\multirow{4}{*}{Query time [s]}
  & 128 & \texttt{-} &   1.8 (1.0) &   2.1 (1.2) &   2.9 (1.6) &   4.2 (2.4) &   7.0 (3.9) &  12.6 (7.0) &  23.7 (13.2) &  52.9 (29.4)\\
  & 128 & \texttt{+} &   2.2 (1.0) &   2.2 (1.0) &   2.4 (1.1) &   2.4 (1.1) &   2.9 (1.3) &   3.0 (1.3) &   4.2 (1.9) &   6.3 (2.8)\\
  & 256 & \texttt{-} &   2.0 (1.0) &   2.6 (1.3) &   3.8 (1.8) &   6.1 (3.0) &  10.5 (5.2) &  19.6 (9.6) &  46.7 (22.9) & 109.3 (53.7)\\
  & 256 & \texttt{+} &   3.4 (1.0) &   3.6 (1.1) &   3.9 (1.2) &   4.6 (1.4) &   5.2 (1.6) &   8.8 (2.6) &  13.4 (4.0) &  22.1 (6.6)\\
\addlinespace
\multirow{4}{*}{Memory size [MiB]}
  & 128 & \texttt{-} &   1.3 (1.0) &   1.8 (1.4) &   2.8 (2.2) &   4.8 (3.7) &   8.8 (6.8) &  16.8 (12.9) &  32.8 (25.2) &  64.7 (49.7)\\
  & 128 & \texttt{+} &   3.0 (1.0) &   4.8 (1.6) &   8.3 (2.8) &  14.8 (5.0) &  27.1 (9.1) &  51.0 (17.1) &  93.0 (31.2) & 183.3 (61.6)\\
  & 256 & \texttt{-} &   1.9 (1.0) &   2.9 (1.6) &   4.9 (2.6) &   8.9 (4.8) &  16.9 (9.0) &  32.9 (17.6) &  64.9 (34.7) & 128.8 (68.9)\\
  & 256 & \texttt{+} &   5.2 (1.0) &   8.9 (1.7) &  15.8 (3.0) &  28.8 (5.5) &  53.7 (10.3) & 101.3 (19.3) & 192.9 (36.8) & 365.9 (69.9)\\
\bottomrule
\end{tabular}
```

In [82]:

```
n_cells = 10000
n_bits = 128
fig, axes = subplots(2, dpi=dpi, figsize=(8, 5))
for i, indexed in enumerate([False, True]):
    ax = axes[i]
    labels = []
    y = 0
    for s in scales:
        n = 2**s
        left = 0
        bars = []
        for j, metric in enumerate(["preproc-time", "search-time", "rank-time"]):
            time = scalability_median[metric].loc[n_bits,n,indexed] / 1000 / n_cells
            bars.append(ax.barh(y, time, left=left, color=f"C{j}"))
            left += time
        labels.append(f"$2^{{{s}}}$")
        y += 1
    ax.set_yticks(range(len(labels)))
    ax.set_yticklabels(labels)
    ax.set_ylabel("database size")
    ax.set_xlabel("elapsed time [μs/cell]")
    ax.legend(bars, ["preprocessing", "searching", "ranking"], title="phase")
    ax.invert_yaxis()
    if indexed:
        ax.set_title("index search")
    else:
        ax.set_title("linear search")
seaborn.despine(fig=fig)
fig.tight_layout();
savefig("scalability-computational-cost")
```
