## Supplementary material for "CellFishing.jl: an ultrafast and scalable cell search method for single-cell RNA-sequencing": Analysis.ipynb

```
# %% [markdown]
# Performance Analysis of CellFishing.jl

# %% [markdown]
## Table of contents

- [Table of contents](#Table-of-contents)
- [Cell-type distributions](#Cell-type-distributions)
- [Randomized SVD](#Randomized-SVD)
- [Similarity estimation via hashing](#Similarity-estimation-via-hashing)
- [Visualization of expression profiles](#Visualization-of-expression-profiles)
- [Self-mapping experiments](#Self-mapping-experiments)
- [Feature selection](#Feature-selection)
- [Cluster-specific scores](#Cluster-specific-scores)
- [Similarities from nearest neighbors](#Similarities-from-nearest-neighbors)
- [Detecting DEGs](#Detecting-DEGs)
- [Mapping across batches](#Mapping-across-batches)
- [Mapping across species](#Mapping-across-species)
- [Mapping across protocols](#Mapping-across-protocols)
- [Benchmarks of saving and loading](#Benachmarks-of-saving-and-loading)
- [Scalability](#Scalability)

# %% [code]
import os
import re
import math
import textwrap
import toml
import numpy
import pandas
import scipy.stats
import sklearn.metrics
from matplotlib.pyplot import subplots
import matplotlib.pyplot
import matplotlib_venn
import seaborn
import umap

# %% [code]
pkgname = "CellFishing"

# %% [code]
ext = "pdf"
dpi = 200
seaborn.set_style("ticks", {"image.cmap": "viridis"})

def savefig(name):
    matplotlib.pyplot.savefig(os.path.join("figures", f"{name}.{ext}"), dpi="figure", bbox_inches="tight")
    
!mkdir -p figures

# %% [markdown]
## Cell-type distributions

# %% [code]
datasets = ["Baron2016-human", "Shekhar2016", "Plass2018", "TabulaMuris-chromium"]

# %% [markdown]
Load cell-type annotations:

# %% [code]
clusters = dict()
for dataset in datasets:
    clusters[dataset] = pandas.read_table(f"data/{dataset}.cluster.tsv", index_col="cell")["cluster"]
clusters["Baron2016-mouse"] = pandas.read_table("data/Baron2016-mouse.cluster.tsv", index_col="cell")["cluster"]
clusters["Baron2016-human"][clusters["Baron2016-human"] == "t_cell"] = "T_cell"  # fix the name to match that of mouse
clusters["TabulaMuris-smart"] = pandas.read_table("data/TabulaMuris-smart.cluster.tsv", index_col="cell")["cluster"]
clusters["1M_neurons"] = pandas.read_table("data/1M_neurons.cluster.tsv", index_col="cell")["cluster"]

# %% [markdown]
Summary of datasets:

# %% [code]
print("{:20s}  {:>9}  {:>9s}".format("dataset", "#cells", "#clusters"))
print("------------------------------------------")
for dataset, cluster in sorted(clusters.items()):
    print("{:20s}  {:>9,d}  {:>9,d}".format(dataset, len(cluster), len(cluster.unique())))

# %% [markdown]
Cluster (cell type) distributions:

# %% [code]
for dataset, cluster in clusters.items():
    print(dataset)
    fig, ax = subplots(dpi=dpi, figsize=(8, 10))
    counts = cluster.value_counts()
    seaborn.countplot(
        y="cluster",
        data=pandas.DataFrame(cluster),
        order=counts.index.values,
        color="C0",
        ax=ax)
    fontsize = 6.5
    for i, c in enumerate(counts.index.values):
        ax.text(counts[c], i, " {:,} ({:.2f}%)".format(counts[c], counts[c] / len(cluster) * 100),
                verticalalignment="center",
                fontsize=fontsize)
    ax.tick_params(labelsize=fontsize)
    ax.set_xlabel("cluster size")
    seaborn.despine(fig=fig)
    savefig(f"{dataset}-clusters")

# %% [markdown]
## Randomized SVD

# %% [markdown]
Load benchmark results:

# %% [code]
def expand_singular_values(data):
    data = pandas.concat([
        pandas.DataFrame({"k": numpy.arange(1, r["n-dims"]+1), **dict(r)}) for ix, r in data.iterrows()], ignore_index=True)
    return data.rename(columns={"singular-values": "singular-value"})

svd = pandas.DataFrame()
for dataset in datasets:
    print(dataset)
    out = toml.load(f"results/{dataset}.svd.toml")
    print(out["date-time"])
    print(out["version-info"])
    expanded = expand_singular_values(pandas.DataFrame(out["experiment"]))
    expanded["dataset"] = dataset
    svd = svd.append(expanded, ignore_index=True)

# %% [markdown]
Matrix sizes:

# %% [code]
svd.groupby(["dataset"]).first()[["n-rows", "n-columns"]].loc[datasets,:]

# %% [markdown]
Summary of benchmarks:

# %% [code]
svd[svd["k"]==1].groupby(["dataset", "algorithm"]).describe()["time-svd"].loc[datasets,:]

# %% [markdown]
Plot the elapsed time:

# %% [code]
fig, ax = subplots(dpi=dpi, figsize=(8, 4))
seaborn.swarmplot(x="dataset", y="time-svd", hue="algorithm", data=svd[svd["k"]==1], dodge=True, size=3)
ax.set_ylabel("elapsed time [s]")
ax.set_xlabel("")
ax.set_ylim(0, None)
ax.locator_params(nbins=5)
ax.grid(axis="y", alpha=0.3)
seaborn.despine(fig=fig)
fig.tight_layout()
savefig(f"svd-benchmark")

# %% [markdown]
Plot the singular values:

# %% [code]
fig, axes = subplots(4, sharex=True, dpi=dpi, figsize=(8, 4))
for i, dataset in enumerate(datasets):
    ax = axes[i]
    seaborn.pointplot(x="k", y="singular-value", hue="algorithm", data=svd[svd["dataset"]==dataset], scale=0.5, ci="sd", ax=ax)
    if i == 3:
        ax.set_xlabel("rank")
    else:
        ax.set_xlabel("")
    ax.set_ylim(0, None)
    for k, label in enumerate(ax.xaxis.get_ticklabels()):
        if k % 2 == 1:
            label.set_visible(False)
    ax.legend_.remove()
    ax.locator_params(axis="y", nbins=1)
    ax.set_ylabel("$\sigma$")
    ax.set_title(dataset)
    ax.grid(axis="y", alpha=0.3)
seaborn.despine(fig=fig)
fig.tight_layout()
savefig(f"svd-singular-values");

# %% [markdown]
Compute the relative errors of the singular values:

# %% [code]
svals = svd[svd["algorithm"]=="full"]["singular-value"].values
relerror = svd[svd["algorithm"]=="rand"]\
    .assign(**{"relative-error": lambda x: numpy.abs(1 - x["singular-value"] / svals)})

# %% [markdown]
Plot the relative errors:

# %% [code]
fig, axes = subplots(4, sharex=True, dpi=dpi, figsize=(8, 4))
for i, dataset in enumerate(datasets):
    ax = axes[i]
    seaborn.pointplot(x="k", y="relative-error", data=relerror[relerror["dataset"]==dataset], scale=0.5, errwidth=1.0, ci="sd", ax=ax)
    if i == 3:
        ax.set_xlabel("rank")
    else:
        ax.set_xlabel("")
    ax.set_ylim(0, None)
    for k, label in enumerate(ax.xaxis.get_ticklabels()):
        if k % 2 == 1:
            label.set_visible(False)
    ax.set_ylabel("rel. error")
    ax.set_title(dataset)
    ax.grid(axis="y", alpha=0.3)
seaborn.despine(fig=fig)
fig.tight_layout()
savefig(f"svd-relative-errors");

# %% [markdown]
## Similarity estimation via hashing

# %% [code]
print("{:20s}  {:>6s}  {:>8s}  {:>7s}  {:>6s}".format("dataset", "n-bits", "superbit", "mean", "std"))
for dataset in datasets:
    fig, axes = subplots(4, dpi=dpi, figsize=(8, 5), sharex=True)
    for i, n_bits in enumerate([64, 128, 256, 512]):
        ax = axes[i]
        ax.set_title(f"n-bits = {n_bits}")
        ax.set_xlim(-0.75, +0.75)
        for j, superbit in enumerate([1, 50]):
            for r in range(1, 6):
                cosdist = numpy.loadtxt(f"results/{dataset}.estimator/n_bits-{n_bits}.superbit-{superbit}.{r}.cos.tsv.gz")
                angle = numpy.arccos(-cosdist+1)
                hamdist = numpy.loadtxt(f"results/{dataset}.estimator/n_bits-{n_bits}.superbit-{superbit}.{r}.ham.tsv.gz")
                approx = hamdist * math.pi / n_bits
                values = (approx - angle).ravel()
                print("{:20s}  {:>6d}  {:>8d}  {:>+.4f}  {:>.4f}".format(dataset, n_bits, superbit, numpy.mean(values), numpy.std(values)))
                kde = scipy.stats.gaussian_kde(values)
                x = numpy.linspace(values.min(), values.max(), num=200)
                ax.plot(x, kde(x), lw=1, color=f"C{j}")
    axes[-1].set_xlabel("estimation error [rad]")
    seaborn.despine(fig=fig)
    fig.tight_layout()
    savefig(f"{dataset}-estimator")

# %% [code]
def n_positives(hamdist, cosdist, k):
    hamord = numpy.argsort(hamdist, axis=1)[:,0:k+1]
    cosord = numpy.argsort(cosdist, axis=1)[:,1]
    n = 0
    for i in range(hamord.shape[0]):
        n += numpy.sum(numpy.isin(cosord[i], hamord[i]))
    return n

n_cells = 6
x_max = 0.3  # upper limit of cosine similarity
print("{:20s}  {:>6s}  {:>2s}  {:>11s}".format("dataset", "n-bits", "k", "n-positives"))
for dataset in datasets:
    fig, axes = subplots(4, n_cells, dpi=dpi, figsize=(8, 5), sharex=True, sharey=True)
    for i, n_bits in enumerate([64, 128, 256, 512]):
        cosdist = numpy.loadtxt(f"results/{dataset}.estimator/n_bits-{n_bits}.superbit-50.1.cos.tsv.gz")
        hamdist = numpy.loadtxt(f"results/{dataset}.estimator/n_bits-{n_bits}.superbit-50.1.ham.tsv.gz")
        for k in [10, 30, 50]:
            print("{:20s}  {:>6d}  {:>2d}  {:>11d}".format(dataset, n_bits, k, n_positives(hamdist, cosdist, k)))
        axes[i,0].set_ylabel(f"n-bits = {n_bits}\n\nHamming dist.")
        for j in range(n_cells):
            ax = axes[i,j]
            x = cosdist[j,:]
            y = hamdist[j,:] / n_bits
            ax.scatter(x[x<x_max], y[x<x_max], s=0.3, lw=0, c="C1")
            ax.set_xlim(0, x_max)
            ax.set_ylim(0, None)
    for j in range(n_cells):    
        axes[3,j].set_xlabel("cosine dist.")
    seaborn.despine(fig=fig)
    fig.tight_layout(pad=0.5)
    savefig(f"{dataset}-correlation")

# %% [markdown]
## Visualization of expression profiles

# %% [markdown]
cell types and query/database plots

# %% [code]
def readlines(filename):
    with open(filename) as f:
        return f.read().splitlines()

cluster = clusters["Shekhar2016"]
tab20 = matplotlib.cm.get_cmap("tab20")
for query in ["1", "2"]:
    print(f"query batch: {query}")
    log = toml.load(f"results/Shekhar2016.hash.batch-{query}.toml")
    print(log["date-time"])
    print(log["version-info"])
    outdir = f"results/Shekhar2016.hash.batch-{query}"
    for metric in ["euclidean", "cosine"]:
        fig, axes = subplots(2, 2, dpi=dpi, figsize=(8, 7))
        for ax in axes.ravel():
            ax.xaxis.set_ticklabels([])
            ax.yaxis.set_ticklabels([])
        for i, project in enumerate(["false", "true"]):
            cells_query = readlines(f"{outdir}/cells.query.project-{project}.txt")
            X_query = numpy.loadtxt(f"{outdir}/X.query.project-{project}.tsv").T
            cells_database = readlines(f"{outdir}/cells.database.project-{project}.txt")
            X_database = numpy.loadtxt(f"{outdir}/X.database.project-{project}.tsv").T
            X = numpy.vstack([X_query, X_database])
            X_umap = umap.UMAP(metric=metric, random_state=1234).fit_transform(X)
            shuf = numpy.arange(X_umap.shape[0])
            numpy.random.RandomState(1234).shuffle(shuf)
            labels = cluster[cells_query + cells_database]
            label2color = {label: tab20(k) for k, label in enumerate(numpy.sort(labels.unique()))}
            color = numpy.array([label2color[label] for label in labels])
            axes[i,0].scatter(X_umap[shuf,0], X_umap[shuf,1], s=0.5, lw=0, c=color[shuf])
            color = numpy.array(["C1"] * len(cells_query) + ["C0"] * len(cells_database))
            axes[i,1].scatter(X_umap[shuf,0], X_umap[shuf,1], s=0.5, lw=0, c=color[shuf])
        axes[0,0].set_title("cell type")
        axes[0,1].set_title("query/database")
        axes[0,0].set_ylabel("projected = false")
        axes[1,0].set_ylabel("projected = true")
        #fig.suptitle(f"query = {query}", va="bottom")
        seaborn.despine(fig=fig)
        fig.tight_layout(pad=0.5)
        savefig(f"Shekhar2016-hash-query-{query}-metric-{metric}")

# %% [code]
%load_ext Cython

# %% [code]
%%cython

import numpy
cimport numpy
cimport cython

cdef extern int __builtin_popcount(unsigned int)

@cython.boundscheck(False)
@cython.wraparound(False)
cdef distance_matrix(numpy.ndarray[numpy.uint8_t,ndim=1] rows,
                     numpy.ndarray[numpy.uint8_t,ndim=1] cols,
                     int nbytes,
                     numpy.ndarray[numpy.uint16_t,ndim=2] output):
    cdef int i, j, k
    cdef int d
    cdef int m = rows.shape[0] // nbytes
    cdef int n = cols.shape[0] // nbytes
    for i in range(m):
        for j in range(n):
            d = 0
            for k in range(nbytes):
                d += __builtin_popcount(rows[i*nbytes+k] ^ cols[j*nbytes+k])
            output[i,j] = d

            
def hamming_distance_matrix(rows, cols, nbytes):
    assert rows.shape[0] % nbytes == 0
    assert cols.shape[0] % nbytes == 0
    m = rows.shape[0] // nbytes
    n = cols.shape[0] // nbytes
    output = numpy.zeros((m, n), dtype=numpy.uint16)
    distance_matrix(rows, cols, nbytes, output)
    return output

# %% [code]
nbytes = 64 // 8
cluster = clusters["Shekhar2016"]
tab20 = matplotlib.cm.get_cmap("tab20")
n = len(cluster)
shuf = numpy.arange(n)
numpy.random.RandomState(1234).shuffle(shuf)
for query in ["1", "2"]:
    fig, axes = subplots(2, 2, dpi=dpi, figsize=(8, 8))
    axes = [axes[0,0], axes[0,1], axes[1,0], axes[1,1]]
    for ax in axes:
        ax.xaxis.set_ticklabels([])
        ax.yaxis.set_ticklabels([])
    outdir = f"results/Shekhar2016.hash.batch-{query}"
    cells_query = readlines(f"{outdir}/cells.query.project-true.txt")
    X_query = numpy.loadtxt(f"{outdir}/X.query.project-true.tsv").T
    cells_database = readlines(f"{outdir}/cells.database.project-true.txt")
    X_database = numpy.loadtxt(f"{outdir}/X.database.project-true.tsv").T
    X = numpy.vstack([X_query, X_database])
    X_umap = umap.UMAP(random_state=1234, metric="cosine").fit_transform(X)
    # plot
    labels = cluster[cells_query + cells_database]
    label2color = {label: tab20(k) for k, label in enumerate(numpy.sort(labels.unique()))}
    color = numpy.array([label2color[label] for label in labels])
    ax = axes.pop(0)
    ax.scatter(X_umap[shuf,0], X_umap[shuf,1], s=0.5, lw=0, c=color[shuf])
    ax.set_title("cosine distance (original)")
    # hashed profiles
    D = numpy.zeros((n, n), dtype=numpy.uint16)
    for l in range(4):
        # compute distance matrix
        Z_query = numpy.fromfile(f"{outdir}/Z.query.project-true.{l+1}.bin", dtype=numpy.uint8)
        Z_database = numpy.fromfile(f"{outdir}/Z.database.project-true.{l+1}.bin", dtype=numpy.uint8)
        Z = numpy.hstack([Z_query, Z_database])
        D += hamming_distance_matrix(Z, Z, nbytes)
        nbits = nbytes * (l+1) * 8
        if nbits in (64, 128, 256):
            D_umap = umap.UMAP(random_state=1234, metric="precomputed").fit_transform(D)         
            ax = axes.pop(0)
            ax.scatter(D_umap[shuf,0], D_umap[shuf,1], s=0.5, lw=0, c=color[shuf])
            ax.set_title(f"Hamming distance (n-bits = {nbits})")
    seaborn.despine(fig=fig)
    fig.tight_layout(pad=0.5)
    savefig(f"Shekhar2016-hash-query-{query}")

# %% [markdown]
## Self-mapping experiments

# %% [code]
def load_knn(filename):
    return pandas.read_table(filename, index_col="cell")

# Compute the consistency, Cohen's kappa, and the adjusted Rand scores.
def cluster_metrics(filename, cluster, k=1, cluster_nn=None):
    if cluster_nn is None:
        cluster_nn = cluster
    knn = load_knn(filename)
    cell = cluster[knn.index].values
    nn = cluster_nn[knn[f"N{k}"]].values
    return dict(
        consistency=sklearn.metrics.accuracy_score(cell, nn),
        cohen_kappa=sklearn.metrics.cohen_kappa_score(cell, nn),
        adjusted_rand=sklearn.metrics.adjusted_rand_score(cell, nn),)

# %% [code]
knn_cv = pandas.DataFrame()
knn_cv_scmap = pandas.DataFrame()
for dataset in datasets:
    print(dataset)
    # CellFishing
    log = toml.load(f"results/{dataset}.knn-cv.toml")
    print(log["date-time"])
    print(log["version-info"])
    exp = pandas.DataFrame(log["experiment"])
    exp["dataset"] = dataset
    exp = pandas.concat(
        [exp, pandas.DataFrame([cluster_metrics(x, clusters[dataset], k=1) for x in exp["filename"]])],
        axis=1)
    knn_cv = knn_cv.append(exp, ignore_index=True)
    # scmap
    log = toml.load(f"results/{dataset}.knn-cv.scmap.toml")
    print(log["date-time"])
    print(log["session-info"])
    exp = pandas.DataFrame(log["experiment"])
    exp["dataset"] = dataset
    exp = pandas.concat(
        [exp, pandas.DataFrame([cluster_metrics(x, clusters[dataset], k=1) for x in exp["filename"]])],
        axis=1)
    knn_cv_scmap = knn_cv_scmap.append(exp, ignore_index=True)

# %% [markdown]
Select columns:

# %% [code]
knn_cv = knn_cv[["dataset", "n-bits", "n-lshashes", "transformer", "filename", "consistency", "cohen_kappa", "adjusted_rand", "index-time", "query-time"]]
knn_cv_scmap = knn_cv_scmap[["dataset", "n-features", "n-centroids-factor", "filename", "consistency", "cohen_kappa", "adjusted_rand", "index-time", "query-time"]]

# %% [markdown]
Summary:

# %% [code]
pandas.options.display.max_columns = 99
knn_cv.groupby(["dataset", "n-bits", "n-lshashes", "transformer"]).describe()

# %% [code]
knn_cv_scmap.groupby(["dataset", "n-features", "n-centroids-factor"]).describe()

# %% [markdown]
Compare the elapsed time with the default parameters:

# %% [code]
knn_cv_elapsed = knn_cv[(knn_cv["n-bits"]==128)&(knn_cv["n-lshashes"]==4)&(knn_cv["transformer"]=="log1p")][["dataset", "index-time", "query-time"]].groupby("dataset").median()
knn_cv_elapsed

# %% [code]
knn_cv_scmap_elapsed = knn_cv_scmap[(knn_cv_scmap["n-features"]==500)&(knn_cv_scmap["n-centroids-factor"]==1)][["dataset", "index-time", "query-time"]].groupby("dataset").median()
knn_cv_scmap_elapsed

# %% [code]
knn_cv_scmap_elapsed / knn_cv_elapsed

# %% [markdown]
Plot consistency scores, Cohen's kappa coefficients, index times, and query times:

# %% [code]
metric_labels = {
    "consistency": "consistency",
    "cohen_kappa": "Cohen's kappa",
    "index-time": "index time [s]",
    "query-time": "query time [s]"}
transformer = "log1p"
for metric in ["consistency", "cohen_kappa", "index-time", "query-time"]:
    fig, axes = subplots(len(datasets), 2, dpi=dpi, figsize=(8, 7))
    for i, dataset in enumerate(datasets):
        if i > 0:
            axes[i,0].get_shared_x_axes().join(axes[0,0], axes[i,0])
            axes[i,1].get_shared_x_axes().join(axes[0,1], axes[i,1])
        axes[i,0].get_shared_y_axes().join(axes[i,0], axes[i,1])
        data = knn_cv[(knn_cv["dataset"]==dataset)&(knn_cv["transformer"]==transformer)]
        data_scmap = knn_cv_scmap[knn_cv_scmap["dataset"]==dataset]
        ymin = min(data[metric].min(), data_scmap[metric].min())
        ymax = max(data[metric].max(), data_scmap[metric].max())
        ydiff = ymax - ymin
        markerscale = 0.5
        labelspacing = 0.2
        if metric.endswith("-time"):
            seaborn.swarmplot(x="n-bits", hue="n-lshashes", y=metric, data=data, ax=axes[i,0], size=3, dodge=True)
            seaborn.swarmplot(x="n-centroids-factor", hue="n-features", y=metric, data=data_scmap, ax=axes[i,1], size=3, dodge=True)
            axes[i,0].set_ylim(0, None)
            axes[i,0].legend(title="n-lshashes", fontsize="small", markerscale=markerscale)
            axes[i,1].legend(title="n-features", fontsize="small", markerscale=markerscale, loc="upper left", labelspacing=labelspacing)
        else:
            axes[i,0].set_ylim(ymin - ydiff * 0.02, ymax + ydiff * 0.02)  # need to set ylim before plotting
            seaborn.swarmplot(x="n-bits", hue="n-lshashes", y=metric, data=data, ax=axes[i,0], size=3, dodge=True)
            seaborn.swarmplot(x="n-centroids-factor", hue="n-features", y=metric, data=data_scmap, ax=axes[i,1], size=3, dodge=True)
            axes[i,0].legend(title="n-lshashes", fontsize="small", markerscale=markerscale, loc="lower right")
            axes[i,1].legend(title="n-features", fontsize="small", markerscale=markerscale, labelspacing=labelspacing)
        if i > 0:
            axes[i,0].legend_.remove()
            axes[i,1].legend_.remove()
        axes[i,0].set_ylabel(dataset + "\n\n" + metric_labels[metric])
        axes[i,1].set_ylabel("")
        axes[i,1].set_yticklabels([])
        for j in range(2):
            axes[i,j].locator_params(axis="y", nbins=4)
            axes[i,j].grid(axis="y", alpha=0.3)
        if i < len(datasets) - 1:
            for j in range(2):
                axes[i,j].set_xlabel("")
                axes[i,j].set_xticklabels([])
    axes[0,0].set_title("CellFishing")
    axes[0,1].set_title("scmap-cell")
    seaborn.despine(fig=fig)
    fig.tight_layout(h_pad=0.2)
    savefig(f"selfmapping-{metric}")

# %% [markdown]
Summarize the adjusted Rand scores from SC3.

# %% [code]
data = []
for dataset in datasets:
    # The TOML files were broken due to the noisy outputs from SC3.
    cluster = clusters[dataset]
    k = len(cluster.unique())
    for r in range(5):
        sc3 = pandas.read_table(f"results/{dataset}.clustering-sc3/k-{k}.{r+1}.tsv.gz", index_col="cell")["cluster"]
        assert sc3.index.equals(cluster.index)
        score = sklearn.metrics.adjusted_rand_score(cluster, sc3)
        data.append((dataset, score))
sc3 = pandas.DataFrame(data, columns=["dataset", "adjusted_rand"])
sc3.groupby("dataset").describe()

# %% [markdown]
## Feature selection

# %% [markdown]
Compare default features:

# %% [code]
def readlines(filename):
    with open(filename) as file:
        return [line.rstrip() for line in file.readlines()]

fig, axes = subplots(2, 2, dpi=dpi)
for i, dataset in enumerate(datasets):
    features = readlines(f"results/{dataset}.select-features.n_features-default.txt")
    features_scmap = readlines(f"results/{dataset}.select-features.scmap.n_features-500.txt")
    ax = axes[i//2,i%2]
    matplotlib_venn.venn2([set(features), set(features_scmap)], set_labels=("", ""), ax=ax)
    ax.set_title(dataset)

# %% [code]
for dataset in datasets:
    fig, axes = subplots(2, 2, dpi=dpi)
    for i, n_features in enumerate([500, 1000, 2000, 4000]):
        features = readlines(f"results/{dataset}.select-features.n_features-{n_features}.txt")
        features_scmap = readlines(f"results/{dataset}.select-features.scmap.n_features-{n_features}.txt")
        ax = axes[i//2,i%2]
        matplotlib_venn.venn2([set(features), set(features_scmap)], set_labels=("", ""), ax=ax)
        ax.set_title(f"n-features = {n_features}")
    fig.suptitle(dataset)

# %% [code]
knn_cv_features = pandas.DataFrame({"dataset": [], "consistency": [], "cohen_kappa": []})
for dataset in datasets:
    for n_features in [500, 1000, 2000, 4000]:
        filename = f"results/{dataset}.knn-cv-features.n_features-{n_features}.toml"
        out = toml.load(filename)
        for exp in out["experiment"]:
            row = cluster_metrics(exp["filename"], clusters[dataset])
            row["dataset"] = dataset
            row["n-features"] = n_features
            knn_cv_features = knn_cv_features.append(row, ignore_index=True)

# %% [code]
knn_cv_features.groupby(["dataset", "n-features"]).describe()

# %% [code]
knn_cv_features_scmap = pandas.DataFrame({"dataset": [], "consistency": [], "cohen_kappa": []})
for dataset in datasets:
    for n_min_features in ["default", 500, 1000, 2000, 4000]:
        filename = f"results/{dataset}.knn-cv-features.scmap.n_features-{n_min_features}.toml"
        out = toml.load(filename)
        for exp in out["experiment"]:
            row = cluster_metrics(exp["filename"], clusters[dataset])
            row["dataset"] = dataset
            row["n-min-features"] = n_min_features
            with open(exp["features-file"]) as f:
                row["n-features"] = len(f.readlines())
            knn_cv_features_scmap = knn_cv_features_scmap.append(row, ignore_index=True)

# %% [code]
knn_cv_features_scmap.groupby(["dataset", "n-min-features"]).describe()

# %% [markdown]
## Cluster-specific scores

# %% [code]
import collections

def count_assignments(cell, nn, cluster):
    consistent = cluster[cell].values == cluster[nn].values
    cluster = cluster[cell]
    # make counters
    count_all = collections.Counter(cluster)
    count_consistent = collections.Counter({c: 0 for c in count_all.keys()})
    count_consistent.update(cluster[consistent])
    count_inconsistent = collections.Counter({c: 0 for c in count_all.keys()})
    count_inconsistent.update(cluster[~consistent])
    # join counts
    counts = pandas.DataFrame({"all": pandas.Series(count_all), "consistent": pandas.Series(count_consistent), "inconsistent": pandas.Series(count_inconsistent)})
    counts = counts.assign(consistency=lambda x: x["consistent"] / x["all"])[["all", "consistent", "inconsistent", "consistency"]]
    counts.sort_values(by="all", ascending=False, inplace=True)
    counts.index.name = "cluster"
    return counts


def plot_assignments_matrix(cell, nn, cluster, cluster2=None, overlapping=False, figsize=(12,12)):
    cluster1 = cluster
    if cluster2 is None:
        cluster2 = cluster1
    labels1 = cluster1.value_counts().index.values
    labels2 = cluster2.value_counts().index.values
    if overlapping:
        labels2 = labels1
    assignments = pandas.crosstab(
        pandas.Categorical(cluster1[cell].values, categories=labels1),
        pandas.Categorical(cluster2[nn].values, categories=labels2),
        rownames=["query"],
        colnames=["neighbor"],
        dropna=False,
        normalize="index")
    assignments = assignments.loc[labels1,labels2]
    fig, ax = subplots(dpi=dpi, figsize=figsize)
    im = ax.imshow(assignments, vmin=0, vmax=1)
    ax.set_ylabel("Query's label")
    ax.set_xlabel("Neighbor's label")
    ax.set_yticks(numpy.arange(assignments.shape[0]))
    ax.set_xticks(numpy.arange(assignments.shape[1]))
    ax.set_yticklabels(assignments.index)
    ax.set_xticklabels(assignments.columns, rotation=90)
    ax.set_yticks(numpy.arange(assignments.shape[0]+1)-.5, minor=True)
    ax.set_xticks(numpy.arange(assignments.shape[1]+1)-.5, minor=True)
    ax.grid(False)  # turn off seaborn's grid
    ax.grid(which="minor", color="gray", linestyle='-', linewidth=0.8)
    ax.tick_params(which="minor", bottom=False, left=False)
    ax.tick_params(length=0)
    cbar = fig.colorbar(im, fraction=0.046, pad=0.02, shrink=0.3, aspect=30)
    cbar.outline.set_visible(False)
    for x in ["left", "right", "top", "bottom"]:
        ax.spines[x].set_visible(False)
    fig.tight_layout()
    return fig, ax

# %% [code]
# wrap long labels
def wraplabel(label, width=16):
    return "\n".join(textwrap.wrap(label.replace("_", " "), width=width))

# %% [code]
for dataset in datasets:
    print(dataset)
    cluster = clusters[dataset]
    # CellFishing
    filenames = knn_cv[(knn_cv["dataset"]==dataset)&(knn_cv["n-bits"]==128)&(knn_cv["n-lshashes"]==4)&(knn_cv["transformer"]=="log1p")]["filename"]
    knn = [load_knn(x) for x in filenames]
    counts1 = [count_assignments(x.index, x["N1"], cluster) for x in knn]
    fig, ax = plot_assignments_matrix(knn[0].index, knn[0]["N1"], cluster)
    savefig(f"{dataset}-selfmapping-matrix")
    # scmap
    filenames = knn_cv_scmap[(knn_cv_scmap["dataset"]==dataset)&(knn_cv_scmap["n-features"]==500)&(knn_cv_scmap["n-centroids-factor"]==1)]["filename"]
    knn = [load_knn(x) for x in filenames]
    counts2 = [count_assignments(x.index, x["N1"], cluster) for x in knn]
    fig, ax = plot_assignments_matrix(knn[0].index, knn[0]["N1"], cluster)
    savefig(f"{dataset}-selfmapping-matrix-scmap")
    # merge
    x = pandas.concat([x["consistency"].reset_index() for x in counts1], ignore_index=True)
    x["method"] = "CellFishing"
    y = pandas.concat([x["consistency"].reset_index() for x in counts2], ignore_index=True)
    y["method"] = "scmap-cell";
    data = pandas.concat([x, y], ignore_index=True)
    cluster_unique = cluster.value_counts().index.values
    n_cols = 3
    y_max = math.ceil(len(cluster_unique) / n_cols)
    fig, axes = subplots(1, n_cols, dpi=dpi, figsize=(8, math.ceil(y_max / 2)))
    for j in range(n_cols):
        if n_cols == 1:
            ax = axes
        else:
            ax = axes[j]
        ax.set_xlim(0, 1.02)
        ax.set_xticks([0, 0.2, 0.4, 0.6, 0.8, 1.0])
        y = cluster_unique[j*y_max:min((j+1)*y_max, len(cluster_unique))]
        seaborn.swarmplot(x="consistency", y="cluster", hue="method", data=data[data["cluster"].isin(y)], order=y, orient="h", dodge=False, size=2.4, ax=ax)
        ax.tick_params(labelsize="small")
        ax.set_ylabel("")
        ax.set_yticklabels([wraplabel(x.get_text()) for x in ax.get_yticklabels()])
        ax.grid(axis="x", alpha=0.3)
        if j > 0:
            ax.legend_.remove()
        else:
            ax.legend(loc="upper left", fontsize="small").set_title("")
    fig.tight_layout(w_pad=0.5)
    seaborn.despine(fig=fig)
    savefig(f"{dataset}-selfmapping-clusters")

# %% [markdown]
## Similarities from nearest neighbors

# %% [code]
def plot_similarities(dataset, cluster, targets):
    fig, axes = subplots(1, len(targets), dpi=dpi, figsize=(8, 4), sharey=True)
    for i, target in enumerate(targets):
        target_escaped = target.replace('/', ';')
        log = toml.load(f"results/{dataset}.knn-similarities.target-{target_escaped}.toml")
        similarities_with_target_cells = pandas.read_table(log["experiment"][0]["filename-similarities-with-target-cells"], index_col="cell")
        similarities_without_target_cells = pandas.read_table(log["experiment"][0]["filename-similarities-without-target-cells"], index_col="cell")
        cells_target = cluster[(cluster == target).values].index.values
        #cells_nontarget = cluster[(cluster != target).values].index.values
        # melt data
        df1 = similarities_with_target_cells.loc[cells_target].reset_index().melt(id_vars="cell", var_name="rank", value_name="similarity")
        df1["removed"] = False
        df2 = similarities_without_target_cells.loc[cells_target].reset_index().melt(id_vars="cell", var_name="rank", value_name="similarity")
        df2["removed"] = True
        df = df1.append(df2)
        df["rank"] = [int(x[1]) for x in df["rank"]]
        df = df[df["rank"]<=3]
        # plot
        ax = axes[i]
        seaborn.violinplot(x="rank", y="similarity", hue="removed", data=df, split=True, inner="quart", linewidth=0.6, ax=ax)
        #seaborn.boxplot(x="rank", y="similarity", hue="removed", data=df, linewidth=0.8, fliersize=0.5, ax=ax)
        ax.set_title(wraplabel(target), fontsize="small")
        ax.legend(loc="lower left")
        ax.legend_.remove()
        ax.grid(axis="y", alpha=0.3)
        if i > 0:
            ax.set_ylabel("")
    axes[0].legend(title="removed", loc="lower left", fontsize="small", title_fontsize="small")
    seaborn.despine(fig=fig)
    fig.tight_layout()
    savefig(f"{dataset}-selfmapping-remove")


def list_nearest_neighbors(dataset, cluster, targets):
    for target in targets:
        target_escaped = target.replace('/', ';')
        log = toml.load(f"results/{dataset}.knn-similarities.target-{target_escaped}.toml")
        #knn_with_target_cells = pandas.read_table(log["experiment"][0]["filename-knn-with-target-cells"], index_col="cell")
        knn_without_target_cells = pandas.read_table(log["experiment"][0]["filename-knn-without-target-cells"], index_col="cell")
        cells_target = cluster[(cluster == target).values].index.values
        value_counts = cluster.loc[knn_without_target_cells.loc[cells_target,"N1"]].value_counts()
        print(target)
        for i, (label, n) in enumerate(value_counts.items()):
            if i > 4:
                break
            print("    {}: {}".format(label, n))

# %% [code]
dataset = "Shekhar2016"
targets = ["BC2", "BC5D", "BC3A", "BC5B", "BC4", "BC8/9 (mixture of BC8 and BC9)", "AC (Amacrine cell)"]
plot_similarities(dataset, clusters[dataset], targets)

# %% [code]
list_nearest_neighbors(dataset, clusters[dataset], targets)

# %% [code]
dataset = "TabulaMuris-chromium"
targets = ["basal cell", "alveolar macrophage", "fibroblast", "immature B cell", "basophil", "kidney cell", "endothelial cell of hepatic sinusoid"]
plot_similarities(dataset, clusters[dataset], targets)

# %% [code]
list_nearest_neighbors(dataset, clusters[dataset], targets)

# %% [markdown]
## Detecting DEGs

# %% [code]
def load_degs(cluster, target, k):
    knns = pandas.read_table(f"results/TabulaMuris-chromium.knn-degs.target-{target}/knn.k-{k}.tsv.gz", index_col="cell")
    degs = pandas.read_table(f"results/TabulaMuris-chromium.knn-degs.target-{target}/degs.k-{k}.tsv.gz", index_col=["cell", "gene"])
    return degs.join(cluster).join(knns["N1"]).join(cluster.rename_axis("N1"), on="N1", rsuffix="N1").drop("N1", axis=1)


def top_genes(by, n=10000, upper=-4):
    def fetch(df):
        x = df.sort_values(by).head(n)
        return x[x[by] <= upper].reset_index("cell", drop=True)
    return fetch


def make_crosstable(cluster, celltypes, target, k, by):
    degs = load_degs(cluster, target, k)
    x = degs.groupby("cell").apply(top_genes(by)).reset_index("gene")
    return pandas.crosstab(x["gene"], x["clusterN1"]).loc[:,celltypes]

# %% [code]
celltype_query = "immature B cell"
celltypes = ['early pro-B cell', 'late pro-B cell', 'B cell',]
cluster = clusters["TabulaMuris-chromium"]

# %% [code]
load_degs(cluster, celltype_query, 5).groupby("cell").first()["clusterN1"].value_counts()

# %% [markdown]
List top DEGs.

# %% [code]
for celltype in celltypes:
    print(celltype)
    print("-"*len(celltype))
    for k in [5, 10, 20]:
        print(f"k = {k}")
        negative = make_crosstable(cluster, celltypes, celltype_query, k, "negative")
        positive = make_crosstable(cluster, celltypes, celltype_query, k, "positive")
        print("  negative: ", end="")
        degs = negative.sort_values(celltype, ascending=False)[celltype]
        for i in range(10):
            if degs.iloc[i] == 0:
                break
            print("{:s} {:d}".format(degs.index[i], degs.iloc[i]), end=", ")
        print()
        print("  positive: ", end="")
        degs = positive.sort_values(celltype, ascending=False)[celltype]
        for i in range(10):
            if degs.iloc[i] == 0:
                break
            print("{:s} {:d}".format(degs.index[i], degs.iloc[i]), end=", ")
        print()
    print()

# %% [markdown]
Generate LaTeX tables.

# %% [code]
n = 6  # the number of top DEGs

def capitalize(s):
    return s[0].upper() + s[1:]

for k in [5, 10, 20]:
    negative = make_crosstable(cluster, celltypes, celltype_query, k, "negative")
    positive = make_crosstable(cluster, celltypes, celltype_query, k, "positive")
    print(f"% k = {k}")
    print(r"\begin{tabular}{" + "lr" * (len(celltypes) * 2) + "}")
    print(r"\toprule")

    for j, celltype in enumerate(celltypes):
        print(r"\multicolumn{4}{c}{", capitalize(celltype), r"}", end=" & " if j < len(celltypes)-1 else r" \\")
    print()
    for j in range(len(celltypes)):
        print(r"\cmidrule(lr){", 4 * j + 1, "-", 4 * j + 4, r"}", end=" ")
    print()

    for j in range(len(celltypes)):
        print(r"\multicolumn{2}{c}{Negative} & \multicolumn{2}{c}{Positive}", end=" & " if j < len(celltypes)-1 else r" \\")
    print()
    for j in range(len(celltypes)):
        print(r"\cmidrule(lr){", 4 * j + 1, "-", 4 * j + 2, r"}", end=" ")
        print(r"\cmidrule(lr){", 4 * j + 3, "-", 4 * j + 4, r"}", end=" ")
    print()

    # Gene count
    for i in range(n):
        for j, celltype in enumerate(celltypes):
            df = negative.sort_values(celltype)
            row = df.iloc[-(i+1)]
            print("\\textit{{{:10s}}} & {:3d}".format(row.name if row[celltype] > 0 else "--", row[celltype]), end=" & ")
            df = positive.sort_values(celltype)
            row = df.iloc[-(i+1)]
            print("\\textit{{{:10s}}} & {:3d}".format(row.name if row[celltype] > 0 else "--", row[celltype]), end=" & " if j < len(celltypes)-1 else r" \\")
        print()

    # Total count
    print(r"\addlinespace")
    for j, celltype in enumerate(celltypes):
        print("Total & {:3d}".format(negative[celltype].sum()), end=" & ")
        print("Total & {:3d}".format(positive[celltype].sum()), end=" & " if j < len(celltypes)-1 else r" \\")
    
    print()
    print(r"\bottomrule")
    print(r"\end{tabular}")
    print()

# %% [markdown]
## Mapping across batches

# %% [code]
knn_batch_shekhar2016 = pandas.DataFrame()
knn_batch_shekhar2016_scmap = pandas.DataFrame()
cluster = clusters["Shekhar2016"]
for query in ["1", "2"]:
    print(f"query = {query}")
    # CellFishing
    log = toml.load(f"results/Shekhar2016.knn-batch.batch-{query}.toml")
    print(log["date-time"])
    print(log["version-info"])
    exp = pandas.DataFrame(log["experiment"])
    exp["query"] = query
    exp = pandas.concat(
        [exp,
         pandas.DataFrame([cluster_metrics(x, cluster) for x in exp["filename"]])],
        axis=1)
    knn_batch_shekhar2016 = knn_batch_shekhar2016.append(exp, ignore_index=True)
    # scmap
    log = toml.load(f"results/Shekhar2016.knn-batch.scmap.batch-{query}.toml")
    print(log["date-time"])
    print(log["session-info"])
    exp = pandas.DataFrame(log["experiment"])
    exp["query"] = query
    exp = pandas.concat(
        [exp,
         pandas.DataFrame([cluster_metrics(x, cluster) for x in exp["filename"]])],
        axis=1)
    knn_batch_shekhar2016_scmap = knn_batch_shekhar2016_scmap.append(exp, ignore_index=True)

# %% [code]
knn_batch_plass2018 = pandas.DataFrame()
knn_batch_plass2018_scmap = pandas.DataFrame()
cluster = clusters["Plass2018"]
for query in ["plan1", "plan2"]:
    print(f"query = {query}")
    # CellScape
    log = toml.load(f"results/Plass2018.knn-batch.batch-{query}.toml")
    print(log["date-time"])
    print(log["version-info"])
    exp = pandas.DataFrame(log["experiment"])
    exp["query"] = query
    exp = pandas.concat(
        [exp,
         pandas.DataFrame([cluster_metrics(x, cluster) for x in exp["filename"]])],
        axis=1)
    knn_batch_plass2018 = knn_batch_plass2018.append(exp, ignore_index=True)
    # scmap
    log = toml.load(f"results/Plass2018.knn-batch.scmap.batch-{query}.toml")
    print(log["date-time"])
    print(log["session-info"])
    exp = pandas.DataFrame(log["experiment"])
    exp["query"] = query
    exp = pandas.concat(
        [exp,
         pandas.DataFrame([cluster_metrics(x, cluster) for x in exp["filename"]])],
        axis=1)
    knn_batch_plass2018_scmap = knn_batch_plass2018_scmap.append(exp, ignore_index=True)

# %% [code]
knn_batch_shekhar2016 = knn_batch_shekhar2016[["query", "n-bits", "n-lshashes", "transformer", "filename", "consistency", "cohen_kappa"]]
knn_batch_shekhar2016_scmap = knn_batch_shekhar2016_scmap[["query", "n-features", "n-centroids-factor", "filename", "consistency", "cohen_kappa"]]

# %% [code]
knn_batch_plass2018 = knn_batch_plass2018[["query", "n-bits", "n-lshashes", "transformer", "filename", "consistency", "cohen_kappa"]]
knn_batch_plass2018_scmap = knn_batch_plass2018_scmap[["query", "n-features", "n-centroids-factor", "filename", "consistency", "cohen_kappa"]]

# %% [code]
knn_batch_shekhar2016.groupby(["query", "n-bits", "n-lshashes", "transformer"]).describe()

# %% [code]
knn_batch_shekhar2016_scmap.groupby(["query", "n-features", "n-centroids-factor"]).describe()

# %% [code]
knn_batch_plass2018.groupby(["query", "n-bits", "n-lshashes", "transformer"]).describe()

# %% [code]
knn_batch_plass2018_scmap.groupby(["query", "n-features", "n-centroids-factor"]).describe()

# %% [code]
n_bits = 128
n_lshashes = 4
transformer = "log1p"
n_features = 500
n_centroids_factor = 1
for metric in ["consistency", "cohen_kappa"]:
    fig, axes = subplots(2, 2, dpi=dpi, figsize=(8, 6))
    for i, (dataset, data, data_scmap) in enumerate([
            ("Shekhar2016", knn_batch_shekhar2016, knn_batch_shekhar2016_scmap),
            ("Plass2018", knn_batch_plass2018, knn_batch_plass2018_scmap)]):
        if i > 0:
            axes[i,0].get_shared_x_axes().join(axes[0,0], axes[i,0])
            axes[i,1].get_shared_x_axes().join(axes[0,1], axes[i,1])
        axes[i,0].get_shared_y_axes().join(axes[i,0], axes[i,1])
        data = data[(data["n-bits"]==n_bits)&(data["n-lshashes"]==n_lshashes)&(data["transformer"]==transformer)]
        mean = knn_cv[(knn_cv["dataset"]==dataset)&(knn_cv["n-bits"]==n_bits)&(knn_cv["n-lshashes"]==n_lshashes)&(knn_cv["transformer"]==transformer)][metric].mean()
        data_scmap = data_scmap[(data_scmap["n-features"]==n_features)&(data_scmap["n-centroids-factor"]==n_centroids_factor)]
        mean_scmap = knn_cv_scmap[(knn_cv_scmap["dataset"]==dataset)&(knn_cv_scmap["n-features"]==n_features)&(knn_cv_scmap["n-centroids-factor"]==n_centroids_factor)][metric].mean()
        ymin = min(data[metric].min(), data_scmap[metric].min())
        ymax = max(data[metric].max(), data_scmap[metric].max())
        ydiff = ymax - ymin
        axes[i,0].set_ylim(ymin - ydiff * 0.02, ymax + ydiff * 0.02)  # need to set ylim before plotting
        seaborn.swarmplot(x="query", y=metric, data=data, ax=axes[i,0])
        axes[i,0].axhline(mean, color="orangered", lw=0.5)
        seaborn.swarmplot(x="query", y=metric, data=data_scmap, ax=axes[i,1])
        axes[i,1].axhline(mean_scmap, color="orangered", lw=0.5)
        axes[i,0].set_ylabel(dataset + "\n\n" + metric_labels[metric])
        axes[i,1].set_ylabel("")
        axes[i,1].set_yticklabels([])
        for j in range(2):
            axes[i,j].locator_params(axis="y", nbins=4)
            axes[i,j].grid(axis="y", alpha=0.3)
    axes[0,0].set_title("CellFishing")
    axes[0,1].set_title("scmap-cell")
    seaborn.despine(fig=fig)
    fig.tight_layout()
    savefig(f"mapping-{metric}")

# %% [code]
batch1 = pandas.read_table("data/Shekhar2016.batch-1.txt", names=["cell"])["cell"]
batch2 = pandas.read_table("data/Shekhar2016.batch-2.txt", names=["cell"])["cell"]
cluster = clusters["Shekhar2016"]
fig, ax = subplots(dpi=dpi, figsize=(8, 8))
seaborn.countplot(y="cluster", hue="batch", data=pandas.concat([
    pandas.DataFrame(cluster[batch1]).assign(batch=1),
    pandas.DataFrame(cluster[batch2]).assign(batch=2),
]), order=cluster.value_counts().index.values, log=False, ax=ax)
ax.set_xlabel("cluster size")
seaborn.despine(fig=fig)
fig.tight_layout()
savefig(f"Shekhar2016-clusters-batches")

# %% [code]
batch1 = pandas.read_table("data/Plass2018.batch-plan1.txt", names=["cell"])["cell"]
batch2 = pandas.read_table("data/Plass2018.batch-plan2.txt", names=["cell"])["cell"]
cluster = clusters["Plass2018"]
fig, ax = subplots(dpi=dpi, figsize=(8, 8))
seaborn.countplot(y="cluster", hue="batch", data=pandas.concat([
    pandas.DataFrame(cluster[batch1]).assign(batch="plan1"),
    pandas.DataFrame(cluster[batch2]).assign(batch="plan2"),
]), order=cluster.value_counts().index.values, log=False, ax=ax)
ax.set_xlabel("cluster size")
seaborn.despine(fig=fig)
fig.tight_layout()
savefig(f"Plass2018-clusters-batches")

# %% [markdown]
## Mapping across species

# %% [code]
knn_species = toml.load("results/Baron2016.knn-species.toml")
print(knn_species["date-time"])
print(knn_species["version-info"])
knn_species = pandas.DataFrame(knn_species["experiment"])

# %% [code]
items = []
for row in knn_species.itertuples():
    if row.query == "mouse":
        query = "Baron2016-mouse"
        reference = "Baron2016-human"
    else:
        query = "Baron2016-human"
        reference = "Baron2016-mouse"
    items.append(cluster_metrics(row.filename, clusters[query], cluster_nn=clusters[reference]))
knn_species = knn_species.join(pandas.DataFrame(items))
knn_species

# %% [code]
knn_species_scmap = toml.load("results/Baron2016.knn-species.scmap.toml")
print(knn_species_scmap["date-time"])
print(knn_species_scmap["session-info"])
knn_species_scmap = pandas.DataFrame(knn_species_scmap["experiment"])

# %% [code]
items = []
for row in knn_species_scmap.itertuples():
    if row.query == "mouse":
        query = "Baron2016-mouse"
        reference = "Baron2016-human"
    else:
        query = "Baron2016-human"
        reference = "Baron2016-mouse"
    items.append(cluster_metrics(row.filename, clusters[query], cluster_nn=clusters[reference]))
knn_species_scmap = knn_species_scmap.join(pandas.DataFrame(items))
knn_species_scmap

# %% [code]
knn_species.groupby(["query", "infer-stats"]).describe()[["consistency","cohen_kappa"]]

# %% [code]
knn_species_scmap.groupby(["query"]).describe()[["consistency","cohen_kappa"]]

# %% [markdown]
## Mapping across protocols

# %% [code]
len(clusters["TabulaMuris-chromium"].unique())

# %% [code]
len(clusters["TabulaMuris-smart"].unique())

# %% [code]
overlappings = set(clusters["TabulaMuris-chromium"].unique()) & set(clusters["TabulaMuris-smart"].unique())
len(overlappings)

# %% [code]
nns = load_knn("results/TabulaMuris.knn-protocol/n_nns-10.1.tsv.gz")
plot_assignments_matrix(nns.index.values, nns["N1"].values, clusters["TabulaMuris-smart"], clusters["TabulaMuris-chromium"], figsize=(12, 14))
savefig("TabulaMuris-smart-chromium-matrix")

# %% [code]
plot_assignments_matrix(nns.index.values, nns["N1"].values,
                        clusters["TabulaMuris-smart"][clusters["TabulaMuris-smart"].isin(overlappings)],
                        clusters["TabulaMuris-chromium"][clusters["TabulaMuris-chromium"].isin(overlappings)],
                        overlapping=True, figsize=(12, 14))
savefig("TabulaMuris-smart-chromium-matrix-overlapping")

# %% [code]
nns = load_knn("results/TabulaMuris.knn-protocol.scmap/n_nns-10.1.tsv.gz")
plot_assignments_matrix(nns.index.values, nns["N1"].values, clusters["TabulaMuris-smart"], clusters["TabulaMuris-chromium"], figsize=(12, 14))
savefig("TabulaMuris-smart-chromium-matrix-scmap")

# %% [code]
plot_assignments_matrix(nns.index.values, nns["N1"].values,
                        clusters["TabulaMuris-smart"][clusters["TabulaMuris-smart"].isin(overlappings)],
                        clusters["TabulaMuris-chromium"][clusters["TabulaMuris-chromium"].isin(overlappings)],
                        overlapping=True, figsize=(12, 14))
savefig("TabulaMuris-smart-chromium-matrix-scmap-overlapping")

# %% [markdown]
## Benachmarks of saving and loading

# %% [code]
saveload = pandas.DataFrame()
for dataset in datasets:
    out = toml.load(f"results/{dataset}.save-load.toml")
    print(out["date-time"])
    print(out["version-info"])
    exp = pandas.DataFrame(out["experiment"])
    exp["dataset"] = dataset
    for x in exp.columns:
        if x.endswith("-size"):
            exp[x] /= 1024**2  # B => MiB
        if x.endswith("-time"):
            exp[x] *= 1000  # s => ms
    saveload = saveload.append(exp)

# %% [code]
saveload_median = saveload\
    .groupby(["dataset", "n-bits", "keep-counts"])\
    .median()\
    [["n-cells", "n-genes",
      "ram-size", "file-size",
      "save-time", "load-time",]]
saveload_median

# %% [markdown]
Generate a LaTeX table.

# %% [code]
print(r"""
\begin{tabular}{l cc rrrr}
\toprule
Data set & \#bits & Raw counts & \multicolumn{2}{c}{Size [MiB]} & \multicolumn{2}{c}{Time [ms]} \\
\cmidrule(lr){4-5} \cmidrule(lr){6-7}
         &        &            & Memory & File                    & Save & Load \\""")
for i, dataset in enumerate(datasets):
    if i == 0:
        print(r"\midrule")
    else:
        print(r"\addlinespace")
    print(r"\multirow{4}{*}{", dataset, "}")
    for n_bits in [128, 256]:
        print(r"  & \multirow{2}{*}{", n_bits, "} ", end="")
        for keep_counts in [False, True]:
            if keep_counts:
                print("  &                        ", end="")
            print(" &", r"\texttt{+}" if keep_counts else r"\texttt{-}", end="")
            for col in ["ram-size", "file-size", "save-time", "load-time"]:
                row = saveload_median.loc[(dataset, n_bits, keep_counts)]
                print(" & {:4.1f} ".format(row[col]), end="")
            print(r"\\")
print(r"""\bottomrule
\end{tabular}
""")

# %% [markdown]
## Scalability

# %% [code]
dataset = "1M_neurons"

# %% [code]
out = toml.load(f"results/{dataset}.scalability.toml")
print(out["date-time"])
print(out["version-info"])
scalability = pandas.DataFrame(out["experiment"])
scalability["ram-size"] /= 1024**2  # byte => MiB
scalability = pandas.concat(
    [scalability, pandas.DataFrame([cluster_metrics(x, clusters[dataset], k=1) for x in scalability["filename"]])],
    axis=1)
scalability = scalability.drop(["n-cells-query", "n-genes", "n-nns"], axis=1)  # drop constant columns

# %% [code]
scalability.groupby(["n-bits", "n-cells-database", "indexed"]).median()

# %% [code]
fig, axes = subplots(4, 2, dpi=dpi, figsize=(8, 6), sharex=True)
for i, metric in enumerate(["index-time", "query-time", "ram-size", "consistency"]):
    for j, n_bits in enumerate([128, 256]):
        ax = axes[i,j]
        for indexed in [False, True]:
            data = scalability[(scalability["n-bits"]==n_bits)&(scalability["indexed"]==indexed)].groupby("n-cells-database").median().reset_index()
            ax.plot(data["n-cells-database"], data[metric], ".-", label="index" if indexed else "linear")
        ax.set_xscale("log", basex=2)
        if metric != "consistency":
            ax.set_yscale("log")
        if i == 0:
            ax.set_title(f"n-bits = {n_bits}")
        if i == 3:
            ax.set_xlabel("database size")
        if j == 0:
            if metric == "index-time":
                ax.set_ylabel("index time [s]")
            elif metric == "query-time":
                ax.set_ylabel("query time [s]")
            elif metric == "ram-size":
                ax.set_ylabel("memory size [MiB]")
            else:
                ax.set_ylabel("consistency")
        if i == 0 and j == 0:
            ax.legend()
seaborn.despine(fig=fig)
fig.tight_layout()
savefig("scalability-metrics")

# %% [markdown]
Generate a LaTeX table.

# %% [code]
scalability_median = scalability.groupby(["n-bits", "n-cells-database", "indexed"]).median()
indexed = True
print(r"""
\begin{tabular}{l cc rrrrrrrr}
\toprule
& \#bits  & Index & \multicolumn{8}{c}{Database size $N$}\\
\cmidrule{4-11}
&         &       & $2^{13}$ (1.0) & $2^{14}$ (2.0) & $2^{15}$ (4.0) & $2^{16}$ (8.0) & $2^{17}$ (16.0) & $2^{18}$ (32.0) & $2^{19}$ (64.0) & $2^{20}$ (128.0)\\
""")
scales = numpy.arange(13, 21)
for i, metric in enumerate(["index-time", "query-time", "ram-size"]):
    if i == 0:
        print(r"\midrule")
    else:
        print(r"\addlinespace")
    print(r"\multirow{4}{*}{", end="")
    if metric == "index-time":
        print("Index time [s]", end="")
    elif metric == "query-time":
        print("Query time [s]", end="")
    else:
        print("Memory size [MiB]", end="")
    print(r"}")
    for n_bits in [128, 256]:
        for indexed in [False, True]:
            print(r"  &", n_bits, "&", r"\texttt{+}" if indexed else r"\texttt{-}", end="")
            base = scalability_median[metric].loc[n_bits,2**scales[0],indexed]
            for s in scales:
                n = 2**s
                val = scalability_median[metric].loc[n_bits,n,indexed]
                print(" & {:5.1f} ({:.1f})".format(val, val/base), end="")
            print(r"\\")
print(r"\bottomrule")
print(r"\end{tabular}")

# %% [code]
n_cells = 10000
n_bits = 128
fig, axes = subplots(2, dpi=dpi, figsize=(8, 5))
for i, indexed in enumerate([False, True]):
    ax = axes[i]
    labels = []
    y = 0
    for s in scales:
        n = 2**s
        left = 0
        bars = []
        for j, metric in enumerate(["preproc-time", "search-time", "rank-time"]):
            time = scalability_median[metric].loc[n_bits,n,indexed] / 1000 / n_cells
            bars.append(ax.barh(y, time, left=left, color=f"C{j}"))
            left += time
        labels.append(f"$2^{{{s}}}$")
        y += 1
    ax.set_yticks(range(len(labels)))
    ax.set_yticklabels(labels)
    ax.set_ylabel("database size")
    ax.set_xlabel("elapsed time [μs/cell]")
    ax.legend(bars, ["preprocessing", "searching", "ranking"], title="phase")
    ax.invert_yaxis()
    if indexed:
        ax.set_title("index search")
    else:
        ax.set_title("linear search")
seaborn.despine(fig=fig)
fig.tight_layout();
savefig("scalability-computational-cost")
```
