## Supplementary material for "CellFishing.jl: an ultrafast and scalable cell search method for single-cell RNA-sequencing": Dockerfile

```
FROM bioconductor/release_base2:R3.5.0_Bioc3.7

WORKDIR /workspace

RUN apt update && apt -y install libhdf5-dev
RUN Rscript -e 'install.packages(c("h5", "matrixStats", "readr")); source("https://bioconductor.org/biocLite.R"); biocLite(c("scmap", "scater", "SC3", "SingleCellExperiment"))'

RUN useradd -U agata
USER agata

CMD ["Rscript"]
```
