## Supplementary material for "CellFishing.jl: an ultrafast and scalable cell search method for single-cell RNA-sequencing": Project.toml

```
[deps]
CellFishing = "5ab3512e-c64d-48f6-b1c0-509c1121fdda"
Dates = "ade2ca70-3891-5945-98fb-dc099432e06a"
DelimitedFiles = "8bb1440f-4735-579b-a4ab-409b98df4dab"
Distances = "b4f34e82-e78d-54a5-968a-f98e89d6e8f7"
HDF5 = "f67ccb44-e63f-5c2f-98bd-6dc0ccc4ba2f"
InteractiveUtils = "b77e0a4c-d291-57a0-90e8-8db25a27a240"
LinearAlgebra = "37e2e46d-f89d-539d-b4ee-838fcccc9c8e"
Random = "9a3f8284-a2c9-5f02-9a11-845980a1fd5c"
SparseArrays = "2f01184e-e22b-5df5-ae63-d93ebab69eaf"
Statistics = "10745b16-79ce-11e8-11f9-7d13ad32a3b2"
```
