## Supplementary material for "CellFishing.jl: an ultrafast and scalable cell search method for single-cell RNA-sequencing": README.md

# Experiments of CellFishing.jl

The script files to reproduce our <https://github.com/bicycle1885/CellFishing.jl> experiments.

## Prerequisites

- Julia (v1.0): to run experiments of CellFishing.jl.
- Pipenv: to run data processing and experiments.
- Docker: to create a Docker image including scmap and other R tools.
- Data: see below.

## Data

Download `clust_retinal_bipolar.txt` from
<https://portals.broadinstitute.org/single_cell/study/retinal-bipolar-neuron-drop-seq>
(need to sign in) and place the file in the `./data/raw/Shekhar2016` directory.
All other files will be downloaded in the Snakemake file.

**md5**:
```
1M_neurons/1M_neurons.h5                              f97b8c67e57a534816b9b2bccf31476b
1M_neurons/1M_neurons_analysis.tar.gz                 afa6abb31c0a2aa2a92f02ab62fea3cc
Baron2016/GSE84133_RAW.tar                            e9171075e0c45ab1aa1c2f7374e006d3
Plass2018/dge.txt.gz                                  8eca2fd3c10e40976a8e1b960001fe5b
Plass2018/R_annotation.txt                            03c36f364ca162c561a0056ed1ce5d45
Shekhar2016/clust_retinal_bipolar.txt                 c826450fdfc9ebac81280d294f6a3b86
Shekhar2016/GSE81904_BipolarUMICounts_Cell2016.txt.gz d85edfc0752bb85a176fa6e567b34c70
TabulaMuris/chromium/5968960.zip                      db808018ef9c28e70913842a982a1e48
TabulaMuris/smart/5829687.zip                         e1bcf40812c4e1a7ec0a17827e74e54c
```

## How to run experiments.

### 1. Install CellFishing.jl.

(TODO: Update the workflow; this will change in Julia 1.0.)

    julia -e 'using Pkg; Pkg.clone("https://github.com/bicycle1885/CellFishing.jl")'

### 2. Create a Docker image from the Docker file.

    docker build -t scmap .
    docker save -o scmap.tar scmap

### 3. Set up the Python environment from Pipenv.lock.

    pipenv install .

### 4. Run experiments.

    pipenv run snakemake all

All results and plots will be emitted into the `./results` and `./figures`
directories, respectively. `./results/Analysis.html` is an HTML report of the
experiments.
