## Supplementary material for "CellFishing.jl: an ultrafast and scalable cell search method for single-cell RNA-sequencing": Baron2016.loomify.py

```
import sys
import pandas
import loompy

clusterfile = sys.argv[1]
inputfiles = sys.argv[2:-1]
outputfile = sys.argv[-1]
cellnames = pandas.read_table(clusterfile)["cell"].values
data = []
for filename in inputfiles:
    df = pandas.read_csv(filename, index_col=0)
    data.append(df.drop(["barcode", "assigned_cluster"], axis=1))
data = pandas.concat(data).T
data = data.loc[:,cellnames]
matrix = data.astype("int32").values
loompy.create(outputfile, matrix, {"Gene": data.index.values}, {"CellID": data.columns.values})
```
