## Supplementary material for "CellFishing.jl: an ultrafast and scalable cell search method for single-cell RNA-sequencing": clustering-sc3.R

```
library(readr)
library(SingleCellExperiment)
library(scater)
library(SC3)
library(h5)

experiment <- function(outdir, data, k, r) {
    timer <- Sys.time()
    # preprocess
    sce <- SingleCellExperiment(assays=list(counts=data))
    sce <- normalize(sce)
    rowData(sce)$feature_symbol <- rownames(data)
    # run SC3
    sce <- sc3(sce, ks=k, svm_num_cells=5000, n_cores=1)
    # run SVM (hybrid approach)
    sce <- sc3_run_svm(sce, ks=k)
    elapsed <- as.double(Sys.time() - timer, units="secs")
    clusters <- data.frame(
        cell=rownames(colData(sce)),
        cluster=colData(sce)[,paste0("sc3_", k, "_clusters")])
    filename <- file.path(outdir, paste0("k-", k, ".", r, ".tsv"))
    system2("mkdir", args=c("-p", outdir))
    write_tsv(clusters, filename)
    system2("gzip", args=c("-f", filename))
    # report
    cat("[[experiment]]\n")
    cat("k = ", k, "\n", sep="")
    cat("elapsed-time = ", elapsed, "  # s\n", sep="")
    cat("filename = '", filename, ".gz'\n", sep="")
    cat("\n")
    flush.console()
}

starttime <- Sys.time()
cat("date-time = ", format(Sys.time(), "%Y-%m-%dT%H:%M:%S\n"), sep="")
cat("session-info = '''\n")
print(sessionInfo())
cat("'''\n\n")

args <- commandArgs(trailingOnly=T)
outdir <- args[1]
datafile <- args[2]
clusterfile <- args[3]

# Load data.
h5data <- h5file(datafile, "r")
data <- h5data["/matrix"][]
rownames(data) <- h5data["/row_attrs/Gene"][]
colnames(data) <- h5data["/col_attrs/CellID"][]
h5close(h5data)

# Set the number of clusters.
clusters <- read_tsv(clusterfile)
n_clusters <- length(unique(clusters$cluster))

# Run experiments.
set.seed(1234)
for (r in 1:5) {
    experiment(outdir, data, n_clusters, r)
}
cat("# END (", capture.output(print(Sys.time() - starttime)), ")\n", sep="")
```
