## Supplementary material for "CellFishing.jl: an ultrafast and scalable cell search method for single-cell RNA-sequencing": estimator.jl

```
using CellFishing
using Distances

include("utils.jl")

function experiment(outdir, data, subset, n_bits, superbit, r)
    counts = convert(Matrix{Float32}, data.counts)
    m, n = size(counts)
    features = CellFishing.selectfeatures(counts, data.featurenames)
    database = CellFishing.CellIndex(counts, features, n_bits=n_bits, superbit=superbit, n_lshashes=1)
    lshash = database.lshashes[1]
    X = CellFishing.preprocess(database.preproc, CellFishing.ExpressionMatrix(counts, data.featurenames), :database)
    Z = fill(zero(CellFishing.bitvectype(lshash)), n)
    CellFishing.sketch!(Z, X, lshash)
    H = hamming_distance(Z, subset)
    mkpath(outdir)
    # cosine distance
    filename_cos = joinpath(outdir, "n_bits-$(n_bits).superbit-$(superbit).$(r).cos.tsv")
    writedlm(filename_cos, pairwise(CosineDist(), X[:,subset], X))
    run(`gzip -f $(filename_cos)`)
    # approximated hamming distance
    filename_ham = joinpath(outdir, "n_bits-$(n_bits).superbit-$(superbit).$(r).ham.tsv")
    writedlm(filename_ham, H)
    run(`gzip -f $(filename_ham)`)
    # report
    println("[[experiment]]")
    println("n-genes = ", m)
    println("n-cells = ", n)
    println("n-cells-subset = ", length(subset))
    println("n-bits = ", n_bits)
    println("superbit = ", superbit)
    println("filename-cos = '", filename_cos, ".gz'")
    println("filename-ham = '", filename_ham, ".gz'")
    println()
    flush(stdout)
end

function hamming_distance(Z, subset)
    H = zeros(Int32, length(subset), length(Z))
    for j in 1:length(Z), i in 1:length(subset)
        H[i,j] = count_ones(xor(Z[subset[i]], Z[j]))
    end
    return H
end

@experiments begin
    outdir, datafile, subsetfile = ARGS
    data = loadloom(datafile)
    name2idx = Dict(c => i for (i, c) in enumerate(data.cellnames))
    subset = [name2idx[c] for c in readlines(subsetfile)]
    for n_bits in [64, 128, 256, 512], superbit in [1, 50]
        Random.seed!(RANDSEED)
        for r in 1:5; experiment(outdir, data, subset, n_bits, superbit, r); end
    end
end
```
