## Supplementary material for "CellFishing.jl: an ultrafast and scalable cell search method for single-cell RNA-sequencing": hash.jl

```
using CellFishing

include("utils.jl")

function experiment(outdir, data, cells_query, cells_database, project)
    cellnames = data.cellnames
    featurenames = data.featurenames
    counts_query = convert(Matrix{Float32}, data.counts[:,cells_query])
    counts_database = convert(Matrix{Float32}, data.counts[:,cells_database])
    if project
        counts = counts_database
    else
        counts = hcat(counts_database, counts_query)
    end
    features = CellFishing.selectfeatures(counts, featurenames)
    database = CellFishing.CellIndex(counts, features, n_bits=64, n_lshashes=4)
    mkpath(outdir)
    # cell names
    filename_cells_query = joinpath(outdir, "cells.query.project-$(project).txt")
    writelines(filename_cells_query, cellnames[cells_query])
    filename_cells_database = joinpath(outdir, "cells.database.project-$(project).txt")
    writelines(filename_cells_database, cellnames[cells_database])
    # reduced matrix
    preprocess(counts) = CellFishing.preprocess(database.preproc, CellFishing.ExpressionMatrix(counts, featurenames), :both)
    filename_X_query = joinpath(outdir, "X.query.project-$(project).tsv")
    X_query = preprocess(counts_query)
    writedlm(filename_X_query, X_query)
    filename_X_database = joinpath(outdir, "X.database.project-$(project).tsv")
    X_database = preprocess(counts_database)
    writedlm(filename_X_database, X_database)
    # bit vectors
    filenames_Z_query = String[]
    filenames_Z_database = String[]
    for (l, lshash) in enumerate(database.lshashes)
        # query
        Z = fill(zero(CellFishing.bitvectype(lshash)), length(cells_query))
        CellFishing.sketch!(Z, X_query, lshash)
        filename = joinpath(outdir, "Z.query.project-$(project).$(l).bin")
        writehash(filename, Z)
        push!(filenames_Z_query, filename)
        # database
        Z = fill(zero(CellFishing.bitvectype(lshash)), length(cells_database))
        CellFishing.sketch!(Z, X_database, lshash)
        filename = joinpath(outdir, "Z.database.project-$(project).$(l).bin")
        writehash(filename, Z)
        push!(filenames_Z_database, filename)
    end
    # report
    println("[[experiment]]")
    println("project = ", project)
    println("n-genes = ", size(data.counts, 1))
    println("n-query-cells = ", size(counts_query, 2))
    println("n-database-cells = ", size(counts_database, 2))
    println("n-selected-features = ", CellFishing.nselected(features))
    println("filename-cells-query = '", filename_cells_query, "'")
    println("filename-cells-database = '", filename_cells_database, "'")
    println("filename-X-query = '", filename_X_query, "'")
    println("filename-X-database = '", filename_X_database, "'")
    println("filenames-Z-query = '", filenames_Z_query, "'")
    println("filenames-Z-database = '", filenames_Z_database, "'")
    println()
    flush(stdout)
end

function writelines(filename, cellnames)
    open(filename, "w") do file
        for name in cellnames
            println(file, name)
        end
    end
end

function writehash(filename, Z)
    @assert sizeof(eltype(Z)) == 8  # 64 bits
    open(filename, "w") do file
        for z in Z
            write(file, reinterpret(UInt64, z))
        end
    end
end

@experiments begin
    outdir, datafile, queryfile = ARGS
    databasefiles = ARGS[4:end]
    data = loadloom(datafile)
    name2idx = Dict(c => i for (i, c) in enumerate(data.cellnames))
    cells_query = [name2idx[c] for c in readlines(queryfile)]
    cells_database = [name2idx[c] for databasefile in databasefiles for c in readlines(databasefile)]
    @assert all(cells_query .> 0)
    @assert all(cells_database .> 0)
    @assert isempty(cells_query ∩ cells_database)
    for project in [false, true]
        Random.seed!(RANDSEED)
        experiment(outdir, data, cells_query, cells_database, project)
    end
end
```
