## Supplementary material for "CellFishing.jl: an ultrafast and scalable cell search method for single-cell RNA-sequencing": knn-batch.jl

```
using CellFishing

include("utils.jl")

function experiment(outdir, data, cells_query, cells_database, k, n_bits, n_lshashes, transformer, r)
    counts_query = data.counts[:,cells_query]
    counts_database = data.counts[:,cells_database]
    index_time = @elapsed begin
        features = CellFishing.selectfeatures(counts_database, data.featurenames)
        database = CellFishing.CellIndex(
            counts_database,
            features,
            n_bits=n_bits,
            n_lshashes=n_lshashes,
            transformer=transformer)
    end
    query_time = @elapsed nns = CellFishing.findneighbors(k, counts_query, data.featurenames, database)
    # save k-NNs
    filename = joinpath(outdir, "n_nns-$(k).n_bits-$(n_bits).n_lshashes-$(n_lshashes).trans-$(transformer).$(r).tsv")
    writeknn(filename, data.cellnames[cells_query], data.cellnames[cells_database], nns.indexes)
    run(`gzip -f $(filename)`)
    # report
    println("[[experiment]]")
    println("n-genes = ", size(data.counts, 1))
    println("n-cells-query = ", length(cells_query))
    println("n-cells-database = ", length(cells_database))
    println("n-nns = ", k)
    println("n-bits = ", n_bits)
    println("n-lshashes = ", n_lshashes)
    println("transformer = '", transformer, "'")
    println("index-time = ", index_time, "  # s")
    println("query-time = ", query_time, "  # s")
    println("preproc-time = ", database.rtstats.preproc_time, "  # ns")
    println("search-time = ", database.rtstats.search_time, "  # ns")
    println("rank-time = ", database.rtstats.rank_time, "  # ns")
    println("filename = '", filename, ".gz'")
    println()
    flush(stdout)
end

@experiments begin
    outdir, datafile, queryfile = ARGS
    databasefiles = ARGS[4:end]
    data = loadloom(datafile)
    name2idx = Dict(c => i for (i, c) in enumerate(data.cellnames))
    cells_query = [name2idx[c] for c in readlines(queryfile)]
    cells_database = [name2idx[c] for databasefile in databasefiles for c in readlines(databasefile)]
    @assert all(cells_query .> 0)
    @assert all(cells_database .> 0)
    @assert isempty(cells_query ∩ cells_database)
    k = 10
    for n_bits in [128, 256], n_lshashes in [1, 4, 16], transformer in [:log1p, :ftt]
        Random.seed!(RANDSEED)
        for r in 1:5; experiment(outdir, data, cells_query, cells_database, k, n_bits, n_lshashes, transformer, r); end
    end
end
```
