## Supplementary material for "CellFishing.jl: an ultrafast and scalable cell search method for single-cell RNA-sequencing": knn-batch.scmap.R

```
library(readr)
library(SingleCellExperiment)
library(scater)
library(scmap)
library(matrixStats)
library(h5)

experiment <- function(outdir, data, cells_query, cells_database, k, n_features, n_centroids_factor, r) {
    m <- nrow(data)
    n <- ncol(data)
    data_query <- data[,cells_query]
    data_database <- data[,cells_database]
    n_centroids <- floor(sqrt(ncol(data_database)) * n_centroids_factor)  # the default value times the factor
    timer <- Sys.time()
    sce_database <- make_sce(data_database)
    sce_database <- selectFeatures(sce_database, suppress_plot=T, n_features=n_features)
    sce_database <- indexCell(sce_database, k=n_centroids)
    index_time <- as.double(Sys.time() - timer, units="secs")
    timer <- Sys.time()
    sce_query <- make_sce(data_query)
    results <- scmapCell(sce_query, list(db=metadata(sce_database)$scmap_cell_index), w=k)
    query_time <- as.double(Sys.time() - timer, units="secs")
    nns <- list()
    for (i in 1:length(cells_query)) {
        # sort cells by similarity
        nns[[i]] <- cells_database[results$db$cells[order(results$db$similarities[,i], decreasing=T),i]]
    }
    nns <- data.frame(t(matrix(colnames(data)[unlist(nns)], nrow=k)))
    colnames(nns) <- paste0("N", 1:k)
    nns <- cbind(data.frame(cell=colnames(data)[cells_query]), nns)
    system2("mkdir", args=c("-p", outdir))
    filename <- file.path(outdir, paste0("n_nns-", k, ".n_features-", n_features, ".n_centroids_factor-", n_centroids_factor, ".", r, ".tsv"))
    write_tsv(nns, filename)
    system2("gzip", args=c("-f", filename))
    # report
    cat("[[experiment]]\n")
    cat("n-genes = ", m, "\n", sep="")
    cat("n-cells-query = ", length(cells_query), "\n", sep="")
    cat("n-cells-database = ", length(cells_database), "\n", sep="")
    cat("n-nns = ", k, "\n", sep="")
    cat("n-features = ", n_features, "\n", sep="")
    cat("n-centroids-factor = ", n_centroids_factor, "\n", sep="")
    cat("index-time = ", index_time, "  # s\n", sep="")
    cat("query-time = ", query_time, "  # s\n", sep="")
    cat("filename = '", filename, ".gz'\n", sep="")
    cat("\n")
    flush.console()
}

make_sce <- function(data) {
    sce <- SingleCellExperiment(assays=list(counts=data))
    sce <- normalize(sce, centre_size_factors=F)
    rowData(sce)$feature_symbol <- rownames(data)
    return(sce)
}

starttime <- Sys.time()
cat("date-time = ", format(Sys.time(), "%Y-%m-%dT%H:%M:%S\n"), sep="")
cat("session-info = '''\n")
print(sessionInfo())
cat("'''\n\n")

args <- commandArgs(trailingOnly=T)
outdir <- args[1]
datafile <- args[2]
queryfile <- args[3]
databasefiles <- args[4:length(args)]

# Load data.
h5data <- h5file(datafile, "r")
data <- h5data["/matrix"][]
rownames(data) <- h5data["/row_attrs/Gene"][]
colnames(data) <- h5data["/col_attrs/CellID"][]
h5close(h5data)

# Find cell indexes.
cells_query <- match(read_lines(queryfile), colnames(data))
cells_database <- match(unlist(lapply(databasefiles, read_lines)), colnames(data))
stopifnot(length(intersect(cells_query, cells_database)) == 0)

# Run experiments
k <- 10
for (n_features in c(500, 1000, 2000, 4000)) {
    for (n_centroids_factor in c(1, 4)) {
        set.seed(1234)
        for (r in 1:5) {
            experiment(outdir, data, cells_query, cells_database, k, n_features, n_centroids_factor, r)
        }
    }
}
cat("# END (", capture.output(print(Sys.time() - starttime)), ")\n", sep="")
```
