## Supplementary material for "CellFishing.jl: an ultrafast and scalable cell search method for single-cell RNA-sequencing": knn-cv-features.jl

```
using CellFishing

include("utils.jl")

const K = 5

function experiment(outdir, data, k, featuresfile, r)
    counts, featurenames = data.counts, data.featurenames
    m, n = size(counts)
    blocks = make_random_blocks(n, K)
    index_time = query_time = 0.0
    preproc_time = search_time = rank_time = UInt64(0)
    nns = zeros(Int, k, n)
    for j in 1:K
        cells_query = blocks[j]
        cells_database = vcat([blocks[l] for l in 1:K if l != j]...)
        counts_query = counts[:,cells_query]
        counts_database = counts[:,cells_database]
        index_time += @elapsed begin
            features = CellFishing.Features(featurenames)
            CellFishing.addfeatures!(features, readlines(featuresfile))
            # drop features with no variability
            CellFishing.dropfeatures!(features, featurenames[vec(minimum(counts_database, dims=2) .== maximum(counts_database, dims=2))])
            database = CellFishing.CellIndex(counts_database, features)
        end
        query_time += @elapsed nns′ = CellFishing.findneighbors(
            k,
            counts_query,
            featurenames,
            database)
        preproc_time += database.rtstats.preproc_time
        search_time += database.rtstats.search_time
        rank_time += database.rtstats.rank_time
        for i in 1:size(counts_query, 2)
            nns[:,cells_query[i]] = cells_database[nns′.indexes[:,i]]
        end
    end
    @assert all(i->i!=0, nns)
    # save k-NNs
    filename = joinpath(outdir, "n_nns-$(k).$(r).tsv")
    writeknn(filename, data.cellnames, nns)
    run(`gzip -f $(filename)`)
    # report
    println("[[experiment]]")
    println("n-genes = ", m)
    println("n-cells = ", n)
    println("n-nns = ", k)
    println("features-file = '", featuresfile, "'")
    println("index-time = ", index_time, "  # s")
    println("query-time = ", query_time, "  # s")
    println("preproc-time = ", preproc_time, "  # ns")
    println("search-time = ", search_time, "  # ns")
    println("rank-time = ", rank_time, "  # ns")
    println("filename = '", filename, ".gz'")
    println()
    flush(stdout)
end

@experiments begin
    outdir, datafile, featuresfile = ARGS
    data = loadloom(datafile)
    k = 10
    Random.seed!(RANDSEED)
    for r in 1:5; experiment(outdir, data, k, featuresfile, r); end
end
```
