## Supplementary material for "CellFishing.jl: an ultrafast and scalable cell search method for single-cell RNA-sequencing": knn-cv-features.scmap.R

```
library(readr)
library(SingleCellExperiment)
library(scater)
library(scmap)
library(matrixStats)
library(h5)

# The parameter of K-fold cross validation.
K <- 5

# K-fold cross validation.
experiment <- function(outdir, data, k, features_file, n_centroids_factor, r) {
    m <- nrow(data)
    n <- ncol(data)
    features <- read_lines(features_file)
    blocks <- make_random_blocks(n, K)
    index_time <- 0.0
    query_time <- 0.0
    nns <- list()
    for (j in 1:K) {
        cells_query <- blocks[[j]]
        cells_index <- unlist(blocks[-j])
        data_query <- data[,cells_query]
        data_index <- data[,cells_index]
        n_centroids <- floor(sqrt(ncol(data_index)) * n_centroids_factor)  # the default value times the factor
        timer <- Sys.time()
        sce_index <- make_sce(data_index)
        sce_index <- setFeatures(sce_index, features)
        sce_index <- indexCell(sce_index, k=n_centroids)
        index_time <- index_time + as.double(Sys.time() - timer, units="secs")
        timer <- Sys.time()
        sce_query <- make_sce(data_query)
        results <- scmapCell(sce_query, list(self=metadata(sce_index)$scmap_cell_index), w=k)
        query_time <- query_time + as.double(Sys.time() - timer, units="secs")
        for (i in 1:length(cells_query)) {
            # sort cells by similarity
            nns[[cells_query[i]]] <- cells_index[results$self$cells[order(results$self$similarities[,i], decreasing=T),i]]
        }
    }
    nns <- data.frame(t(matrix(colnames(data)[unlist(nns)], nrow=k)))
    colnames(nns) <- paste0("N", 1:k)  # 'N' is neighbor.
    nns <- cbind(data.frame(cell=colnames(data)), nns)
    system2("mkdir", args=c("-p", outdir))
    filename <- file.path(outdir, paste0("n_nns-", k, ".n_centroids_factor-", n_centroids_factor, ".", r, ".tsv"))
    write_tsv(nns, filename)
    # report
    system2("gzip", args=c("-f", filename))
    cat("[[experiment]]\n")
    cat("n-genes = ", m, "\n", sep="")
    cat("n-cells = ", n, "\n", sep="")
    cat("n-nns = ", k, "\n", sep="")
    cat("features-file = '", features_file, "'\n", sep="")
    cat("n-centroids-factor = ", n_centroids_factor, "\n", sep="")
    cat("index-time = ", index_time, "  # s\n", sep="")
    cat("query-time = ", query_time, "  # s\n", sep="")
    cat("filename = '", filename, ".gz'\n", sep="")
    cat("\n")
    flush.console()
}

make_sce <- function(data) {
    sce <- SingleCellExperiment(assays=list(counts=data))
    sce <- normalize(sce, centre_size_factors=F)
    rowData(sce)$feature_symbol <- rownames(data)
    return(sce)
}

make_random_blocks <- function(n, k) {
    cells <- sample(1:n)
    blocks <- list()
    q <- n %/% k
    r <- n %% k
    i <- 0
    l <- 1
    while (i < n) {
        sz <- q
        if (r > 0) {
            sz <- sz + 1
            r <- r - 1
        }
        blocks[[l]] <- cells[(i+1):(i+sz)]
        i <- i + sz
        l <- l + 1
    }
    return(blocks)
}

starttime <- Sys.time()
cat("date-time = ", format(Sys.time(), "%Y-%m-%dT%H:%M:%S\n"), sep="")
cat("session-info = '''\n")
print(sessionInfo())
cat("'''\n\n")

args <- commandArgs(trailingOnly=T)
outdir <- args[1]
datafile <- args[2]
features_file <- args[3]

# Load data.
h5data <- h5file(datafile, "r")
data <- h5data["/matrix"][]
rownames(data) <- h5data["/row_attrs/Gene"][]
colnames(data) <- h5data["/col_attrs/CellID"][]
h5close(h5data)

# Run experiments.
k <- 10
n_centroids_factor <- 1
set.seed(1234)
for (r in 1:5) {
    experiment(outdir, data, k, features_file, n_centroids_factor, r)
}
cat("# END (", capture.output(print(Sys.time() - starttime)), ")\n", sep="")
```
