## Supplementary material for "CellFishing.jl: an ultrafast and scalable cell search method for single-cell RNA-sequencing": knn-degs.jl

```
using CellFishing

include("utils.jl")

function experiment(outdir, data, labels, target, k)
    counts, featurenames, cellnames = data.counts, data.featurenames, data.cellnames
    m, n = size(counts)
    cells_target = findall(labels .== target)
    cells_database = findall(labels .!= target)
    counts_target = counts[:,cells_target]
    # without target cells
    counts = counts[:,cells_database]
    features = CellFishing.selectfeatures(counts, featurenames)
    database = CellFishing.CellIndex(counts, features, keep_counts=true)
    neighbors = CellFishing.findneighbors(10, counts_target, featurenames, database)
    filename_knn = joinpath(outdir, "knn.k-$(k).tsv")
    writeknn(filename_knn, cellnames[cells_target], cellnames[cells_database], neighbors.indexes)
    run(`gzip -f $(filename_knn)`)
    degs = CellFishing.finddegs(counts_target, featurenames, neighbors.indexes[1,:], database, k=k)
    filename_degs = joinpath(outdir, "degs.k-$(k).tsv")
    writedegs(filename_degs, degs, cellnames[cells_target])
    run(`gzip -f $(filename_degs)`)
    # report
    println("[[experiment]]")
    println("target = '$(target)'")
    println("k = $(k)")
    println("filename-knn = '$(filename_knn).gz'")
    println("filename-degs = '$(filename_degs).gz'")
    println()
    flush(stdout)
end

function writedegs(filename, degs, cellnames)
    m, n = size(degs.means)
    featurenames = degs.names
    @assert m == length(featurenames)
    @assert n == length(cellnames)
    open(filename, "w") do file
        println(file, "cell\tgene\tnegative\tpositive\tmean")
        for j in 1:n
            for i in 1:m
                println(file, cellnames[j], '\t', featurenames[i], '\t', degs.negatives[i,j], '\t', degs.positives[i,j], '\t', degs.means[i,j])
            end
        end
    end
end

@experiments begin
    outdir, datafile, labelfile, target = ARGS
    data = loadloom(datafile)
    labels = [String(split(line, '\t')[2]) for line in eachline(labelfile)][2:end]
    target = replace(target, ';' => '/')
    for k in [5, 10, 20]
        Random.seed!(RANDSEED)
        experiment(outdir, data, labels, target, k)
    end
end
```
