## Supplementary material for "CellFishing.jl: an ultrafast and scalable cell search method for single-cell RNA-sequencing": knn-protocol.jl

```
using CellFishing

include("utils.jl")

function experiment(outdir, querydata, databasedata, k, r)
    index_time = @elapsed begin
        features = CellFishing.selectfeatures(databasedata.counts, databasedata.featurenames)
        database = CellFishing.CellIndex(databasedata.counts, features)
    end
    query_time = @elapsed nns = CellFishing.findneighbors(k, querydata.counts, querydata.featurenames, database)
    # save k-NNs
    filename = joinpath(outdir, "n_nns-$(k).$(r).tsv")
    writeknn(filename, querydata.cellnames, databasedata.cellnames, nns.indexes)
    run(`gzip -f $(filename)`)
    # report
    println("[[experiment]]")
    println("n-cells-query = ", length(querydata.cellnames))
    println("n-cells-database = ", length(databasedata.cellnames))
    println("n-nns = ", k)
    println("index-time = ", index_time, "  # s")
    println("query-time = ", query_time, "  # s")
    println("preproc-time = ", database.rtstats.preproc_time, "  # ns")
    println("search-time = ", database.rtstats.search_time, "  # ns")
    println("rank-time = ", database.rtstats.rank_time, "  # ns")
    println("filename = '", filename, ".gz'")
    println()
    flush(stdout)
end

@experiments begin
    outdir, queryfile, databasefile = ARGS
    querydata = loadloom(queryfile)
    databasedata = loadloom(databasefile)
    k = 10
    Random.seed!(RANDSEED)
    for r in 1:5; experiment(outdir, querydata, databasedata, k, r); end
end
```
