## Supplementary material for "CellFishing.jl: an ultrafast and scalable cell search method for single-cell RNA-sequencing": knn-protocol.scmap.R

```
library(readr)
library(SingleCellExperiment)
library(scater)
library(scmap)
library(matrixStats)
library(h5)

experiment <- function(outdir, data_query, data_index, k, r) {
    timer <- Sys.time()
    sce_index <- make_sce(data_index)
    sce_index <- selectFeatures(sce_index, suppress_plot=T)
    sce_index <- indexCell(sce_index)
    index_time <- as.double(Sys.time() - timer, units="secs")
    timer <- Sys.time()
    sce_query <- make_sce(data_query)
    results <- scmapCell(sce_query, list(self=metadata(sce_index)$scmap_cell_index), w=k)
    query_time <- as.double(Sys.time() - timer, units="secs")
    nns <- list()
    for (i in 1:ncol(data_query)) {
        # sort cells by similarity
        nns[[i]] <- results$self$cells[order(results$self$similarities[,i], decreasing=T),i]
    }
    nns <- data.frame(t(matrix(colnames(data_index)[unlist(nns)], nrow=k)))
    colnames(nns) <- paste0("N", 1:k)  # 'N' is neighbor.
    nns <- cbind(data.frame(cell=colnames(data_query)), nns)
    filename <- file.path(outdir, paste0("n_nns-", k, ".", r, ".tsv"))
    system2("mkdir", args=c("-p", outdir))
    write_tsv(nns, filename)
    system2("gzip", args=c("-f", filename))
    # report
    cat("[[experiment]]\n")
    cat("n-genes = ", nrow(data_index), "\n", sep="")
    cat("n-cells-query = ", ncol(data_query), "\n", sep="")
    cat("n-cells-database = ", ncol(data_index), "\n", sep="")
    cat("n-nns = ", k, "\n", sep="")
    cat("index-time = ", index_time, "  # s\n", sep="")
    cat("query-time = ", query_time, "  # s\n", sep="")
    cat("filename = '", filename, ".gz'\n", sep="")
    cat("\n")
    flush.console()
}

make_sce <- function(data) {
    sce <- SingleCellExperiment(assays=list(counts=data))
    sce <- normalize(sce, centre_size_factors=F)
    rowData(sce)$feature_symbol <- rownames(data)
    return(sce)
}

load_loom <- function(filename) {
    h5data <- h5file(filename, "r")
    data <- h5data["/matrix"][]
    rownames(data) <- h5data["/row_attrs/Gene"][]
    colnames(data) <- h5data["/col_attrs/CellID"][]
    h5close(h5data)
    return(data)
}

starttime <- Sys.time()
cat("date-time = ", format(Sys.time(), "%Y-%m-%dT%H:%M:%S\n"), sep="")
cat("session-info = '''\n")
print(sessionInfo())
cat("'''\n\n")

args <- commandArgs(trailingOnly=T)
outdir <- args[1]
file_query <- args[2]
file_index <- args[3]

# Load data.
data_query <- load_loom(file_query)
data_index <- load_loom(file_index)

# Run experiments.
k <- 10
set.seed(1234)
for (r in 1:5) {
    experiment(outdir, data_query, data_index, k, r)
}
cat("# END (", capture.output(print(Sys.time() - starttime)), ")\n", sep="")
```
