## Supplementary material for "CellFishing.jl: an ultrafast and scalable cell search method for single-cell RNA-sequencing": knn-similarity.jl

```
using CellFishing

include("utils.jl")

const K = 5

function experiment(outdir, data, labels, target)
    counts, featurenames, cellnames = data.counts, data.featurenames, data.cellnames
    k = 10
    m, n = size(counts)
    blocks = make_random_blocks(n, K)
    cells_target = findall(labels .== target)
    knn_with_target_cells = fill(0, k, n)
    knn_without_target_cells = fill(0, k, n)
    similarities_with_target_cells = fill(NaN32, k, n)
    similarities_without_target_cells = fill(NaN32, k, n)
    for j in 1:K
        cells_query = blocks[j]
        cells_database = vcat([blocks[l] for l in 1:K if l != j]...)
        counts_query = counts[:,cells_query]
        # with target cells
        counts_database = counts[:,cells_database]
        features = CellFishing.selectfeatures(counts_database, featurenames)
        database = CellFishing.CellIndex(counts_database, features)
        neighbors = CellFishing.findneighbors(k, counts_query, featurenames, database)
        knn_with_target_cells[:,cells_query] = cells_database[neighbors.indexes]
        similarities_with_target_cells[:,cells_query] = CellFishing.similarities(neighbors, database)
        # without target cells
        cells_database = setdiff(cells_database, cells_target)
        counts_database = counts[:,cells_database]
        features = CellFishing.selectfeatures(counts_database, featurenames)
        database = CellFishing.CellIndex(counts_database, features)
        neighbors = CellFishing.findneighbors(k, counts_query, featurenames, database)
        knn_without_target_cells[:,cells_query] = cells_database[neighbors.indexes]
        similarities_without_target_cells[:,cells_query] = CellFishing.similarities(neighbors, database)
    end
    @assert all(knn_with_target_cells .> 0)
    @assert all(knn_without_target_cells .> 0)
    @assert all(isfinite.(similarities_with_target_cells))
    @assert all(isfinite.(similarities_without_target_cells))
    filename_knn_with_target_cells = joinpath(outdir, "knn.with-target.tsv")
    writeknn(filename_knn_with_target_cells, cellnames, knn_with_target_cells)
    run(`gzip -f $(filename_knn_with_target_cells)`)
    filename_similarities_with_target_cells = joinpath(outdir, "similarities.with-target.tsv")
    writesim(filename_similarities_with_target_cells, cellnames, similarities_with_target_cells)
    run(`gzip -f $(filename_similarities_with_target_cells)`)
    filename_knn_without_target_cells = joinpath(outdir, "knn.without-target.tsv")
    writeknn(filename_knn_without_target_cells, cellnames, knn_without_target_cells)
    run(`gzip -f $(filename_knn_without_target_cells)`)
    filename_similarities_without_target_cells = joinpath(outdir, "similarities.without-target.tsv")
    writesim(filename_similarities_without_target_cells, cellnames, similarities_without_target_cells)
    run(`gzip -f $(filename_similarities_without_target_cells)`)
    # report
    println("[[experiment]]")
    println("n-genes = ", m)
    println("n-cells = ", n)
    println("target = '", target, "'")
    println("filename-knn-with-target-cells = '", filename_knn_with_target_cells, ".gz'")
    println("filename-similarities-with-target-cells = '", filename_similarities_with_target_cells, ".gz'")
    println("filename-knn-without-target-cells = '", filename_knn_without_target_cells, ".gz'")
    println("filename-similarities-without-target-cells = '", filename_similarities_without_target_cells, ".gz'")
    println()
    flush(stdout)
end

function writesim(filename, cellnames, similarities)
    mkpath(dirname(filename))
    k, n = size(similarities)
    open(filename, "w") do file
        # header
        println(file, join(["cell"; string.("N", 1:k)], '\t'))
        # body
        for j in 1:n
            print(file, cellnames[j])
            for i in 1:k
                print(file, '\t', similarities[i,j])
            end
            println(file)
        end
    end
end

@experiments begin
    outdir, datafile, labelfile, target = ARGS
    data = loadloom(datafile)
    labels = [String(split(line, '\t')[2]) for line in eachline(labelfile)][2:end]
    target = replace(target, ';' => '/')
    Random.seed!(RANDSEED)
    experiment(outdir, data, labels, target)
end
```
