## Supplementary material for "CellFishing.jl: an ultrafast and scalable cell search method for single-cell RNA-sequencing": save-load.jl

```
using CellFishing

include("utils.jl")

function experiment(data, n_bits, keep_counts)
    m, n = size(data.counts)
    println("[[experiment]]")
    println("n-genes = ", m)
    println("n-cells = ", n)
    println("n-bits = ", n_bits)
    println("keep-counts = ", keep_counts)
    println("index-time = ", @elapsed begin
        features = CellFishing.selectfeatures(data.counts, data.featurenames)
        database = CellFishing.CellIndex(
            data.counts, features,
            metadata=data.cellnames,
            n_bits=n_bits, keep_counts=keep_counts)
    end)
    println("ram-size = ", Base.summarysize(database))
    mktempdir() do tmpdir
        tmpfile = joinpath(tmpdir, "database")
        println("save-time = ", @elapsed CellFishing.save(tmpfile, database))
        println("file-size = ", filesize(tmpfile))
        println("load-time = ", @elapsed CellFishing.load(tmpfile))
    end
    println()
    flush(stdout)
end

@experiments begin
    datafile, = ARGS
    data = loadloom(datafile)
    for n_bits in [128, 256], keep_counts in [false, true]
        Random.seed!(RANDSEED)
        for _ in 1:10; experiment(data, n_bits, keep_counts); end
    end
end
```
