## Supplementary material for "CellFishing.jl: an ultrafast and scalable cell search method for single-cell RNA-sequencing": scalability.jl

```
using CellFishing

include("utils.jl")

function experiment(outdir, data, cells_query, k, index, n_bits, db_size, r)
    counts, featurenames = data.counts, data.featurenames
    m, n = size(counts)
    @assert length(featurenames) == m
    @assert n - length(cells_query) ≥ db_size
    cells_database = shuffle!(setdiff(1:n, cells_query))[1:db_size]
    counts_database = counts[:,cells_database]
    counts_query = counts[:,cells_query]
    index_time = @elapsed begin
        features = CellFishing.selectfeatures(counts_database, featurenames)
        database = CellFishing.CellIndex(counts_database, features, n_bits=n_bits, index=index)
    end
    query_time = @elapsed nns = CellFishing.findneighbors(k, counts_query, featurenames, database)
    preproc_time = database.rtstats.preproc_time
    search_time = database.rtstats.search_time
    rank_time = database.rtstats.rank_time
    ram_size = Base.summarysize(database)
    # save k-NNs
    filename = joinpath(outdir, "n_nns-$(k).index-$(index).n_bits-$(n_bits).db_size-$(db_size).$(r).tsv")
    writeknn(filename, data.cellnames[cells_query], data.cellnames[cells_database], nns.indexes)
    run(`gzip -f $(filename)`)
    # report
    println("[[experiment]]")
    println("n-genes = ", m)
    println("n-cells-database = ", length(cells_database))
    println("n-cells-query = ", length(cells_query))
    println("indexed = ", index)
    println("n-nns = ", k)
    println("n-bits = ", n_bits)
    println("index-time = ", index_time, "  # s")
    println("query-time = ", query_time, "  # s")
    println("preproc-time = ", preproc_time, "  # ns")
    println("search-time = ", search_time, "  # ns")
    println("rank-time = ", rank_time, "  # ns")
    println("ram-size = ", ram_size, "  # bytes")
    println("filename = '", filename, ".gz'")
    println()
    flush(stdout)
end

@experiments begin
    outdir, datafile, queryfile = ARGS
    data = loadchromium(datafile, "/mm10")
    name2idx = Dict(c => i for (i, c) in enumerate(data.cellnames))
    cells_query = [name2idx[c] for c in readlines(queryfile)]
    k = 10
    for index in false:true, n_bits in [128, 256], db_size in 2 .^ (13:20)
        Random.seed!(RANDSEED)
        for r in 1:5; experiment(outdir, data, cells_query, k, index, n_bits, db_size, r); end
    end
end
```
