## Supplementary material for "CellFishing.jl: an ultrafast and scalable cell search method for single-cell RNA-sequencing": select-features.jl

```
using CellFishing

include("utils.jl")

datafile, n_min_features = ARGS
data = loadloom(datafile)
n_min_features = parse(Int, n_min_features)

# Write selected features.
Random.seed!(RANDSEED)
if n_min_features == 0  # default
    features = CellFishing.selectfeatures(data.counts, data.featurenames)
else
    features = CellFishing.selectfeatures(data.counts, data.featurenames, n_min_features=n_min_features)
end
for gene in CellFishing.selectedfeatures(features)
    println(gene)
end
```
