## Supplementary material for "CellFishing.jl: an ultrafast and scalable cell search method for single-cell RNA-sequencing": select-features.scmap.R

```
library(SingleCellExperiment)
library(scater)
library(scmap)
library(h5)

make_sce <- function(data) {
    sce <- SingleCellExperiment(assays=list(counts=data))
    sce <- normalize(sce, centre_size_factors=F)
    rowData(sce)$feature_symbol <- rownames(data)
    return(sce)
}

args <- commandArgs(trailingOnly=T)
datafile <- args[1]
n_features <- as.integer(args[2])

# Load data.
h5data <- h5file(datafile, "r")
data <- h5data["/matrix"][]
rownames(data) <- h5data["/row_attrs/Gene"][]
colnames(data) <- h5data["/col_attrs/CellID"][]
h5close(h5data)

# Write selected features.
set.seed(1234)
sce <- make_sce(data)
sce <- selectFeatures(sce, suppress_plot=T, n_features=n_features)
selected_features <- rowData(sce)$feature_symbol[rowData(sce)$scmap_features]
for (gene in selected_features) {
    cat(gene, "\n", sep="")
}
```
