## Supplementary material for "CellFishing.jl: an ultrafast and scalable cell search method for single-cell RNA-sequencing": svd.jl

```
using CellFishing

include("utils.jl")

function experiment(X, n_dims, algorithm)
    if algorithm == :full
        time_svd = @elapsed _, svals, _ = svd(X)
        svals = svals[1:n_dims]
    elseif algorithm == :trunc
        time_svd = @elapsed _, svals, _ = CellFishing.tsvd(X, n_dims)
    elseif algorithm == :rand
        time_svd = @elapsed _, svals, _ = CellFishing.rsvd(X, n_dims)
    else
        @assert(false)
    end
    # report
    println("[[experiment]]")
    println("n-dims = ", n_dims)
    println("n-rows = ", size(X, 1))
    println("n-columns = ", size(X, 2))
    println("algorithm = '", algorithm, "'")
    println("singular-values = [", join(svals, ", "), "]")
    println("time-svd = ", time_svd, "  # s")
    println()
    flush(stdout)
end

BLAS.set_num_threads(2)

@experiments begin
    datafile, = ARGS
    data = loadloom(datafile)

    # Preprocess data.
    features = CellFishing.selectfeatures(data.counts, data.featurenames)
    X = convert(Matrix{Float32}, data.counts[features.selected,:])
    X .*= 1.0f4 ./ sum(X, dims=1)
    CellFishing.transform!(CellFishing.LogT(1.0f0), X)
    X .-= mean(X, dims=2)
    X ./= std(X, dims=2)
    @assert X isa Matrix{Float32}

    n_dims = 50
    for algorithm in [:full, :trunc, :rand]
        Random.seed!(RANDSEED)
        for _ in 1:10; experiment(X, n_dims, algorithm); end
    end
end
```
