## Supplementary material for "CellFishing.jl: an ultrafast and scalable cell search method for single-cell RNA-sequencing": TabulaMuris-chromium.loomify.py

```
import os
import sys
import numpy
import scipy.io
import pandas
import loompy

clusterfile = sys.argv[1]
rootdir = sys.argv[2]
outputfile = sys.argv[3]

cellnames = pandas.read_table(clusterfile)["cell"].values
colnames = []
genes = []
Ms = []
firstfile = True
for dropletdir in os.listdir(rootdir):
    if dropletdir.startswith("."):
        continue
    dirname = dropletdir.split("-")[1]
    dropletdir = os.path.join(rootdir, dropletdir)
    print("  reading", dropletdir)
    with open(os.path.join(dropletdir, "barcodes.tsv")) as f:
        barcodes = []
        for line in f:
            barcodes.append(line.split("-")[0])
    with open(os.path.join(dropletdir, "genes.tsv")) as f:
        for j, line in enumerate(f):
            x, y = line.rstrip().split("\t")
            assert x == y
            if firstfile:
                # initialize gene list
                genes.append(x)
            else:
                # check consistency
                assert x == genes[j]
    M = scipy.io.mmread(os.path.join(dropletdir, "matrix.mtx")).astype(numpy.int16)
    n_detected_genes = numpy.array((M > 0).sum(axis=0)).ravel()
    n_umi_counts = numpy.array(M.sum(axis=0)).ravel()
    # QC filter taken from http://tabula-muris.ds.czbiohub.org/
    passed = (n_detected_genes > 500) & (n_umi_counts > 1000)
    # make cell names like: 10X_P7_8_AAACGGGTCTCGTATT
    colnames.extend([dirname + "_" + barcode for barcode in numpy.array(barcodes)[passed]])
    M = M.tocsr()[:,passed]
    Ms.append(M)
    firstfile = False

Y = numpy.hstack([M.toarray() for M in Ms])
data = pandas.DataFrame(Y, index=pandas.Series(genes, name="gene"), columns=colnames)
artificial = data.index.str.startswith("ERCC-") | data.index.str.endswith("_transgene")
data = data.loc[~artificial,cellnames].astype("int32")
loompy.create(outputfile, data.values, {"Gene": data.index.values}, {"CellID": data.columns.values})
```
