## Supplementary material for "CellFishing.jl: an ultrafast and scalable cell search method for single-cell RNA-sequencing": TabulaMuris-smart.loomify.py

```
import os
import sys
import numpy
import pandas
import loompy

clusterfile = sys.argv[1]
rootdir = sys.argv[2]
outputfile = sys.argv[3]

cellnames = pandas.read_table(clusterfile)["cell"].values
genes = None
data = []
for countsfile in os.listdir(rootdir):
    if not countsfile.endswith(".csv"):
        continue
    counts = pandas.read_csv(os.path.join(rootdir, countsfile), index_col=0)
    print("  reading", countsfile)
    if genes is None:
        genes = counts.index.values
    assert numpy.all(genes == counts.index.values)
    data.append(counts)

data = pandas.concat(data, axis=1)
artificial = data.index.str.startswith("ERCC-") | data.index.str.endswith("_transgene")
data = data.loc[~artificial,cellnames].astype("int32")
loompy.create(outputfile, data.values, {"Gene": data.index.values}, {"CellID": data.columns.values})
```
