## Supplementary material for "CellFishing.jl: an ultrafast and scalable cell search method for single-cell RNA-sequencing": transformation.jl

```
using CellFishing

include("utils.jl")

function experiment(X::Matrix{Float32}, t)
    CellFishing.transform!(t, copy(X))  # JIT compile
    X = copy(X)
    time = @elapsed CellFishing.transform!(t, X)
    println("[[experiment]]")
    println("size = [", size(X, 1), ", ", size(X, 2), "]")
    println("transformation = '", t, "'")
    println("elapsed-time = ", time )
    println()
    flush(stdout)
end

@experiments begin
    m = 3000
    n = 10000
    Random.seed!(RANDSEED)
    X = rand(Float32, m, n) .* 10
    for t in [CellFishing.LogT(1.0f0), CellFishing.FTT()]
        for _ in 1:100; experiment(X, t); end
    end
end
```
