## Supplementary material for "CellFishing.jl: an ultrafast and scalable cell search method for single-cell RNA-sequencing": utils.jl

```
using Dates
using DelimitedFiles
using InteractiveUtils
using Random
using SparseArrays
using Statistics
using LinearAlgebra
using HDF5

function loadloom(datafile, features=nothing)
    HDF5.h5open(datafile) do file
        counts = Matrix(read(file, "/matrix")')
        featurenames = read(file, "/row_attrs/Gene")
        cellnames = read(file, "/col_attrs/CellID")
        if features != nothing
            rows = [findfirst(isequal(x), featurenames) for x in features]
            counts = counts[rows,:]
            featurenames = featurenames[rows]
        end
        return (counts=counts, featurenames=featurenames, cellnames=cellnames)
    end
end

function loadchromium(datafile, group)
    HDF5.h5open(datafile) do file
        data = file[group]
        featurenames = read(data["genes"])
        cellnames = read(data["barcodes"])
        m, n = read(data["shape"])
        counts = SparseMatrixCSC(
            m, n,
            read(data["indptr"]).+1,
            read(data["indices"]).+1,
            convert(Vector{Int32}, read(data["data"])))
        SparseArrays.sortSparseMatrixCSC!(counts)
        return (counts=counts, featurenames=featurenames, cellnames=cellnames)
    end
end

function make_random_blocks(n, k)
    cells = shuffle(1:n)
    blocks = Vector{Int}[]
    q, r = divrem(n, k)
    i = 0
    while i < n
        sz = q
        if r > 0
            sz += 1
            r -= 1
        end
        push!(blocks, cells[i+1:i+sz])
        i += sz
    end
    @assert length(blocks) == k
    return blocks
end

writeknn(filename, cellnames, neighbors) = writeknn(filename, cellnames, cellnames, neighbors)
function writeknn(filename, cellnames_query, cellnames_reference, neighbors)
    mkpath(dirname(filename))
    k, n = size(neighbors)
    open(filename, "w") do file
        # header
        println(file, join(["cell"; string.("N", 1:k)], '\t'))
        # body
        for j in 1:n
            print(file, cellnames_query[j])
            for i in 1:k
                print(file, '\t', cellnames_reference[neighbors[i,j]])
            end
            println(file)
        end
    end
end

const RANDSEED = 1234

macro experiments(blk)
    quote
        starttime = now()
        println("program-file = '", PROGRAM_FILE, "'")
        println("date-time = ", starttime)
        println("version-info = '''\n", sprint(versioninfo), "'''")
        println()
        $(esc(blk))
        println("# END (", Dates.canonicalize(Dates.CompoundPeriod(now() - starttime)), ")")
    end
end
```
