## Supplementary material for "CellFishing.jl: an ultrafast and scalable cell search method for single-cell RNA-sequencing": Snakefile

```
DATASETS = [
    "Baron2016-human",
    "Shekhar2016",
    "Plass2018",
    "TabulaMuris-chromium",
]

Shekhar2016_BATCHES = ["1", "2"]

Plass2018_BATCHES = [
    "plan1",
    "plan2",
    "ds2501rerun",
    "ds2502rerun",
    "gfp",
    "h2b",
    "p7mayrerun",
    "x1mayrerun",
    "cutd0",
    "cutd2",
    "cutd3",
]


# Experiments
# -----------

rule all:
    input:
        [[f"results/{dataset}.estimator.toml",
          f"results/{dataset}.svd.toml",
          f"results/{dataset}.knn-cv.toml",
          f"results/{dataset}.knn-cv.scmap.toml",
          f"results/{dataset}.clustering-sc3.toml",
          f"results/{dataset}.save-load.toml",] for dataset in DATASETS],
        [f"results/{dataset}.select-features.n_features-default.txt" for dataset in DATASETS],
        [[f"results/{dataset}.select-features.n_features-{n_features}.txt",
          f"results/{dataset}.select-features.scmap.n_features-{n_features}.txt",] for dataset in DATASETS for n_features in [500, 1000, 2000, 4000]],
        [f"results/{dataset}.knn-cv-features.n_features-{n_features}.toml" for dataset in DATASETS for n_features in [500, 1000, 2000, 4000]],
        [f"results/{dataset}.knn-cv-features.scmap.n_features-{n_features}.toml" for dataset in DATASETS for n_features in ["default", 500, 1000, 2000, 4000]],
        [[f"results/Shekhar2016.hash.batch-{batch}.toml",
          f"results/Shekhar2016.knn-batch.batch-{batch}.toml",
          f"results/Shekhar2016.knn-batch.scmap.batch-{batch}.toml",] for batch in ["1", "2"]],
        [f"results/Shekhar2016.knn-similarities.target-{target}.toml"
         # semicolon (;) is a replacement of slash (/) here
         for target in ["BC2", "BC5D", "BC3A", "BC5B", "BC4", "BC8;9 (mixture of BC8 and BC9)", "AC (Amacrine cell)"]],
        [[f"results/Plass2018.knn-batch.batch-{batch}.toml",
          f"results/Plass2018.knn-batch.scmap.batch-{batch}.toml",] for batch in ["plan1", "plan2"]],
        [f"results/TabulaMuris-chromium.knn-similarities.target-{target}.toml"
         for target in ["basal cell", "alveolar macrophage", "fibroblast", "immature B cell", "basophil", "kidney cell", "endothelial cell of hepatic sinusoid"]],
        "results/TabulaMuris-chromium.knn-degs.target-immature B cell.toml",
        "results/Baron2016.knn-species.toml",
        "results/Baron2016.knn-species.scmap.toml",
        "results/TabulaMuris.knn-protocol.toml",
        "results/TabulaMuris.knn-protocol.scmap.toml",
        "results/1M_neurons.scalability.toml",
        "results/transformation.toml",
    output:
        "results/Analysis.html"
    log:
        "results/Analysis.log"
    shell:
        "pipenv run jupyter nbconvert --execute --ExecutePreprocessor.timeout=-1 --stdout Analysis.ipynb >{output} 2>{log}"

rule:
    output:
        "results/{dataset}.svd.toml"
    log:
        "results/{dataset}.svd.log"
    input:
        "data/{dataset}.dge.loom"
    shell:
        "julia scripts/svd.jl {input} >{output} 2>{log}"

rule:
    output:
        "results/{dataset}.estimator.toml"
    log:
        "results/{dataset}.estimator.log"
    input:
        "data/{dataset}.dge.loom",
        "data/{dataset}.random-100.txt"
    shell:
        "julia scripts/estimator.jl results/{wildcards.dataset}.estimator/ {input} >{output} 2>{log}"

rule:
    output:
        "results/transformation.toml"
    log:
        "results/transformation.log"
    shell:
        "julia scripts/transformation.jl >{output} 2>{log}"

rule:
    output:
        "results/Shekhar2016.hash.batch-{batch}.toml"
    log:
        "results/Shekhar2016.hash.batch-{batch}.log"
    input:
        "data/Shekhar2016.dge.loom",
        "data/Shekhar2016.batch-{batch}.txt",
        lambda wildcards: [f"data/Shekhar2016.batch-{batch}.txt" for batch in Shekhar2016_BATCHES if batch != wildcards.batch]
    shell:
        "julia scripts/hash.jl results/Shekhar2016.hash.batch-{wildcards.batch}/ {input} >{output} 2>{log}"

rule:
    output:
        "results/{dataset}.knn-cv.toml"
    log:
        "results/{dataset}.knn-cv.log"
    input:
        "data/{dataset}.dge.loom"
    shell:
        "julia scripts/knn-cv.jl results/{wildcards.dataset}.knn-cv/ {input} >{output} 2>{log}"

rule:
    output:
        "results/{dataset}.knn-cv.scmap.toml"
    log:
        "results/{dataset}.knn-cv.scmap.log"
    input:
        "data/{dataset}.dge.loom"
    shell:
        r"""
        /usr/bin/docker run --rm -u $(id -u):$(id -g) -v $(pwd):/workspace ksato/scmap Rscript --vanilla scripts/knn-cv.scmap.R results/{wildcards.dataset}.knn-cv.scmap/ {input} >{output} 2>{log}
        """

rule:
    output:
        "results/{dataset}.clustering-sc3.toml"
    log:
        "results/{dataset}.clustering-sc3.log"
    input:
        "data/{dataset}.dge.loom",
        "data/{dataset}.cluster.tsv"
    shell:
        r"""
        /usr/bin/docker run --rm -u $(id -u):$(id -g) -v $(pwd):/workspace ksato/scmap Rscript --vanilla scripts/clustering-sc3.R results/{wildcards.dataset}.clustering-sc3/ {input} >{output} 2>{log}
        """
rule:
    output:
        "results/{dataset}.knn-similarities.target-{target}.toml"
    log:
        "results/{dataset}.knn-similarities.target-{target}.log"
    input:
        "data/{dataset}.dge.loom",
        "data/{dataset}.cluster.tsv"
    shell:
        """
        julia scripts/knn-similarity.jl 'results/{wildcards.dataset}.knn-similarities.target-{wildcards.target}/' {input} '{wildcards.target}' >'{output}' 2>'{log}'
        """

rule:
    output:
        "results/{dataset}.knn-degs.target-{target}.toml"
    log:
        "results/{dataset}.knn-degs.target-{target}.log"
    input:
        "data/{dataset}.dge.loom",
        "data/{dataset}.cluster.tsv"
    shell:
        """
        julia scripts/knn-degs.jl 'results/{wildcards.dataset}.knn-degs.target-{wildcards.target}/' {input} '{wildcards.target}' >'{output}' 2>'{log}'
        """

rule:
    output:
        "results/{dataset}.save-load.toml"
    log:
        "results/{dataset}.save-load.log"
    input:
        "data/{dataset}.dge.loom"
    shell:
        "julia scripts/save-load.jl {input} >{output} 2>{log}"

rule:
    output:
        "results/{dataset}.select-features.n_features-default.txt"
    log:
        "results/{dataset}.select-features.n_features-default.log"
    input:
        "data/{dataset}.dge.loom"
    shell:
        "julia scripts/select-features.jl {input} 0 >{output} 2>{log}"

rule:
    output:
        "results/{dataset}.select-features.n_features-{n_features}.txt"
    log:
        "results/{dataset}.select-features.n_features-{n_features}.log"
    input:
        "data/{dataset}.dge.loom"
    wildcard_constraints:
        n_features="\d+"
    shell:
        "julia scripts/select-features.jl {input} {wildcards.n_features} >{output} 2>{log}"

rule:
    output:
        "results/{dataset}.select-features.scmap.n_features-{n_features}.txt"
    log:
        "results/{dataset}.select-features.scmap.n_features-{n_features}.log"
    input:
        "data/{dataset}.dge.loom"
    shell:
        r"""
        /usr/bin/docker run --rm -u $(id -u):$(id -g) -v $(pwd):/workspace ksato/scmap Rscript --vanilla scripts/select-features.scmap.R {input} {wildcards.n_features} >{output} 2>{log}
        """

rule:
    output:
        "results/{dataset}.knn-cv-features.n_features-{n_features}.toml"
    log:
        "results/{dataset}.knn-cv-features.n_features-{n_features}.log"
    input:
        "data/{dataset}.dge.loom",
        "results/{dataset}.select-features.scmap.n_features-{n_features}.txt",
    shell:
        "julia scripts/knn-cv-features.jl results/{wildcards.dataset}.knn-cv-features.n_features-{wildcards.n_features}/ {input} >{output} 2>{log}"

rule:
    output:
        "results/{dataset}.knn-cv-features.scmap.n_features-{n_features}.toml"
    log:
        "results/{dataset}.knn-cv-features.scmap.n_features-{n_features}.log"
    input:
        "data/{dataset}.dge.loom",
        "results/{dataset}.select-features.n_features-{n_features}.txt",
    shell:
        r"""
        /usr/bin/docker run --rm -u $(id -u):$(id -g) -v $(pwd):/workspace ksato/scmap Rscript --vanilla scripts/knn-cv-features.scmap.R results/{wildcards.dataset}.knn-cv-features.scmap.n_features-{wildcards.n_features}/ {input} >{output} 2>{log}
        """

rule:
    output:
        "results/Shekhar2016.knn-batch.batch-{batch}.toml"
    log:
        "results/Shekhar2016.knn-batch.batch-{batch}.log"
    input:
        "data/Shekhar2016.dge.loom",
        "data/Shekhar2016.batch-{batch}.txt",
        lambda wildcards: [f"data/Shekhar2016.batch-{batch}.txt" for batch in Shekhar2016_BATCHES if batch != wildcards.batch]
    shell:
        "julia scripts/knn-batch.jl results/Shekhar2016.knn-batch.batch-{wildcards.batch}/ {input} >{output} 2>{log}"

rule:
    output:
        "results/Plass2018.knn-batch.batch-{batch}.toml"
    log:
        "results/Plass2018.knn-batch.batch-{batch}.log"
    input:
        "data/Plass2018.dge.loom",
        "data/Plass2018.batch-{batch}.txt",
        lambda wildcards: [f"data/Plass2018.batch-{batch}.txt" for batch in Plass2018_BATCHES if batch != wildcards.batch]
    shell:
        "julia scripts/knn-batch.jl results/Plass2018.knn-batch.batch-{wildcards.batch}/ {input} >{output} 2>{log}"

rule:
    output:
        "results/Shekhar2016.knn-batch.scmap.batch-{batch}.toml"
    log:
        "results/Shekhar2016.knn-batch.scmap.batch-{batch}.log"
    input:
        "data/Shekhar2016.dge.loom",
        "data/Shekhar2016.batch-{batch}.txt",
        lambda wildcards: [f"data/Shekhar2016.batch-{batch}.txt" for batch in Shekhar2016_BATCHES if batch != wildcards.batch]
    shell:
        r"""
        /usr/bin/docker run --rm -u $(id -u):$(id -g) -v $(pwd):/workspace ksato/scmap Rscript --vanilla scripts/knn-batch.scmap.R results/Shekhar2016.knn-batch.scmap.batch-{wildcards.batch}/ {input} >{output} 2>{log}
        """

rule:
    output:
        "results/Plass2018.knn-batch.scmap.batch-{batch}.toml"
    log:
        "results/Plass2018.knn-batch.scmap.batch-{batch}.log"
    input:
        "data/Plass2018.dge.loom",
        "data/Plass2018.batch-{batch}.txt",
        lambda wildcards: [f"data/Plass2018.batch-{batch}.txt" for batch in Plass2018_BATCHES if batch != wildcards.batch]
    shell:
        r"""
        /usr/bin/docker run --rm -u $(id -u):$(id -g) -v $(pwd):/workspace ksato/scmap Rscript --vanilla scripts/knn-batch.scmap.R results/Plass2018.knn-batch.scmap.batch-{wildcards.batch}/ {input} >{output} 2>{log}
        """

rule:
    output:
        "results/Baron2016.knn-species.toml"
    log:
        "results/Baron2016.knn-species.log"
    input:
        "data/Baron2016.mouse_human_homgenes.tsv",
        "data/Baron2016-mouse.dge.loom",
        "data/Baron2016-human.dge.loom"
    shell:
        "julia scripts/knn-species.jl results/Baron2016.knn-species/ {input} >{output} 2>{log}"

rule:
    output:
        "results/Baron2016.knn-species.scmap.toml"
    log:
        "results/Baron2016.knn-species.scmap.log"
    input:
        "data/Baron2016.mouse_human_homgenes.tsv",
        "data/Baron2016-mouse.dge.loom",
        "data/Baron2016-human.dge.loom"
    shell:
        r"""
        /usr/bin/docker run --rm -u $(id -u):$(id -g) -v $(pwd):/workspace ksato/scmap Rscript --vanilla scripts/knn-species.scmap.R results/Baron2016.knn-species.scmap/ {input} >{output} 2>{log}
        """

rule:
    output:
        "results/TabulaMuris.knn-protocol.toml"
    log:
        "results/TabulaMuris.knn-protocol.log"
    input:
        "data/TabulaMuris-smart.dge.loom",
        "data/TabulaMuris-chromium.dge.loom"
    shell:
        "julia scripts/knn-protocol.jl results/TabulaMuris.knn-protocol/ {input} >{output} 2>{log}"

rule:
    output:
        "results/TabulaMuris.knn-protocol.scmap.toml"
    log:
        "results/TabulaMuris.knn-protocol.scmap.log"
    input:
        "data/TabulaMuris-smart.dge.loom",
        "data/TabulaMuris-chromium.dge.loom"
    shell:
        r"""
        /usr/bin/docker run --rm -u $(id -u):$(id -g) -v $(pwd):/workspace ksato/scmap Rscript --vanilla scripts/knn-protocol.scmap.R results/TabulaMuris.knn-protocol.scmap/ {input} >{output} 2>{log}
        """

rule:
    output:
        "results/1M_neurons.scalability.toml"
    log:
        "results/1M_neurons.scalability.log"
    input:
        "data/raw/1M_neurons/1M_neurons.h5",
        "data/1M_neurons.random-10000.txt"
    shell:
        "julia scripts/scalability.jl results/1M_neurons.scalability/ {input} >{output} 2>{log}"


# Data
# ----

rule data:
    input:
        "data/Baron2016-human.dge.loom",
        "data/Baron2016-human.cluster.tsv",
        "data/Baron2016-mouse.dge.loom",
        "data/Baron2016-mouse.cluster.tsv",
        "data/Shekhar2016.dge.loom",
        "data/Shekhar2016.cluster.tsv",
        "data/Plass2018.dge.loom",
        "data/Plass2018.cluster.tsv",
        "data/TabulaMuris-chromium.dge.loom",
        "data/TabulaMuris-chromium.cluster.tsv",
        "data/TabulaMuris-smart.dge.loom",
        "data/TabulaMuris-smart.cluster.tsv",

rule:
    output:
        "data/{dataset}.random-100.txt"
    input:
        "data/{dataset}.cluster.tsv"
    shell:
        r"""cat {input} | perl -F'\t' -anle 'print $F[0] if $. > 1' | perl -MList::Util=shuffle -e 'srand(1234); print((shuffle(<>))[0..99])' >{output}"""

rule:
    output:
        "data/{dataset}.featurenames.txt"
    input:
        "data/{dataset}.dge.loom"
    shell:
        "python scripts/featurenames.py {input} >{output}"


# Baron2016
# ---------

Baron2016_HUMAN = [
    "GSM2230757_human1_umifm_counts.csv.gz",
    "GSM2230758_human2_umifm_counts.csv.gz",
    "GSM2230759_human3_umifm_counts.csv.gz",
    "GSM2230760_human4_umifm_counts.csv.gz",
]
Baron2016_MOUSE = [
    "GSM2230761_mouse1_umifm_counts.csv.gz",
    "GSM2230762_mouse2_umifm_counts.csv.gz",
]

rule:
    output:
        "data/Baron2016-human.dge.loom"
    input:
        "data/Baron2016-human.cluster.tsv",
        [f"data/tmp/Baron2016/{name}" for name in Baron2016_HUMAN]
    shell:
        """
        python scripts/Baron2016.loomify.py {input} {output}
        """

rule:
    output:
        "data/Baron2016-mouse.dge.loom"
    input:
        "data/Baron2016-mouse.cluster.tsv",
        [f"data/tmp/Baron2016/{name}" for name in Baron2016_MOUSE]
    shell:
        """
        python scripts/Baron2016.loomify.py {input} {output}
        """

rule:
    output:
        "data/Baron2016-human.cluster.tsv"
    input:
        [f"data/tmp/Baron2016/{name}" for name in Baron2016_HUMAN]
    shell:
        r"""
        cat {input} | gzip -cd | perl -F, -anle 'BEGIN {{print "cell\tcluster"}} print "$F[0]\t$F[2]" if $F[0] ne ""' >{output}
        """

rule:
    output:
        "data/Baron2016-mouse.cluster.tsv"
    input:
        [f"data/tmp/Baron2016/{name}" for name in Baron2016_MOUSE]
    shell:
        r"""
        cat {input} | gzip -cd | perl -F, -anle 'BEGIN {{print "cell\tcluster"}} print "$F[0]\t$F[2]" if $F[0] ne ""' >{output}
        """

rule:
    output:
        [f"data/tmp/Baron2016/{name}" for name in Baron2016_HUMAN + Baron2016_MOUSE]
    input:
        "data/raw/Baron2016/GSE84133_RAW.tar"
    shell:
        """
        tar xf {input} -C data/tmp/Baron2016/
        touch {output}
        """

rule:
    output:
        "data/raw/Baron2016/GSE84133_RAW.tar"
    shell:
        "curl -o {output} 'https://www.ncbi.nlm.nih.gov/geo/download/?acc=GSE84133&format=file'"

rule:
    output:
        "data/Baron2016.mouse_human_homgenes.tsv"
    input:
        "data/raw/Baron2016/HOM_MouseHumanSequence.tsv",
        "data/Baron2016-mouse.featurenames.txt",
        "data/Baron2016-human.featurenames.txt"
    shell:
        r"""
        export LC_ALL=C
        sort -c {input[1]}
        TMP_MOUSE=$(mktemp)
        cat {input[0]} | perl -F'\t' -anle 'print "$F[3]\t$F[0]" if $F[2] eq "10090"' | sort -k1 | join -t $'\t' {input[1]} - | perl -F'\t' -anle 'print "$F[1]\t$F[0]"' | sort -k1 >$TMP_MOUSE
        sort -c {input[2]}
        TMP_HUMAN=$(mktemp)
        cat {input[0]} | perl -F'\t' -anle 'print "$F[3]\t$F[0]" if $F[2] eq "9606"'  | sort -k1 | join -t $'\t' {input[2]} - | perl -F'\t' -anle 'print "$F[1]\t$F[0]"' | sort -k1 >$TMP_HUMAN
        TMP_UNIQUE=$(mktemp)
        join -t $'\t' $TMP_MOUSE $TMP_HUMAN | cut -f1 | uniq -u >$TMP_UNIQUE
        join -t $'\t' $TMP_UNIQUE $TMP_MOUSE | join -t $'\t' - $TMP_HUMAN | cut -f2-3 >{output}
        rm -f $TMP_MOUSE $TMP_HUMAN $TMP_UNIQUE
        """

rule:
    output:
        "data/raw/Baron2016/HOM_MouseHumanSequence.tsv"
    shell:
        "curl -o {output} http://www.informatics.jax.org/downloads/reports/HOM_MouseHumanSequence.rpt"


# Shekhar2016
# -----------

rule:
    output:
        "data/Shekhar2016.dge.loom"
    input:
        "data/Shekhar2016.cluster.tsv",
        "data/raw/Shekhar2016/GSE81904_BipolarUMICounts_Cell2016.txt.gz"
    shell:
        "python scripts/Shekhar2016.loomify.py {input} {output}"

rule:
    output:
        "data/Shekhar2016.cluster.tsv"
    input:
        "data/raw/Shekhar2016/clust_retinal_bipolar.txt"
    shell:
        r"""cat {input} | perl -F'\t' -anle 'BEGIN {{print "cell\tcluster"}} print "$F[0]\t$F[1]" if $. > 2' >{output}"""

rule:
    output:
        "data/Shekhar2016.batch-1.txt"
    input:
        "data/Shekhar2016.cluster.tsv"
    shell:
        r"""cat {input} | perl -F'\t' -anle 'print $F[0] if $F[0] =~ /^Bipolar[1234]_/' >{output}"""

rule:
    output:
        "data/Shekhar2016.batch-2.txt"
    input:
        "data/Shekhar2016.cluster.tsv"
    shell:
        r"""cat {input} | perl -F'\t' -anle 'print $F[0] if $F[0] =~ /^Bipolar[56]_/' >{output}"""

rule:
    output:
        "data/raw/Shekhar2016/GSE81904_BipolarUMICounts_Cell2016.txt.gz"
    shell:
        "curl -o {output} 'https://www.ncbi.nlm.nih.gov/geo/download/?acc=GSE81904&format=file&file=GSE81904%5FBipolarUMICounts%5FCell2016%2Etxt%2Egz'"


# Plass2018
# ---------

rule:
    output:
        "data/Plass2018.dge.loom"
    input:
        "data/Plass2018.cluster.tsv",
        "data/raw/Plass2018/dge.txt.gz"
    shell:
        "python scripts/Plass2018.loomify.py {input} {output}"

rule:
    output:
        "data/Plass2018.cluster.tsv"
    input:
        "data/raw/Plass2018/dge.txt.gz",
        "data/raw/Plass2018/R_annotation.txt"
    shell:
        r"""
        set +o pipefail  # SIGPIPE
        gzip -cd <{input[0]} | head -1 | perl -ple 's/\t/\n/g' | (perl -le 'print "cell\tcluster"'; paste - {input[1]}) >{output}
        """

rule:
    output:
        "data/Plass2018.batch-{name}.txt"
    input:
        "data/Plass2018.cluster.tsv"
    shell:
        r"""cat {input} | perl -F'\t' -anle 'print $F[0] if $F[0] =~ /^{wildcards.name}_/' >{output}"""

rule:
    output:
        "data/raw/Plass2018/dge.txt.gz"
    shell:
        "curl -o {output} 'http://bimsbstatic.mdc-berlin.de/rajewsky/PSCA/dge.txt.gz'"

rule:
    output:
        "data/raw/Plass2018/R_annotation.txt"
    shell:
        "curl -o {output} 'http://bimsbstatic.mdc-berlin.de/rajewsky/PSCA/R_annotation.txt'"


# TabulaMuris
# -----------

rule:
    output:
        "data/TabulaMuris-chromium.dge.loom"
    input:
        "data/TabulaMuris-chromium.cluster.tsv",
        "data/tmp/TabulaMuris/chromium/droplet.zip"
    shell:
        r"""
        unzip -od data/tmp/TabulaMuris/chromium/ {input[1]}
        python scripts/TabulaMuris-chromium.loomify.py {input[0]} data/tmp/TabulaMuris/chromium/droplet/ {output}
        """

rule:
    output:
        "data/TabulaMuris-smart.dge.loom"
    input:
        "data/TabulaMuris-smart.cluster.tsv",
        "data/tmp/TabulaMuris/smart/FACS.zip"
    shell:
        r"""
        unzip -od data/tmp/TabulaMuris/smart/ {input[1]}
        python scripts/TabulaMuris-smart.loomify.py {input[0]} data/tmp/TabulaMuris/smart/FACS/ {output}
        """

rule:
    output:
        "data/TabulaMuris-chromium.cluster.tsv"
    input:
        "data/tmp/TabulaMuris/chromium/annotations_droplet.csv"
    shell:
        r"""cat {input} | sed 1d | perl -F, -anle 'BEGIN {{print "cell\tcluster"}} print "$F[0]\t$F[1]" if $F[1] ne ""' >{output}"""

rule:
    output:
        "data/TabulaMuris-smart.cluster.tsv"
    input:
        "data/tmp/TabulaMuris/smart/annotations_FACS.csv"
    shell:
        r"""cat {input} | sed 1d | perl -F, -anle 'BEGIN {{print "cell\tcluster"}} print "$F[2]\t$F[3]" if $F[3] ne ""' >{output}"""

rule:
    output:
        "data/tmp/TabulaMuris/chromium/annotations_droplet.csv",
        "data/tmp/TabulaMuris/chromium/droplet.zip"
    input:
        "data/raw/TabulaMuris/chromium/5968960.zip"
    shell:
        """
        unzip -od data/tmp/TabulaMuris/chromium/ {input}
        touch {output}
        """

rule:
    output:
        "data/tmp/TabulaMuris/smart/annotations_FACS.csv",
        "data/tmp/TabulaMuris/smart/FACS.zip"
    input:
        "data/raw/TabulaMuris/smart/5829687.zip"
    shell:
        """
        unzip -od data/tmp/TabulaMuris/smart/ {input}
        touch {output}
        """

rule:
    output:
        "data/raw/TabulaMuris/chromium/5968960.zip"
    shell:
        "curl -o {output} 'https://ndownloader.figshare.com/articles/5968960/versions/2'"

rule:
    output:
        "data/raw/TabulaMuris/smart/5829687.zip"
    shell:
        "curl -o {output} 'https://ndownloader.figshare.com/articles/5829687/versions/7'"


# 1M_neurons
# ----------

rule:
    output:
        "data/1M_neurons.random-10000.txt"
    input:
        "data/1M_neurons.cluster.tsv"
    shell:
        r"""cat {input} | perl -F'\t' -anle 'print $F[0] if $. > 1' | perl -MList::Util=shuffle -e 'srand(1234); print((shuffle(<>))[0..9999])' >{output}"""

rule:
    output:
        "data/1M_neurons.cluster.tsv"
    input:
        "data/raw/1M_neurons/1M_neurons_analysis.tar.gz"
    shell:
        r"""tar -xOf {input} analysis/clustering/graphclust/clusters.csv | perl -F, -anle 'BEGIN {{print "cell\tcluster"}} print "$F[0]\t$F[1]" if $. > 1' >{output}"""

rule:
    output:
        "data/raw/1M_neurons/1M_neurons.h5"
    shell:
        "curl -o {output} 'http://cf.10xgenomics.com/samples/cell-exp/1.3.0/1M_neurons/1M_neurons_filtered_gene_bc_matrices_h5.h5'"

rule:
    output:
        "data/raw/1M_neurons/1M_neurons_analysis.tar.gz"
    shell:
        "curl -o {output} 'http://cf.10xgenomics.com/samples/cell-exp/1.3.0/1M_neurons/1M_neurons_analysis.tar.gz'"
```
