## Supplementary figures and images for "CellFishing.jl: an ultrafast and scalable cell search method for single-cell RNA-sequencing"

### schematic.pdf

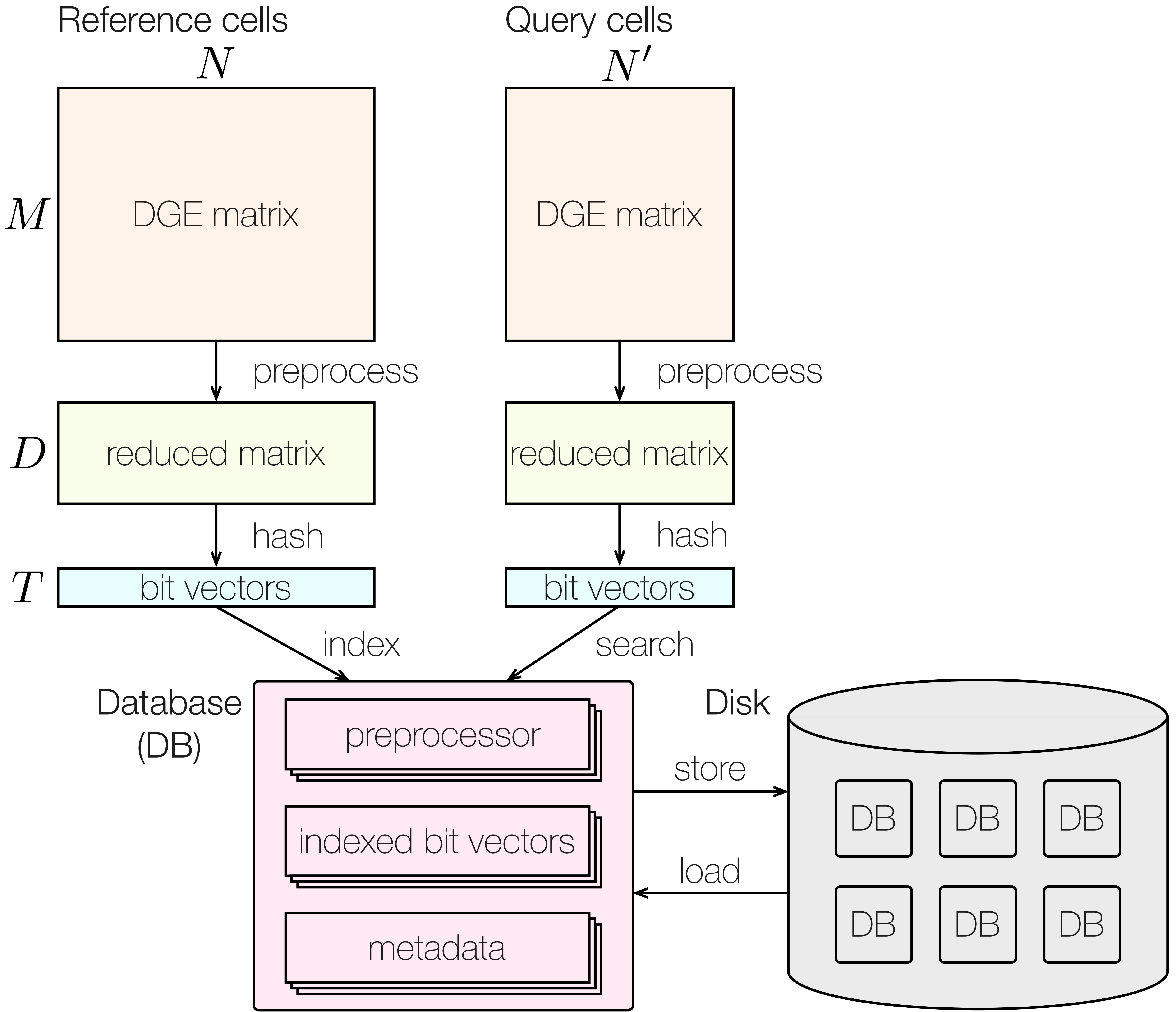

### subindex.pdf

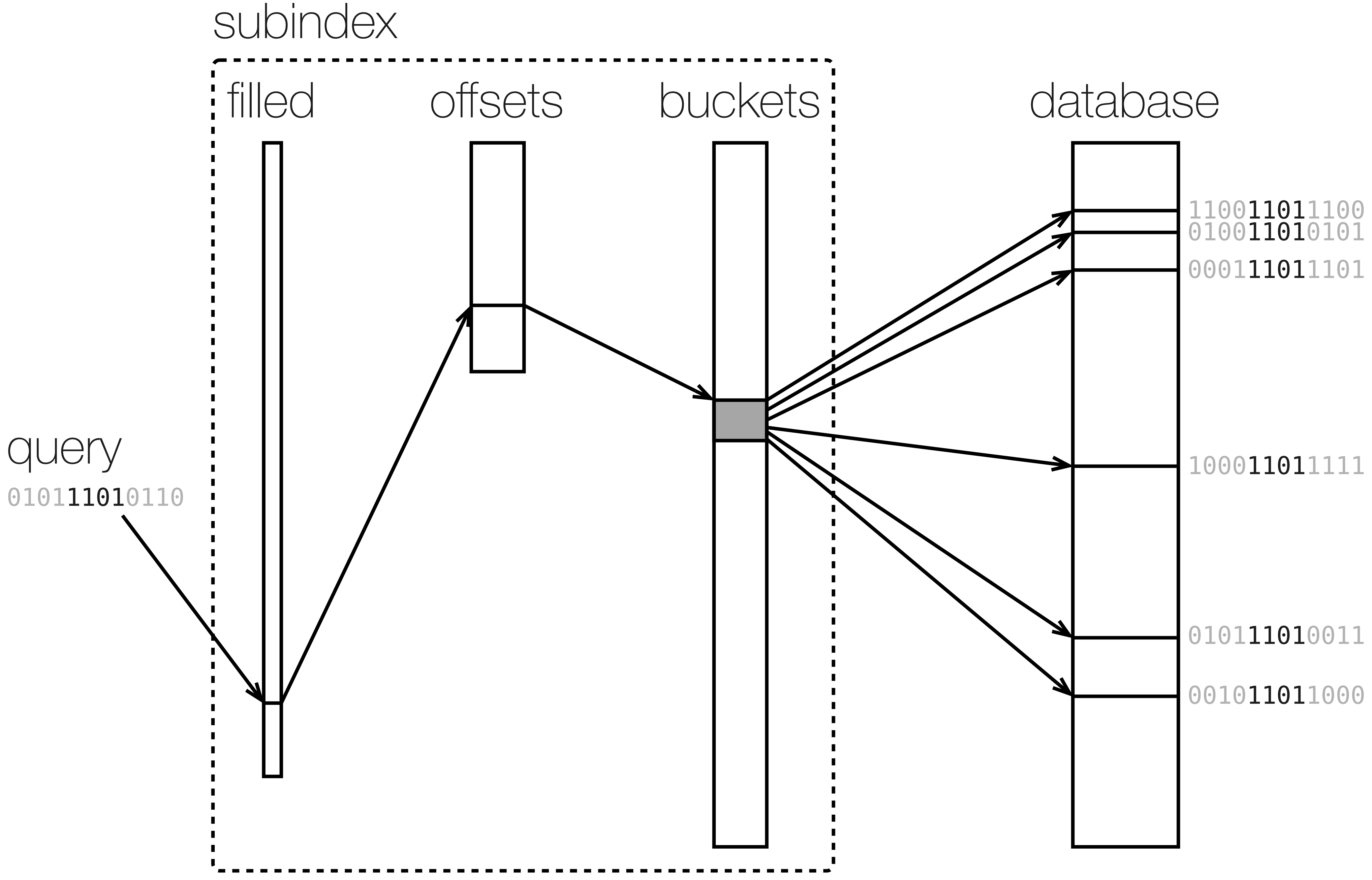
